# The global and promoter-centric 3D genome organization temporally resolved during a circadian cycle

**DOI:** 10.1101/2020.07.23.217992

**Authors:** Masami Ando-Kuri, Rodrigo G. Arzate-Mejía, Jorg Morf, Jonathan Cairns, Cesar A. Poot-Hernández, Simon Andrews, Csilla Várnai, Boo Virk, Steven W. Wingett, Peter Fraser, Mayra Furlan-Magaril

## Abstract

Circadian gene expression is essential for organisms to adjust cellular responses and anticipate daily changes in the environment. In addition to its physiological importance, the clock circuit represents an ideal, temporally resolved, system to study transcription regulation. Here, we analysed changes in spatial mouse liver chromatin conformation using genome-wide and promoter-capture Hi-C alongside daily oscillations in gene transcription in mouse liver. We found circadian topologically associated domains switched assignments to the transcriptionally active, open chromatin compartment and the inactive compartment at different hours of the day while their boundaries stably maintain their structure over time. Individual circadian gene promoters displayed maximal chromatin contacts at times of peak transcriptional output and the expression of circadian genes and contacted transcribed regulatory elements, or other circadian genes, was phase-coherent. Anchor sites of promoter chromatin loops were enriched in binding sites for liver nuclear receptors and transcription factors, some exclusively present in either rhythmic or stable contacts. The circadian 3D chromatin maps provided here identify the scales of chromatin conformation that parallel oscillatory gene expression and protein factors specifically associated with circadian or stable chromatin configurations.

## Introduction

Circadian variation of gene expression in the liver is essential to temporally coordinate metabolic processes including lipid and glycogen metabolism, and maintain organism homeostasis. Considerable progress has been made in understanding circadian transcription regulation (Takahashi, 2017; Zhang, et al., 2014), however, the impact of 3D chromatin configuration dynamics over the course of a day in circadian oscillations of gene expression has not been fully characterized.

Previous work on chromatin contacts restricted to individual, candidate genomic loci has provided evidence that the circadian gene *Dbp* forms inter-chromosomal contacts with ∼200kb genome blocks, which fluctuate in strength over the course of a day in cultured cells (Aguilar-Arnal et al., 2013). At higher resolution enhancer-promoter contacts for core-clock gene *Cry1* and clock output genes *Mreg, Slc45a3, Gys2* have been shown to oscillate in a daily and *Arntl*-dependent manner in liver (Mermet et al., 2018; Yeung & Naef, 2018). In contrast, analysis of *Arntl* cistrome showed quite stable contacts between *Arntl* occupied regulatory elements (Beytebiere et al., 2018). Also *Nr1d1* circadian gene forms invariant contacts to a nearby super-enhancer with the help of Cohesin throughout the circadian cycle (Xu et al., 2016). Finally, genome-wide Hi-C studies at two time points of a day-night cycle suggested that circadian target genes of the Nr1d1 repressor protein form contacts within their respective Topological Associating Domains (TADs), that can be dynamic over time (Y. H. Kim et al., 2018b).

Even if there is evidence of dynamic and stable contacts from candidate circadian genes, we still lack an understanding of what factors distinguish rhythmic and constant genomic contacts formed by circadian genes with maxima of transcriptional output (acrophases) at different times, and how common these types of chromatin contacts are when analysing all the circadian gene promoters in the genome. Also, a high-resolution genome-wide promoter centric panorama of all circadian contacts resolved in time is still missing. Here we present data from in-nucleus Hi-C and Promoter Capture Hi-C (P-CHi-C) at 4 time points during a day in mouse adult liver. We provide for the first time a temporally resolved genome-wide contact analysis at different scales, encompassing genomic compartments (A, B compartments), mega to kilo-base domains (TADs) and high-resolution coverage of contacts from all individual gene promoters including circadian gene promoters. We identify instances in which genomic A/B compartment assignment changes in parallel to circadian modulation of histone modifications in chromatin and oscillatory gene expression. Many of the circadian genes with accompanying changes in A/B assignment are found in TADs that remain constant during the day. Exploring gene promoter interactions at restriction fragment resolution through P-CHi-C, we found circadian gene promoters form dynamic and stable contacts, increasing their number of genomic contacts at the acrophase of the corresponding transcriptional units. Furthermore, we found a set of liver nuclear receptors (i.e., Nr5a2) binding motifs enriched at both dynamic and constant circadian contacting regions and some transcription factor (TF) binding motifs unique for dynamic or constant promoter contacts (i.e. Immediate early factors Tcfap2 and Fos:Jun). The contacts formed by diurnal and nocturnal circadian gene promoters were found biased towards enhancers and other circadian promoters with day and night time transcriptional activity, respectively. Finally we found that core clock associated gene promoters engage in more dynamic interactomes than output circadian genes in the liver. Together our results provide evidence that genome conformation dynamics are coupled with circadian transcriptional fluctuations at different genomic scales and constitute a detailed 3D map of the liver gene promoter-interactome over a 24 hours cycle.

## Results

### Circadian A/B Chromatin Compartments switch between open and closed configurations throughout the day

To study global genome architecture during a circadian cycle we performed in-nucleus Hi-C (see methods) on mouse adult liver with three replicates at four different time points during a cycle (ZT0, 6, 12 and 18), with ZT0 and ZT12 being the beginning of the light and dark phase, respectively. A section of individual livers from pool a and b were processed in parallel for RNA-seq. We produced high-quality Hi-C data sets characterized by the elevated percentage of valid pairs (∼80%), low PCR duplicates (less than 2%) and high cis:trans interaction ratios obtained (∼80:20%) (Table S1). In total we obtained ∼2 billion valid Hi-C read pairs from mouse adult liver across a circadian cycle (Table S1).

To detect open, transcriptionally active and closed, silent genomic compartments (A and B compartments, respectively) we performed PCA analysis (Lieberman-Aiden et al., 2009) on Hi-C data at different time points throughout the circadian cycle, at 100kb bin resolution. Changes in chromatin compartments have been associated with changes in transcription and chromatin states during cell differentiation and mouse early development (Bonev et al., 2017; Dixon et al., 2015). As expected PC1 values partitioned the liver genome into chromatin compartments (Figure 1A,B Figure S1 A,B). We then compared the eigenvectors of the different time points and identified changes in the sign of regional PC1 values, indicative of compartment switching between all time point pairs (Figure 1A,B Figure S1C one-way ANOVA p-value <2e-16). These genomic regions, termed Oscillatory Chromatin Compartments (OCCs) spanned 440.4 Mb of the mouse genome (Figure 1A,B individual replicates and merged data, respectively). The rest of the genome retained the same compartment identity during the 24 hour cycle (Figure S1A,B individual replicates and merged data, respectively, S1D). We found OCCs with compartment assignments ZT0=A, ZT6=A, ZT12=B, ZT18=A (AABA) being the most abundant type in the genome covering 194.7Mb (Figure S1E).

**Figure 1.**
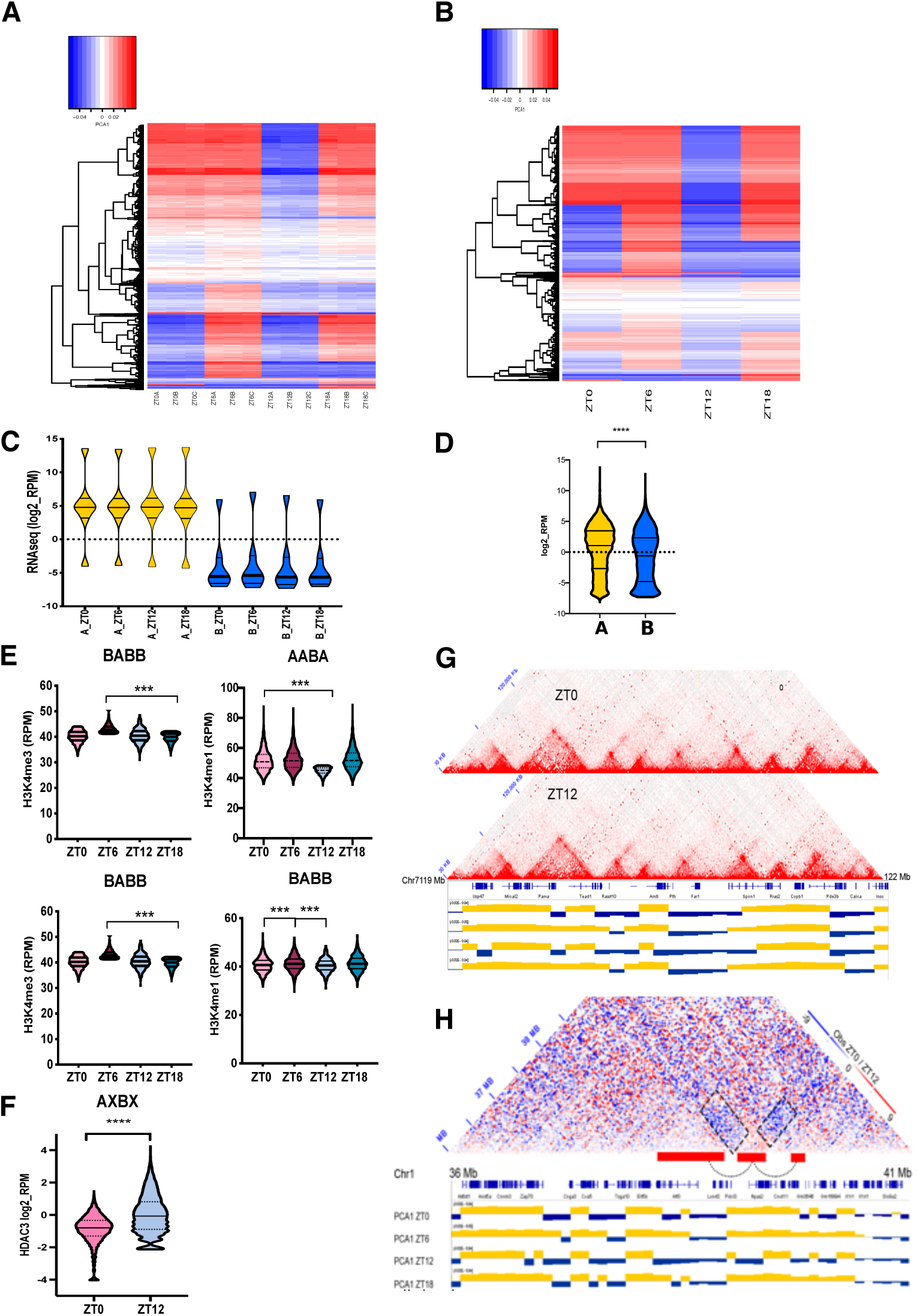
Chromatin compartments change during the 24 hours. A. Heatmap of PC1 values for OCCs (independent replicates, significant variance p-value < 2e-16, one-way ANOVA). B. Heatmap of PC1 values for OCCs (merged replicates, significant variance p-value < 2e-16, one-way ANOVA). C. RNA-seq reads coverage (log2_RPM) in all A vs all B compartments across time points during the circadian cycle (p < 0.005, Kluskal-Wallis test) D. RNA-seq reads coverage (log2_RPM) in OCCs A vs B compartments (p< 0.0001, Mann-Whitney test). E. H3K4me3 and H3K4me1 RPM ChIP-seq signal in OCCs across time points. BABB (ZT0=B, ZT6=A, ZT12=B, ZT18=B) and AABA (ZT0=A, ZT6=A, ZT12=B, ZT18=A) (all p-values < 0.001, one way ANOVA test, Tukey post hoc test). F. HDAC3 ChIPseq log2_RPM signal in OCCs at ZT0 and ZT12 (p-value < 0.0001, Wilcoxon rank test) G. Observed Hi-C contacts from ZT0 and ZT12 liver samples at the *Antl1* cTAD. PC1 values plotted underneath. *Arntl* cTAD switches chromatin compartment at ZT12. H. ZT0/ZT12 differential Hi-C contact matrix of a region including the *Npas2* cTAD switching chromatin compartment at ZT12. PC1 values plotted underneath.

To relate chromatin compartments with transcription we performed stranded ribosomal RNA-depleted total RNA-seq. We obtained ∼ 500,000,000 number of 150 bp reads per timepoint (4 biological replicates each) reaching ∼2,000,000,000 total reads (Table S2). Spearman correlation analysis showed good correlation between the 4 biological replicates (Figure S2G, ZT0 and 12 shown). Genes with differential expression between at least one pair of time points were identified (q-value <0.01) and classified as circadian. In total we detected 1257 circadian gene transcripts (Figure S2A Table S3). Inspection of individual gene expression profiles from our RNA-seq data for known circadian genes (both core clock and liver output genes) showed good agreement with their reported acrophase (Figure S2B,E). Gene Ontology and KEGG pathway analysis identified circadian rhythm and metabolism as significantly enriched categories in our identified circadian gene set (Figure S2C,D) confirming efficient detection of circadian oscillating genes. We also confirmed circadian gene expression of mRNA and primary mRNA of candidate genes by RT-qPCR confirming the expected expression profiles for these genes (Figure S2F). Comparing RNA content in A and B compartments we found a significant difference in RNA abundance between the two types of compartments, with A compartments being RNA-rich and B compartments RNA-poor (Figure 1C, p < 0.005, Kluskal-Wallis test) at all time points. We then examined RNA content in OCCs and also found a significant difference in abundance between regions falling into A or B at different times of the day (Figure 1D, p< 0.0001, Mann-Whitney test). In addition to RNA abundance we analysed time resolved H3K4me3 and H3K4me1 histone modifications enrichment (Koike et al., 2012a) reflecting open chromatin in OCCs around the clock. Both histone marks were significantly enriched in regions at times of A compartment assignment compared to B compartment across all time points (Figure 1E, AABA [ZT0=A, ZT6=A, ZT12=B, ZT18=A] and BABB [ZT0=B, ZT6=A, ZT12=B, ZT18=B], S1F AABB, ABBB, BABA. p<0.001, one-way ANOVA test, Tukey post hoc test). HDAC3 binding reflects deacetylated closed chromatin and it has been measured before in the mouse liver at ZT22 and ZT10 (Sun et al., 2013). We measured HDAC3 enrichment at OCCs at ZT0 and 12 (our closest time points to ZT22 and 10). HDAC3 binding was higher when regions fell into the B compartment at ZT12 compared to the A compartment at ZT0 (Figure 1F, Wilcoxon p<0.001). Together these results show that transcription and chromatin state both fluctuate in accordance with compartment switching during a circadian cycle.

### Topologically Associated Domains spatially partition temporal gene expression control but remain structurally invariant during a circadian cycle

To identify TADs we assigned TAD insulation scores to Hi-C data (Crane et al., 2015) and examined them across time points (see methods). TADs displayed little variation across time points as has been previously observed (Y. H. Kim et al., 2018b). Of a total of 4358 TADs, 2936 were preserved throughout the day and 952 at least between two time points (Figure 2A and Figure S3A,B). To confirm insulation of TADs across time points we selected a random set of 1000 TADs detected at ZT0 and plotted their median Observed/Expected contacts using Hi-C data from ZT0 and ZT12 (see methods). We recovered higher contact frequencies within TADs compared to outside of domains at both time points confirming large domain structure preservation (Figure 2B left panel). The same result held true for TADs detected at ZT6, 12 and 18 hours (Figure S3C).

**Figure 2.**
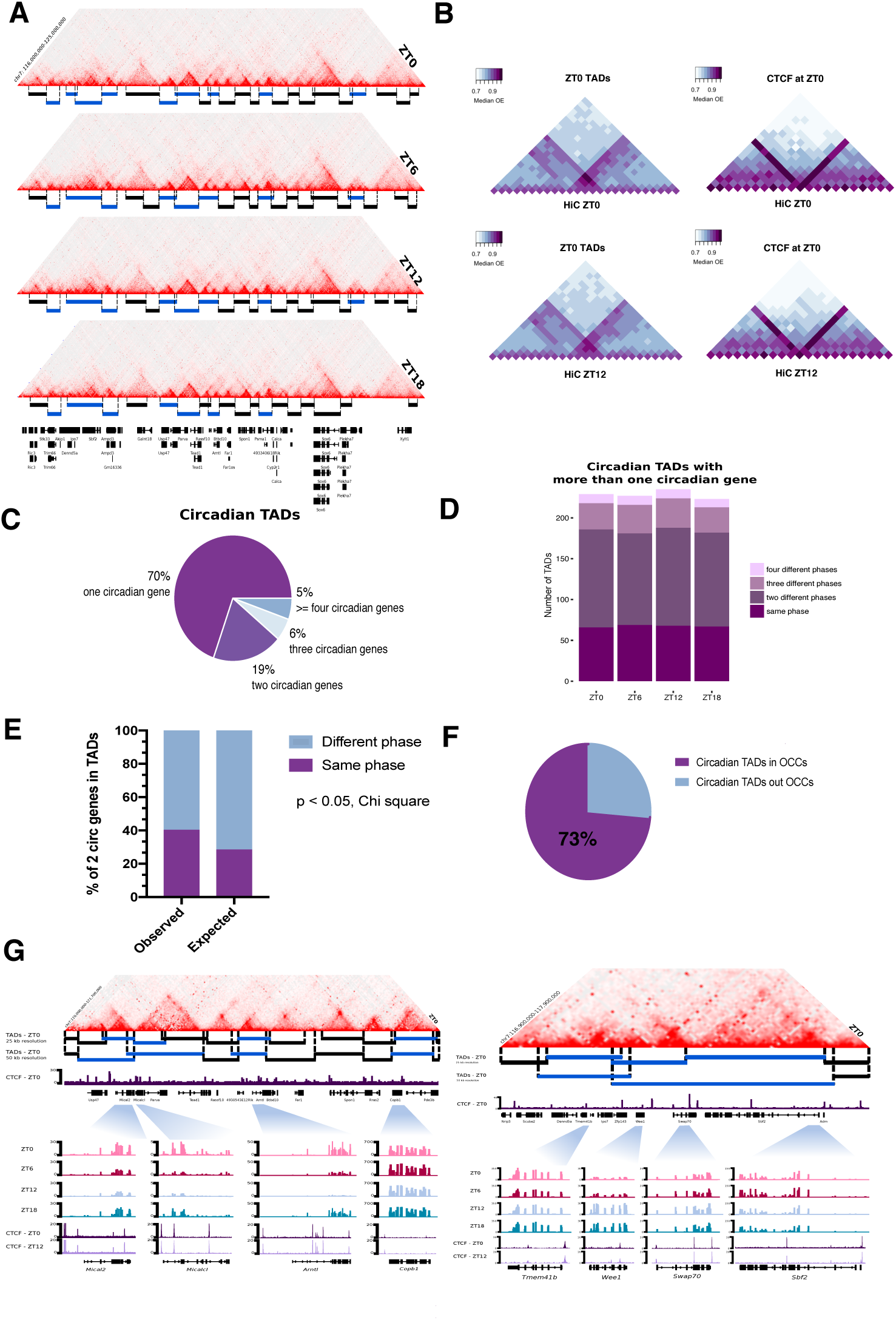
Circadian TADs isolate circadian genes with shared time of transcription. A. Observed Hi-C contact matrices showing the TADs landscape of the genomic region including the *Arntl1* gene locus at four time points during the circadian cycle. B. Left, 50kb resolution Median Observed/Expected ZT0 and ZT12 Hi-C signal around 1000 randomly selected TADs from ZT0 plotted on ZT0 and ZT12 Hi-Cs. TADs were scaled to fit the five central bins. Right, the same metaplots but for 1000 randomly CTCF peaks found at ZT0. CTCF peaks are at the central bin of the metaplot. C. Proportion of cTADs harbouring 1, 2, 3 or 4 circadian genes. D. Phase distribution of circadian genes sharing TADs. E. Observed and expected proportion of TADs with 2 circadian genes sharing transcriptional peak phase (p-value <0.05, Chi square test). F. Proportion of circadian TADs overlapping and non-overlapping OCCs (p-value<0.0001, t test) G. Examples of cTAD Hi-C contact matrices from ZT0. Left, cTADs allocating *Mical2, Micalcl, Arntl1* and *Copb1* circadian genes. Right, cTADs allocating *Tmem41b, Wee1, Swap70* and *Sbf2* circadian genes. Close up to each circadian gene with genomic tracks showing RNA-seq signal at all time points and CTCF ChIP-seq peaks at ZT0 and ZT12.

CTCF functions as an architectural protein that establishes chromatin domains together with cohesin in mammalian chromatin (Elphege, 2017; Fudenberg et al., 2016; Merkenschlager & Nora, 2016). We performed ChIP-seq against CTCF at ZT0 and ZT12 from the same samples as above. 33,262 CTCF sites (75.3%), out of a total of 44163 sites, were shared between ZT0 and 12 and CTCF showed similar chromatin occupancy between the two time points (Figure S3D). As expected, CTCF-bound regions exhibited robust contact insulation properties, independent of the time point examined (Figure 2B right panel and S3E top panels). Additionally, we assessed preservation of CTCF insulation between mouse ES cells and liver cells by overlaying regions occupied by CTCF, as identified in mESCs (Shen et al., 2012), onto our liver Hi-C data sets. Robust insulation was observed when using either ZT0 or ZT12 Hi-C data suggesting large agreement of CTCF chromatin occupancy and insulation properties between mESCs and adult liver tissue (Figure S3E bottom panels).

Next, we assigned genes to TADs. We found that on average TADs harboured 7.2 genes. TADs containing one or more circadian genes (cTADs) were larger than non-circadian TADs and on average contained more genes (14.2) (Figure S3F, all p-values < 2.2e-16, Wilcoxon rank sum test). We observed that 70% of cTADs contained only one circadian gene (Figure 2C). The remaining cTADs contained more than one circadian gene and remarkably, the circadian genes sharing TADs exhibited peak transcriptional expression at shared times during the day (40% for TADs with 2 circadian genes compared to the expected 28%, 18% of TADs with 3 circadian genes compared to the expected 8.8%, p < 0.0001, Chi square test) (Figure 2C,E, only data from cTADs with 2 circadian genes shown). Examples of cTADs containing one circadian gene (cTADs with *Mical2, MicalcI, Arntl* and *Copb1*, respectively), or more than one circadian gene (cTAD with *Wee1* and *Swap70*) are shown in Figure 2G (top and bottom respectively). We next analysed whether TADs encoding circadian genes displayed chromatin compartment switching over the circadian cycle. Indeed, the majority of cTADs (73%) overlapped with OCCs more than expected when compared to a random set of the same number of non-cTADs (Wilcoxon test, p< 0.0001) (Figure 2F). Examples of cTADs overlapping OCCs are shown for *Arntl* (Figure 1G) and *Npas2* (Figure 1H). These results show that cTADs often set phase coherence between multiple circadian genes in the same TAD. While most TADs maintain their structural boundaries over time, chromatin compartments overlapping cTADs switch between active and silent states throughout a circadian cycle.

### Gene Promoter-Promoter networks in the liver and its circadian component

To gain insights into chromatin contacts at the level of individual circadian genes we measured genome-wide promoter-promoter and promoter-regulatory element contacts at four time points during a circadian cycle using Promoter-CHi-C (Figure 3A) (Rubin, et al., 2017; Schoenfelder, et al., 2015; Schoenfelder, et al., 2015). Promoter-containing ligation products from Hi-C libraries were efficiently captured (∼ 70%) using 39,021 RNA probes, which hybridise to 22,225 genomic restriction fragments covering all annotated gene promoters in the mouse genome. We produced ∼1,560,000,000 total valid read pairs from the three biological replicates for the four time points, thus obtaining ∼390,000,000 valid ligation products per time point (Table S4). Capture of gene promoters increased the number of valid ligation products per promoter to ∼10-15 fold compared to Hi-C. An example of this enrichment is shown for the *Arntl* gene locus comparing a virtual 4C from *Arntl* gene promoter performed on the Hi-C versus the Promoter-CHi-C chromatin contact data sequenced at equivalent depth (Figure S4A close view, and S4B comparison of raw paired reads per restriction fragment in CHi-C vs Hi-C p<0.0001, Wilcoxon rank test). We estimated the statistical significance of interactions between pairs of promoters and between promoter and other, potentially gene regulatory genomic regions, using the CHiCAGO pipeline for each time point (methods) (Cairns et al., 2016). We obtained ∼150,000 statistically significant interactions per time point resulting in ∼600,000 statistically significant promoter interactions in total (Table S4).

**Figure 3.**
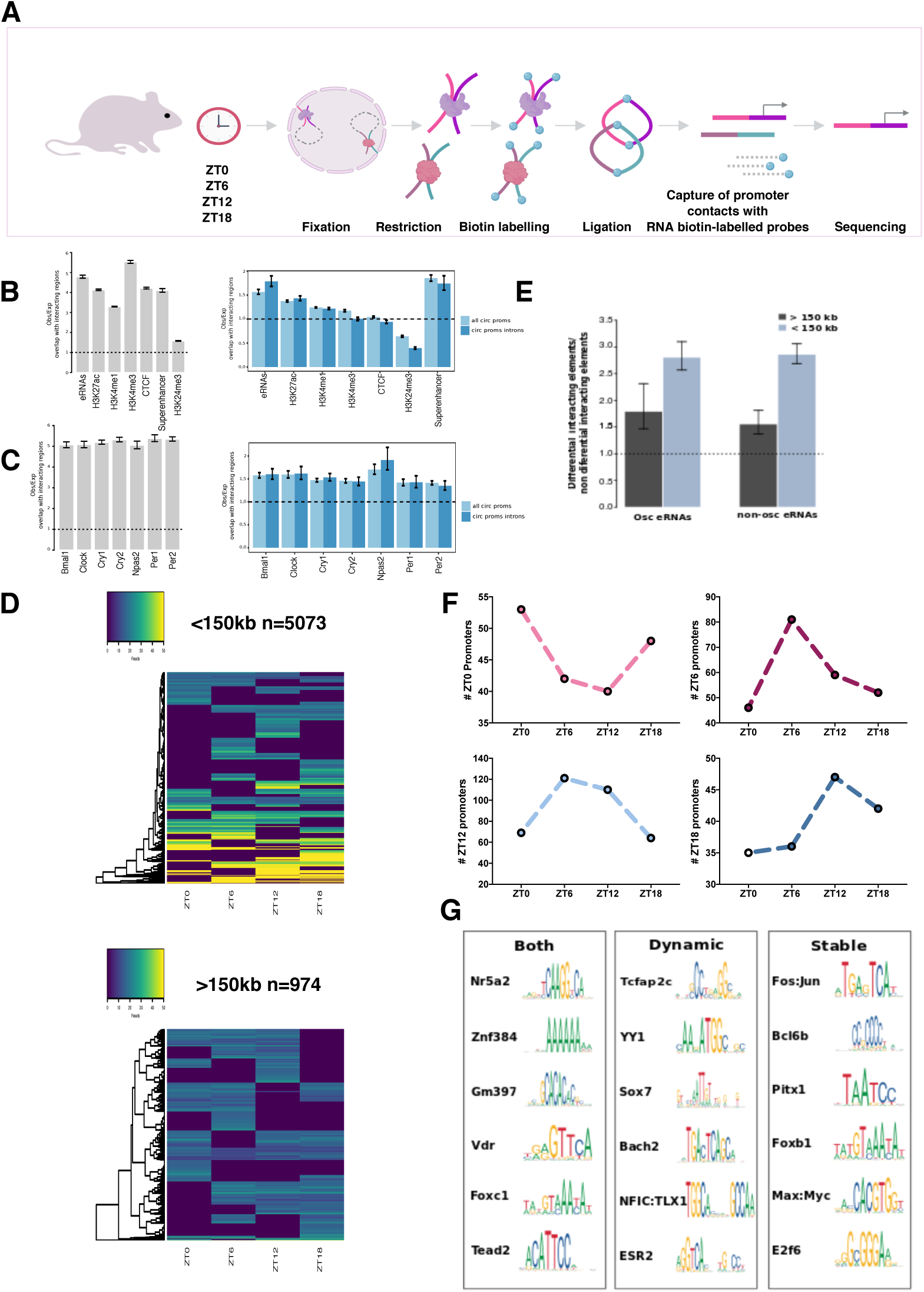
Promoter Capture-Hi-C and chromatin contact dynamics during a circadian cycle. A. Summary of the experimental workflow of Promoter-CHi-C technology. Cells are fixed, digested, filled and biotin labelled in nucleus. Pull down with streptavidin beads is then performed and Hi-C libraries prepared for sequencing. Using the Hi-C material as a template hybiridization is performed using the designed RNA biotinilated probes to capture promoters. A second pull down is performed to recover the hybrid molecules, DNA recovered and sequenced. B. Left, Obs/Exp signal ratio at promoter interacting regions of liver chromatin features including enhancers producing eRNAs, H3K27ac, H3K4me1, H3K4me3, CTCF, superenhancers and H3K27me3 (left) (all p-values < 8.632642e-123, t test). Obs/Exp signal ratio of the same chromatin features but for interacting regions for all oscillating gene promoters and intronically oscillating gene promoters compared to a random set of non-oscillating gene promoters (all p-values < 2.525639e-42 except H3K27me3). C. Right, the same as in D but for circadian TFs enrichment including Bmal1, Clock, Cry1, Cry2, Npas2, Per1 and Per2 for all gene promoters and left, oscillatory gene promoters (all p-values < pval<3.870992e-205, t test). D. Heat maps for read counts at detected dynamic contacts below and above 150 kbs (differential interactions supported by at least 15 reads, FDR< 0.1 and logFC > 1 or logFC < -1). E. Obs/Exp enrichment of enhancers producing eRNAs at dynamic contacts over stable contacts (p-value < 2e-16, t.test). F. Number of circadian gene promoters making the maximum number of contacts at ZT0, 6, 12 and 18 (p-value < 0.001, Chi square test). G. Transcription Factor DNA binding motifs significantly enriched at dynamic, stable or both circadian gene promoter chromatin contacts (E-value < 1.00e-002).

First, we focused on all gene promoter-promoter contacts. We built promoter-promoter networks at all time points and found large disconnected networks as expected for promoters scattered across the different chromosomes (Figure S5A, ZT0 shown). We then evaluated the larger clusters of the promoter-promoter network containing hundreds of connected gene promoters (Figure S5C, the 4 larger clusters shown including inter and intra chromosomal promoter-promoter interactions) and performed gene enrichment analysis on these using gProfiler (Raudvere et al., 2019) (Figure S5D). We found the Major Histocompatibility Complex forming one cluster with genes encoded on chromosomes 7 and 17. MHC genes are lowly expressed in the healthy liver and constitute dense, highly connected chromatin (Spengler et al., 1988). Similarly, a transcriptionally repressed cluster is formed between olfactory receptor genes on chromosome 7. In addition to repressed genes clustering in spatial proximity, we identified actively transcribed genes involved in glutathione synthesis and amino acid metabolism essential for liver detoxification function arranged in a promoter network. Another cluster encompassed constitutive histone genes located on chromosomes 3, 13, 11, 18, 12 and 7, among others (Figure S5C,D). Thus, prominent constitutive and liver specific promoter-promoter networks both transcriptionally active and inactive were identified in the adult liver.

Next we examined the circadian component in promoter-promoter networks. Circadian promoters are dispersed across chromosomes and networks described above as can be seen in Figure S5B. Nevertheless, we observed significantly increased contacts between circadian promoters in comparison to non-circadian genes (Figure S5E). When we analysed the maxima in mRNA abundance of circadian genes whose promoters were contacting each other, we found that diurnal and nocturnal circadian genes significantly contact each other respectively and this was even more striking when analysing circadian genes from our data and oscillating at the intronic level as defined by (Koike et al., 2012) or detected through GROseq (Fang et al., 2014) reflecting primary transcription (Figure S5F for the intronic gene set and Figure S4E for the GRO-seq circadian gene set, all p-values < 0.0001 Wilcoxon signed rank test). This is exemplified by the *Tef* gene promoter, which contacts the *Aco2* gene promoter ∼45 kb apart and both their shared pre-messenger expression peak at ZT12 (Figure S5G). Likewise, promoters of *Rorc* and *Cgn*, which form spatial contacts bypassing a genomic distance of ∼360 kb, were both found to drive peak expression around ZT18 (Figure S4F). These results show that circadian promoters physically interacting in the nuclear space have maximal transcriptional activity at similar times during the day.

### Regulatory elements form dynamic and stable chromatin contacts with circadian gene promoters

We next examined contacts between promoters and non-promoter genomic regions. Genomic regions significantly interacting with gene promoters including circadian gene promoters in mouse liver, showed enrichments for histone modifications characteristic for open chromatin and regulatory elements such as H3K27ac, H3K4me1, H3K4me3, H3K27me3 as well as the structural protein CTCF (Yue Feng et al., 2014) compared to distance matched non-interacting regions (Figure 3B all histone modifications p value < 8e-166, t test, CTCF, p value < 8e-18; t test). A set of enhancers from which enhancer RNAs (eRNAs) are produced have been described in the liver (Fang et al., 2014). We found a significant enrichment of these enhancers at the gene promoter contacted regions (Figure 3E p value < 9e-165; t test). Overall, the chromatin features at promoter-contacting regions demonstrate the efficient recovery of elements with possible structural and/or regulatory functions by P-CHi-C in our experiments.

Besides enhancer and open chromatin marks, we found significant enrichment of core clock transcription factor occupancy (Koike et al., 2012a) at promoter interacting regions (Figure 3C, pval<8e-100, t test). When measuring the same enrichments at only circadian gene promoters interacting regions and comparing them with non-circadian gene promoter contacts, we found a significant preference for circadian gene promoters to contact with enhancers both detected by eRNA transcription or histone modifications, as well as regions occupied by core clock TF (Figure 3C and D; all pval<8e-216, t test). We next assessed the rhythmicity of contacts between regulatory elements and circadian promoters during a circadian cycle. We identified dynamic genomic contacts involving circadian promoters using two distance regimes as described (see methods) (Figure 3D). In total we found 13,782 stable and 6,047 dynamic contacts for 1,195 circadian promoters and found enhancers significantly enriched at dynamically contacted regions (Figure 3E). We next analysed the number of interactions made by circadian promoters at different time points. We found that circadian gene promoters form a maximal number of contacts during or around the phase with maximal mRNA level, (Chi square, p < 0.001) (Figure 3F) suggesting more contacts are made together with increased transcription (see different examples of circadian gene promoter virtual 4C signal across the paper Figure 4E-G Figures S4F, S5G, S6A-F).

**Figure 4.**
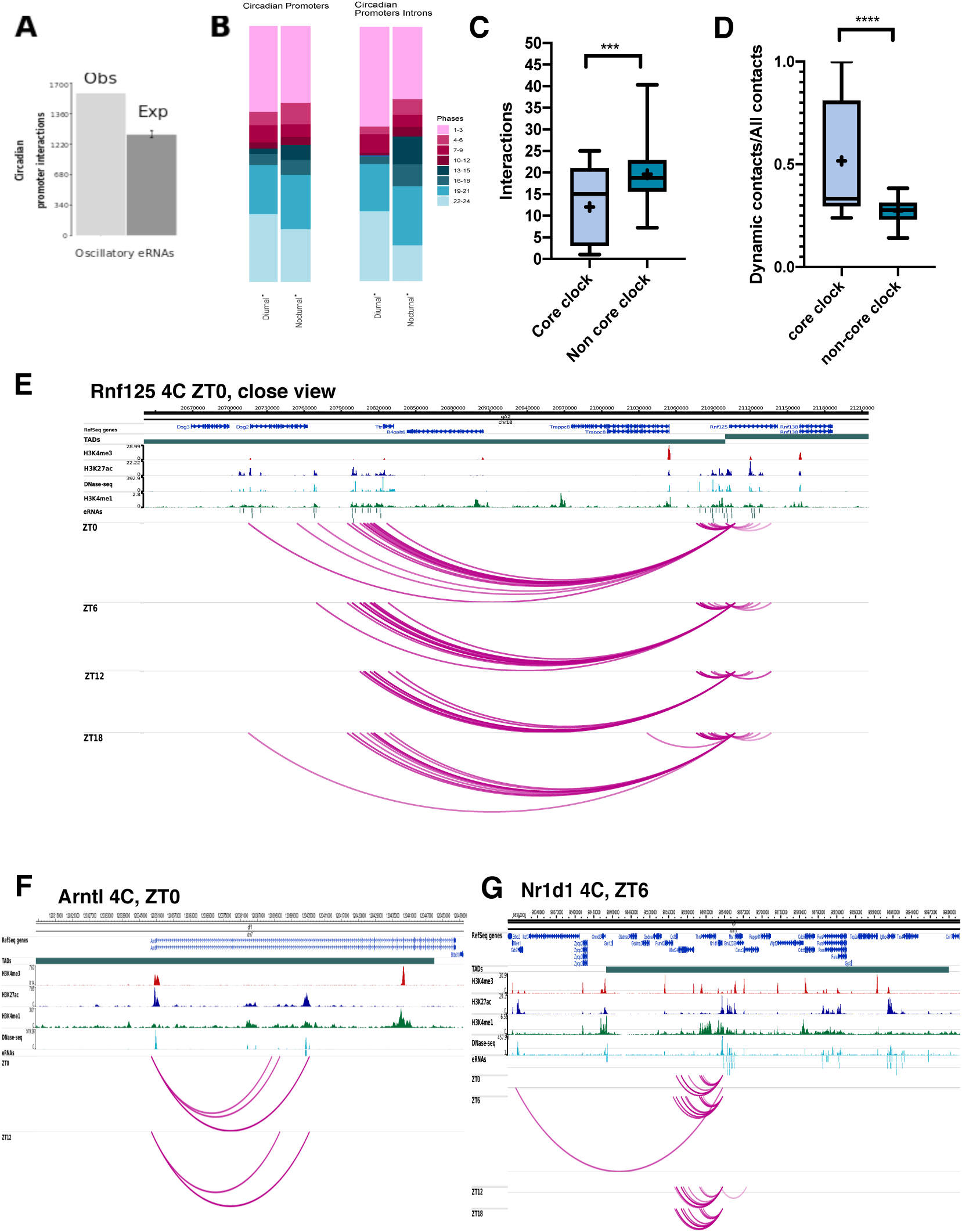
Transcriptional phase coherence between circadian genes and transcribed enhancers and the core-clock circadian genes display highly dynamic chromatin contacts. A. Circadian gene promoter observed and expected contacts with enhancers producing eRNA (p-value < 0.001, t test). B. Phase distribution of eRNAs produced from enhancers contacting all diurnal and nocturnal circadian promoters and circadian genes oscillating at the intronic level (all p-values < 0.001, Wilcoxon ranked sum test). C. Number of total significant interactions for core clock circadian gene promoters and a random control set of circadian gene promoters (p-value < 0.0001, Mann Whitney test). D. Proportion of dynamic contacts for core clock genes and a random control set of circadian gene promoters (p-value < 0.0001, Mann Whitney test). E. Partial virtual 4C landscape of the *Rnf125* gene promoter at the four time points during the day. Significant contacts with enhancers producing oscillatory eRNA are shown. The majority of the eRNAs present a peak in transcription at ZT0 as *Rnf125* does. Acrophase is written next to the gene name. Genomic tracks show significant contacts as arcs and chromatin features including liver H3K4me3, H3K4me1, H3K27ac, DNaseI, eRNAs and TADs. F. Virtual 4C for *Arntl and Nr1d1* (G) core clock gene promoters at all time points during the day. Features displayed are the same described in E.

Next we aimed to identify transcription factor binding motifs at genomic regions involved in constant or rhythmic interactions with circadian promoters in an unbiased manner. Using the MEME suite (methods) (Bailey et al., 2015), we found several binding motifs significantly enriched in both types of contacts as well as binding motifs specifically occurring in dynamic or constant contacts (Figure 3G, Table S5). Transcription factor binding sites associated with both constant and dynamic contacts were shared between promoters of circadian genes expressed at different phases. Among such ubiquitous sites the nuclear receptor Nr5a2/LRH-1 binding motif was highly enriched. Nr5a2 is a nuclear receptor that acts as a key metabolic sensor by regulating genes involved in bile acid synthesis and cholesterol homeostasis trough regulation of *Cyp7a1* and *Cyp8a1* circadian genes, triglyceride synthesis and lipid composition and metabolism (Chen et al., 2003; Chong, et al., 2012; Matsukuma, et al.,, 2007; Wu et al., 2016). Interestingly, deficiencies in Nr5a2 are linked to non-alcoholic fatty liver disease where it seems to have an anti-inflammatory role (Schwaderer et al., 2020). In addition, lack of Nr5a2 in the adult liver leads to disruption of hepatic lipid homeostasis and composition (Miranda et al., 2018). Our data implies that Nr5a2 could be important in circadian chromatin loop formation. Other binding sites for nuclear receptors enriched in regions contacting circadian promoters included VDR (Vitamin D Receptor) and ESR2 (Estrogen Receptor 2).

Tcfap2c/AP-2 gamma binding motif was found to be highly enriched at dynamic interacting regions of circadian gene promoters. In the liver this TF has been associated with repression of fatty acid synthesis pathways (Holl et al., 2011) and was identified as a key TF involved in lipid droplets biogenesis (Scott et al., 2018). Fos:Jun/AP1 binding sites were found in genomic regions forming stable contacts with circadian promoters. AP1 factors are a well-characterized immediate early transcription factors induced in response to signals in the serum and which regulate the expression of circadian genes in liver and cultured cells (Balsalobre et al., 1998) as well as the SCN (Y. Chen et al., 2018; Guido et al.,1999; Schwartz et al., 2000). Recently, AP-1 has been implicated in stable and dynamic loop formation during macrophage development bringing together key macrophage genes and enhancers (Phanstiel et al., 2017).

In summary, our results identified a distinct set of DNA binding sites for nuclear receptors and immediate early genes in regions contacting circadian promoters, which could function in the wiring of the circadian promoter 3D interactome in the liver.

### Diurnal and nocturnal circadian gene promoters contact diurnal and nocturnal enhancers in the nuclear space

A set of enhancers has been shown to be transcribed in a circadian fashion in mouse liver (Fang et al., 2014). When analysing the interactions from this subset of rhythmic liver enhancers we found they preferentially contacted circadian gene promoters over other gene promoters, suggesting that rhythmically transcribed genomic regions, protein-coding and non-coding, contact each other in nuclear space (Figure 4A). The same finding resulted when we restricted our analysis to promoters of circadian genes whose intronic portions oscillated in a circadian manner or detected through GRO-seq (Fang et al., 2014) reflecting primary transcriptional oscillation (Figure S4C).

We then compared the transcriptional phases between promoters of circadian genes and their corresponding contacted enhancer elements with rhythmic transcription. We found a significant contact preference between diurnal and nocturnal promoters and diurnal and nocturnal enhancers, respectively. The phase bias was more pronounced when analysing circadian genes, which oscillated at the intronic level and detected through GRO-seq (Figure 4B our intronic set, Figure S4D, detected through GRO-seq. all p-values < 0.0001 Wilcoxon signed rank test). For example, the *Rnf125* circadian gene promoter with peak transcription at ZT0 contacts 12 rhythmically expressed enhancers with acrophases between 19 and 1 hours during the circadian cycle. Furthermore, as it can be observed, the number of contacts with the enhancers, increase during the acrophase (Figure 4E).

### The core clock gene promoter contacts

Finally, we focused on the genomic significant interactions formed by circadian core clock gene promoters including *Npas2, Clock, Arntl, Cry1, Cry2, Per1, Per2, Rorc, Nr1d1, Nr1d2* as defined by (Anafi et al., 2014). Notably, all core clock genes displayed fewer overall contacts compared to a random set of the same number of other circadian genes in the liver (12 vs 19.6 mean number of contacts for core-clock vs other circadian genes, Figure 4C, p<0.0001, t test). The contacts formed by the core clock gene promoters however, were significantly more dynamic than a random set of the same number of contacts for other circadian genes in the liver (42.3% vs 26.8% mean proportion of dynamic contacts for core clock vs other circadian genes, Figure 4D, p<0.0001, t test). For instance the promoter regulating *Arntl1* expression engages in contacts with two enhancer elements at ZT18, the time when *Arntl* expression increases, in three contacts at ZT0, at maximal transcriptional output, and with no significant contacts at both ZT6 and ZT12, when *Arntl1* transcription ceases (Figure 4F). The *Nr1d1* gene promoter exhibited higher connectivity at the gene’s expression acrophase around ZT6 (Figure 4G). In contrast to core clock contact profiles (additional examples are shown for *Per2, Nr1d2* and *Npas2*, Figure S6A-D), promoters of circadian output genes form contacts with significantly more genomic regions with a larger contribution of constantly interacting elements as exemplified by *PTG* and *Dhr3* gene promoters (Figure S6E,F). In conclusion, core-clock gene promoters engage in more dynamic genomic contacts with fewer genomic elements compared to other circadian genes in the liver.

## Discussion

Circadian fluctuations in gene expression in the adult liver orchestrate essential physiological metabolic responses in the body. While molecular mechanisms underlying the circadian clock circuitry have been described at transcriptional and post-transcriptional levels, less is understood of how different genome structures contribute to or reflect cyclic gene expression (Yeung & Naef, 2018). Here we analysed genome conformation in mouse adult liver throughout a circadian cycle and report the properties of the circadian cis-regulatory chromatin landscape at different genomic scales associated with circadian rhythmicity in gene expression.

We found 17% of the genome fluctuates between A and B compartments during a 24-hour cycle. The genomic regions with changing compartment assignments throughout the day (OCCs) overlap with circadian TADs whose domain boundaries in contrast remain unchanged during a cycle. Switches between closed and open states of genomic compartments have been reported during organism development, cell differentiation and cell cycle (Bonev et al., 2017; Dixon et al., 2015; Nagano et al., 2017). However, our results reveal that dynamic changes between compartment states occur also within hours and without cells dividing or changing their identity (Figure 5).

**Figure 5.**
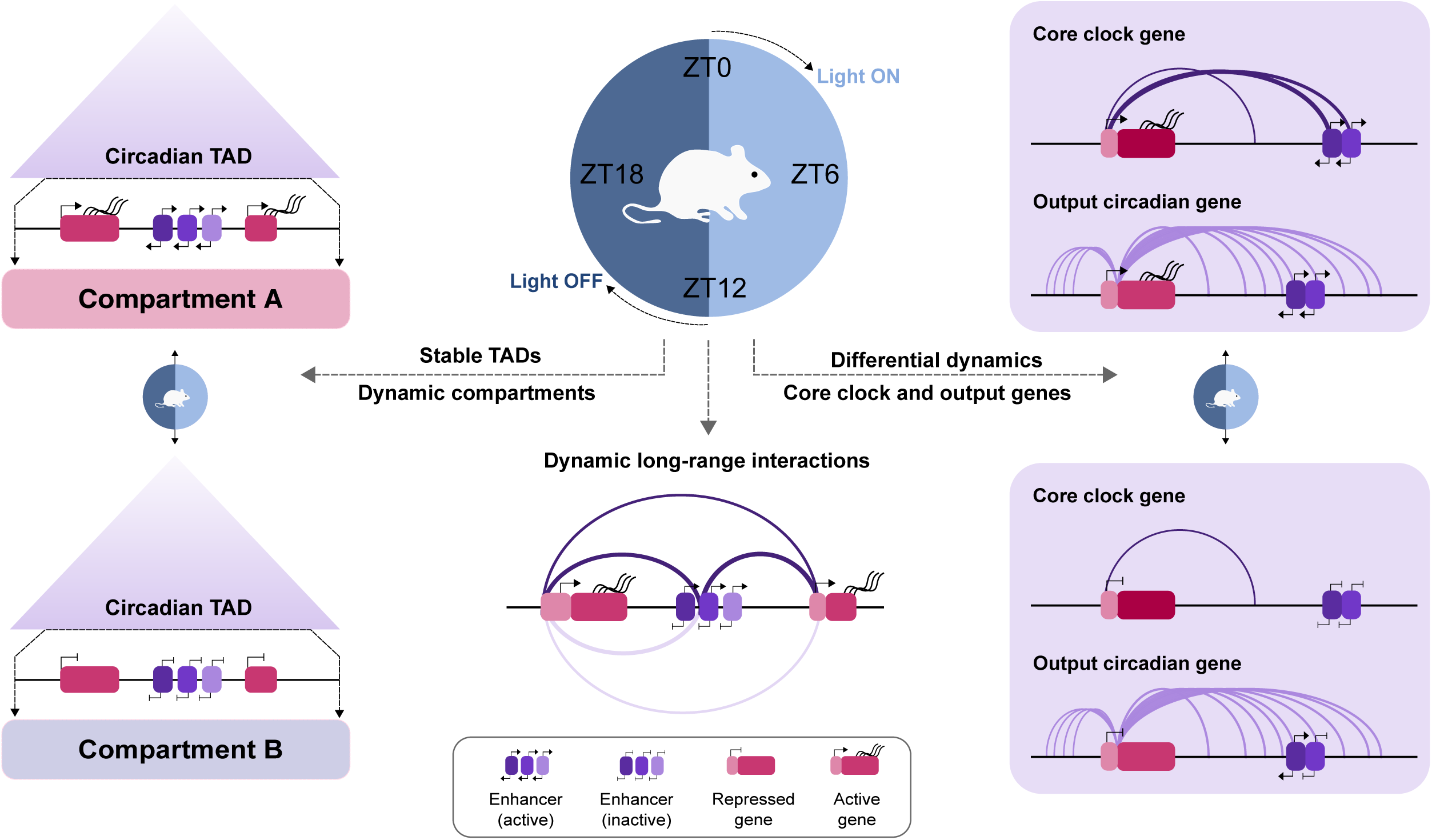
Chromatin conformation dynamics during a circadian cycle. Circadian TADs containing one or more circadian genes remain stable but switch chromatin compartments in correlation with transcriptional activation and gain of open histone modifications marks during the 24 hours (left). Inside cTADs, circadian genes and regulatory elements with similar acrophases contact each other in the nuclear space with interactions increasing at the time of peak transcription (middle). Core clock genes display highly dynamic contacts during the 24 hours. In contrast circadian output genes have more saturated contact profiles that remain stable during the day (right).

Within TADs encompassing multiple transcribed loci, circadian genes tend to be alone or sharing the TAD with other circadian genes and transcribed regulatory elements, with similar transcriptional acrophase suggesting spatial isolation of temporal transcription control. However, the contacts formed by circadian gene promoters can be constantly maintained or dynamically changed over time (Figure 5).

We performed an unbiased identification of TF binding motifs to discover candidate protein factors enriched at the anchor sites of chromatin loops involving circadian genes engaged in both dynamic and stable contacts. We found metabolic liver nuclear receptors (Nr5a2/LRH-1) and immediate early genes AP2 gamma and Fos:Jun enriched in chromatin regions contacting circadian gene promoters in both, dynamic and stable fashions. This suggests that specific nuclear receptors together with immediate early factors participate in shaping the circadian 3D cistrome.

Finally, by comparing the interaction profiles between core clock and output circadian genes we discovered that core clock genes tend to contact fewer different genomic elements and that core clock interactomes are far more dynamic compared to output circadian genes in the liver. This suggests a robust regulation of core clock gene transcription by a few specific regulatory elements, which dynamically contact the core clock promoters. Alternatively, the control of core clock gene expression might rely primarily on their respective promoters and fewer distal genomic elements than for output genes. On the contrary, output circadian genes have more complex and constantly maintained contact profiles, which suggests the participation of an increased number of regulatory elements required to control their expression in a pre-formed genome architecture. Our genome-wide observation is in line with and extends previous candidate-scale 4C chromosome capture experiments for a set of core clock and output genes (Mermet, Yeung, & Naef, 2020).

In conclusion, our results provide evidence that chromatin architecture dynamics during a circadian cycle take place in parallel with circadian oscillations in transcription and provide a genome-wide atlas of the liver genome conformation resolved in time.

## Acknowledgements

M.F.-M. is supported by UNAM Technology Innovation and Research Support Program (PAPIIT) IN207319 and was supported by MODHEP (grant agreement no. 259743). J.C., S.W.W, and P.F. were supported by BBSRC grant BB/J004480/1, and C.V. was supported by ERC advanced grant 340152 DEVOCHROMO to P.F.

## Author Contributions

M.F.-M. designed the project, performed the experiments, analysed data and wrote the manuscript with contribution from all authors specially R.A.M. and J.M. M.A.-K. analysed data. R.G.A.-M. analysed data. J.M. assisted with sample collection, liver RNA extraction and performed RT-PCRs. J.C. ran CHiCAGO to detect significant interactions from CHi-C. A.C.P.-H. and B.V. performed promoter-promoter network analysis. C.V. performed initial analysis of Hi-C data sets. S.A. performed initial analysis of RNA-seq data sets and provided bioinformatics advice. S.W.W. ran HiCUP pipeline in Hi-C and CHi-C data sets. P.F. helped with project design and data interpretation.

## Methods

### Key resources table

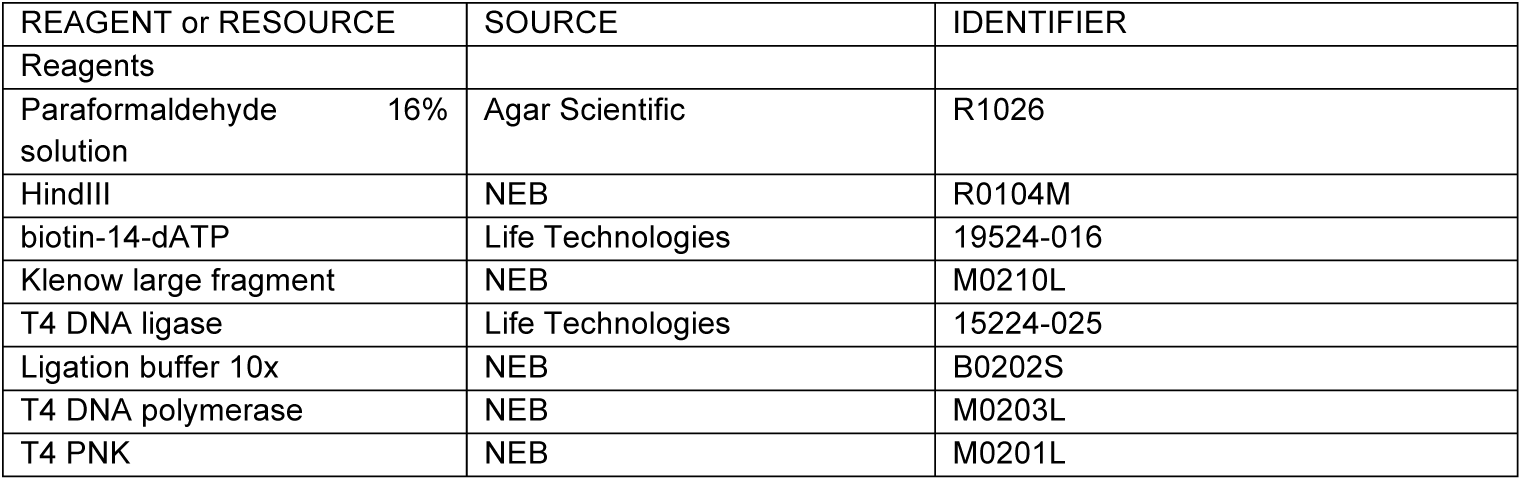
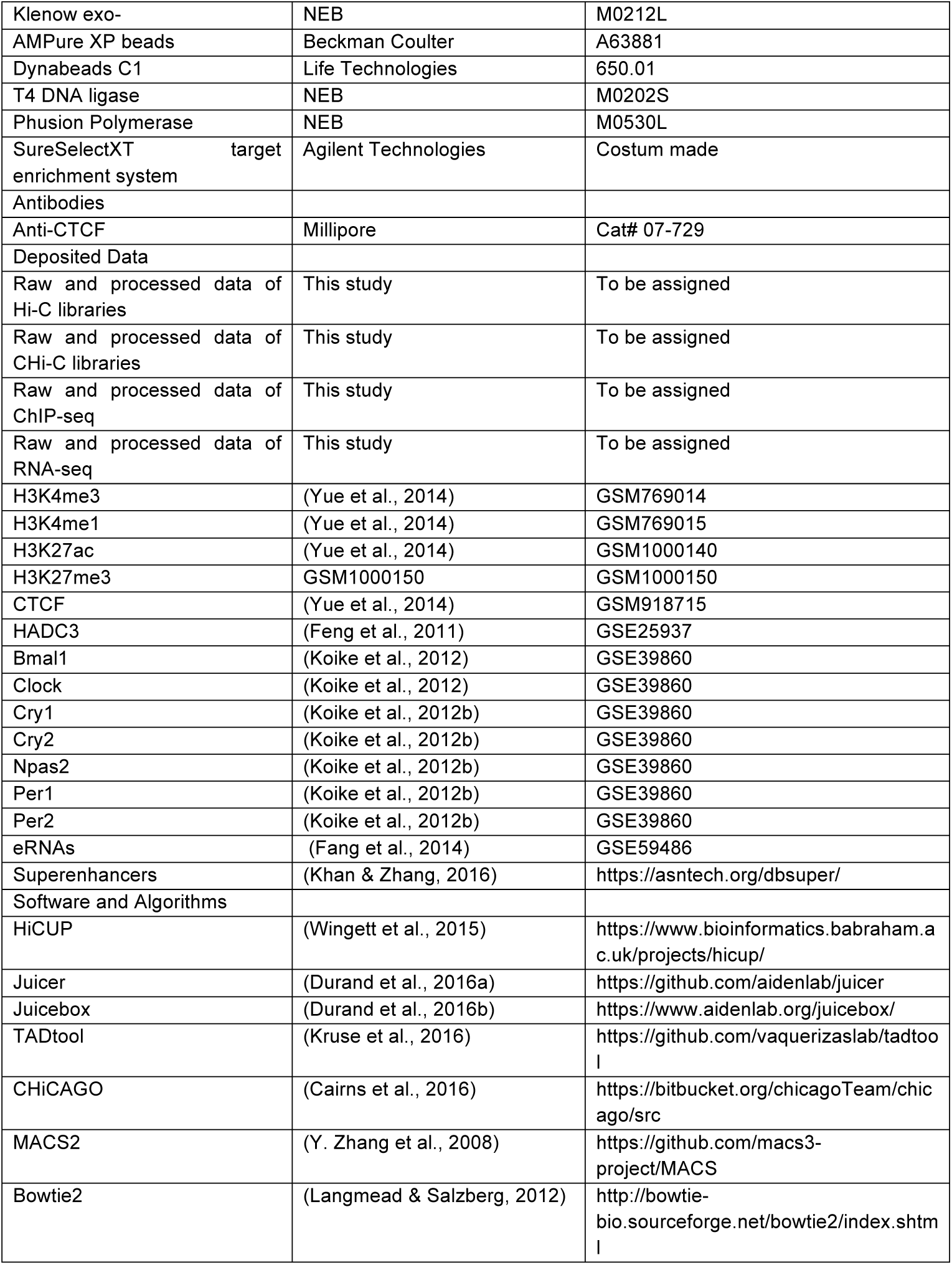
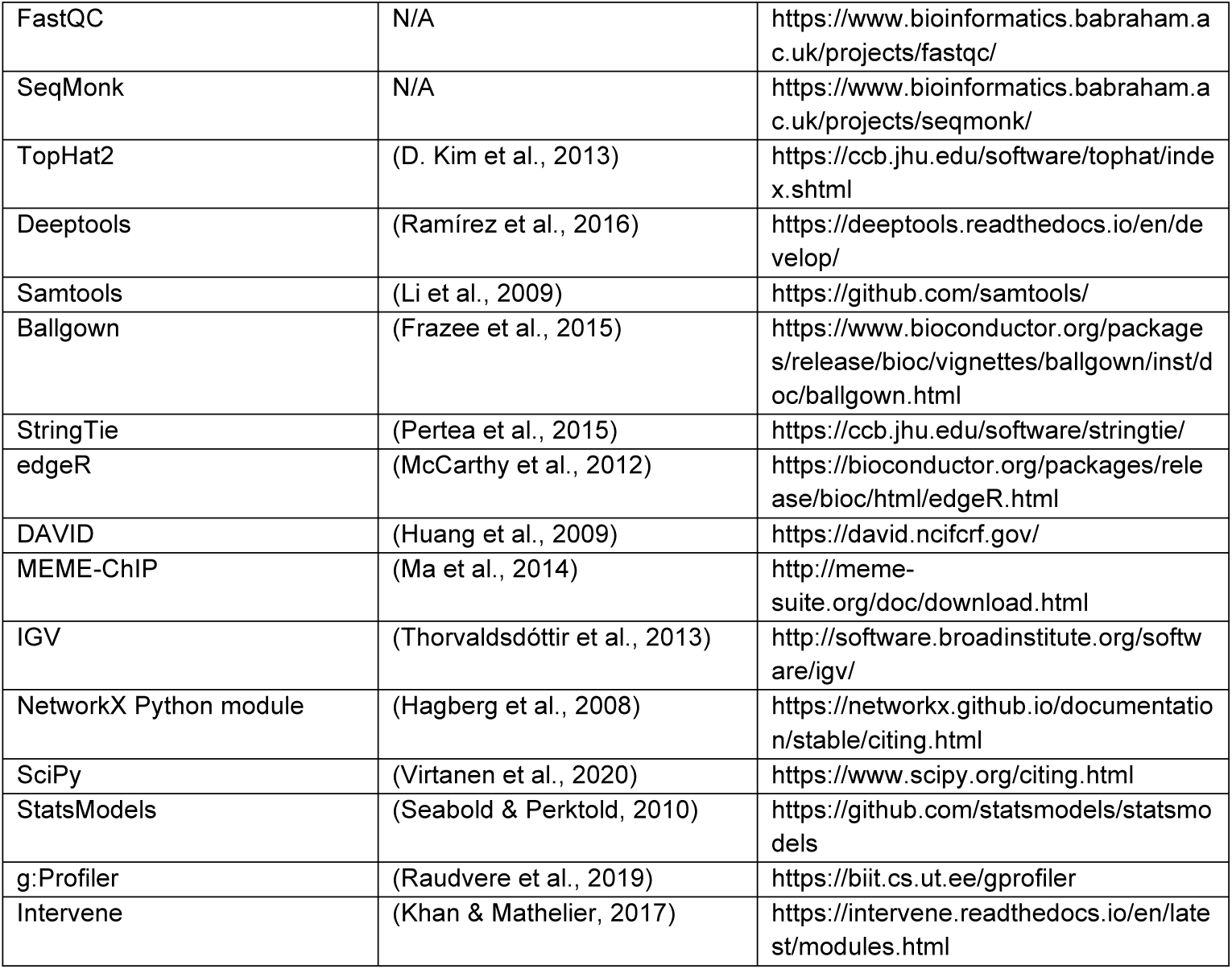

| REAGENT or RESOURCE | SOURCE | IDENTIFIER |
| --- | --- | --- |
| Reagents |  |  |
| Paraformaldehyde 16% solution | Agar Scientific | R1026 |
| HindIII | NEB | R0104M |
| biotin-14-dATP | Life Technologies | 19524-016 |
| Klenow large fragment | NEB | M0210L |
| T4 DNA ligase | Life Technologies | 15224-025 |
| Ligation buffer 10x | NEB | B0202S |
| T4 DNA polymerase | NEB | M0203L |
| T4 PNK | NEB | M0201L |
| Klenow exo- | NEB | M0212L |
| AMPure XP beads | Beckman Coulter | A63881 |
| Dynabeads C1 | Life Technologies | 650.01 |
| T4 DNA ligase | NEB | M0202S |
| Phusion Polymerase | NEB | M0530L |
| SureSelectXT target enrichment system | Agilent Technologies | Costum made |
| Antibodies |  |  |
| Anti-CTCF | Millipore | Cat# 07-729 |
| Deposited Data |  |  |
| Raw and processed data of Hi-C libraries | This study | To be assigned |
| Raw and processed data of CHi-C libraries | This study | To be assigned |
| Raw and processed data of ChIP-seq | This study | To be assigned |
| Raw and processed data of RNA-seq | This study | To be assigned |
| H3K4me3 | (Yue et al., 2014) | GSM769014 |
| H3K4me1 | (Yue et al., 2014) | GSM769015 |
| H3K27ac | (Yue et al., 2014) | GSM1000140 |
| H3K27me3 | GSM1000150 | GSM1000150 |
| CTCF | (Yue et al., 2014) | GSM918715 |
| HADC3 | (Feng et al., 2011) | GSE25937 |
| Bmal1 | (Koike et al., 2012) | GSE39860 |
| Clock | (Koike et al., 2012) | GSE39860 |
| Cry1 | (Koike et al., 2012b) | GSE39860 |
| Cry2 | (Koike et al., 2012b) | GSE39860 |
| Npas2 | (Koike et al., 2012b) | GSE39860 |
| Per1 | (Koike et al., 2012b) | GSE39860 |
| Per2 | (Koike et al., 2012b) | GSE39860 |
| eRNAs | (Fang et al., 2014) | GSE59486 |
| Superenhancers | (Khan & Zhang, 2016) | https://asntech.org/dbsuper/ |
| Software and Algorithms |  |  |
| HiCUP | (Wingett et al., 2015) | https://www.bioinformatics.babraham.ac.uk/projects/hicup/ |
| Juicer | (Durand et al., 2016a) | https://github.com/aidenlab/juicer |
| Juicebox | (Durand et al., 2016b) | https://www.aidenlab.org/juicebox/ |
| TADtool | (Kruse et al., 2016) | https://github.com/vaquerizaslab/tadtool |
| CHiCAGO | (Cairns et al., 2016) | https://bitbucket.org/chicagoTeam/chicago/src |
| MACS2 | (Y. Zhang et al., 2008) | https://github.com/macs3-project/MACS |
| Bowtie2 | (Langmead & Salzberg, 2012) | http://bowtie-bio.sourceforge.net/bowtie2/index.shtml |
| FastQC | N/A | https://www.bioinformatics.babraham.ac.uk/projects/fastqc/ |
| SeqMonk | N/A | https://www.bioinformatics.babraham.ac.uk/projects/seqmonk/ |
| TopHat2 | (D. Kim et al., 2013) | https://ccb.jhu.edu/software/tophat/index.shtml |
| Deeptools | (Ramírez et al., 2016) | https://deeptools.readthedocs.io/en/develop/ |
| Samtools | (Li et al., 2009) | https://github.com/samtools/ |
| Ballgown | (Frazee et al., 2015) | https://www.bioconductor.org/packages/release/bioc/vignettes/ballgown/inst/doc/ballgown.html |
| StringTie | (Pertea et al., 2015) | https://ccb.jhu.edu/software/stringtie/ |
| edgeR | (McCarthy et al., 2012) | https://bioconductor.org/packages/release/bioc/html/edgeR.html |
| DAVID | (Huang et al., 2009) | https://david.ncifcrf.gov/ |
| MEME-ChIP | (Ma et al., 2014) | http://meme-suite.org/doc/download.html |
| IGV | (Thorvaldsdóttir et al., 2013) | http://software.broadinstitute.org/software/igv/ |
| NetworkX Python module | (Hagberg et al., 2008) | https://networkx.github.io/documentation/stable/citing.html |
| SciPy | (Virtanen et al., 2020) | https://www.scipy.org/citing.html |
| StatsModels | (Seabold & Perktold, 2010) | https://github.com/statsmodels/statsmodels |
| g:Profiler | (Raudvere et al., 2019) | https://biit.cs.ut.ee/gprofiler |
| Intervene | (Khan & Mathelier, 2017) | https://intervene.readthedocs.io/en/latest/modules.html |

### Contact for reagent and resource sharing

Further information and requests for resources and reagents should be directed and will be fulfilled by the Lead Contact Mayra Furlan-Magaril, Phone: +52 55 56225739

## Experimental model and subject details

### Method details

#### Mice and Tissue isolation

C57BL/6 male mice were maintained in the Babraham Institute Animal Facility and all applied procedures were approved considering the animal welfare practices according to the Home Office in the UK. 8 week-old male mice were maintained under a 12 hours light:12 hours dark cycle for two weeks and fed *ad libitum*. Livers were dissected from three biological replicate pools (Pool A, B and C) composed of 2 livers each at four time points ZT0, 6,12 and 18, with ZT0=lights on and ZT12=lights off. The liver tissue was chopped into 5-6 mm^3^ pieces and directly fixed in 2% formaldehyde for 10 minutes.

#### *In nucleus* Hi-C

In nucleus Hi-C library generation was performed as previously described (Rubin et al., 2017a). Briefly, fixed adult liver tissue from three biological replicates at ZT0 0, 6, 12 and 18 was sieved through a 70µM cell strainer and dounce homogenized in 10 ml of ice-cold lysis buffer with a tight pestle for a total of 30 strokes on ice. Nuclei were washed and permeabilized with 0.3% SDS for 45 minutes at 37 °C and then incubated overnight with HindIII at 37 °C, DNA ends were labeled with biotin-14-dATP (Life Technologies) in a Klenow end-filling reaction and ligated in nuclei overnight. DNA was purified by phenol-chloroform the concentration was measured using Quant-iT PicoGreen (Life Technologies). A total of 10 µg of DNA was sheared to an average size of 400 bp using a Covaris machine and following the manufacturer’s instructions. The sheared DNA was end-repaired, adenine tailed, and subject to a double size selection using AMPure XP beads to isolate DNA ranging from 250 to 550 bp in size. Ligated fragments marked by biotin-14-dATP were pulled-down using MyOne Streptavidin C1 DynaBeads (Invitrogen) and ligated to paired-end adaptors (Illumina). The in nucleus Hi-C libraries were amplified using the PE PCR 1.0 and PE PCR 2.0 primers (Illumina) using 6–9 PCR amplification cycles as required.

#### Promoter Capture in nucleus Hi-C (Chi-C)

Promoter Capture was performed as previously described (Rubin, et al., 2017; Schoenfelder et al., 2015; Schoenfelder et al., 2015). Briefly, Biotinylated 120-mer RNA baits were designed to target both ends of HindIII restriction fragments overlapping the Ensembl promoters of protein-coding and noncoding transcripts and UCEs as described in detail in Schoenfelder et al., 2015a. Promoter Capture was carried out using *in nucleus* Hi-C libraries derived from three biological replicates at ZT0 0, 6, 12, and 18 with the SureSelect target enrichment system and the biotinylated RNA bait library according to the manufacturer’s instructions (Agilent Technologies). After library enrichment, a post-capture PCR amplification step was carried out using the PE PCR 1.0 and PE PCR 2.0 primers (Illumina) with 4–6 PCR amplification cycles as required. *In nucleus* Hi-C and CHi-C libraries were sequenced on the Illumina HiSeq 2000 platform.

#### ChIP-seq

For ChIP–seq, liver tissue for two biological replicates at ZT0 and ZT12 was dissected as processed as for Hi-C and then fixed in 1% formaldehyde for 5 minutes. Chromatin immunoprecipitation was performed as described (Rubin et al., 2017) using 10 µg of α-CTCF (Millipore, 07-729). DNA was purified using Zymo Research DNA purification columns. Sequencing libraries were prepared with the NEBNext ChIP–seq library prep kit (NEB) according to manufacturer’s instructions. DNA was purified using AMPure beads (Agencourt).

#### RNA-seq

For RNA-seq libraries, total RNA was purified from the same livers processed for *in nucleus* Hi-C from four biological replicates at ZT0, 6, 12, and 18. Sequencing libraries were prepared with TruSeq Stranded Total RNA Gold Library Prep Kit v2 (Illumina).

### Data Processing

#### Hi-C analysis

Hi-C sequenced reads were mapped to the mouse genome (mm9) using HiCUP (Wingett et al., 2015) with default parameters. Downstream processing was done using Juicer (Durand, et al., 2016) and data was visualized using Juicebox (Durand, Robinson, et al., 2016). Hi-C heatmaps at different bin resolutions were created and normalized using Knight-Ruiz (KR) matrix balancing algorithm from Juicer. Statistics for each library can be found in Table S1.

Metaplots. The metaplots were created using python custom scripts. Briefly, the script takes a feature of interest and calculates the frequency of interactions around it using as input the KR normalized Obs/Exp Hi-C matrices at different resolutions (10, 25, or 50 Kb) from different time points (ZT0,6,12,18). The final metaplot is the median value of all the plots for the list of anchors. For the TAD-anchored metaplots, Hi-C normalized matrices at 50kb resolution were used. Each TAD (see TAD calling) was scaled to fit into 5 bins, and only 1,000 TADs from all datasets and using all chromosomes were randomly chosen to reduce computing time. For the CTCF-anchored metaplots, Hi-C normalized matrices at 10kb resolution were used. Each CTCF peak (see ChIP-Seq analysis) was scaled to fit into a single bin, and only 1000 CTCF peaks identified at ZT0 and ZT12 were randomly chosen to reduce computing time. The matrices generated were plotted using heatmap.2 from the package plots.

#### TAD calling

For all time points, we retrieved Knight-Ruiz normalized contact matrices from Juicer for all chromosomes at 25kb and 50kb resolution. TADs were identified using TADtool (Kruse et al., 2016) with the insulation score algorithm. To find appropriate parameters for TAD identification we called TADs for chromosome 1 across all time points using contact matrices at 25kb and 50kb resolution and a window size of 100, 150, 155, 175, 195 and 200kb over threshold values from 70 to 200. For all data sets at 50kb resolution, we called TADs with a window size value of 200kb and a threshold value of 140 while for all data sets at 25 kb resolution, we called TADs with a window size value of 100kb and a threshold value of 76. We found that these parameters show good agreement between identified TADs and visual inspection of Hi-C datasets in Juicer. Of note, visual inspection of Hi-C datasets with TADs identified at 25kb resolution reveals that these represent sub-TADs contained within TADs identified at 50kb resolution.

#### TAD analysis

##### Size distribution

The number of TADs per time point and the size distribution of TADs across a circadian cycle was calculated using a custom R script using a Wilcoxon rank sum test.

##### Circadian TADs

Circadian TADs were defined as previously described (Y. H. Kim et al., 2018). TADs containing at least one circadian gene as identified by our RNA-seq analysis were classified as Circadian TADs. Each Circadian TAD was further categorized depending on the number of circadian genes contained within the TAD. Then we considered the transcriptional phases of the circadian genes within Circadian TADs and classified them as same or different. The probability to have the same or different transcriptional phases within cTADs was calculated considering the total number of circadian genes per phase. The observed over expected significance was estimated performing a Chi square test.

##### Overlap

To determine the number of unique and shared TADs between the time points we calculated the overlap in different pair-wise comparisons using the Venn module of Intervene (Khan & Mathelier, 2017). A TAD was considered to be shared between time points if more than 80% of the genomic domain region overlapped with a domain from a different Hi-C data set.

#### Compartment analysis

Compartments were identified applying PCA to the normalized interaction matrices at a 100 Kb resolution using Juicer (Durand, et al., 2016). PCA1 was used to assign A and B compartments. To verify the reproducibility of the compartment call, the PCA analysis was applied on the separate replicates and just the merged data was used for downstream analysis. A custom script and publicly available ChIP-seq BAM files for H3K4me3 (Koike et al., 2012) were used to set the sign to the compartments identified by Juicer. A total of ∼20,000 compartments were identified at each time point. We identified significantly changing compartments as those genomic regions with a change in PCA1 across different time points consistently in the three biological replicates through a one-way ANOVA test.

##### Transcription in A and B compartments

To relate Compartment A and B with transcription we calculated the log2 RPM (Reads Per Million) values for all regions assigned to Compartment A and B per time point (ZT0,6,12,18) using SeqMonk and RNA-seq BAM files (see *RNA-seq analysis*) per time point as input and to applied a Kluskal Wallis test for all compartments and a Mann-Whitney test for OCCs. The distribution of log2 RPM values per compartment type at each time point is presented as Violin Plots.

##### Correlation with histone marks

To relate changes in compartment status with the enrichment of histone post-translational modifications we calculated the RPM (Reads Per Million) values for all regions assigned to Compartment A and B per time point (ZT0,6,12,18) using SeqMonk and publicly available ChIP-seq datasets for the histone post-translational modifications H3K4me3 and H3K4me1 (Yue et al., 2014) per time point as input and applied a one way ANOVA test and a Tukey post hoc test.

##### Correlation with HDAC3

To relate changes in compartment status with the enrichment of HDAC3 we calculated the log2 RPM (Reads Per Million) values for all regions assigned to Compartment A at ZT0 and that change to Compartment B at Z12 using SeqMonk and publicly available ChIP-seq datasets for the histone deacetylase HDAC3 at ZT0 and ZT12 (Feng et al., 2011) and applied a Wilcoxon test.

#### Promoter CHi-C

The sequenced reads were processed using HiCUP (Wingett et al., 2015). The filtering and identification of significant interactions were performed with CHiCAGO (Cairns et al., 2016). To identify differential interactions the script implemented by (Rubin et al., 2017) was used. This script can identify the differential interactions from a Promoter Capture Hi-C dataset, using the edgeR (McCarthy et al., 2012) package to statistically quantify changes in reads for the interactions. To increase the confidence in dynamic interactions, we filtered the dataset only including baits overlapping circadian genes. To account for the distance bias in the read count, we also divided the CHi-C interactions into greater or less than 150 Kb groups. These preliminary results were filtered by FDR and fold change; both distance regimes were combined. To plot long-range interactions we used the Washington Epigenome Browser (http://epigenomegateway.wustl.edu/browser/) using the mouse genome version mm9 and as input properly formatted CHiCAGO output files.

##### Characterization of interacting regions

To characterize the type of genomic element that promoters contact derived from our Promoter CHi-C we calculated the observed/expected number of overlaps between the otherends (the genomic segment interacting with a promoter) and a set of genomic regions occupied by Transcription Factors or enriched for histone post-translational modifications using a custom python script. The expected number of overlaps is calculated by generating a random distribution of otherends CHi-C fragments considering two conditions: 1) the length of the otherend-fragment observed in the original dataset; 2) the distance between the baits (promoter) and otherends. With this random set, we repeat the overlap with the features of interest and keep the number of “expected” overlaps by chance and repeat the process at least 100 times. To plot the results the mean was calculated for all the iterations of the expected values. The significant differences were calculated using a t-test between the distribution of the expected values and the observed number of overlaps.

##### Interactions per promoter

To calculate the number of interactions per promoter at each time point python custom scripts were used. Circadian promoters were defined as genes with circadian transcription as identified by our RNA-seq analysis. For core clock promoter interaction analysis, we used the classic core clock list described in (Anafi et al., 2014). A list of enhancers with circadian transcription (eRNAs) was used derived from (Fang et al., 2014). The comparison with non-core-clock-circadian genes includes all the other circadian genes determined from our RNA-seq analysis. To calculate the distribution of the expected number of interactions made by non-core-clock-circadian genes we randomly selected 11 genes from the entire set, this procedure was repeated 100 times. The comparison between the observed number of interactions made by the core clock genes and the distribution of expected interactions of non-core-clock-circadian genes was made using a Mann Whitney test.

##### Analysis of interactions between circadian promoters and enhancers producing eRNAs

The enhancer regions with transcribed eRNAs from (Fang et al., 2014) were assigned to the otherends (interacting region) and then the bait (promoter) from our Promoter CHi-C datasets. To calculate the observed/expected number of interactions between promoters and enhancers with oscillatory eRNAs (osceRNAs) and non-oscillatory eRNAs (nonosceRNAs) that map to the CHi-C dataset, the expected number was calculated by taking the same number of osceRNAs from the nonosceRNAs set and count the number of interactions between the restriction fragment containing an enhancer transcribed into eRNAs and the fragment containing circadian promoters derived from the CHi-C dataset. This process is repeated at least 100 times. Each enhancer region with transcribed eRNAs was assigned with a phase (maximal transcription during a circadian cycle) (Fang et al., 2014) as well as the promoter of the CHi-C datasets using our RNA-seq analysis. To make eRNAs phases (Fang et al., 2014) more comparable to the circadian promoters identified by our RNA-seq analysis, we grouped them into eight groups each containing three-time points. First, we mapped the osceRNAs to the otherends of the CHi-C, then, retrieved the bait fragment associated with that eRNA and filtered the fragments overlapping with the set of circadian promoters identified by our RNA-seq analysis. Then, we divided the elements in diurnal and nocturnal to have a better understanding of the interaction profiles and perform a Wilcoxon rank sum test per pair of elements for each category.

### Virtual 4C

The output BAM files from HiCUP for the different promoter CHi-C libraries per time point were used as input for SeqMonk to create Virtual-4C plots using promoters of interest as viewpoints to display raw promoter Chi-C counts as for the Arntl1 virtual 4C (Figure S4A). Alternatively the view point of interest and its significant interaction partners were filtered from the CHiCAGO output file and visualized in the Washington Epigenome Browser (http://epigenomegateway.wustl.edu/browser/) using the mouse genome version mm9.

### Construction of promoter-promoter interaction networks

The graphs were constructed using the output generated by CHiCAGO. The raw interaction files for each time point were processed to adjust gene names using *ad hoc* Python scripts due to many transcripts variants presented in those files. Also, a nomenclature was established to represent Ultraconserved Elements (UCEs). Each UCE was represented according to its locus using the following format: uce_[Chromosome number_[Position of first nucleotide]. Each undirected graph was constructed, analyzed and visualized using the NetworkX Python module (Hagberg et al., 2008). The graphs generated were disconnected and contained a large number of small components. For this reason, the components containing 3 or fewer nodes were filtered out for presentation purposes. The statistical analysis used to identify differences in node properties between timepoints was carried out using SciPy (Virtanen et al., 2020) and StatsModels (Seabold & Perktold, 2010) modules. Finally, the ontology enrichment analysis was conducted with the g:Profiler web tool (Raudvere et al., 2019) using the POST request API against the *Mus musculus* genome (*mmusculu*s) with other parameters kept as default. To evaluate the phase coherence between circadian gene promoters contacting each other we assigned the phases of circadian genes with our RNAseq and plotted the phase distribution of circadian promoters contacted by either diurnal or nocturnal circadian promoters. We applied a Wilcoxon rank sum test to calculate the difference between pairs of the two categories.

### ChIP-seq data analysis

Raw sequencing data files for all samples were first processed with FastQC for general quality controls. Sequencing reads were mapped against the mouse genome (NCBIM37/mm9) using Bowtie2 (Langmead & Salzberg, 2012) with default parameters for single and paired reads. Mapped reads were filtered by map quality (-q 30) using samtools (samtools view). Bam files were sorted (samtools sort) and indexed (samtools index). Duplicates were removed with Pickard. Bam files were imported to deeTools v3.3.1 (Ramírez et al., 2016) to create signal tracks with bamCoverage. Signal tracks for all data were visualized using IGV (Thorvaldsdóttir et al., 2013). Peak calling was performed using MACS2 callpeak function with default parameters (Y. Zhang et al., 2008). Peak overlap analysis was performed using the Venn module of Intervene (Khan & Mathelier, 2017).The MEME-ChIP tool (Ma et al., 2014) was used for motif analysis using the fasta sequences from peaks detected by MACS2 with default parameters.

### RNA-seq data processing

Raw sequencing data files for all samples were first processed with FastQC for general quality controls. Sequencing reads were mapped against the mouse genome (NCBIM37/mm9) using TopHat (D. Kim et al., 2013) with default parameters. Deeptools (Ramírez et al., 2016) and BAM files for all samples were used to calculate Spearman’s correlation between all biological replicates for each time point. All samples were highly correlated (Spearman’s correlation >0.85). Heatmaps of correlations were created using the Deeptools plotCorrelation. To create bigWig signal tracks for all time points we merged all biological replicates per time point using samtools (Li et al., 2009). Merged bam files per time point were processed with Deeptools bamCoverage to create strand-specific and RPKM normalized signal tracks suited for comparison. Visualization of signal tracks was done using IGV genome browser (Thorvaldsdóttir et al., 2013). Mapped reads for all samples were then used to assemble and quantify expressed genes and transcripts using StringTie (Pertea et al., 2015) with default parameters. Differential expression was performed by Ballgown (Frazee et al., 2015) in R using StringTie table counts for all samples. Genes with differential expression between at least one pair of time points were identified after correction for multiple hypothesis testing with a q-value <0.01. We classified a gene as circadian if it was differentially expressed between at least a pair of time points. We assigned a phase for each differentially expressed gene to the time point with the highest average FPKM value. Heatmap of circadian genes was created using pheatmap function in R using as input a list of 1256 differentially expressed genes at a q-value <0.01 and ordered by phase. Expression values for all genes were Z-score corrected. Plots of FPKM expression over time for selected examples were generated using library ggplot2 in R.

### Gene Ontology analysis

Gene ontology enrichment analysis and pathway enrichment were done using DAVID (Huang et al., 2009). All significant biological processes and pathways had a *p-value* <0.01. Barplots were generated with ggplot2 in R.

### Motif analysis

The fasta sequence from the HindIII fragments of the *otherend* of stable, dynamic and both types of chromatin contacts was downloaded using the mouse genome version mm9. The MEME-ChIP tool (Ma et al., 2014) was used for motif analysis with default parameters.

### Data availability

Data sets are available upon request. Custom scripts are available without restrictions upon request.

