## Supplementary figures and tables for "The global and promoter-centric 3D genome organization temporally resolved during a circadian cycle"

**Figure legends**

**Figure S1. OCCs present fluctuations in histone modifications.** A. Heatmap of PC1 values for stable compartments from the three independent Hi-C biological replicates or the merged replicates (B). C. PC1 eigenvector correlations between every pair of time points (Kruskal-Wallis rank sum test p-value <2e-16). D. Proportion of OCCs and constant compartments in the mouse genome. E. Barplot of frequency of OCCs categories. F. Chromatin features (H3Kme4 and H3Kme1) at OCCs across time points (AABB, ABBB and ABAB OCCs are shown) (all p-values < 0.001 except for the H3K4me1 AABB, one way ANOVA and Tukey post hoc test).

**Figure S2.** Total RNA-seq **analysis at four time points during the circadian cycle.** A. Heat map of the relative transcription (z scores) of 1,257 oscillating genes sorted by oscillation phase. B. Genome browser tracks showing examples of transcription for *Arntl, Per1, Nr1d1, Cry1, Ppp1r3c* and *Gsk3a* oscillating genes and FPKM quantified at all time points (n=4, q-value <0.01). C. GO analysis for the detected oscillating genes. Significantly enriched categories are shown. D. KEGG analysis for detected oscillating genes. Significantly enriched categories are shown. E. Additional individual transcriptional profiles from the RNA-seq for *Rorc*, *Clock*, *Npas2*, *Cry2*, *Per2*, *Per3* circadian genes. F. RT-PCR validation for candidate circadian genes both at mRNA and pre-mRNA level (p-values < 0.0001, one-way ANOVA test). G. Spearman correlation analysis of the four biological replicates from ZT0 and ZT12 ($r \geq 0.85$).

**Figure S3. TAD structure and CTCF binding is preserved during the 24 hours.** A. TAD size distribution at all time points. B. TADs overlap between time points. C. 50kb resolution Median Observed/Expected Hi-C signal around 1000 randomly selected TADs from ZT6, 12 and 18 plotted on the indicated time points. TADs were scaled to fit the five central bins. D. CTCF ChIP-seq motif analysis and peak overlap between ZT0 and 12. Genomic tracks of CTCF ChIP seq signal at candidate regions. E. The same metaplots as in C but for 1000 randomly CTCF peaks found at ZT12 plotted using ZT12 and ZT0 Hi-C contacts (above). Below, metaplot using CTCF peaks from mESC and plotted using liver ZT0 and 12 Hi-C contacts. CTCF peaks are at the central bin of the metaplot. F. TAD and cTAD sizes (p-value < 2.2e-16, Wilcoxon rank sum test).

**Figure S4. P-CHi-C provides detailed promoter interactomes.** A. Partial view of a virtual 4C from the Arntl1 gene promoter using the Hi-C and P-CHi-C raw valid pairs. Histograms of read counts per restriction fragment around the bait region corresponding to the captured promoter are shown. B. Number of total read counts comparing Hi-C and P-CHi-C recovered using the Arntl1 gene promoter as bait on a virtual 4C (restriction fragments with at least 5 reads in the CHi-C experiment where used) (p-value < 0.0001, Wilcoxon ranked test). C.

Observed and Expected contacts for circadian promoters oscillating at the intronic level or detected trough GRO-seq (Fang et al., 2014), with enhancers producing eRNAs. D. Phase distributions of rhythmic eRNAs producing enhancers contacted by circadian promoters detected through GRO-seq. E. Phase distribution of circadian promoters contacted by other circadian promoters detected through GRO-seq. F. Virtual 4C for *Rorc and Cgn* circadian gene promoters from CHi-C data at all time points during the day. The two promoters with shared transcriptional peak contact each other in the nuclear space. Acrophases are written next to the gene names. Genomic tracks show significant contacts as arcs and chromatin features including liver H3K4me3, H3K4me1, H3K27ac, DNAseI, eRNAs and TADs.

**Figure S5. Promoter-Promoter networks in the mouse liver over a circadian cycle.** A. Promoter-promoter contact network at ZT0. Each color represents a chromosome. Nodes with degree=1 are not included for simplicity. B. Promoter-promoter contact network as in A but with circadian genes marked in blue. C. The four main subcomponents of the graph at the different time points during the day. D. Significantly enriched GO categories for the genes on the most prominent subcomponents in C. E. Above, number of total network edges at ZT0, 6, 12 and 18 hours of the day for all gene promoters and circadian gene promoters (blue line). Below, z-score for circadian gene promoter contacts compared to random sampling. F. Phase distributions for nocturnal and diurnal circadian promoter-promoters contacts (all p-values < 0.0001, Wilcoxon signed rank test). G. Virtual 4C for *Tef* gene promoter from CHi-C data at all time points during the day. Acrophase is written next to the gene name. Genomic tracks show significant contacts as arcs and chromatin features including liver H3K4me3, H4K4me1, H3K27ac, DNAseI, eRNAs and TADs. *Tef* gene promoter contacts *Aco2* gene promoter both with acrophase at ZT12.

**Figure S6. Core clock gene promoter vs output circadian genes interaction landscapes.** A-D. Virtual 4C for *Per2*, *Npas2, Nr1d2,* core clock gene promoters from CHi-C data at all time points during the day. E-F Virtual 4C for *Dhrs3* and *PTG* liver output circadian genes. Acrophases are written next to the gene names. Genomic tracks show significant contacts as arcs and chromatin features including liver H3K4me3, H3K4me1, H3K27ac, DNAseI, eRNAs and TADs. E-F. The interaction profiles of core clock genes display less contacts that dynamically change over time. The two output gene contact profiles show more saturated contacts that are mostly constant during the 24 hours.

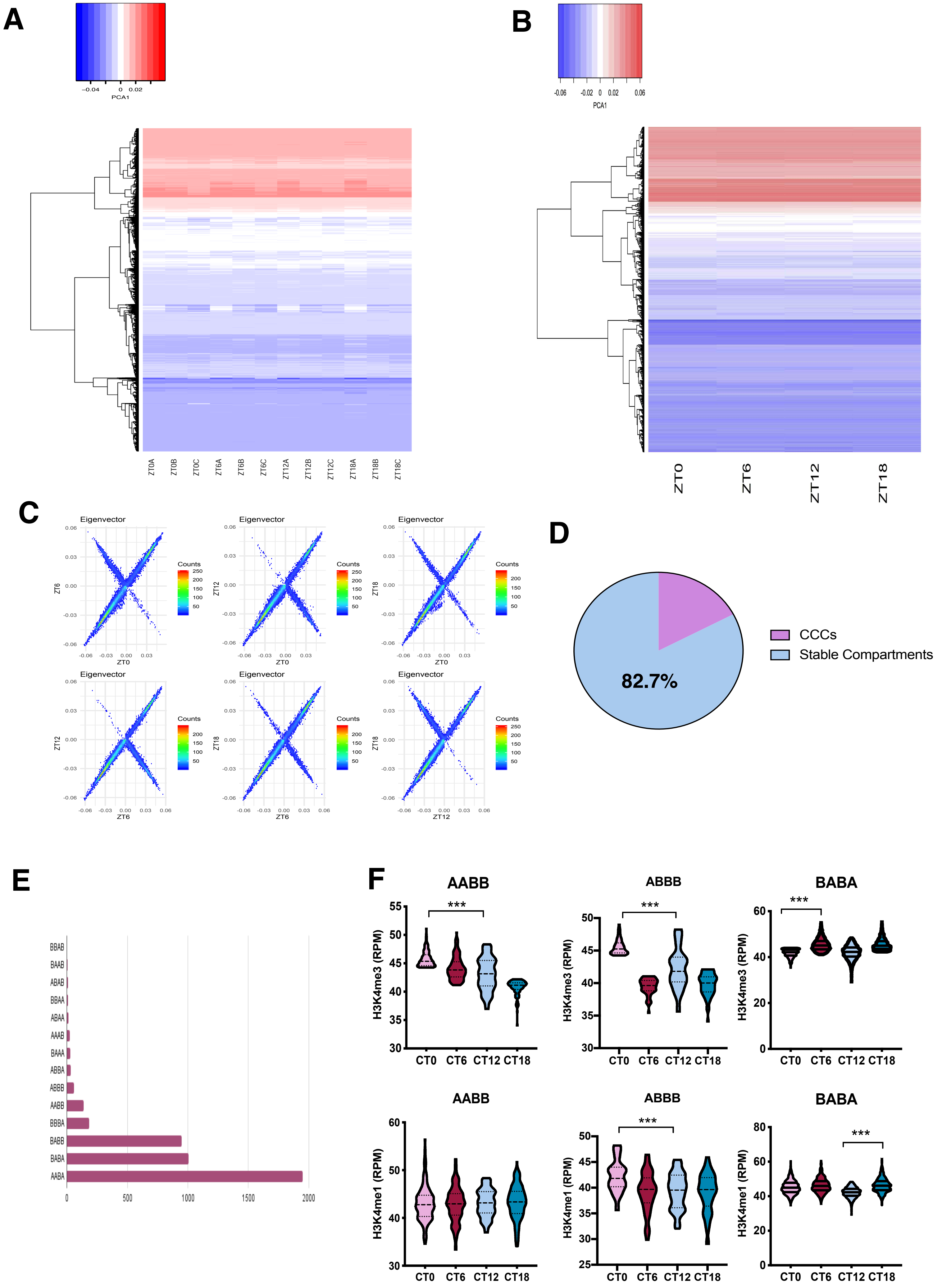

A
PCA1
-0.04
0
0.02
ZT0A
ZT0B
ZT0C
ZT6A
ZT6B
ZT6C
ZT12A
ZT12B
ZT12C
ZT18A
ZT18B
ZT18C
B
PCA1
-0.06
-0.02
0
0.02
0.06
ZT0
ZT6
ZT12
ZT18
C
Eigenvector
Counts
ZT0
ZT6
ZT12
ZT18
D
CCCs
Stable Compartments
82.7%
E
BBAB
BAAB
ABAB
BBAA
ABAA
AAAB
BAAA
ABBA
ABBB
AABB
BBBA
BABB
BABA
AABA
0
500
1000
1500
2000
F
AABB
ABBB
BABA
H3K4me3 (RPM)
H3K4me1 (RPM)
CT0
CT6
CT12
CT18
***

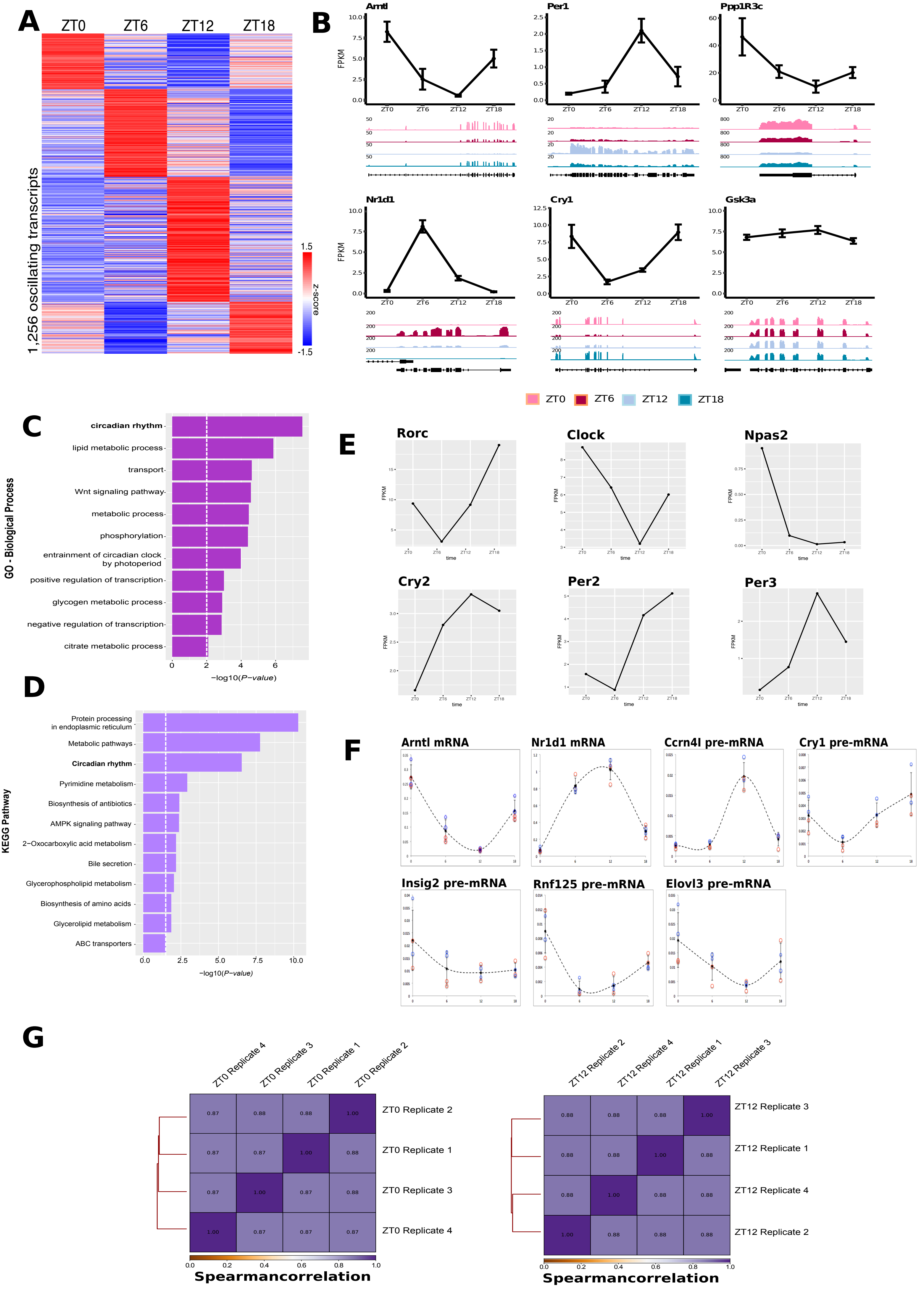
A
ZT0
ZT6
ZT12
ZT18
1,256 oscillating transcripts
Z-score
B
Arntl
Per1
Ppp1R3c
Nr1d1
Cry1
Gsk3a
FPKM
ZT0
ZT6
ZT12
ZT18
C
GO - Biological Process
circadian rhythm
lipid metabolic process
transport
Wnt signaling pathway
metabolic process
phosphorylation
entrainment of circadian clock by photoperiod
positive regulation of transcription
glycogen metabolic process
negative regulation of transcription
citrate metabolic process
−log10(P−value)
D
KEGG Pathway
Protein processing in endoplasmic reticulum
Metabolic pathways
Circadian rhythm
Pyrimidine metabolism
Biosynthesis of antibiotics
AMPK signaling pathway
2−Oxocarboxylic acid metabolism
Bile secretion
Glycerophospholipid metabolism
Biosynthesis of amino acids
Glycerolipid metabolism
ABC transporters
−log10(P−value)
E
Rorc
Clock
Npas2
Cry2
Per2
Per3
time
F
Arntl mRNA
Nr1d1 mRNA
Ccrn4l pre-mRNA
Cry1 pre-mRNA
Insig2 pre-mRNA
Rnf125 pre-mRNA
Elovl3 pre-mRNA
G
ZT0 Replicate 4
ZT0 Replicate 3
ZT0 Replicate 1
ZT0 Replicate 2
ZT12 Replicate 2
ZT12 Replicate 4
ZT12 Replicate 1
ZT12 Replicate 3
Spearmancorrelation
Spearmancorrelation

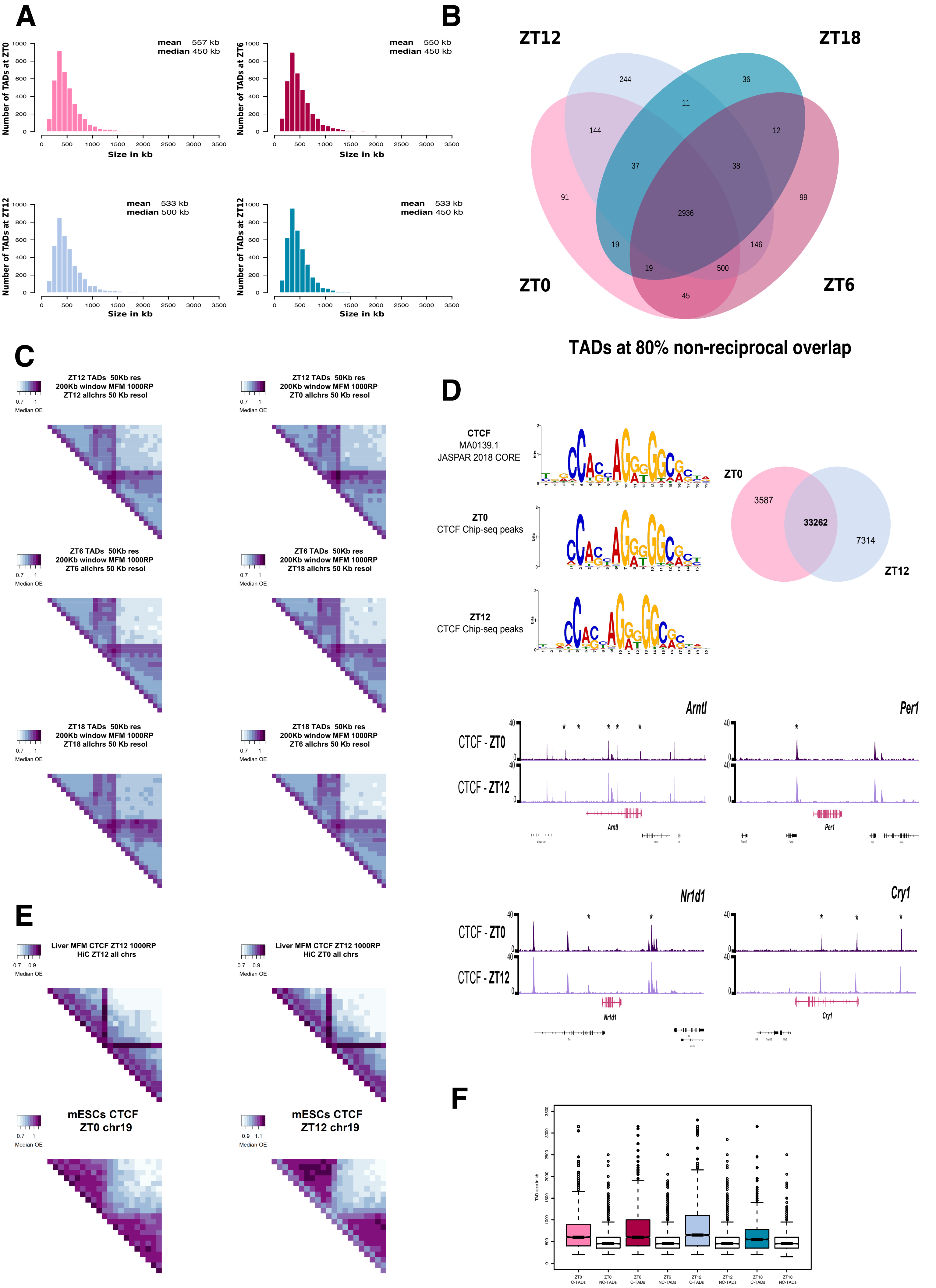
A
mean 557 kb
median 450 kb
Number of TADs at ZT0
Size in kb
mean 550 kb
median 450 kb
Number of TADs at ZT6
mean 533 kb
median 500 kb
Number of TADs at ZT12
mean 533 kb
median 450 kb
B
ZT12
ZT18
ZT0
ZT6
244
36
11
144
12
37
38
91
2936
99
19
146
19
500
45
TADs at 80% non-reciprocal overlap
C
ZT12 TADs 50Kb res
200Kb window MFM 1000RP
ZT12 allchrs 50 Kb resol
ZT12 TADs 50Kb res
200Kb window MFM 1000RP
ZT0 allchrs 50 Kb resol
ZT6 TADs 50Kb res
200Kb window MFM 1000RP
ZT6 allchrs 50 Kb resol
ZT6 TADs 50Kb res
200Kb window MFM 1000RP
ZT18 allchrs 50 Kb resol
ZT18 TADs 50Kb res
200Kb window MFM 1000RP
ZT18 allchrs 50 Kb resol
ZT18 TADs 50Kb res
200Kb window MFM 1000RP
ZT6 allchrs 50 Kb resol
Median OE
D
CTCF
MA0139.1
JASPAR 2018 CORE
ZT0
CTCF Chip-seq peaks
ZT12
CTCF Chip-seq peaks
ZT0
3587
33262
7314
ZT12
Arntl
Per1
CTCF - ZT0
CTCF - ZT12
Nr1d1
Cry1
E
Liver MFM CTCF ZT12 1000RP
HiC ZT12 all chrs
Liver MFM CTCF ZT12 1000RP
HiC ZT0 all chrs
mESCs CTCF
ZT0 chr19
mESCs CTCF
ZT12 chr19
F
TAD size in kb
ZT0 C-TADs
ZT0 NC-TADs
ZT6 C-TADs
ZT6 NC-TADs
ZT12 C-TADs
ZT12 NC-TADs
ZT18 C-TADs
ZT18 NC-TADs

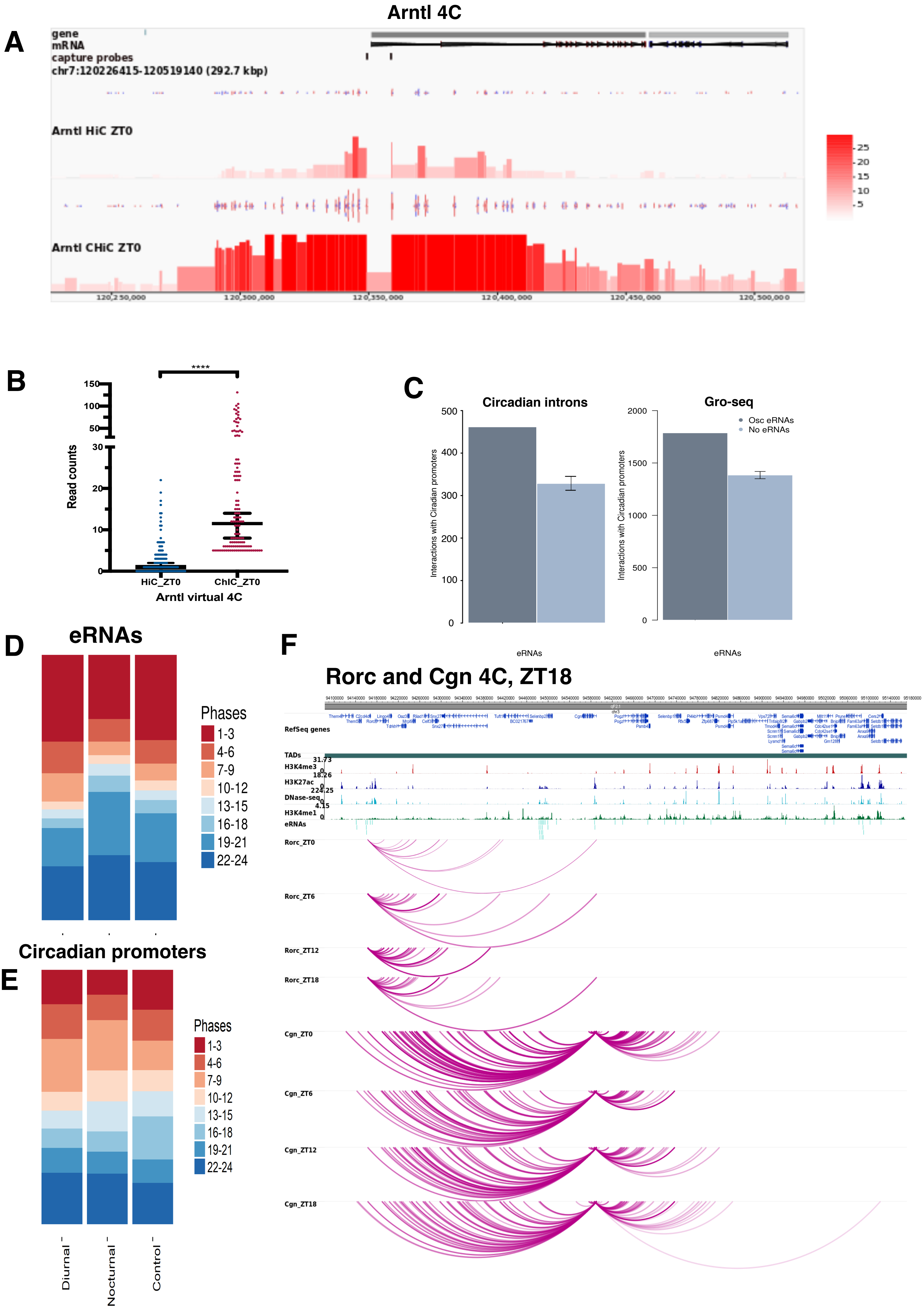

A
Arntl 4C
gene
mRNA
capture probes
chr7:120226415-120519140 (292.7 kbp)
Arntl HiC ZT0
Arntl CHiC ZT0
25
20
15
10
5
120,250,000
120,300,000
120,350,000
120,400,000
120,450,000
120,500,000
B
****
Read counts
150
100
50
30
20
10
0
HiC_ZT0
ChIC_ZT0
Arntl virtual 4C
C
Circadian introns
Gro-seq
Interactions with Ciradian promoters
Interactions with Circadian promoters
500
400
300
200
100
0
2000
1500
1000
500
0
Osc eRNAs
No eRNAs
eRNAs
eRNAs
D
eRNAs
Phases
1-3
4-6
7-9
10-12
13-15
16-18
19-21
22-24
Circadian promoters
E
Phases
1-3
4-6
7-9
10-12
13-15
16-18
19-21
22-24
Diurnal
Nocturnal
Control
F
Rorc and Cgn 4C, ZT18
RefSeq genes
TADs
H3K4me3
H3K27ac
DNase-seq
H3K4me1
eRNAs
31.73
18.26
224.25
4.15
Rorc_ZT0
Rorc_ZT6
Rorc_ZT12
Rorc_ZT18
Cgn_ZT0
Cgn_ZT6
Cgn_ZT12
Cgn_ZT18

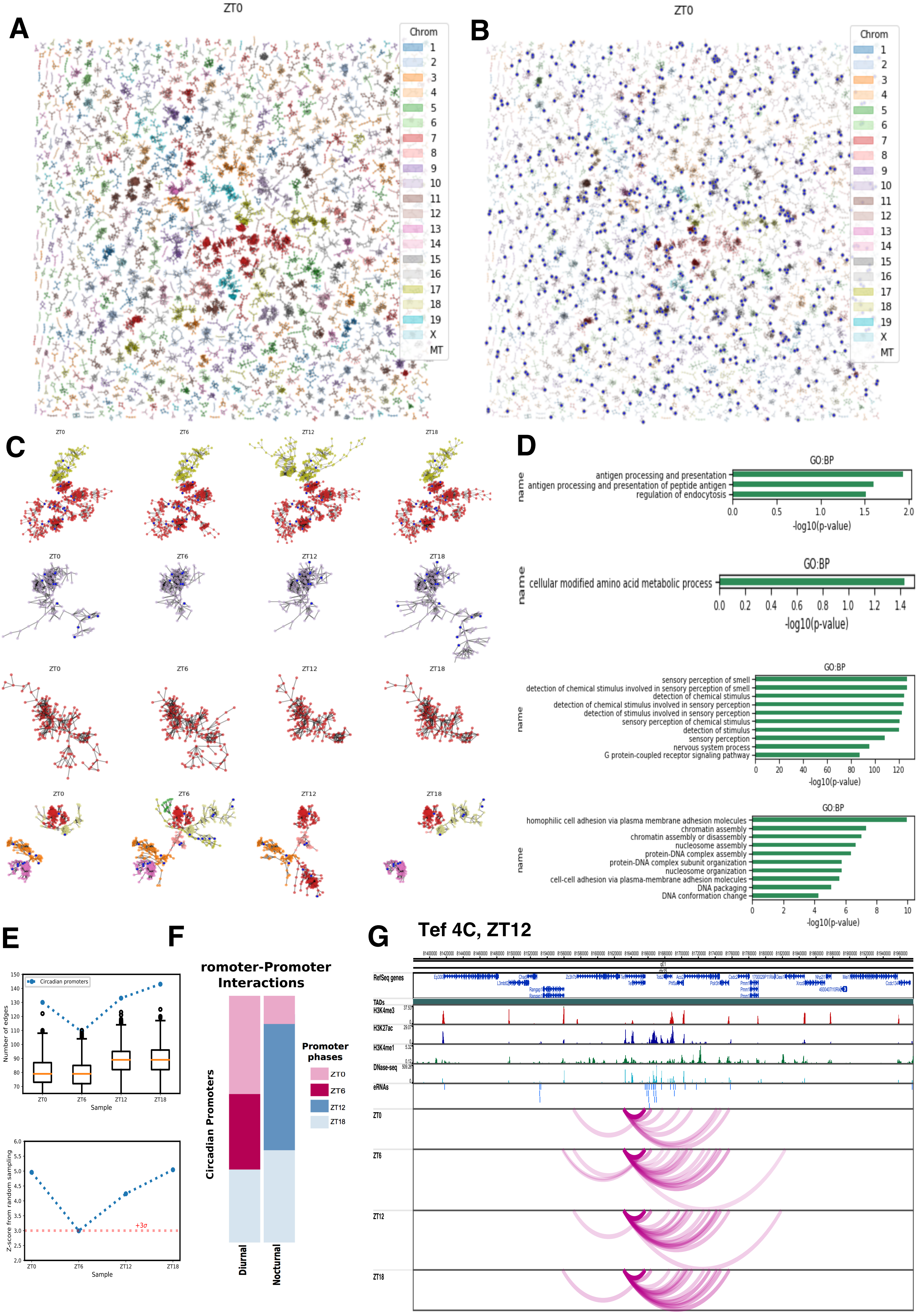

A
ZT0
Chrom
X
MT
B
ZT0
C
ZT0
ZT6
ZT12
ZT18
D
GO:BP
antigen processing and presentation
antigen processing and presentation of peptide antigen
regulation of endocytosis
-log10(p-value)
cellular modified amino acid metabolic process
sensory perception of smell
detection of chemical stimulus involved in sensory perception of smell
detection of chemical stimulus
detection of chemical stimulus involved in sensory perception
detection of stimulus involved in sensory perception
sensory perception of chemical stimulus
detection of stimulus
sensory perception
nervous system process
G protein-coupled receptor signaling pathway
homophilic cell adhesion via plasma membrane adhesion molecules
chromatin assembly
chromatin assembly or disassembly
nucleosome assembly
protein-DNA complex assembly
protein-DNA complex subunit organization
nucleosome organization
cell-cell adhesion via plasma-membrane adhesion molecules
DNA packaging
DNA conformation change
name
E
Circadian promoters
Number of edges
Sample
Z-score from random sampling
+3σ
F
romoter-Promoter Interactions
Circadian Promoters
Promoter phases
ZT0
ZT6
ZT12
ZT18
Diurnal
Nocturnal
G
Tef 4C, ZT12
RefSeq genes
TADs
H3K4me3
H3K27ac
H3K4me1
DNase-seq
eRNAs
ZT0
ZT6
ZT12
ZT18

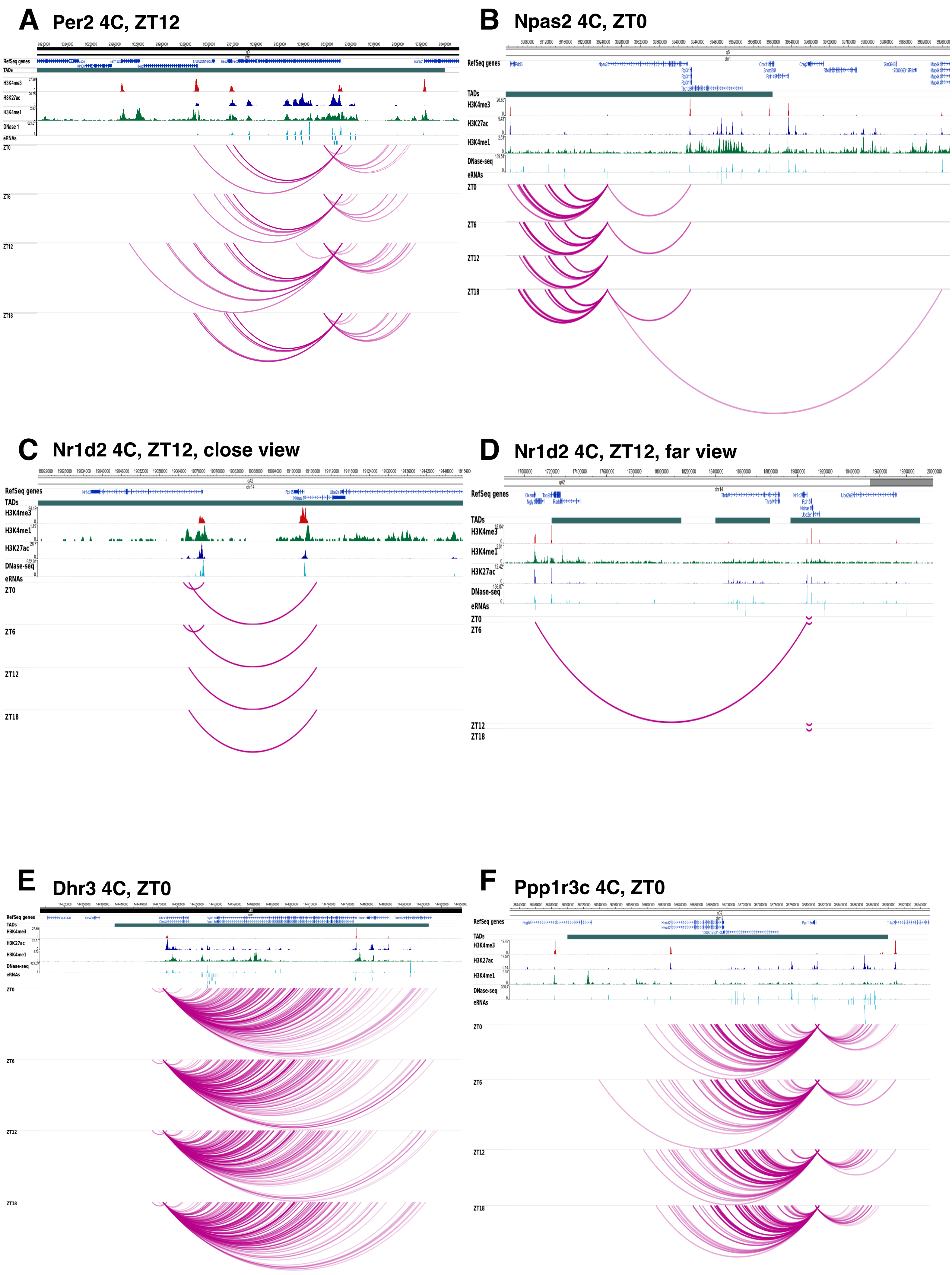
A
Per2 4C, ZT12
RefSeq genes
TADs
H3K4me3
H3K27ac
H3K4me1
DNase 1
eRNAs
ZT0
ZT6
ZT12
ZT18
B
Npas2 4C, ZT0
RefSeq genes
TADs
H3K4me3
H3K27ac
H3K4me1
DNase-seq
eRNAs
ZT0
ZT6
ZT12
ZT18
C
Nr1d2 4C, ZT12, close view
RefSeq genes
TADs
H3K4me3
H3K4me1
H3K27ac
DNase-seq
eRNAs
ZT0
ZT6
ZT12
ZT18
D
Nr1d2 4C, ZT12, far view
RefSeq genes
TADs
H3K4me3
H3K4me1
H3K27ac
DNase-seq
eRNAs
ZT0
ZT6
ZT12
ZT18
E
Dhr3 4C, ZT0
RefSeq genes
TADs
H3K4me3
H3K27ac
H3K4me1
DNase-seq
eRNAs
ZT0
ZT6
ZT12
ZT18
F
Ppp1r3c 4C, ZT0
RefSeq genes
TADs
H3K4me3
H3K27ac
H3K4me1
DNase-seq
eRNAs
ZT0
ZT6
ZT12
ZT18

#### Supplementary tables

**Table S1. HiC statistics.** HiCUP summary results for the independent HiC replicates.

| Sample | Total number of read pairs | Unique paired alignments | Total number of valid read pairs | % Valid pairs | % Unique valid pairs | % cis reads | % trans reads |
|---|---|---|---|---|---|---|---|
| HiC ZT0 Pool A | 421,930,174 | 274,526,514 | 138,334,919 | 50.4 | 99.25 | 84 | 16 |
| HiC ZT0 Pool B | 218,523,760 | 147,339,543 | 123,675,798 | 84 | 99.2 | 84.2 | 15.8 |
| HiC ZT0 Pool C | 266,336,305 | 165,258,505 | 135,220,903 | 81.8 | 98.9 | 81.3 | 18.8 |
| HiC ZT6 Pool A | 413,884,648 | 269,112,742 | 114,851,884 | 42.7 | 99.4 | 82.5 | 17.4 |
| HiC ZT6 Pool B | 197,750,025 | 131,191,846 | 109,998,946 | 83.8 | 99.2 | 83.2 | 16.8 |
| HiC ZT6 Pool C | 277,055,047 | 170,871,915 | 152,207,672 | 89.1 | 99 | 82.1 | 17.9 |
| HiC ZT12 Pool A | 443,824,919 | 294,318,745 | 108,372,640 | 36.8 | 99.4 | 83.3 | 16.7 |
| HiC ZT12 Pool B | 215,571,675 | 144,746,635 | 125,558,474 | 86.7 | 99.2 | 83.7 | 16.3 |
| HiC ZT12 Pool C | 271,236,635 | 165,859,847 | 145,192,886 | 87.5 | 98.9 | 81.6 | 18.4 |
| HiC ZT18 Pool A | 467,586,439 | 306,126,645 | 126,673,109 | 41.7 | 99.5 | 83.2 | 16.9 |
| HiC ZT18 Pool B | 224,661,794 | 151,845,907 | 134,589,293 | 88.6 | 99.2 | 83.3 | 16.7 |
| HiC ZT18 Pool C | 287,472,226 | 176,450,565 | 155,424,860 | 88.1 | 98.6 | 81 | 18.9 |

Table S2. **Total stranded RNAseq number of unique read pairs**

| Sample | Total unique 150bp read pairs |
|---|---|
| ZT0_replicate_1 | 125,163,820 |
| ZT0_replicate_2 | 107,586,456 |
| ZT0_replicate_3 | 133,618,265 |
| ZT0_replicate_4 | 112,081,774 |
| ZT6_replicate_1 | 154,897,216 |
| ZT6_replicate_2 | 88,379,045 |
| ZT6_replicate_3 | 167,923,190 |
| ZT6_replicate_4 | 131,189,401 |
| ZT12_replicate_1 | 171,532,500 |
| ZT12_replicate_2 | 136,197,578 |
| ZT12_replicate_3 | 183,736,327 |
| ZT12_replicate_4 | 100,236,131 |
| ZT18_replicate_1 | 93,867,568 |
| ZT18_replicate_2 | 143,856,427 |
| ZT18_replicate_3 | 138,638,802 |
| ZT18_replicate_4 | 108,275,521 |

Table S3. **Circadian genes detected from total RNAseq.**

| gene_id | gene_name | chr | start | end | strand | B_pvalue | B_qvalue | FPKM_ZT0 | FPKM_ZT6 | FPKM_ZT12 | FPKM_ZT18 | Max FPKM | ZT_Max_FPKM | Koike2012_exon | Koike2012_intron |
|---|---|---|---|---|---|---|---|---|---|---|---|---|---|---|---|
| ENSMUSG00000026077 | Npas2 | 1 | 39250576 | 39420079 | + | 3.12E-11 | 3.96E-07 | 0.95186357 | 0.09751367 | 0.01401603 | 0.03311566 | 0.95186357 | ZT0 | Present_Koike2012 | Present_Koike2012 |
| ENSMUSG00000034917 | Tjp3 | 10 | 80735952 | 80754012 | - | 1.56E-10 | 7.92E-07 | 10.061522 | 5.19245224 | 2.35376323 | 7.18746534 | 10.061522 | ZT0 | Present_Koike2012 | Present_Koike2012 |
| ENSMUSG00000038754 | Elovl3 | 19 | 46206387 | 46210184 | + | 3.68E-09 | 7.81E-06 | 307.024413 | 84.219419 | 14.6589042 | 88.0869945 | 307.024413 | ZT0 | Present_Koike2012 | Present_Koike2012 |
| ENSMUSG00000058672 | Tubb2a | 13 | 34166170 | 34169877 | - | 1.23E-08 | 1.84E-05 | 81.5520268 | 26.2687995 | 11.8542283 | 41.5215913 | 81.5520268 | ZT0 | Present_Koike2012 | Present_Koike2012 |
| ENSMUSG00000002250 | Ppard | 17 | 28339699 | 28438414 | + | 1.56E-08 | 2.20E-05 | 1.29052408 | 0.20472718 | 0.22981426 | 0.34584421 | 1.29052408 | ZT0 | Present_Koike2012 | Present_Koike2012 |
| ENSMUSG00000003721 | Insig2 | 1 | 123200930 | 123229166 | - | 4.50E-08 | 3.95E-05 | 46.2658036 | 31.4196732 | 9.74865394 | 21.9382875 | 46.2658036 | ZT0 | Present_Koike2012 | Present_Koike2012 |
| ENSMUSG00000021871 | Pnp | 14 | 51563977 | 51573087 | + | 5.20E-08 | 4.38E-05 | 21.7649202 | 15.8889106 | 10.4577973 | 15.7904487 | 21.7649202 | ZT0 | Present_Koike2012 | Absent_Koike2012 |
| ENSMUSG00000042363 | 1110067D22 | 11 | 20723579 | 20731059 | - | 7.03E-08 | 4.97E-05 | 8.11479221 | 2.79616217 | 1.70890543 | 4.9646021 | 8.11479221 | ZT0 | Present_Koike2012 | Absent_Koike2012 |
| ENSMUSG00000015357 | Clpx | 9 | 65142067 | 65178465 | + | 7.59E-08 | 5.08E-05 | 58.8832347 | 33.578119 | 12.7716998 | 25.1968144 | 58.8832347 | ZT0 | Present_Koike2012 | Present_Koike2012 |
| ENSMUSG00000091780 | Sco2 | 15 | 89202113 | 89202880 | - | 7.42E-08 | 5.08E-05 | 16.3311958 | 14.8847304 | 5.80182402 | 6.8831432 | 16.3311958 | ZT0 | Absent_Koike2012 | Absent_Koike2012 |
| ENSMUSG00000045136 | Tubb2b | 13 | 34218877 | 34222223 | - | 8.95E-08 | 5.69E-05 | 10.5208668 | 3.16785075 | 1.5474625 | 5.06874325 | 10.5208668 | ZT0 | Absent_Koike2012 | Absent_Koike2012 |
| ENSMUSG00000066026 | Dhrs3 | 4 | 144482730 | 144518112 | + | 1.72E-07 | 8.01E-05 | 25.4383862 | 21.7177849 | 11.8953316 | 17.7445933 | 25.4383862 | ZT0 | Present_Koike2012 | Absent_Koike2012 |
| ENSMUSG00000025504 | Eps8l2 | 7 | 148524779 | 148549819 | + | 1.54E-07 | 8.01E-05 | 9.18786911 | 9.09890706 | 3.09478326 | 6.03765677 | 9.18786911 | ZT0 | Present_Koike2012 | Present_Koike2012 |
| ENSMUSG00000025229 | Pitx3 | 19 | 46210175 | 46222865 | - | 1.82E-07 | 8.14E-05 | 2.23195346 | 0.36445203 | 0.05181715 | 0.69421886 | 2.23195346 | ZT0 | Absent_Koike2012 | Absent_Koike2012 |
| ENSMUSG00000039395 | Mreg | 1 | 72205807 | 72258881 | - | 2.03E-07 | 8.32E-05 | 23.54987 | 17.2465685 | 9.28149125 | 17.188919 | 23.54987 | ZT0 | Present_Koike2012 | Present_Koike2012 |
| ENSMUSG00000002957 | Ap2a2 | 7 | 148748079 | 148818910 | + | 2.23E-07 | 8.42E-05 | 24.461143 | 20.1702038 | 17.01984 | 22.0223898 | 24.461143 | ZT0 | Present_Koike2012 | Absent_Koike2012 |
| ENSMUSG00000020123 | Avpr1a | 10 | 121885555 | 121890509 | + | 2.16E-07 | 8.42E-05 | 11.7198363 | 7.07136425 | 2.98471225 | 6.51900825 | 11.7198363 | ZT0 | Present_Koike2012 | Absent_Koike2012 |
| ENSMUSG00000038366 | Lasp1 | 11 | 97660314 | 97700078 | + | 2.20E-07 | 8.42E-05 | 14.2953969 | 12.4783937 | 7.43623596 | 12.965683 | 14.2953969 | ZT0 | Absent_Koike2012 | Absent_Koike2012 |
| ENSMUSG00000024913 | Lrp5 | 19 | 3584828 | 3686564 | - | 2.94E-07 | 9.83E-05 | 6.95280543 | 5.91584915 | 3.74165342 | 4.56685856 | 6.95280543 | ZT0 | Absent_Koike2012 | Present_Koike2012 |
| ENSMUSG00000026489 | Adck3 | 1 | 182095369 | 182129733 | - | 3.21E-07 | 0.00010344 | 34.3012544 | 27.8843378 | 10.8638568 | 26.4293929 | 34.3012544 | ZT0 | Present_Koike2012 | Present_Koike2012 |
| ENSMUSG00000054814 | Usp46 | 5 | 74394487 | 74464455 | - | 3.85E-07 | 0.00011155 | 3.28139579 | 2.22344085 | 2.83872361 | 3.05263994 | 3.28139579 | ZT0 | Absent_Koike2012 | Absent_Koike2012 |
| ENSMUSG00000037379 | Spon2 | 5 | 33556167 | 33560887 | - | 4.07E-07 | 0.0001138 | 3.90750115 | 2.2945821 | 0.68846032 | 1.53851463 | 3.90750115 | ZT0 | Present_Koike2012 | Present_Koike2012 |
| ENSMUSG00000026179 | Pnkd | 1 | 74331504 | 74400268 | + | 3.99E-07 | 0.0001138 | 9.68778729 | 5.9482753 | 3.69291525 | 7.48250118 | 9.68778729 | ZT0 | Present_Koike2012 | Absent_Koike2012 |
| ENSMUSG00000036078 | Sigmar1 | 4 | 41685366 | 41703030 | - | 4.32E-07 | 0.00011704 | 12.4483434 | 11.761291 | 6.8716455 | 11.7671674 | 12.4483434 | ZT0 | Present_Koike2012 | Absent_Koike2012 |
| ENSMUSG00000027184 | Caprin1 | 2 | 103603098 | 103637806 | - | 4.96E-07 | 0.00012943 | 23.1236556 | 16.4092627 | 11.640241 | 18.1066484 | 23.1236556 | ZT0 | Present_Koike2012 | Present_Koike2012 |
| ENSMUSG00000064254 | Ethe1 | 7 | 25372562 | 25393944 | + | 5.62E-07 | 0.00013341 | 31.8978495 | 22.8506343 | 11.414395 | 17.872829 | 31.8978495 | ZT0 | Present_Koike2012 | Present_Koike2012 |
| ENSMUSG00000031147 | Magix | X | 7250290 | 7258377 | - | 5.63E-07 | 0.00013341 | 3.03426442 | 2.96218422 | 1.99893213 | 2.56995074 | 3.03426442 | ZT0 | Absent_Koike2012 | Absent_Koike2012 |
| ENSMUSG00000090877 | Hspa1b | 17 | 35093381 | 35096183 | - | 5.55E-07 | 0.00013341 | 9.85178087 | 1.14284878 | 3.20906372 | 4.58821951 | 9.85178087 | ZT0 | Absent_Koike2012 | Absent_Koike2012 |
| ENSMUSG00000010609 | Psen2 | 1 | 182157135 | 182193569 | - | 6.91E-07 | 0.00015034 | 28.8768927 | 21.9385219 | 12.9399937 | 24.8971015 | 28.8768927 | ZT0 | Present_Koike2012 | Present_Koike2012 |
| ENSMUSG00000039620 | 6430573F11I | 8 | 37520602 | 37577374 | + | 7.03E-07 | 0.00015123 | 0.95178431 | 0.6729576 | 0.24749086 | 0.40213544 | 0.95178431 | ZT0 | Present_Koike2012 | Present_Koike2012 |
| ENSMUSG00000005312 | Ubqln1 | 13 | 58277524 | 58316997 | - | 7.19E-07 | 0.00015123 | 36.5711357 | 30.222391 | 26.7520298 | 32.1032822 | 36.5711357 | ZT0 | Absent_Koike2012 | Present_Koike2012 |
| ENSMUSG00000021607 | Mrpl36 | 13 | 73468457 | 73469623 | + | 7.23E-07 | 0.00015123 | 18.9433228 | 14.2714808 | 15.0163408 | 17.3772103 | 18.9433228 | ZT0 | Present_Koike2012 | Absent_Koike2012 |
| ENSMUSG00000053559 | Smagp | 15 | 100451773 | 100467296 | - | 9.17E-07 | 0.00017284 | 10.746439 | 8.37894714 | 4.49804962 | 5.11637624 | 10.746439 | ZT0 | Present_Koike2012 | Absent_Koike2012 |
| ENSMUSG00000028327 | 1300002K09I | 4 | 45861536 | 45899880 | + | 1.10E-06 | 0.00019425 | 10.9361511 | 9.50922925 | 5.57961166 | 6.78209417 | 10.9361511 | ZT0 | Present_Koike2012 | Present_Koike2012 |
| ENSMUSG00000039982 | Dtx4 | 19 | 12540831 | 12576486 | - | 1.18E-06 | 0.00019741 | 2.97216581 | 2.08988485 | 0.65811408 | 1.92399266 | 2.97216581 | ZT0 | Present_Koike2012 | Present_Koike2012 |
| ENSMUSG00000027579 | Srms | 2 | 180940267 | 180947890 | - | 1.19E-06 | 0.00019773 | 0.58049301 | 0.23476977 | 0.14798475 | 0.56697256 | 0.58049301 | ZT0 | Absent_Koike2012 | Absent_Koike2012 |
| ENSMUSG00000086103 | Gm11832 | 4 | 11538286 | 11542154 | - | 1.30E-06 | 0.0002057 | 1.10891875 | 0.66316875 | 0.043526 | 0.2709865 | 1.10891875 | ZT0 | Absent_Koike2012 | Absent_Koike2012 |
| ENSMUSG00000045092 | S1pr1 | 3 | 115413351 | 115417973 | - | 1.30E-06 | 0.0002057 | 22.2282049 | 13.0450044 | 11.2053812 | 18.7727249 | 22.2282049 | ZT0 | Present_Koike2012 | Absent_Koike2012 |
| ENSMUSG00000029817 | Tra2a | 6 | 49193920 | 49214051 | - | 1.35E-06 | 0.00021037 | 4.7416364 | 1.98910568 | 1.93867112 | 4.02424768 | 4.7416364 | ZT0 | Absent_Koike2012 | Absent_Koike2012 |
| ENSMUSG00000055116 | Arntl | 7 | 120350979 | 120457636 | + | 1.42E-06 | 0.00021852 | 8.26126531 | 2.53373778 | 0.55343384 | 5.02131242 | 8.26126531 | ZT0 | Present_Koike2012 | Present_Koike2012 |
| ENSMUSG00000036957 | Lrfn3 | 7 | 31140533 | 31147791 | - | 1.59E-06 | 0.0002381 | 2.953302 | 1.656391 | 0.68498375 | 2.2586295 | 2.953302 | ZT0 | Absent_Koike2012 | Present_Koike2012 |
| ENSMUSG00000023147 | Wrb | 16 | 96367014 | 96379459 | + | 1.81E-06 | 0.00025798 | 1.07835336 | 0.69547473 | 0.86760473 | 1.00071125 | 1.07835336 | ZT0 | Absent_Koike2012 | Absent_Koike2012 |
| ENSMUSG00000039956 | Mrap | 16 | 90738569 | 90750030 | + | 2.03E-06 | 0.0002812 | 21.8758003 | 12.3402379 | 12.706166 | 21.105728 | 21.8758003 | ZT0 | Present_Koike2012 | Present_Koike2012 |
| ENSMUSG00000022550 | Adck5 | 15 | 76406788 | 76426246 | + | 2.40E-06 | 0.00031817 | 5.35314955 | 5.28266085 | 2.95088065 | 3.3682726 | 5.35314955 | ZT0 | Present_Koike2012 | Present_Koike2012 |
| ENSMUSG00000001983 | Taco1 | 11 | 105927375 | 105934926 | + | 2.56E-06 | 0.00033609 | 9.3575505 | 8.75728975 | 6.8462895 | 8.58429225 | 9.3575505 | ZT0 | Absent_Koike2012 | Absent_Koike2012 |
| ENSMUSG00000033147 | Slc22a15 | 3 | 101659701 | 101728376 | - | 2.63E-06 | 0.00034158 | 0.82711518 | 0.80390551 | 0.51994723 | 0.71293528 | 0.82711518 | ZT0 | Absent_Koike2012 | Absent_Koike2012 |
| ENSMUSG00000027478 | Dnmt3b | 2 | 153475186 | 153513466 | + | 2.78E-06 | 0.0003535 | 0.93299587 | 0.33893731 | 0.17377503 | 0.51136162 | 0.93299587 | ZT0 | Absent_Koike2012 | Present_Koike2012 |
| ENSMUSG00000032249 | Anp32a | 9 | 62189100 | 62226619 | + | 2.99E-06 | 0.00037095 | 13.2272783 | 12.8202687 | 7.7783849 | 12.0725076 | 13.2272783 | ZT0 | Present_Koike2012 | Absent_Koike2012 |
| ENSMUSG00000038244 | Mical2 | 7 | 119369370 | 119498708 | + | 3.14E-06 | 0.00038752 | 2.35460439 | 1.22237638 | 0.91171015 | 1.91062076 | 2.35460439 | ZT0 | Absent_Koike2012 | Absent_Koike2012 |
| ENSMUSG00000026614 | Slc30a10 | 1 | 187278727 | 187292641 | + | 3.20E-06 | 0.00039165 | 10.1076559 | 5.27947315 | 2.62448007 | 6.1181391 | 10.1076559 | ZT0 | Present_Koike2012 | Present_Koike2012 |
| ENSMUSG00000019916 | P4ha1 | 10 | 58786044 | 58836052 | + | 3.30E-06 | 0.0003969 | 4.71762693 | 1.59055822 | 2.55217433 | 3.87114383 | 4.71762693 | ZT0 | Absent_Koike2012 | Present_Koike2012 |
| ENSMUSG00000033107 | Rnf125 | 18 | 21103126 | 21142349 | + | 3.57E-06 | 0.00041684 | 74.553585 | 66.7390656 | 25.855586 | 45.2087403 | 74.553585 | ZT0 | Present_Koike2012 | Present_Koike2012 |
| ENSMUSG00000091971 | Hspa1a | 17 | 35106144 | 35109069 | - | 3.87E-06 | 0.00044151 | 3.943756 | 0.418735 | 1.320209 | 1.77006225 | 3.943756 | ZT0 | Absent_Koike2012 | Absent_Koike2012 |
| ENSMUSG00000053286 | 1190005F20I | 1 | 153275778 | 153305312 | + | 4.06E-06 | 0.00045035 | 6.406282 | 3.83913375 | 3.7731985 | 5.16487075 | 6.406282 | ZT0 | Absent_Koike2012 | Present_Koike2012 |
| ENSMUSG00000067787 | Blcap | 2 | 157382098 | 157397010 | - | 4.03E-06 | 0.00045035 | 6.67469274 | 4.47070684 | 3.46443125 | 5.34795161 | 6.67469274 | ZT0 | Present_Koike2012 | Absent_Koike2012 |
| ENSMUSG00000029238 | Clock | 5 | 76639992 | 76733817 | - | 4.95E-06 | 0.00048968 | 8.71843714 | 6.41628511 | 3.20038883 | 6.01493701 | 8.71843714 | ZT0 | Present_Koike2012 | Present_Koike2012 |
| ENSMUSG00000022512 | Cldn1 | 16 | 26356728 | 26371927 | - | 6.09E-06 | 0.00056173 | 33.5607285 | 30.3514495 | 9.58974825 | 27.2695803 | 33.5607285 | ZT0 | Present_Koike2012 | Present_Koike2012 |
| ENSMUSG00000029003 | Mad2l2 | 4 | 147504493 | 147519808 | + | 6.35E-06 | 0.00057938 | 2.65805961 | 2.09409149 | 1.11732012 | 1.77831348 | 2.65805961 | ZT0 | Present_Koike2012 | Absent_Koike2012 |
| ENSMUSG00000028669 | 1110049F12I | 4 | 135531375 | 135543179 | - | 7.11E-06 | 0.00063268 | 6.44821066 | 5.56352971 | 4.47270548 | 5.18234175 | 6.44821066 | ZT0 | Present_Koike2012 | Absent_Koike2012 |
| ENSMUSG00000090107 | Gm11492 | 11 | 87380155 | 87382752 | + | 7.30E-06 | 0.00063613 | 7.64646604 | 5.17836451 | 4.64828651 | 6.80701662 | 7.64646604 | ZT0 | Absent_Koike2012 | Absent_Koike2012 |
| ENSMUSG00000031901 | Dus2l | 8 | 108515237 | 108577740 | + | 7.52E-06 | 0.00065065 | 1.82584798 | 1.40547269 | 0.99231379 | 1.40609112 | 1.82584798 | ZT0 | Absent_Koike2012 | Absent_Koike2012 |
| ENSMUSG00000055401 | Fbxo6 | 4 | 147519825 | 147526249 | - | 7.67E-06 | 0.00066154 | 8.86668176 | 8.24648419 | 5.94620119 | 6.42594218 | 8.86668176 | ZT0 | Present_Koike2012 | Absent_Koike2012 |
| ENSMUSG00000031549 | Ido2 | 8 | 25642364 | 25686805 | - | 8.07E-06 | 0.00068033 | 8.14568573 | 6.1941267 | 5.55419293 | 8.05874522 | 8.14568573 | ZT0 | Absent_Koike2012 | Absent_Koike2012 |
| ENSMUSG00000053334 | Ficd | 5 | 114185791 | 114190609 | + | 8.42E-06 | 0.00070026 | 4.02678875 | 1.94185125 | 2.371184 | 2.3191565 | 4.02678875 | ZT0 | Absent_Koike2012 | Present_Koike2012 |
| ENSMUSG00000030036 | Mogs | 6 | 83065490 | 83068892 | + | 8.77E-06 | 0.00071011 | 14.5702767 | 11.3703359 | 12.7256144 | 14.4922329 | 14.5702767 | ZT0 | Absent_Koike2012 | Absent_Koike2012 |
| ENSMUSG00000001229 | Dpp9 | 17 | 56326105 | 56358312 | - | 9.12E-06 | 0.00072718 | 9.70015659 | 8.16390281 | 5.65554754 | 7.32106037 | 9.70015659 | ZT0 | Present_Koike2012 | Absent_Koike2012 |
| ENSMUSG00000091956 | C2cd4b | 9 | 67607244 | 67608736 | + | 9.79E-06 | 0.00076677 | 0.0413435 | 0 | 0 | 0 | 0.0413435 | ZT0 | Absent_Koike2012 | Absent_Koike2012 |
| ENSMUSG00000021559 | Dapk1 | 13 | 60703308 | 60864546 | + | 1.04E-05 | 0.00079946 | 11.5285234 | 9.88960208 | 6.85683107 | 10.690715 | 11.5285234 | ZT0 | Present_Koike2012 | Present_Koike2012 |

| | | | | | | | | | | | | | | | |
|---|---|---|---|---|---|---|---|---|---|---|---|---|---|---|---|
| ENSMUSG00000025185 | Loxl4 | 19 | 42668472 | 42687303 | - | 1.05E-05 | 0.00079994 | 2.50343055 | 0.60480703 | 0.25374709 | 0.59224423 | 2.50343055 | ZT0 | Present_Koike2012 | Absent_Koike2012 |
| ENSMUSG00000018547 | Pip4k2b | 11 | 97576471 | 97606018 | - | 1.28E-05 | 0.0009025 | 4.24392714 | 3.12916793 | 3.24945005 | 3.98349374 | 4.24392714 | ZT0 | Absent_Koike2012 | Absent_Koike2012 |
| ENSMUSG00000085990 | Gm16731 | 4 | 46548773 | 46552674 | + | 1.29E-05 | 0.0009093 | 1.12042606 | 0.43111657 | 0.35321177 | 0.7503769 | 1.12042606 | ZT0 | Absent_Koike2012 | Absent_Koike2012 |
| ENSMUSG00000024160 | Spsb3 | 17 | 25023588 | 25029097 | + | 1.36E-05 | 0.00093173 | 1.74508925 | 1.25106955 | 0.89625499 | 1.37859769 | 1.74508925 | ZT0 | Absent_Koike2012 | Absent_Koike2012 |
| ENSMUSG00000009566 | Fpgs | 2 | 32538129 | 32559657 | - | 1.45E-05 | 0.00096032 | 15.4183269 | 12.8310155 | 6.66469293 | 13.2352295 | 15.4183269 | ZT0 | Present_Koike2012 | Present_Koike2012 |
| ENSMUSG00000030762 | Aqp8 | 7 | 130605808 | 130611517 | + | 1.50E-05 | 0.0009807 | 68.2536187 | 33.6261808 | 12.9932621 | 48.2970862 | 68.2536187 | ZT0 | Present_Koike2012 | Present_Koike2012 |
| ENSMUSG00000040888 | Gfer | 17 | 24830136 | 24833101 | - | 1.50E-05 | 0.0009807 | 12.9132933 | 9.4582185 | 8.430769 | 10.223789 | 12.9132933 | ZT0 | Present_Koike2012 | Absent_Koike2012 |
| ENSMUSG00000006356 | Crip2 | 12 | 114378447 | 114383715 | + | 1.66E-05 | 0.00105505 | 23.8976345 | 18.831934 | 9.87064775 | 15.181146 | 23.8976345 | ZT0 | Present_Koike2012 | Absent_Koike2012 |
| ENSMUSG00000024812 | Tjp2 | 19 | 24169013 | 24299444 | - | 1.80E-05 | 0.00111876 | 5.13095247 | 4.57357952 | 3.36530878 | 4.73857479 | 5.13095247 | ZT0 | Absent_Koike2012 | Absent_Koike2012 |
| ENSMUSG00000052751 | Repin1 | 6 | 48543882 | 48549081 | + | 1.82E-05 | 0.00112572 | 7.02175379 | 6.48042201 | 5.11773021 | 6.51988996 | 7.02175379 | ZT0 | Absent_Koike2012 | Absent_Koike2012 |
| ENSMUSG00000035498 | Cdcp1 | 9 | 123079942 | 123125156 | - | 1.88E-05 | 0.00114201 | 1.18643363 | 0.53177209 | 0.40447502 | 0.81138473 | 1.18643363 | ZT0 | Absent_Koike2012 | Present_Koike2012 |
| ENSMUSG00000038150 | Ormdl3 | 11 | 98442570 | 98448682 | - | 1.93E-05 | 0.00116159 | 30.2645908 | 19.7908853 | 20.7244723 | 29.5470888 | 30.2645908 | ZT0 | Present_Koike2012 | Absent_Koike2012 |
| ENSMUSG00000004931 | Apba3 | 10 | 80730917 | 80735991 | + | 2.22E-05 | 0.00129776 | 12.7877019 | 9.19159836 | 9.91044804 | 12.4902254 | 12.7877019 | ZT0 | Absent_Koike2012 | Absent_Koike2012 |
| ENSMUSG00000071633 | Gm4952 | 19 | 12674506 | 12702106 | + | 2.28E-05 | 0.00130849 | 45.1638546 | 42.6459072 | 16.1079334 | 18.9520612 | 45.1638546 | ZT0 | Present_Koike2012 | Present_Koike2012 |
| ENSMUSG00000015971 | Actr8 | 14 | 30791523 | 30806407 | + | 2.30E-05 | 0.00130891 | 8.19413695 | 7.92802868 | 8.11746421 | 6.99385585 | 8.19413695 | ZT0 | Absent_Koike2012 | Absent_Koike2012 |
| ENSMUSG00000039220 | Ppp1r10 | 17 | 36053379 | 36069226 | + | 2.64E-05 | 0.00143416 | 5.29161135 | 2.84121062 | 4.84236334 | 4.73140287 | 5.29161135 | ZT0 | Present_Koike2012 | Absent_Koike2012 |
| ENSMUSG00000031217 | Efnb1 | X | 96331469 | 96344330 | + | 2.84E-05 | 0.00150069 | 3.53495894 | 2.66999631 | 1.98926601 | 3.16676234 | 3.53495894 | ZT0 | Absent_Koike2012 | Absent_Koike2012 |
| ENSMUSG00000056749 | Nfil3 | 13 | 53062578 | 53076442 | - | 3.06E-05 | 0.00156349 | 16.1906875 | 6.14833684 | 5.8185476 | 10.6176113 | 16.1906875 | ZT0 | Present_Koike2012 | Present_Koike2012 |
| ENSMUSG00000022040 | Ephx2 | 14 | 66703214 | 66743337 | - | 3.69E-05 | 0.00178641 | 320.40709 | 256.928578 | 283.475146 | 301.566788 | 320.40709 | ZT0 | Absent_Koike2012 | Absent_Koike2012 |
| ENSMUSG00000087679 | C330006A16 | 2 | 25992334 | 25996041 | - | 3.69E-05 | 0.00178641 | 18.5271415 | 15.9395919 | 13.6144287 | 17.1489867 | 18.5271415 | ZT0 | Absent_Koike2012 | Absent_Koike2012 |
| ENSMUSG00000025227 | Tmem180 | 19 | 46415611 | 46449742 | + | 3.73E-05 | 0.0017947 | 0.61562834 | 0.46805515 | 0.31989624 | 0.53013848 | 0.61562834 | ZT0 | Absent_Koike2012 | Absent_Koike2012 |
| ENSMUSG00000035845 | Alg12 | 15 | 88635674 | 88649748 | - | 3.77E-05 | 0.00180628 | 2.76234456 | 1.43217401 | 1.62136842 | 2.26549942 | 2.76234456 | ZT0 | Absent_Koike2012 | Absent_Koike2012 |
| ENSMUSG00000024790 | Sac3d1 | 19 | 6116004 | 6118586 | - | 3.91E-05 | 0.0018616 | 2.44029375 | 1.488117 | 2.198361 | 2.29161975 | 2.44029375 | ZT0 | Absent_Koike2012 | Absent_Koike2012 |
| ENSMUSG00000022769 | Sdf2l1 | 16 | 17130231 | 17132476 | - | 4.01E-05 | 0.00189478 | 33.3310723 | 18.063526 | 12.9048868 | 29.6304983 | 33.3310723 | ZT0 | Present_Koike2012 | Absent_Koike2012 |
| ENSMUSG00000042992 | Loh12cr1 | 6 | 134589645 | 134661202 | + | 4.19E-05 | 0.00194776 | 4.00837847 | 3.1182027 | 3.07639146 | 3.80891298 | 4.00837847 | ZT0 | Absent_Koike2012 | Present_Koike2012 |
| ENSMUSG00000041782 | Lad1 | 1 | 137715175 | 137729918 | + | 4.19E-05 | 0.00194776 | 2.24715975 | 1.219896 | 1.03409075 | 1.780007 | 2.24715975 | ZT0 | Absent_Koike2012 | Absent_Koike2012 |
| ENSMUSG00000001774 | Chordc1 | 9 | 18096711 | 18118444 | + | 4.34E-05 | 0.0019957 | 16.966674 | 8.88831625 | 13.460493 | 16.4075005 | 16.966674 | ZT0 | Absent_Koike2012 | Present_Koike2012 |
| ENSMUSG00000029467 | Atp2a2 | 5 | 122903531 | 122952183 | - | 4.39E-05 | 0.00199897 | 30.4664968 | 20.2442186 | 21.0054018 | 29.1257642 | 30.4664968 | ZT0 | Absent_Koike2012 | Present_Koike2012 |
| ENSMUSG00000036120 | Rfxank | 8 | 72654711 | 72663096 | - | 4.39E-05 | 0.00199897 | 3.35726623 | 1.89196428 | 1.26795181 | 2.1395486 | 3.35726623 | ZT0 | Present_Koike2012 | Absent_Koike2012 |
| ENSMUSG00000011305 | Plin5 | 17 | 56251024 | 56257019 | - | 4.47E-05 | 0.00202635 | 16.1279484 | 8.06655431 | 11.2868161 | 13.5111471 | 16.1279484 | ZT0 | Absent_Koike2012 | Absent_Koike2012 |
| ENSMUSG00000030847 | Bag3 | 7 | 135667097 | 135690491 | + | 4.62E-05 | 0.00206788 | 8.715283 | 3.267241 | 4.12863125 | 5.89655925 | 8.715283 | ZT0 | Present_Koike2012 | Present_Koike2012 |
| ENSMUSG00000030871 | Ears2 | 7 | 129180727 | 129210777 | - | 5.01E-05 | 0.0021599 | 1.79430467 | 1.33548598 | 1.30345891 | 1.71953504 | 1.79430467 | ZT0 | Absent_Koike2012 | Absent_Koike2012 |
| ENSMUSG00000021751 | Acox2 | 14 | 9058025 | 9091867 | - | 5.33E-05 | 0.00226662 | 47.1533567 | 42.2786344 | 32.7915972 | 38.9826618 | 47.1533567 | ZT0 | Present_Koike2012 | Present_Koike2012 |
| ENSMUSG00000049922 | Slc35c1 | 2 | 92292921 | 92300695 | - | 5.43E-05 | 0.00227744 | 2.04751589 | 1.37370839 | 1.43257342 | 1.9106433 | 2.04751589 | ZT0 | Absent_Koike2012 | Absent_Koike2012 |
| ENSMUSG00000056724 | Nbeal2 | 9 | 110527293 | 110556665 | - | 5.58E-05 | 0.00232321 | 1.032297 | 0.64608148 | 0.65262383 | 1.00815803 | 1.032297 | ZT0 | Absent_Koike2012 | Present_Koike2012 |
| ENSMUSG00000024958 | Gpr137 | 19 | 7012559 | 7016940 | - | 5.63E-05 | 0.0023292 | 2.509365 | 1.98771625 | 1.75943413 | 2.02633889 | 2.509365 | ZT0 | Present_Koike2012 | Absent_Koike2012 |
| ENSMUSG00000046962 | Zfp295 | 16 | 98168599 | 98183786 | - | 5.72E-05 | 0.00235429 | 2.54350581 | 1.08272107 | 1.32516627 | 2.12257857 | 2.54350581 | ZT0 | Present_Koike2012 | Absent_Koike2012 |
| ENSMUSG00000001761 | Smo | 6 | 29685694 | 29711365 | + | 5.77E-05 | 0.00236138 | 1.4337555 | 1.290581 | 0.82061075 | 0.818594 | 1.4337555 | ZT0 | Absent_Koike2012 | Absent_Koike2012 |
| ENSMUSG00000031024 | St5 | 7 | 116667425 | 116760661 | - | 5.82E-05 | 0.00236871 | 5.69611233 | 5.12358441 | 1.91064918 | 4.973142 | 5.69611233 | ZT0 | Present_Koike2012 | Present_Koike2012 |
| ENSMUSG00000090611 | C330013E15I | 15 | 100444571 | 100445541 | - | 6.04E-05 | 0.00244134 | 1.230444 | 0.7567835 | 0.65188425 | 1.13234875 | 1.230444 | ZT0 | Absent_Koike2012 | Absent_Koike2012 |
| ENSMUSG00000086844 | B230206H07 | 7 | 148545040 | 148551009 | - | 6.24E-05 | 0.00248712 | 3.16397979 | 3.1418143 | 1.06437824 | 1.80920279 | 3.16397979 | ZT0 | Absent_Koike2012 | Absent_Koike2012 |
| ENSMUSG00000062762 | Ei24 | 9 | 36586740 | 36604938 | - | 6.23E-05 | 0.00248712 | 50.9676062 | 45.8347423 | 40.3430146 | 43.7838562 | 50.9676062 | ZT0 | Present_Koike2012 | Present_Koike2012 |
| ENSMUSG00000044847 | Lsm11 | 11 | 45741771 | 45758437 | - | 6.27E-05 | 0.00249862 | 0.53589525 | 0.3997815 | 0.425309 | 0.52845225 | 0.53589525 | ZT0 | Absent_Koike2012 | Absent_Koike2012 |
| ENSMUSG00000092563 | Gm3617 | 1 | 39371744 | 39372298 | + | 6.29E-05 | 0.00249929 | 0.0816645 | 0.00628775 | 0.00380125 | 0.01409925 | 0.0816645 | ZT0 | Absent_Koike2012 | Absent_Koike2012 |
| ENSMUSG00000052214 | Opa3 | 7 | 19813683 | 19841892 | + | 6.31E-05 | 0.00250194 | 6.94280822 | 5.30082922 | 6.65389449 | 6.76275261 | 6.94280822 | ZT0 | Absent_Koike2012 | Absent_Koike2012 |
| ENSMUSG00000085031 | Gm16982 | 7 | 148795965 | 148799646 | - | 6.84E-05 | 0.00263289 | 0.7362705 | 0.55334475 | 0.49195875 | 0.6343195 | 0.7362705 | ZT0 | Absent_Koike2012 | Absent_Koike2012 |
| ENSMUSG00000049555 | Tmie | 9 | 110768215 | 110782617 | - | 6.90E-05 | 0.00264663 | 5.05347963 | 2.56170599 | 2.12519394 | 4.62009106 | 5.05347963 | ZT0 | Present_Koike2012 | Absent_Koike2012 |
| ENSMUSG00000016344 | Ppdpf | 2 | 180921952 | 180923476 | + | 7.21E-05 | 0.00272763 | 20.6506139 | 17.5407558 | 14.0314679 | 17.234305 | 20.6506139 | ZT0 | Absent_Koike2012 | Absent_Koike2012 |
| ENSMUSG00000068876 | Cgn | 3 | 94464507 | 94590414 | - | 7.53E-05 | 0.00281147 | 3.20261436 | 2.45680744 | 1.88886473 | 2.96273291 | 3.20261436 | ZT0 | Absent_Koike2012 | Present_Koike2012 |
| ENSMUSG00000032575 | Manf | 9 | 106740643 | 106794310 | - | 7.61E-05 | 0.00282578 | 22.2080335 | 9.88857762 | 13.86371 | 22.1493819 | 22.2080335 | ZT0 | Absent_Koike2012 | Absent_Koike2012 |
| ENSMUSG00000030771 | Micalcl | 7 | 119511822 | 119556617 | + | 7.63E-05 | 0.00282583 | 0.32761439 | 0.12584661 | 0.07844817 | 0.19958742 | 0.32761439 | ZT0 | Absent_Koike2012 | Absent_Koike2012 |
| ENSMUSG00000047617 | BC029214 | 2 | 25310661 | 25316614 | - | 7.65E-05 | 0.00282825 | 4.68108891 | 1.99675696 | 1.3692421 | 1.95241785 | 4.68108891 | ZT0 | Present_Koike2012 | Absent_Koike2012 |
| ENSMUSG00000030031 | Kbtbd8 | 6 | 95067234 | 95079712 | + | 7.68E-05 | 0.00283476 | 0.98695203 | 0.27639975 | 0.83452888 | 0.68820583 | 0.98695203 | ZT0 | Present_Koike2012 | Absent_Koike2012 |
| ENSMUSG00000057606 | Colq | 14 | 32336268 | 32404529 | - | 7.78E-05 | 0.0028533 | 25.4553213 | 18.007551 | 10.8522873 | 21.0025598 | 25.4553213 | ZT0 | Absent_Koike2012 | Absent_Koike2012 |
| ENSMUSG00000032615 | Nt5m | 11 | 59652949 | 59694470 | + | 8.84E-05 | 0.00313011 | 1.88465485 | 1.32838268 | 1.49819038 | 1.74198316 | 1.88465485 | ZT0 | Absent_Koike2012 | Absent_Koike2012 |
| ENSMUSG00000071711 | Mpst | 15 | 78237142 | 78244432 | + | 9.00E-05 | 0.00316614 | 25.8612751 | 19.5280907 | 19.4734474 | 25.3972032 | 25.8612751 | ZT0 | Present_Koike2012 | Absent_Koike2012 |
| ENSMUSG00000053617 | Sh3pxd2a | 19 | 47334664 | 47538901 | - | 9.43E-05 | 0.00328306 | 4.21758127 | 3.62946207 | 2.19661381 | 3.08520031 | 4.21758127 | ZT0 | Absent_Koike2012 | Absent_Koike2012 |
| ENSMUSG00000052632 | Asap2 | 12 | 20996320 | 21276032 | + | 9.62E-05 | 0.00332302 | 0.97816183 | 0.7160251 | 0.25209607 | 0.53806614 | 0.97816183 | ZT0 | Present_Koike2012 | Present_Koike2012 |
| ENSMUSG00000039616 | Mocos | 18 | 24812192 | 24860057 | + | 9.68E-05 | 0.00333663 | 17.1030163 | 15.8869388 | 13.2553765 | 15.8254198 | 17.1030163 | ZT0 | Present_Koike2012 | Absent_Koike2012 |
| ENSMUSG00000002963 | Pnkp | 7 | 52112509 | 52118295 | + | 9.75E-05 | 0.00334947 | 5.10370847 | 4.36266094 | 3.6223741 | 4.04090299 | 5.10370847 | ZT0 | Present_Koike2012 | Absent_Koike2012 |
| ENSMUSG00000026811 | St6galnac6 | 2 | 32455229 | 32476326 | + | 9.85E-05 | 0.00337067 | 6.86218299 | 5.82107567 | 5.15607283 | 6.74484412 | 6.86218299 | ZT0 | Absent_Koike2012 | Absent_Koike2012 |
| ENSMUSG00000042487 | Leo1 | 9 | 75289331 | 75314239 | + | 9.91E-05 | 0.00337453 | 8.16861675 | 5.0687505 | 3.6655205 | 5.86794475 | 8.16861675 | ZT0 | Present_Koike2012 | Absent_Koike2012 |
| ENSMUSG00000003585 | Sec14l2 | 11 | 3997042 | 4023418 | - | 0.00010064 | 0.00342366 | 121.68178 | 105.595877 | 92.3063358 | 110.813042 | 121.68178 | ZT0 | Present_Koike2012 | Present_Koike2012 |
| ENSMUSG00000026974 | Zmynd19 | 2 | 24805312 | 24817595 | + | 0.00010308 | 0.00348811 | 1.78639738 | 1.18992538 | 1.77312555 | 1.65436529 | 1.78639738 | ZT0 | Absent_Koike2012 | Absent_Koike2012 |
| ENSMUSG00000008435 | Rdh13 | 7 | 4376372 | 4397251 | - | 0.00011139 | 0.00369556 | 1.57085302 | 0.99038879 | 1.56452907 | 1.00596311 | 1.57085302 | ZT0 | Present_Koike2012 | Absent_Koike2012 |
| ENSMUSG00000008855 | Hdac5 | 11 | 102055746 | 102091480 | - | 0.00011473 | 0.00374465 | 2.81383135 | 2.51160067 | 1.81770417 | 2.80291251 | 2.81383135 | ZT0 | Present_Koike2012 | Present_Koike2012 |
| ENSMUSG00000036613 | Tssc1 | 12 | 29436693 | 29552356 | + | 0.00011488 | 0.00374465 | 4.15995612 | 3.16278932 | 2.76488532 | 3.67690912 | 4.15995612 | ZT0 | Present_Koike2012 | Absent_Koike2012 |
| ENSMUSG00000023067 | Cdkn1a | 17 | 29227924 | 29237667 | + | 0.00012009 | 0.00388289 | 2.15319961 | 0.30316184 | 0.31819737 | 0.65713274 | 2.15319961 | ZT0 | Present_Koike2012 | Absent_Koike2012 |
| ENSMUSG00000038055 | Dexi | 16 | 10530300 | 10543147 | - | 0.00012089 | 0.00390384 | 7.71560765 | 5.65834712 | 5.43367838 | 7.33966094 | 7.71560765 | ZT0 | Present_Koike2012 | Absent_Koike2012 |

| | | | | | | | | | | | | | | | |
|---|---|---|---|---|---|---|---|---|---|---|---|---|---|---|---|
| ENSMUSG00000043398 | Gpr135 | 12 | 73170605 | 73171978 | - | 0.00012378 | 0.00396601 | 0.2263175 | 0.09607675 | 0.19847275 | 0.12012425 | 0.2263175 | ZT0 | Absent_Koike2012 | Absent_Koike2012 |
| ENSMUSG00000030748 | Il4ra | 7 | 132695785 | 132722986 | + | 0.00012415 | 0.00396601 | 5.85636597 | 5.36126732 | 4.2678311 | 5.47325061 | 5.85636597 | ZT0 | Present_Koike2012 | Present_Koike2012 |
| ENSMUSG00000024807 | Syvn1 | 19 | 6046576 | 6053712 | + | 0.00012422 | 0.00396601 | 5.45907669 | 2.54896128 | 2.52938066 | 4.17159114 | 5.45907669 | ZT0 | Absent_Koike2012 | Absent_Koike2012 |
| ENSMUSG00000091093 | Gm17302 | 12 | 101447198 | 101448024 | - | 0.00012913 | 0.00406664 | 9.66902325 | 7.3908325 | 7.12056625 | 8.7215535 | 9.66902325 | ZT0 | Absent_Koike2012 | Absent_Koike2012 |
| ENSMUSG00000057778 | Cyb5d2 | 11 | 72590734 | 72609644 | - | 0.00013584 | 0.00420527 | 2.56107058 | 2.1034482 | 1.97584205 | 2.29859902 | 2.56107058 | ZT0 | Absent_Koike2012 | Absent_Koike2012 |
| ENSMUSG00000025408 | Ddit3 | 10 | 126727830 | 126733344 | + | 0.00013796 | 0.00425014 | 1.86310406 | 0.70861404 | 0.78719471 | 1.07386099 | 1.86310406 | ZT0 | Absent_Koike2012 | Absent_Koike2012 |
| ENSMUSG00000075225 | Ccdc162 | 10 | 41258652 | 41429106 | - | 0.00013878 | 0.00427042 | 1.18194745 | 1.06836398 | 0.63615515 | 0.74510087 | 1.18194745 | ZT0 | Present_Koike2012 | Present_Koike2012 |
| ENSMUSG00000005469 | Prkaca | 8 | 86496877 | 86520344 | + | 0.00013935 | 0.00428272 | 16.2811703 | 14.5216148 | 12.8405004 | 14.5161761 | 16.2811703 | ZT0 | Absent_Koike2012 | Absent_Koike2012 |
| ENSMUSG00000019818 | Cd164 | 10 | 41239306 | 41250848 | + | 0.00014277 | 0.00433658 | 87.6229862 | 82.7984913 | 69.1168626 | 70.083134 | 87.6229862 | ZT0 | Absent_Koike2012 | Absent_Koike2012 |
| ENSMUSG00000059708 | Akap17b | X | 34148309 | 34185390 | - | 0.00014242 | 0.00433658 | 1.51154567 | 0.78481801 | 1.18654441 | 1.33475571 | 1.51154567 | ZT0 | Absent_Koike2012 | Absent_Koike2012 |
| ENSMUSG00000031864 | Ints10 | 8 | 71317828 | 71353309 | + | 0.00014587 | 0.00438917 | 1.03415463 | 0.67065615 | 0.78220262 | 0.99051098 | 1.03415463 | ZT0 | Absent_Koike2012 | Absent_Koike2012 |
| ENSMUSG00000045752 | Tssc4 | 7 | 150255154 | 150256998 | + | 0.00014826 | 0.0044235 | 4.22503174 | 3.12721825 | 3.10038566 | 3.67080524 | 4.22503174 | ZT0 | Present_Koike2012 | Absent_Koike2012 |
| ENSMUSG00000028042 | Zbtb7b | 3 | 89181566 | 89198698 | - | 0.00015091 | 0.00448262 | 8.59485531 | 8.16160285 | 6.48334995 | 6.87753642 | 8.59485531 | ZT0 | Present_Koike2012 | Absent_Koike2012 |
| ENSMUSG00000039737 | Prkrip1 | 5 | 136653997 | 136674833 | - | 0.00015183 | 0.00449242 | 1.05497455 | 0.74193619 | 0.82796103 | 0.74430618 | 1.05497455 | ZT0 | Absent_Koike2012 | Absent_Koike2012 |
| ENSMUSG00000004500 | Zfp324 | 7 | 13551187 | 13559585 | + | 0.00015513 | 0.00455834 | 2.68839925 | 1.45636362 | 1.62528883 | 2.05639002 | 2.68839925 | ZT0 | Absent_Koike2012 | Absent_Koike2012 |
| ENSMUSG00000033253 | Szt2 | 4 | 118035348 | 118081878 | - | 0.00016321 | 0.00473558 | 1.19181938 | 0.99143057 | 0.72989453 | 1.02936616 | 1.19181938 | ZT0 | Absent_Koike2012 | Absent_Koike2012 |
| ENSMUSG00000032902 | Slc16a1 | 3 | 104442591 | 104462385 | + | 0.00016664 | 0.00478918 | 17.852793 | 12.8604093 | 9.7450895 | 13.6455803 | 17.852793 | ZT0 | Present_Koike2012 | Absent_Koike2012 |
| ENSMUSG00000029641 | Rasl11a | 5 | 147656647 | 147659302 | + | 0.00016889 | 0.00483065 | 0.29574 | 0.10501075 | 0.0984005 | 0.23524625 | 0.29574 | ZT0 | Absent_Koike2012 | Absent_Koike2012 |
| ENSMUSG00000050445 | Cyp8b1 | 9 | 121823474 | 121825423 | - | 0.00016942 | 0.0048386 | 124.9252 | 88.937525 | 41.3587838 | 99.087616 | 124.9252 | ZT0 | Present_Koike2012 | Absent_Koike2012 |
| ENSMUSG00000020182 | Ddc | 11 | 11714104 | 11798047 | - | 0.00017674 | 0.00497504 | 19.6029127 | 12.5120813 | 8.38535847 | 15.0676374 | 19.6029127 | ZT0 | Present_Koike2012 | Absent_Koike2012 |
| ENSMUSG00000060862 | Zbtb40 | 4 | 136535647 | 136604716 | - | 0.00017852 | 0.00500845 | 1.19275074 | 0.60442535 | 0.87388009 | 0.94476664 | 1.19275074 | ZT0 | Absent_Koike2012 | Absent_Koike2012 |
| ENSMUSG00000026389 | Steap3 | 1 | 122087334 | 122169282 | - | 0.00018205 | 0.00506768 | 2.26298981 | 2.07014735 | 1.33578623 | 1.78158898 | 2.26298981 | ZT0 | Present_Koike2012 | Present_Koike2012 |
| ENSMUSG00000044641 | Pard6b | 2 | 167906504 | 167926703 | + | 0.00019453 | 0.00532857 | 1.942214 | 1.3297255 | 1.24188075 | 1.56945425 | 1.942214 | ZT0 | Absent_Koike2012 | Absent_Koike2012 |
| ENSMUSG00000042210 | Abhd14a | 9 | 106342382 | 106350009 | - | 0.00019578 | 0.0053571 | 4.19149884 | 3.38294487 | 2.6954788 | 3.49348697 | 4.19149884 | ZT0 | Present_Koike2012 | Absent_Koike2012 |
| ENSMUSG00000068114 | Ccdc134 | 15 | 81958352 | 81972630 | + | 0.00019986 | 0.00543983 | 1.5475715 | 0.6726525 | 0.88678275 | 1.07636225 | 1.5475715 | ZT0 | Absent_Koike2012 | Absent_Koike2012 |
| ENSMUSG00000021400 | Wrnip1 | 13 | 32893907 | 32914478 | + | 0.00020683 | 0.00556962 | 11.3397385 | 10.1242525 | 6.1664585 | 9.844837 | 11.3397385 | ZT0 | Present_Koike2012 | Absent_Koike2012 |
| ENSMUSG00000045287 | Rtn4rl1 | 11 | 75007285 | 75081271 | + | 0.00020741 | 0.00557915 | 2.43376325 | 1.853306 | 1.2712845 | 2.13237925 | 2.43376325 | ZT0 | Present_Koike2012 | Absent_Koike2012 |
| ENSMUSG00000067279 | Ppp1r3c | 19 | 36806227 | 36811143 | - | 0.0002083 | 0.00558593 | 46.2904443 | 20.935975 | 9.9763325 | 20.2173048 | 46.2904443 | ZT0 | Present_Koike2012 | Present_Koike2012 |
| ENSMUSG00000039242 | B3galnt2 | 13 | 14046941 | 14091335 | + | 0.00020865 | 0.00558908 | 6.58774 | 4.178858 | 5.33376875 | 6.274695 | 6.58774 | ZT0 | Present_Koike2012 | Present_Koike2012 |
| ENSMUSG00000032607 | Amt | 9 | 108199184 | 108203902 | + | 0.00021228 | 0.00565651 | 11.2430161 | 8.90748129 | 4.87693923 | 5.61539339 | 11.2430161 | ZT0 | Present_Koike2012 | Absent_Koike2012 |
| ENSMUSG00000005483 | Dnajb1 | 8 | 86132074 | 86135801 | + | 0.00021254 | 0.00565739 | 9.67894688 | 3.787362 | 6.42252149 | 7.34630954 | 9.67894688 | ZT0 | Present_Koike2012 | Absent_Koike2012 |
| ENSMUSG00000057134 | Ado | 10 | 67007260 | 67011703 | - | 0.00021348 | 0.00566609 | 4.650578 | 3.65637125 | 3.667706 | 4.4363675 | 4.650578 | ZT0 | Absent_Koike2012 | Absent_Koike2012 |
| ENSMUSG00000047123 | Ticam1 | 17 | 56408742 | 56416209 | - | 0.00022383 | 0.00585111 | 2.98618175 | 2.177516 | 1.85707225 | 1.88140825 | 2.98618175 | ZT0 | Absent_Koike2012 | Absent_Koike2012 |
| ENSMUSG00000080679 | AC112790.1 | 10 | 86308556 | 86308653 | + | 0.00022583 | 0.005888 | 0.05438725 | 0.00573975 | 0 | 0.0062075 | 0.05438725 | ZT0 | Absent_Koike2012 | Absent_Koike2012 |
| ENSMUSG00000022614 | Lmf2 | 15 | 89181435 | 89186090 | - | 0.00023493 | 0.00604067 | 4.23669775 | 3.93024675 | 3.43792475 | 3.687063 | 4.23669775 | ZT0 | Absent_Koike2012 | Present_Koike2012 |
| ENSMUSG00000018102 | Hist1h2bc | 13 | 23776068 | 23784357 | + | 0.00024172 | 0.00612648 | 79.829254 | 60.4135083 | 55.3810108 | 72.978199 | 79.829254 | ZT0 | Present_Koike2012 | Present_Koike2012 |
| ENSMUSG00000036216 | Leap2 | 11 | 53235683 | 53236672 | - | 0.00024363 | 0.00615142 | 84.7195685 | 43.725132 | 36.7163493 | 61.7100785 | 84.7195685 | ZT0 | Present_Koike2012 | Present_Koike2012 |
| ENSMUSG00000078584 | AU022252 | 4 | 118876175 | 118905317 | - | 0.00024479 | 0.0061735 | 4.81769182 | 4.71039889 | 3.58780124 | 4.00455799 | 4.81769182 | ZT0 | Absent_Koike2012 | Present_Koike2012 |
| ENSMUSG00000040128 | Pnrc1 | 4 | 33332398 | 33377138 | - | 0.00024658 | 0.00620437 | 23.612188 | 14.0671178 | 9.87810222 | 15.3196278 | 23.612188 | ZT0 | Present_Koike2012 | Present_Koike2012 |
| ENSMUSG00000020782 | Llgl2 | 11 | 115685363 | 115717094 | + | 0.00024824 | 0.00621735 | 3.48933111 | 3.31593984 | 2.58227306 | 3.37740556 | 3.48933111 | ZT0 | Absent_Koike2012 | Absent_Koike2012 |
| ENSMUSG00000002658 | Gtf2f1 | 17 | 57142826 | 57150723 | - | 0.00025583 | 0.00631908 | 9.33794008 | 6.4845083 | 7.21056083 | 8.35237623 | 9.33794008 | ZT0 | Absent_Koike2012 | Absent_Koike2012 |
| ENSMUSG00000003228 | Grk5 | 19 | 60965602 | 61168456 | + | 0.00025602 | 0.00631908 | 1.74215805 | 0.80339589 | 0.37071029 | 0.69787386 | 1.74215805 | ZT0 | Present_Koike2012 | Absent_Koike2012 |
| ENSMUSG00000001138 | Cnnm3 | 1 | 36568712 | 36585082 | + | 0.00025704 | 0.00632571 | 1.76301081 | 1.64037206 | 1.24247124 | 1.3866442 | 1.76301081 | ZT0 | Absent_Koike2012 | Absent_Koike2012 |
| ENSMUSG00000032121 | Tmem218 | 9 | 37015918 | 37030795 | + | 0.00025728 | 0.00632571 | 2.54263575 | 2.10355675 | 1.3798925 | 1.13642275 | 2.54263575 | ZT0 | Absent_Koike2012 | Absent_Koike2012 |
| ENSMUSG00000024589 | Nedd4l | 18 | 65047410 | 65377480 | + | 0.00025692 | 0.00632571 | 3.47648558 | 2.16186601 | 1.58619881 | 2.97571012 | 3.47648558 | ZT0 | Absent_Koike2012 | Present_Koike2012 |
| ENSMUSG00000040102 | Klhdc5 | 6 | 147039663 | 147061300 | + | 0.00025912 | 0.00634021 | 3.72350105 | 2.5976518 | 2.96126097 | 3.20338817 | 3.72350105 | ZT0 | Absent_Koike2012 | Absent_Koike2012 |
| ENSMUSG00000021497 | Txndc15 | 13 | 55816011 | 55827582 | + | 0.00026069 | 0.00636639 | 43.4030478 | 38.7750053 | 36.0616683 | 40.3503648 | 43.4030478 | ZT0 | Present_Koike2012 | Absent_Koike2012 |
| ENSMUSG00000037892 | Pcdh18 | 3 | 49547218 | 49561238 | - | 0.00026235 | 0.00638864 | 0.4556875 | 0.26230675 | 0.2261505 | 0.3072175 | 0.4556875 | ZT0 | Absent_Koike2012 | Absent_Koike2012 |
| ENSMUSG00000087275 | 1110002J07F | 10 | 66375267 | 66383006 | - | 0.00027293 | 0.006546 | 0.12967725 | 0.1181685 | 0.04597775 | 0.12794675 | 0.12967725 | ZT0 | Absent_Koike2012 | Absent_Koike2012 |
| ENSMUSG00000058756 | Thra | 11 | 98601952 | 98630320 | + | 0.00027745 | 0.00662264 | 1.68580305 | 1.62706801 | 1.09640739 | 1.16500665 | 1.68580305 | ZT0 | Absent_Koike2012 | Absent_Koike2012 |
| ENSMUSG00000044052 | Ccr10 | 11 | 101034311 | 101036757 | - | 0.00027769 | 0.00662264 | 0.0917835 | 0.015882 | 0.048665 | 0.06227425 | 0.0917835 | ZT0 | Absent_Koike2012 | Absent_Koike2012 |
| ENSMUSG00000022246 | Rai14 | 15 | 10498734 | 10644386 | - | 0.00027994 | 0.00666426 | 5.63303472 | 5.07075828 | 3.33477464 | 4.34832556 | 5.63303472 | ZT0 | Absent_Koike2012 | Absent_Koike2012 |
| ENSMUSG00000010045 | Tmem115 | 9 | 107436276 | 107440987 | + | 0.00028417 | 0.00673308 | 9.30578925 | 8.105533 | 7.9463405 | 7.565788 | 9.30578925 | ZT0 | Absent_Koike2012 | Absent_Koike2012 |
| ENSMUSG00000032280 | Tle3 | 9 | 61220190 | 61266297 | + | 0.00029299 | 0.00688697 | 0.80893606 | 0.61197297 | 0.43952799 | 0.65463871 | 0.80893606 | ZT0 | Absent_Koike2012 | Absent_Koike2012 |
| ENSMUSG00000048217 | Nags | 11 | 102006827 | 102010791 | + | 0.00029262 | 0.00688697 | 17.4678453 | 14.2230907 | 14.3909745 | 16.8984002 | 17.4678453 | ZT0 | Present_Koike2012 | Absent_Koike2012 |
| ENSMUSG00000053536 | Cstf2t | 19 | 31157331 | 31161080 | + | 0.00030393 | 0.00706945 | 9.2463365 | 7.8884975 | 6.90729525 | 7.6019665 | 9.2463365 | ZT0 | Absent_Koike2012 | Absent_Koike2012 |
| ENSMUSG00000031327 | Chic1 | X | 100551815 | 100591431 | + | 0.00031318 | 0.00719922 | 1.76954127 | 1.31777816 | 0.67499377 | 1.1380411 | 1.76954127 | ZT0 | Present_Koike2012 | Absent_Koike2012 |
| ENSMUSG00000020741 | 1300001I01R | 11 | 74462997 | 74484349 | + | 0.00032515 | 0.00738758 | 39.782246 | 36.2516504 | 28.7784455 | 29.7726226 | 39.782246 | ZT0 | Absent_Koike2012 | Absent_Koike2012 |
| ENSMUSG00000013150 | Gfod2 | 8 | 108237754 | 108282564 | - | 0.00033622 | 0.00753834 | 1.4688995 | 0.90668716 | 0.6732706 | 1.23613289 | 1.4688995 | ZT0 | Present_Koike2012 | Absent_Koike2012 |
| ENSMUSG00000024026 | Glo1 | 17 | 30729811 | 30749604 | - | 0.00033688 | 0.00753834 | 187.978017 | 173.294995 | 139.49441 | 138.980826 | 187.978017 | ZT0 | Present_Koike2012 | Absent_Koike2012 |
| ENSMUSG00000016520 | Lnx2 | 5 | 147828231 | 147888162 | - | 0.00036242 | 0.00794363 | 4.71586315 | 3.09579915 | 2.80182968 | 3.85389636 | 4.71586315 | ZT0 | Present_Koike2012 | Present_Koike2012 |
| ENSMUSG00000047554 | Tmem41b | 7 | 117115701 | 117130443 | - | 0.000364 | 0.0079621 | 4.04705401 | 2.64731965 | 3.4769466 | 4.03637297 | 4.04705401 | ZT0 | Absent_Koike2012 | Absent_Koike2012 |
| ENSMUSG00000038910 | Plcl2 | 17 | 50648872 | 50827818 | + | 0.00036516 | 0.00796932 | 3.58048375 | 3.55673125 | 3.2222775 | 3.40914325 | 3.58048375 | ZT0 | Absent_Koike2012 | Absent_Koike2012 |
| ENSMUSG00000020857 | Nme2 | 11 | 93811128 | 93817573 | - | 0.00036603 | 0.00797456 | 31.4512155 | 25.8383675 | 28.8591818 | 30.4245766 | 31.4512155 | ZT0 | Absent_Koike2012 | Absent_Koike2012 |
| ENSMUSG00000074102 | Rbm15b | 9 | 106783260 | 106784620 | - | 0.00036729 | 0.00799528 | 4.43362925 | 2.6359335 | 3.138279 | 3.004075 | 4.43362925 | ZT0 | Present_Koike2012 | Absent_Koike2012 |
| ENSMUSG00000058706 | 0610030E20I | 6 | 72297311 | 72303152 | + | 0.00038241 | 0.00824672 | 4.05228291 | 3.7998779 | 2.36122634 | 2.78165421 | 4.05228291 | ZT0 | Absent_Koike2012 | Absent_Koike2012 |
| ENSMUSG00000041959 | S100a10 | 3 | 93359002 | 93368565 | + | 0.00040568 | 0.00856735 | 41.2045077 | 26.9560427 | 16.4819185 | 25.19079 | 41.2045077 | ZT0 | Present_Koike2012 | Present_Koike2012 |
| ENSMUSG00000031788 | Kifc3 | 8 | 97623732 | 97666440 | - | 0.00040886 | 0.00861271 | 4.74275874 | 2.8524686 | 3.22305834 | 4.6987374 | 4.74275874 | ZT0 | Absent_Koike2012 | Absent_Koike2012 |
| ENSMUSG00000000902 | Smarcb1 | 10 | 75359514 | 75384362 | - | 0.00041069 | 0.00863704 | 1.45765663 | 1.26449501 | 1.07776638 | 1.28736916 | 1.45765663 | ZT0 | Absent_Koike2012 | Absent_Koike2012 |

| | | | | | | | | | | | | | | | |
|---|---|---|---|---|---|---|---|---|---|---|---|---|---|---|---|
| ENSMUSG00000026172 | Bcs1l | 1 | 74634863 | 74639017 | + | 0.00041363 | 0.00867783 | 3.4467299 | 2.7452754 | 2.66409245 | 3.11218341 | 3.4467299 | ZT0 | Present_Koike2012 | Absent_Koike2012 |
| ENSMUSG00000022098 | Bmp1 | 14 | 70874364 | 70920067 | - | 0.00041803 | 0.00873566 | 18.1022746 | 14.9129502 | 14.6813756 | 17.7761426 | 18.1022746 | ZT0 | Absent_Koike2012 | Absent_Koike2012 |
| ENSMUSG00000021336 | Slc17a4 | 13 | 23989607 | 24006876 | - | 0.00041904 | 0.00874055 | 10.077714 | 7.73477991 | 5.4616384 | 5.28189527 | 10.077714 | ZT0 | Present_Koike2012 | Absent_Koike2012 |
| ENSMUSG00000002393 | Nr2f6 | 8 | 73898022 | 73905859 | - | 0.0004287 | 0.00883337 | 13.8198213 | 11.9505621 | 9.6599512 | 12.2811992 | 13.8198213 | ZT0 | Present_Koike2012 | Absent_Koike2012 |
| ENSMUSG00000022574 | Naprt1 | 15 | 75721394 | 75724911 | - | 0.00042936 | 0.00883984 | 31.420709 | 26.2840703 | 25.401727 | 30.9696943 | 31.420709 | ZT0 | Present_Koike2012 | Absent_Koike2012 |
| ENSMUSG00000053460 | Ggcx | 6 | 72364332 | 72380706 | + | 0.00044232 | 0.00906265 | 30.311697 | 26.484645 | 24.8898583 | 26.7023583 | 30.311697 | ZT0 | Absent_Koike2012 | Absent_Koike2012 |
| ENSMUSG00000038422 | Hdhd3 | 4 | 62160042 | 62164501 | - | 0.00044269 | 0.00906276 | 13.8590122 | 8.05981537 | 7.14313094 | 12.2925049 | 13.8590122 | ZT0 | Present_Koike2012 | Absent_Koike2012 |
| ENSMUSG00000001763 | Tspan33 | 6 | 29644222 | 29668559 | + | 0.00044326 | 0.00906723 | 2.17392595 | 1.87592397 | 1.4561794 | 2.12115549 | 2.17392595 | ZT0 | Absent_Koike2012 | Absent_Koike2012 |
| ENSMUSG00000015176 | Nolc1 | 19 | 46150353 | 46160018 | + | 0.00045449 | 0.00921547 | 8.55570074 | 3.57218932 | 5.76067836 | 5.35174832 | 8.55570074 | ZT0 | Absent_Koike2012 | Absent_Koike2012 |
| ENSMUSG00000000876 | Pxmp4 | 2 | 154411494 | 154429444 | - | 0.00045785 | 0.009254 | 21.2842867 | 14.5310962 | 10.6602694 | 14.5558648 | 21.2842867 | ZT0 | Present_Koike2012 | Present_Koike2012 |
| ENSMUSG00000071649 | B3gat3 | 19 | 8994864 | 9001724 | + | 0.00047657 | 0.00955915 | 12.1051952 | 10.3917954 | 7.97986644 | 8.53014092 | 12.1051952 | ZT0 | Present_Koike2012 | Absent_Koike2012 |
| ENSMUSG00000026798 | Coq4 | 2 | 29643013 | 29653455 | + | 0.00047777 | 0.00955915 | 1.41977527 | 1.20699946 | 1.13420674 | 1.18981591 | 1.41977527 | ZT0 | Absent_Koike2012 | Absent_Koike2012 |
| ENSMUSG00000042787 | Exog | 9 | 119354041 | 119374635 | + | 0.0004818 | 0.00963111 | 0.43416247 | 0.24706947 | 0.32479848 | 0.41751459 | 0.43416247 | ZT0 | Absent_Koike2012 | Absent_Koike2012 |
| ENSMUSG00000048832 | Vps37c | 19 | 10763305 | 10788909 | + | 0.00050559 | 0.00996563 | 3.108156 | 2.22223375 | 2.574883 | 2.39955625 | 3.108156 | ZT0 | Absent_Koike2012 | Absent_Koike2012 |
| ENSMUSG00000033917 | Gde1 | 7 | 125832059 | 125849292 | - | 6.21E-10 | 2.33E-06 | 19.7178853 | 14.4237364 | 31.2716349 | 31.1072988 | 31.2716349 | ZT12 | Present_Koike2012 | Present_Koike2012 |
| ENSMUSG00000021775 | Nr1d2 | 14 | 19036568 | 19071641 | - | 6.42E-10 | 2.33E-06 | 1.22758566 | 11.3221774 | 13.759182 | 4.48466367 | 13.759182 | ZT12 | Present_Koike2012 | Present_Koike2012 |
| ENSMUSG00000059824 | Dbp | 7 | 52960458 | 52965525 | + | 7.89E-10 | 2.51E-06 | 0.13992848 | 6.98034154 | 8.9046088 | 1.02941633 | 8.9046088 | ZT12 | Present_Koike2012 | Present_Koike2012 |
| ENSMUSG00000033161 | Atp1a1 | 3 | 101380142 | 101408586 | - | 9.76E-10 | 2.76E-06 | 47.6824779 | 48.441279 | 69.6039785 | 67.46604 | 69.6039785 | ZT12 | Present_Koike2012 | Present_Koike2012 |
| ENSMUSG00000021814 | Anxa7 | 14 | 21274483 | 21299355 | - | 1.42E-09 | 3.61E-06 | 18.2429284 | 20.9093816 | 31.410074 | 21.4509163 | 31.410074 | ZT12 | Present_Koike2012 | Present_Koike2012 |
| ENSMUSG00000020572 | Nampt | 12 | 33505200 | 33538234 | + | 9.48E-09 | 1.61E-05 | 10.1382162 | 9.57274355 | 37.2508853 | 23.2560552 | 37.2508853 | ZT12 | Present_Koike2012 | Present_Koike2012 |
| ENSMUSG00000028957 | Per3 | 4 | 150377761 | 150418774 | - | 6.10E-08 | 4.67E-05 | 0.15540634 | 0.76661042 | 2.74713908 | 1.44825597 | 2.74713908 | ZT12 | Present_Koike2012 | Present_Koike2012 |
| ENSMUSG00000022295 | Atp6v1c1 | 15 | 38591659 | 38622198 | + | 6.41E-08 | 4.67E-05 | 10.6031543 | 12.3749485 | 15.2225375 | 10.4526655 | 15.2225375 | ZT12 | Present_Koike2012 | Absent_Koike2012 |
| ENSMUSG00000044345 | Marveld1 | 19 | 42221890 | 42226185 | + | 1.06E-07 | 6.59E-05 | 4.95870525 | 3.51299433 | 7.81424839 | 7.26513391 | 7.81424839 | ZT12 | Present_Koike2012 | Present_Koike2012 |
| ENSMUSG00000009378 | Slc16a12 | 19 | 34742893 | 34821779 | - | 1.13E-07 | 6.84E-05 | 6.47799465 | 5.7939269 | 14.6340935 | 10.2533507 | 14.6340935 | ZT12 | Present_Koike2012 | Present_Koike2012 |
| ENSMUSG00000031016 | Wee1 | 7 | 117265864 | 117285626 | + | 1.27E-07 | 7.53E-05 | 0.829927 | 1.14887175 | 4.779791 | 3.09295375 | 4.779791 | ZT12 | Present_Koike2012 | Present_Koike2012 |
| ENSMUSG00000054321 | Taf4b | 18 | 14941754 | 15058868 | + | 1.47E-07 | 8.01E-05 | 0.41153667 | 0.52111763 | 0.64063123 | 0.33061836 | 0.64063123 | ZT12 | Absent_Koike2012 | Present_Koike2012 |
| ENSMUSG00000040659 | Efhd2 | 4 | 141414057 | 141430835 | - | 1.75E-07 | 8.01E-05 | 4.82692825 | 5.32779975 | 11.713613 | 7.507126 | 11.713613 | ZT12 | Present_Koike2012 | Present_Koike2012 |
| ENSMUSG00000032788 | Pdxk | 10 | 77899489 | 77927720 | - | 1.76E-07 | 8.01E-05 | 9.13227324 | 6.67257317 | 13.4235559 | 13.3764771 | 13.4235559 | ZT12 | Present_Koike2012 | Absent_Koike2012 |
| ENSMUSG00000025981 | Coq10b | 1 | 55109614 | 55129545 | + | 1.65E-07 | 8.01E-05 | 8.85807955 | 10.5499711 | 32.505714 | 11.6805679 | 32.505714 | ZT12 | Present_Koike2012 | Present_Koike2012 |
| ENSMUSG00000035828 | Pim3 | 15 | 88692616 | 88696150 | + | 1.50E-07 | 8.01E-05 | 4.74977147 | 10.1358792 | 18.4570012 | 7.99512913 | 18.4570012 | ZT12 | Absent_Koike2012 | Absent_Koike2012 |
| ENSMUSG00000024066 | Xdh | 17 | 74233248 | 74299522 | - | 1.71E-07 | 8.01E-05 | 26.2767427 | 27.8312969 | 39.5991012 | 32.0511034 | 39.5991012 | ZT12 | Present_Koike2012 | Present_Koike2012 |
| ENSMUSG00000027253 | Lrp4 | 2 | 91297668 | 91354058 | + | 1.63E-07 | 8.01E-05 | 2.25388558 | 1.97780319 | 3.62150904 | 3.48192605 | 3.62150904 | ZT12 | Present_Koike2012 | Absent_Koike2012 |
| ENSMUSG00000028792 | Ak2 | 4 | 128669202 | 128688773 | + | 1.91E-07 | 8.15E-05 | 55.6541155 | 62.8735399 | 99.9199666 | 70.509041 | 99.9199666 | ZT12 | Absent_Koike2012 | Absent_Koike2012 |
| ENSMUSG00000026064 | Ptp4a1 | 1 | 30997148 | 31006615 | - | 1.95E-07 | 8.15E-05 | 12.864789 | 12.3880044 | 23.0934326 | 18.5037814 | 23.0934326 | ZT12 | Present_Koike2012 | Absent_Koike2012 |
| ENSMUSG00000038116 | Phf20 | 2 | 156022202 | 156135688 | + | 2.11E-07 | 8.41E-05 | 0.93588819 | 0.9096287 | 1.27706417 | 1.01675248 | 1.27706417 | ZT12 | Absent_Koike2012 | Absent_Koike2012 |
| ENSMUSG00000052566 | Hook2 | 8 | 87514494 | 87527263 | + | 2.28E-07 | 8.42E-05 | 3.28726619 | 3.71752066 | 6.12780219 | 3.26844807 | 6.12780219 | ZT12 | Present_Koike2012 | Absent_Koike2012 |
| ENSMUSG00000038550 | Gm129 | 3 | 95682426 | 95686174 | - | 2.77E-07 | 9.41E-05 | 0.06355789 | 0.58877772 | 4.15116184 | 0.61517142 | 4.15116184 | ZT12 | Present_Koike2012 | Absent_Koike2012 |
| ENSMUSG00000022462 | Slc38a2 | 15 | 96517825 | 96530129 | - | 3.05E-07 | 0.00010066 | 12.5198968 | 22.1275238 | 34.0685395 | 13.2131988 | 34.0685395 | ZT12 | Present_Koike2012 | Absent_Koike2012 |
| ENSMUSG00000034617 | Mtrr | 13 | 68699657 | 68720998 | - | 3.68E-07 | 0.00011012 | 1.41957633 | 0.81915631 | 1.64690861 | 1.39120871 | 1.64690861 | ZT12 | Absent_Koike2012 | Present_Koike2012 |
| ENSMUSG00000033060 | Lmo7 | 14 | 102129145 | 102349230 | + | 3.65E-07 | 0.00011012 | 0.86928458 | 1.0372942 | 1.62044415 | 1.25467938 | 1.62044415 | ZT12 | Present_Koike2012 | Present_Koike2012 |
| ENSMUSG00000025078 | Nhlrc2 | 19 | 56622751 | 56673336 | + | 3.68E-07 | 0.00011012 | 3.65461125 | 3.719927 | 6.410576 | 5.00297539 | 6.410576 | ZT12 | Present_Koike2012 | Present_Koike2012 |
| ENSMUSG00000023307 | March5 | 19 | 37282033 | 37296629 | + | 3.86E-07 | 0.00011155 | 16.0433522 | 17.9078881 | 21.0892025 | 15.7409335 | 21.0892025 | ZT12 | Present_Koike2012 | Absent_Koike2012 |
| ENSMUSG00000038023 | Atp6v0a2 | 5 | 125079673 | 125204824 | + | 4.04E-07 | 0.0001138 | 6.3597265 | 5.24726725 | 7.05004325 | 6.673545 | 7.05004325 | ZT12 | Absent_Koike2012 | Absent_Koike2012 |
| ENSMUSG00000004018 | Fancl | 11 | 26286135 | 26371876 | + | 4.17E-07 | 0.00011525 | 1.16179707 | 1.5149775 | 1.61069385 | 1.25237616 | 1.61069385 | ZT12 | Absent_Koike2012 | Absent_Koike2012 |
| ENSMUSG00000024085 | Man2a1 | 17 | 64950076 | 65104450 | + | 4.25E-07 | 0.00011627 | 33.5057006 | 36.6324622 | 52.2187526 | 47.4400854 | 52.2187526 | ZT12 | Present_Koike2012 | Absent_Koike2012 |
| ENSMUSG00000024190 | Dusp1 | 17 | 26642535 | 26645464 | - | 4.98E-07 | 0.00012943 | 2.42509655 | 1.99287042 | 7.06146309 | 3.40335727 | 7.06146309 | ZT12 | Absent_Koike2012 | Absent_Koike2012 |
| ENSMUSG00000002068 | Ccne1 | 7 | 38883003 | 38892553 | - | 5.40E-07 | 0.00013341 | 0.22119213 | 0.34334276 | 1.33424314 | 0.23450967 | 1.33424314 | ZT12 | Present_Koike2012 | Absent_Koike2012 |
| ENSMUSG00000073139 | BC023829 | X | 67712928 | 67730355 | - | 5.71E-07 | 0.00013341 | 3.201178 | 3.09461193 | 4.56649099 | 4.100488 | 4.56649099 | ZT12 | Absent_Koike2012 | Absent_Koike2012 |
| ENSMUSG00000022139 | Mbnl2 | 14 | 120674891 | 120830919 | + | 5.19E-07 | 0.00013341 | 6.83933045 | 5.6402722 | 10.1699743 | 8.38203528 | 10.1699743 | ZT12 | Absent_Koike2012 | Present_Koike2012 |
| ENSMUSG00000027433 | Xrn2 | 2 | 146838732 | 146903736 | + | 5.40E-07 | 0.00013341 | 9.28343937 | 11.7776929 | 12.0636999 | 8.26573836 | 12.0636999 | ZT12 | Absent_Koike2012 | Present_Koike2012 |
| ENSMUSG00000038481 | Cdk19 | 10 | 40069114 | 40203624 | + | 5.91E-07 | 0.00013663 | 0.9070275 | 1.37097725 | 1.46476167 | 1.135519 | 1.46476167 | ZT12 | Absent_Koike2012 | Absent_Koike2012 |
| ENSMUSG00000042476 | Abcb4 | 5 | 8893717 | 8959225 | + | 6.17E-07 | 0.00014128 | 40.872058 | 49.8482166 | 51.3666583 | 33.859973 | 51.3666583 | ZT12 | Present_Koike2012 | Absent_Koike2012 |
| ENSMUSG00000091905 | Gm6395 | 12 | 65887298 | 65888021 | + | 6.59E-07 | 0.00014583 | 11.9216928 | 12.2926153 | 18.4048835 | 14.0123633 | 18.4048835 | ZT12 | Absent_Koike2012 | Absent_Koike2012 |
| ENSMUSG00000066357 | Wdr6 | 9 | 108474644 | 108481070 | - | 6.70E-07 | 0.00014699 | 3.870558 | 5.26627926 | 6.14866191 | 3.4313949 | 6.14866191 | ZT12 | Present_Koike2012 | Absent_Koike2012 |
| ENSMUSG00000028878 | Fam76a | 4 | 132455128 | 132478473 | - | 7.11E-07 | 0.00015123 | 2.92808941 | 3.75583067 | 5.07284594 | 3.30622412 | 5.07284594 | ZT12 | Present_Koike2012 | Absent_Koike2012 |
| ENSMUSG00000020733 | Slc9a3r1 | 11 | 115024655 | 115042495 | + | 7.25E-07 | 0.00015123 | 48.1995413 | 52.359916 | 73.0071868 | 67.5312003 | 73.0071868 | ZT12 | Absent_Koike2012 | Absent_Koike2012 |
| ENSMUSG00000039917 | Rhbdd2 | 5 | 136108488 | 136122318 | + | 8.20E-07 | 0.0001633 | 0.70727297 | 0.8045685 | 1.39811652 | 1.204351 | 1.39811652 | ZT12 | Present_Koike2012 | Absent_Koike2012 |
| ENSMUSG00000071551 | Akr1c19 | 13 | 4227946 | 4247927 | + | 8.26E-07 | 0.0001633 | 35.8096003 | 31.3567835 | 76.4573003 | 56.4887034 | 76.4573003 | ZT12 | Present_Koike2012 | Present_Koike2012 |
| ENSMUSG00000025077 | Dclre1a | 19 | 56603649 | 56622516 | - | 8.19E-07 | 0.0001633 | 1.94789114 | 2.26507459 | 3.44386524 | 2.71210306 | 3.44386524 | ZT12 | Present_Koike2012 | Absent_Koike2012 |
| ENSMUSG00000020131 | Pcsk4 | 10 | 79784028 | 79792243 | - | 8.46E-07 | 0.00016551 | 0.38596965 | 1.21344398 | 2.55881366 | 1.09080066 | 2.55881366 | ZT12 | Present_Koike2012 | Present_Koike2012 |
| ENSMUSG00000001436 | Slc19a1 | 10 | 76494986 | 76523747 | + | 9.58E-07 | 0.00017923 | 0.64946599 | 0.68778868 | 1.18233817 | 0.6697676 | 1.18233817 | ZT12 | Present_Koike2012 | Present_Koike2012 |
| ENSMUSG00000051671 | 1810063B05I | 8 | 128946401 | 128949334 | + | 1.04E-06 | 0.00018856 | 6.6526552 | 5.2598213 | 8.31618542 | 7.81383076 | 8.31618542 | ZT12 | Absent_Koike2012 | Absent_Koike2012 |
| ENSMUSG00000029578 | Wipi2 | 5 | 143105496 | 143146542 | + | 1.06E-06 | 0.00018954 | 7.82740799 | 8.61877571 | 8.69642262 | 6.87432511 | 8.69642262 | ZT12 | Absent_Koike2012 | Absent_Koike2012 |
| ENSMUSG00000017774 | Myo1c | 11 | 75464006 | 75488134 | + | 1.14E-06 | 0.00019624 | 5.03324012 | 4.8281504 | 6.4043151 | 5.47725717 | 6.4043151 | ZT12 | Absent_Koike2012 | Absent_Koike2012 |
| ENSMUSG00000015869 | Prpsap1 | 11 | 116332159 | 116355516 | - | 1.14E-06 | 0.00019624 | 6.40803197 | 6.57523128 | 8.41009484 | 5.79529364 | 8.41009484 | ZT12 | Absent_Koike2012 | Absent_Koike2012 |
| ENSMUSG00000020593 | Lpin1 | 12 | 16542475 | 16596576 | - | 1.16E-06 | 0.00019624 | 2.29119615 | 4.33564027 | 36.594845 | 6.85555062 | 36.594845 | ZT12 | Present_Koike2012 | Present_Koike2012 |
| ENSMUSG00000023150 | Ivns1abp | 1 | 153191628 | 153211575 | + | 1.16E-06 | 0.00019624 | 18.1441568 | 14.3047368 | 26.4979029 | 21.6982638 | 26.4979029 | ZT12 | Present_Koike2012 | Present_Koike2012 |
| ENSMUSG00000044749 | Abca6 | 11 | 110038134 | 110113090 | - | 1.20E-06 | 0.00019797 | 34.6888024 | 44.5245902 | 50.6347444 | 36.0938106 | 50.6347444 | ZT12 | Present_Koike2012 | Absent_Koike2012 |
| ENSMUSG00000042510 | AA986860 | 1 | 132628553 | 132641198 | + | 1.25E-06 | 0.00020487 | 0.285131 | 0.1400705 | 1.188622 | 0.32130225 | 1.188622 | ZT12 | Absent_Koike2012 | Absent_Koike2012 |

| | | | | | | | | | | | | | | | |
|---|---|---|---|---|---|---|---|---|---|---|---|---|---|---|---|
| **ENSMUSG00000022517** | Mgrn1 | 16 | 4886252 | 4938296 | + | 1.30E-06 | 0.0002057 | 9.59525732 | 12.0608496 | 12.1402433 | 10.0285269 | 12.1402433 | ZT12 | Present_Koike2012 | Absent_Koike2012 |
| **ENSMUSG00000005615** | Pcyt1a | 16 | 32431007 | 32475156 | + | 1.27E-06 | 0.0002057 | 5.11745751 | 4.8056109 | 6.2917586 | 6.10136182 | 6.2917586 | ZT12 | Absent_Koike2012 | Present_Koike2012 |
| **ENSMUSG00000084839** | Gm14097 | 2 | 132520168 | 132524259 | - | 1.28E-06 | 0.0002057 | 1.43478275 | 6.4824045 | 10.3183505 | 2.5698985 | 10.3183505 | ZT12 | Absent_Koike2012 | Absent_Koike2012 |
| **ENSMUSG00000031015** | Swap70 | 7 | 117365217 | 117427017 | + | 1.54E-06 | 0.00023367 | 1.91614225 | 2.16573775 | 2.74714875 | 2.32464325 | 2.74714875 | ZT12 | Absent_Koike2012 | Present_Koike2012 |
| **ENSMUSG00000014496** | Ankrd28 | 14 | 32513201 | 32643601 | - | 1.59E-06 | 0.0002381 | 5.43972182 | 5.81452015 | 7.28411354 | 6.36203412 | 7.28411354 | ZT12 | Present_Koike2012 | Present_Koike2012 |
| **ENSMUSG00000020649** | Rrm2 | 12 | 25393106 | 25399011 | + | 1.69E-06 | 0.00024774 | 1.29935152 | 0.95325939 | 3.69118714 | 1.92187977 | 3.69118714 | ZT12 | Absent_Koike2012 | Absent_Koike2012 |
| **ENSMUSG00000037580** | Gch1 | 14 | 47773570 | 47809088 | - | 1.69E-06 | 0.00024774 | 32.1231438 | 46.7070685 | 46.881795 | 25.0133725 | 46.881795 | ZT12 | Present_Koike2012 | Absent_Koike2012 |
| **ENSMUSG00000024222** | Fkbp5 | 17 | 28536040 | 28654469 | - | 1.81E-06 | 0.00025798 | 1.71379539 | 2.08033475 | 5.20974381 | 3.52625276 | 5.20974381 | ZT12 | Present_Koike2012 | Present_Koike2012 |
| **ENSMUSG00000019917** | sep-10 | 10 | 58604398 | 58684595 | - | 1.86E-06 | 0.00026234 | 4.50183613 | 5.44403248 | 5.47631254 | 4.5505797 | 5.47631254 | ZT12 | Absent_Koike2012 | Absent_Koike2012 |
| **ENSMUSG00000034254** | Agpat1 | 17 | 34741207 | 34750394 | + | 1.88E-06 | 0.00026431 | 1.34687934 | 1.01780222 | 1.52145228 | 1.47858715 | 1.52145228 | ZT12 | Absent_Koike2012 | Absent_Koike2012 |
| **ENSMUSG00000046404** | Yod1 | 1 | 132613904 | 132618632 | + | 2.12E-06 | 0.00029178 | 3.47790775 | 2.93033375 | 5.81753225 | 3.08343475 | 5.81753225 | ZT12 | Present_Koike2012 | Absent_Koike2012 |
| **ENSMUSG00000058729** | Lin9 | 1 | 182571498 | 182620816 | + | 2.27E-06 | 0.00030922 | 0.61737798 | 0.93913437 | 1.17754957 | 0.5572416 | 1.17754957 | ZT12 | Absent_Koike2012 | Absent_Koike2012 |
| **ENSMUSG00000020893** | Per1 | 11 | 68908719 | 68923462 | + | 2.29E-06 | 0.00030975 | 0.19774642 | 0.40888338 | 2.09966988 | 0.71312173 | 2.09966988 | ZT12 | Present_Koike2012 | Present_Koike2012 |
| **ENSMUSG00000021670** | Hmgcr | 13 | 97418922 | 97440891 | - | 2.36E-06 | 0.00031721 | 11.0723135 | 11.7602437 | 35.4194206 | 25.6176758 | 35.4194206 | ZT12 | Present_Koike2012 | Present_Koike2012 |
| **ENSMUSG00000006641** | Slc5a6 | 5 | 31338409 | 31351297 | - | 2.40E-06 | 0.00031817 | 0.97408027 | 1.00006711 | 2.65956991 | 1.08733564 | 2.65956991 | ZT12 | Present_Koike2012 | Absent_Koike2012 |
| **ENSMUSG00000005514** | Por | 5 | 136145903 | 136211196 | + | 2.38E-06 | 0.00031817 | 20.0344134 | 27.7990022 | 74.6496041 | 41.6845855 | 74.6496041 | ZT12 | Present_Koike2012 | Present_Koike2012 |
| **ENSMUSG00000066170** | E230001N04 | 17 | 28660238 | 28662292 | + | 2.43E-06 | 0.00032015 | 0.017731 | 0.0175915 | 0.1854275 | 0.052916 | 0.1854275 | ZT12 | Absent_Koike2012 | Absent_Koike2012 |
| **ENSMUSG00000028599** | Tnfrsf1b | 4 | 144803366 | 144836773 | - | 2.68E-06 | 0.00034578 | 3.42213925 | 4.14552203 | 6.39379522 | 4.11433649 | 6.39379522 | ZT12 | Absent_Koike2012 | Absent_Koike2012 |
| **ENSMUSG00000034329** | Brip1 | 11 | 85871638 | 86014695 | - | 2.75E-06 | 0.00035199 | 0.19931833 | 0.40384015 | 0.60862587 | 0.25052351 | 0.60862587 | ZT12 | Absent_Koike2012 | Absent_Koike2012 |
| **ENSMUSG00000048756** | Foxo3 | 10 | 41901647 | 41996561 | - | 2.91E-06 | 0.00036477 | 5.21557941 | 7.53126376 | 8.9414127 | 7.03123968 | 8.9414127 | ZT12 | Absent_Koike2012 | Present_Koike2012 |
| **ENSMUSG00000023977** | Ubr2 | 17 | 47065241 | 47147481 | - | 3.16E-06 | 0.00038817 | 6.63222427 | 6.9680926 | 10.237155 | 7.93448286 | 10.237155 | ZT12 | Present_Koike2012 | Present_Koike2012 |
| **ENSMUSG00000029250** | Polr2b | 5 | 77739172 | 77778349 | + | 3.22E-06 | 0.00039165 | 9.56703991 | 8.41085473 | 12.1720528 | 11.6939574 | 12.1720528 | ZT12 | Absent_Koike2012 | Present_Koike2012 |
| **ENSMUSG00000090622** | A930033H14 | 10 | 68672510 | 68675846 | - | 3.33E-06 | 0.0003969 | 0.1586725 | 0.207673 | 1.15370375 | 0.40141875 | 1.15370375 | ZT12 | Absent_Koike2012 | Absent_Koike2012 |
| **ENSMUSG00000055128** | Cgrrf1 | 14 | 47451809 | 47473868 | + | 3.34E-06 | 0.0003969 | 5.44635783 | 4.89372845 | 6.96276267 | 5.90250742 | 6.96276267 | ZT12 | Absent_Koike2012 | Absent_Koike2012 |
| **ENSMUSG00000024588** | Fech | 18 | 64616204 | 64648720 | - | 3.38E-06 | 0.00039952 | 29.7262579 | 37.3911841 | 39.9112917 | 31.2590001 | 39.9112917 | ZT12 | Present_Koike2012 | Absent_Koike2012 |
| **ENSMUSG00000000296** | Tpd52l1 | 10 | 31052186 | 31165727 | - | 3.52E-06 | 0.00041257 | 2.37678336 | 2.62621787 | 3.37045301 | 2.65362775 | 3.37045301 | ZT12 | Absent_Koike2012 | Absent_Koike2012 |
| **ENSMUSG00000024052** | Lpin2 | 17 | 71531900 | 71599157 | + | 3.59E-06 | 0.00041708 | 3.90670218 | 9.11309379 | 19.2312957 | 9.6010578 | 19.2312957 | ZT12 | Present_Koike2012 | Present_Koike2012 |
| **ENSMUSG00000089712** | Gm15889 | 8 | 96901821 | 96903522 | + | 3.66E-06 | 0.00042356 | 0.45009975 | 0.554047 | 1.81982775 | 1.462984 | 1.81982775 | ZT12 | Absent_Koike2012 | Absent_Koike2012 |
| **ENSMUSG00000025280** | Polr3a | 14 | 25267916 | 25306268 | - | 3.76E-06 | 0.0004315 | 3.53333325 | 3.340582 | 3.81255875 | 3.088128 | 3.81255875 | ZT12 | Absent_Koike2012 | Absent_Koike2012 |
| **ENSMUSG00000034667** | Xpot | 10 | 121024436 | 121063372 | - | 3.95E-06 | 0.00044725 | 10.630794 | 11.1664207 | 15.6715578 | 12.4388154 | 15.6715578 | ZT12 | Present_Koike2012 | Absent_Koike2012 |
| **ENSMUSG00000005225** | Plekha8 | 6 | 54545105 | 54595833 | + | 4.05E-06 | 0.00045035 | 5.64628601 | 3.87646074 | 6.74426545 | 6.11244017 | 6.74426545 | ZT12 | Present_Koike2012 | Present_Koike2012 |
| **ENSMUSG00000030747** | Dgat2 | 7 | 106302173 | 106331223 | - | 4.07E-06 | 0.00045035 | 135.981751 | 146.932129 | 199.307148 | 180.920181 | 199.307148 | ZT12 | Present_Koike2012 | Absent_Koike2012 |
| **ENSMUSG00000048007** | Timm8a1 | X | 131071795 | 131076404 | - | 4.19E-06 | 0.00045771 | 15.3945355 | 14.3924265 | 20.5550708 | 17.1684095 | 20.5550708 | ZT12 | Absent_Koike2012 | Present_Koike2012 |
| **ENSMUSG00000023122** | Sult1c2 | 17 | 53968962 | 53985664 | - | 4.17E-06 | 0.00045771 | 1.534964 | 2.4431405 | 2.56847125 | 1.4368495 | 2.56847125 | ZT12 | Absent_Koike2012 | Absent_Koike2012 |
| **ENSMUSG00000037606** | Osbpl5 | 7 | 150874667 | 150942890 | - | 4.26E-06 | 0.00046142 | 0.52104493 | 0.40309733 | 0.82359567 | 0.68159535 | 0.82359567 | ZT12 | Absent_Koike2012 | Absent_Koike2012 |
| **ENSMUSG00000033400** | Agl | 3 | 116442917 | 116511084 | - | 4.39E-06 | 0.0004674 | 3.68002305 | 4.34564164 | 5.61107151 | 4.23840994 | 5.61107151 | ZT12 | Present_Koike2012 | Absent_Koike2012 |
| **ENSMUSG00000030244** | Gys2 | 6 | 142371133 | 142421629 | - | 4.51E-06 | 0.00047761 | 33.4014637 | 46.4631525 | 90.6063571 | 66.3574094 | 90.6063571 | ZT12 | Present_Koike2012 | Absent_Koike2012 |
| **ENSMUSG00000049969** | Plekhf2 | 4 | 10915809 | 10935074 | - | 4.58E-06 | 0.00048067 | 4.50606171 | 5.57802623 | 5.76649294 | 4.92065931 | 5.76649294 | ZT12 | Absent_Koike2012 | Absent_Koike2012 |
| **ENSMUSG00000035354** | Uvrag | 7 | 106035253 | 106289654 | - | 4.59E-06 | 0.00048067 | 6.70370525 | 7.5871305 | 10.15955 | 7.75143425 | 10.15955 | ZT12 | Present_Koike2012 | Present_Koike2012 |
| **ENSMUSG00000060038** | Dhps | 8 | 87595656 | 87599061 | + | 4.66E-06 | 0.0004817 | 2.49448203 | 2.04085042 | 2.88276867 | 2.64244853 | 2.88276867 | ZT12 | Present_Koike2012 | Absent_Koike2012 |
| **ENSMUSG00000040170** | Fmo2 | 1 | 164804448 | 164828857 | - | 4.64E-06 | 0.0004817 | 1.2359664 | 0.96878713 | 2.67563462 | 2.5773089 | 2.67563462 | ZT12 | Present_Koike2012 | Present_Koike2012 |
| **ENSMUSG00000041891** | Lman1 | 18 | 66140408 | 66182234 | - | 4.93E-06 | 0.00048968 | 23.7172709 | 21.8281797 | 28.9741743 | 27.5569717 | 28.9741743 | ZT12 | Present_Koike2012 | Present_Koike2012 |
| **ENSMUSG00000027668** | Mfn1 | 3 | 32428387 | 32478161 | + | 4.94E-06 | 0.00048968 | 6.1426749 | 6.7466346 | 8.07803867 | 6.11624715 | 8.07803867 | ZT12 | Present_Koike2012 | Absent_Koike2012 |
| **ENSMUSG00000015143** | Actn1 | 12 | 81268529 | 81361358 | - | 5.19E-06 | 0.00050374 | 4.61929828 | 5.77699373 | 7.10460855 | 5.12362066 | 7.10460855 | ZT12 | Present_Koike2012 | Absent_Koike2012 |
| **ENSMUSG00000021500** | Ddx46 | 13 | 55736388 | 55782617 | + | 5.23E-06 | 0.00050374 | 3.1821441 | 4.39452574 | 4.43395653 | 2.64858057 | 4.43395653 | ZT12 | Absent_Koike2012 | Absent_Koike2012 |
| **ENSMUSG00000078941** | Taf9 | 13 | 101421298 | 101436368 | + | 5.86E-06 | 0.00054777 | 9.57560651 | 10.2902023 | 10.8357386 | 8.15440452 | 10.8357386 | ZT12 | Absent_Koike2012 | Absent_Koike2012 |
| **ENSMUSG00000017493** | Igfbp4 | 11 | 98902558 | 98915706 | + | 6.17E-06 | 0.00056666 | 133.27944 | 151.299549 | 193.202245 | 178.73078 | 193.202245 | ZT12 | Present_Koike2012 | Absent_Koike2012 |
| **ENSMUSG00000028479** | Gne | 4 | 44046947 | 44097049 | - | 6.21E-06 | 0.00056806 | 9.70573607 | 13.4413081 | 19.4640362 | 11.2085255 | 19.4640362 | ZT12 | Present_Koike2012 | Present_Koike2012 |
| **ENSMUSG00000064145** | Arih2 | 9 | 108505273 | 108551711 | - | 6.42E-06 | 0.00058347 | 7.83888749 | 9.53249351 | 10.0766235 | 7.13556207 | 10.0766235 | ZT12 | Absent_Koike2012 | Absent_Koike2012 |
| **ENSMUSG00000000708** | Kat2b | 17 | 53706186 | 53812045 | + | 6.75E-06 | 0.00060915 | 17.6026723 | 21.5168055 | 23.1357289 | 15.7485502 | 23.1357289 | ZT12 | Present_Koike2012 | Absent_Koike2012 |
| **ENSMUSG00000029804** | Herc3 | 6 | 58781459 | 58870392 | + | 6.86E-06 | 0.00061707 | 0.62529805 | 0.97413253 | 1.11716001 | 0.5775011 | 1.11716001 | ZT12 | Absent_Koike2012 | Absent_Koike2012 |
| **ENSMUSG00000020262** | Adarb1 | 10 | 76753471 | 76881015 | - | 7.06E-06 | 0.00063264 | 1.08730057 | 1.21981791 | 1.70494583 | 1.21755529 | 1.70494583 | ZT12 | Present_Koike2012 | Present_Koike2012 |
| **ENSMUSG00000026259** | Ngef | 1 | 89373409 | 89470445 | - | 7.14E-06 | 0.00063268 | 3.99723539 | 5.08025295 | 8.5599032 | 5.39003493 | 8.5599032 | ZT12 | Present_Koike2012 | Absent_Koike2012 |
| **ENSMUSG00000044026** | Tmem20 | 19 | 38470470 | 38480097 | + | 7.21E-06 | 0.00063372 | 6.075108 | 5.34794471 | 10.4252264 | 5.79137113 | 10.4252264 | ZT12 | Present_Koike2012 | Absent_Koike2012 |
| **ENSMUSG00000039914** | Coq10a | 10 | 127800153 | 127807088 | - | 7.40E-06 | 0.00064303 | 4.0772446 | 4.47206221 | 4.69801106 | 3.17250233 | 4.69801106 | ZT12 | Present_Koike2012 | Absent_Koike2012 |
| **ENSMUSG00000025139** | Tollip | 7 | 149060718 | 149104412 | - | 7.78E-06 | 0.00066916 | 8.18111662 | 9.07696879 | 10.886401 | 9.14133313 | 10.886401 | ZT12 | Absent_Koike2012 | Absent_Koike2012 |
| **ENSMUSG00000032596** | Uba7 | 9 | 107877836 | 107886391 | + | 7.83E-06 | 0.00066997 | 2.09558916 | 2.65229816 | 2.90103836 | 2.08563185 | 2.90103836 | ZT12 | Absent_Koike2012 | Absent_Koike2012 |
| **ENSMUSG00000015846** | Rxra | 2 | 27531960 | 27618477 | + | 7.85E-06 | 0.00066997 | 49.448891 | 58.7395232 | 63.1545245 | 47.113353 | 63.1545245 | ZT12 | Present_Koike2012 | Absent_Koike2012 |
| **ENSMUSG00000030662** | Ipo5 | 14 | 121310416 | 121347264 | + | 8.11E-06 | 0.00068033 | 10.0315618 | 10.1199562 | 13.99092 | 10.7368078 | 13.99092 | ZT12 | Absent_Koike2012 | Present_Koike2012 |
| **ENSMUSG00000003546** | Klc4 | 17 | 46767574 | 46782104 | - | 8.13E-06 | 0.00068033 | 14.2122128 | 12.7144833 | 17.2792713 | 16.0489244 | 17.2792713 | ZT12 | Absent_Koike2012 | Absent_Koike2012 |
| **ENSMUSG00000039050** | Osbpl2 | 2 | 179854011 | 179897385 | + | 8.49E-06 | 0.0007025 | 4.67388084 | 4.56493567 | 5.87906898 | 5.01775875 | 5.87906898 | ZT12 | Absent_Koike2012 | Absent_Koike2012 |
| **ENSMUSG00000023830** | Igf2r | 17 | 12875272 | 12962530 | - | 8.58E-06 | 0.00070511 | 7.68330782 | 8.54599729 | 9.58949639 | 7.62953688 | 9.58949639 | ZT12 | Absent_Koike2012 | Absent_Koike2012 |
| **ENSMUSG00000028452** | Vcp | 4 | 42992835 | 43013379 | - | 8.77E-06 | 0.00071011 | 55.3375831 | 59.0129151 | 62.0523453 | 55.5864663 | 62.0523453 | ZT12 | Absent_Koike2012 | Absent_Koike2012 |
| **ENSMUSG00000034610** | Zcchc11 | 4 | 108132031 | 108232026 | + | 8.73E-06 | 0.00071011 | 1.14067357 | 1.71435578 | 1.98399302 | 1.26293396 | 1.98399302 | ZT12 | Present_Koike2012 | Absent_Koike2012 |
| **ENSMUSG00000022299** | Slc25a32 | 15 | 38925737 | 38944262 | - | 8.79E-06 | 0.00071011 | 3.61082939 | 3.90435605 | 5.62922374 | 3.43192374 | 5.62922374 | ZT12 | Present_Koike2012 | Absent_Koike2012 |
| **ENSMUSG00000017950** | Hnf4a | 2 | 163332544 | 163398646 | + | 8.82E-06 | 0.00071011 | 89.3146546 | 81.0323049 | 109.245463 | 87.3899411 | 109.245463 | ZT12 | Absent_Koike2012 | Absent_Koike2012 |
| **ENSMUSG00000059714** | Flot1 | 17 | 35960175 | 35969736 | + | 8.91E-06 | 0.00071292 | 2.28402378 | 3.00806639 | 3.58778224 | 1.99905607 | 3.58778224 | ZT12 | Absent_Koike2012 | Absent_Koike2012 |
| **ENSMUSG00000029998** | Pcyox1 | 6 | 86336000 | 86347144 | - | 9.19E-06 | 0.00073064 | 28.6840319 | 25.5107975 | 36.0242657 | 33.7409608 | 36.0242657 | ZT12 | Absent_Koike2012 | Absent_Koike2012 |
| **ENSMUSG00000020250** | Txnrd1 | 10 | 82296696 | 82360469 | + | 9.58E-06 | 0.00075458 | 25.4954727 | 24.3878126 | 29.9865669 | 25.4903411 | 29.9865669 | ZT12 | Absent_Koike2012 | Absent_Koike2012 |

| | | | | | | | | | | | | | | | |
|---|---|---|---|---|---|---|---|---|---|---|---|---|---|---|---|
| ENSMUSG00000081535 | Gm15453 | 1 | 152112665 | 152113129 | - | 9.55E-06 | 0.00075458 | 2.4974155 | 4.62289075 | 7.13793925 | 4.5019785 | 7.13793925 | ZT12 | Absent_Koike2012 | Absent_Koike2012 |
| ENSMUSG00000030102 | Itpr1 | 6 | 108163112 | 108501103 | + | 9.75E-06 | 0.00076545 | 7.70545577 | 7.42236426 | 11.2455062 | 9.7548431 | 11.2455062 | ZT12 | Present_Koike2012 | Present_Koike2012 |
| ENSMUSG00000051435 | Fhad1 | 4 | 141446353 | 141570997 | - | 1.00E-05 | 0.0007797 | 0.11436799 | 0.1160393 | 0.21539792 | 0.21352995 | 0.21539792 | ZT12 | Absent_Koike2012 | Absent_Koike2012 |
| ENSMUSG00000000811 | Txnrd3 | 6 | 89593982 | 89625523 | + | 1.00E-05 | 0.0007797 | 2.14848007 | 2.67489375 | 2.99769638 | 1.96795321 | 2.99769638 | ZT12 | Present_Koike2012 | Absent_Koike2012 |
| ENSMUSG00000018736 | Ndel1 | 11 | 68634936 | 68685360 | - | 1.03E-05 | 0.00079259 | 4.43398102 | 4.67282249 | 7.04911829 | 5.01962571 | 7.04911829 | ZT12 | Present_Koike2012 | Present_Koike2012 |
| ENSMUSG00000066245 | Gm10156 | 7 | 115006489 | 115007619 | + | 1.08E-05 | 0.00081049 | 0.95135131 | 2.00633696 | 2.81709773 | 1.87285788 | 2.81709773 | ZT12 | Absent_Koike2012 | Absent_Koike2012 |
| ENSMUSG00000033530 | Ttc7b | 12 | 101538981 | 101759032 | - | 1.08E-05 | 0.00081049 | 3.50740325 | 3.21145675 | 4.75534175 | 4.365017 | 4.75534175 | ZT12 | Absent_Koike2012 | Present_Koike2012 |
| ENSMUSG00000026131 | Dst | 1 | 33965125 | 34365495 | + | 1.15E-05 | 0.00084945 | 4.08761188 | 4.05581852 | 5.80372593 | 5.35110405 | 5.80372593 | ZT12 | Present_Koike2012 | Present_Koike2012 |
| ENSMUSG00000068739 | Sars | 3 | 108227783 | 108248127 | - | 1.17E-05 | 0.00085897 | 8.81475064 | 9.0404987 | 9.33651462 | 7.67399343 | 9.33651462 | ZT12 | Absent_Koike2012 | Absent_Koike2012 |
| ENSMUSG00000032640 | Chsy1 | 7 | 73254401 | 73318675 | + | 1.18E-05 | 0.00086223 | 0.60142934 | 0.61476192 | 0.83001061 | 0.52805725 | 0.83001061 | ZT12 | Present_Koike2012 | Present_Koike2012 |
| ENSMUSG00000018566 | Slc2a4 | 11 | 69756045 | 69761690 | - | 1.18E-05 | 0.00086534 | 0.03591969 | 0.02547168 | 0.64898913 | 0.3067966 | 0.64898913 | ZT12 | Absent_Koike2012 | Absent_Koike2012 |
| ENSMUSG00000026107 | Obfc2a | 1 | 51526342 | 51535243 | - | 1.21E-05 | 0.00087818 | 3.7485232 | 3.51682464 | 5.81858747 | 4.09224339 | 5.81858747 | ZT12 | Present_Koike2012 | Present_Koike2012 |
| ENSMUSG00000074622 | Mafb | 2 | 160189413 | 160192801 | - | 1.21E-05 | 0.00087818 | 2.49262525 | 4.7743915 | 10.567605 | 4.407421 | 10.567605 | ZT12 | Present_Koike2012 | Absent_Koike2012 |
| ENSMUSG00000056612 | Ppp1r14b | 19 | 7049538 | 7051814 | + | 1.22E-05 | 0.00087853 | 10.2621943 | 7.06782175 | 11.5588863 | 11.0145125 | 11.5588863 | ZT12 | Absent_Koike2012 | Absent_Koike2012 |
| ENSMUSG00000035992 | Fnip1 | 11 | 54251701 | 54331737 | + | 1.22E-05 | 0.00087909 | 4.92564599 | 7.45888298 | 7.51616692 | 4.99394848 | 7.51616692 | ZT12 | Present_Koike2012 | Present_Koike2012 |
| ENSMUSG00000029246 | Ppat | 5 | 77342274 | 77380603 | - | 1.23E-05 | 0.00087967 | 4.45358315 | 4.00655265 | 6.73803776 | 3.99907305 | 6.73803776 | ZT12 | Present_Koike2012 | Present_Koike2012 |
| ENSMUSG00000006442 | Srm | 4 | 147965612 | 147969102 | + | 1.24E-05 | 0.00088809 | 9.61438093 | 4.80161566 | 12.3069246 | 11.4082398 | 12.3069246 | ZT12 | Absent_Koike2012 | Absent_Koike2012 |
| ENSMUSG00000032435 | Dync1li1 | 9 | 114597908 | 114633420 | + | 1.25E-05 | 0.00088834 | 5.51151094 | 6.02438369 | 7.12850114 | 5.00736637 | 7.12850114 | ZT12 | Absent_Koike2012 | Absent_Koike2012 |
| ENSMUSG00000025194 | Abcc2 | 19 | 43856682 | 43912441 | + | 1.28E-05 | 0.00090215 | 29.8804976 | 40.4829952 | 53.2427932 | 32.2006972 | 53.2427932 | ZT12 | Present_Koike2012 | Present_Koike2012 |
| ENSMUSG00000078566 | Bnip3 | 7 | 146082519 | 146101202 | - | 1.30E-05 | 0.00091401 | 39.1758728 | 47.0444453 | 76.6111301 | 60.2411983 | 76.6111301 | ZT12 | Present_Koike2012 | Present_Koike2012 |
| ENSMUSG00000037062 | Sh3glb1 | 3 | 144347972 | 144383287 | - | 1.31E-05 | 0.00091581 | 8.52318908 | 8.45840167 | 10.3721196 | 9.36490248 | 10.3721196 | ZT12 | Absent_Koike2012 | Absent_Koike2012 |
| ENSMUSG00000070738 | Dgkd | 1 | 89749862 | 89841064 | + | 1.34E-05 | 0.00092324 | 3.06018536 | 3.37653504 | 3.73500857 | 2.86482721 | 3.73500857 | ZT12 | Present_Koike2012 | Absent_Koike2012 |
| ENSMUSG00000027016 | Zfp385b | 2 | 77248691 | 77657696 | - | 1.35E-05 | 0.0009312 | 0.55563067 | 0.81220166 | 1.17391062 | 0.81097328 | 1.17391062 | ZT12 | Absent_Koike2012 | Present_Koike2012 |
| ENSMUSG00000034203 | Chchd4 | 6 | 91414270 | 91423417 | - | 1.36E-05 | 0.00093173 | 7.9280315 | 5.553895 | 8.80511775 | 6.6256215 | 8.80511775 | ZT12 | Absent_Koike2012 | Absent_Koike2012 |
| ENSMUSG00000078905 | Gm14393 | 2 | 174887050 | 174893282 | - | 1.37E-05 | 0.00093173 | 1.95717806 | 1.68455166 | 2.41445509 | 2.33484622 | 2.41445509 | ZT12 | Absent_Koike2012 | Absent_Koike2012 |
| ENSMUSG00000042622 | Maff | 15 | 79177971 | 79189502 | + | 1.38E-05 | 0.00093776 | 0.25785726 | 1.11962233 | 1.25286919 | 0.42619898 | 1.25286919 | ZT12 | Present_Koike2012 | Absent_Koike2012 |
| ENSMUSG00000032010 | Usp2 | 9 | 43875104 | 43903710 | + | 1.40E-05 | 0.00094206 | 0.54440129 | 0.51436572 | 4.47945242 | 1.71142138 | 4.47945242 | ZT12 | Present_Koike2012 | Present_Koike2012 |
| ENSMUSG00000042246 | Tmc7 | 7 | 125679357 | 125728250 | - | 1.42E-05 | 0.0009538 | 0.61253515 | 0.57674268 | 1.4170063 | 0.95099541 | 1.4170063 | ZT12 | Absent_Koike2012 | Absent_Koike2012 |
| ENSMUSG00000032528 | Vipr1 | 9 | 121551834 | 121582072 | + | 1.43E-05 | 0.0009538 | 5.769915 | 5.31081301 | 6.30024675 | 5.95924005 | 6.30024675 | ZT12 | Absent_Koike2012 | Absent_Koike2012 |
| ENSMUSG00000056501 | Cebpb | 2 | 167514415 | 167515918 | + | 1.43E-05 | 0.0009538 | 8.57316075 | 11.9280503 | 25.1266043 | 15.2442578 | 25.1266043 | ZT12 | Present_Koike2012 | Absent_Koike2012 |
| ENSMUSG00000000056 | Narf | 11 | 121098567 | 121117170 | + | 1.57E-05 | 0.00101283 | 4.64502399 | 5.32768627 | 7.17905238 | 4.56956541 | 7.17905238 | ZT12 | Present_Koike2012 | Absent_Koike2012 |
| ENSMUSG00000052040 | Klf13 | 7 | 71031237 | 71083801 | - | 1.60E-05 | 0.0010258 | 3.6648728 | 4.42962248 | 8.650653 | 4.70653455 | 8.650653 | ZT12 | Present_Koike2012 | Present_Koike2012 |
| ENSMUSG00000045216 | Hs6st1 | 1 | 36125245 | 36163291 | + | 1.64E-05 | 0.00104907 | 16.8536073 | 16.7438063 | 24.254078 | 16.2694235 | 24.254078 | ZT12 | Present_Koike2012 | Absent_Koike2012 |
| ENSMUSG00000030946 | Lhpp | 7 | 139802321 | 139898103 | + | 1.67E-05 | 0.00105721 | 8.37300014 | 11.6018368 | 13.0374161 | 8.33799101 | 13.0374161 | ZT12 | Present_Koike2012 | Absent_Koike2012 |
| ENSMUSG00000035086 | Becn1 | 11 | 101147266 | 101163600 | - | 1.70E-05 | 0.00106931 | 3.8257023 | 3.76883917 | 4.45277333 | 4.02353454 | 4.45277333 | ZT12 | Absent_Koike2012 | Absent_Koike2012 |
| ENSMUSG00000030729 | Pgm2l1 | 7 | 107375904 | 107427378 | + | 1.77E-05 | 0.0010999 | 0.67604616 | 0.5235071 | 1.12747101 | 0.87807071 | 1.12747101 | ZT12 | Present_Koike2012 | Present_Koike2012 |
| ENSMUSG00000029198 | Grpel1 | 5 | 36807647 | 36816729 | + | 1.87E-05 | 0.00113764 | 19.7485915 | 20.6136858 | 21.7147008 | 18.8215433 | 21.7147008 | ZT12 | Present_Koike2012 | Absent_Koike2012 |
| ENSMUSG00000031700 | Gpt2 | 8 | 88016475 | 88051459 | + | 1.86E-05 | 0.00113764 | 34.4042068 | 29.3859644 | 51.1260793 | 43.5462398 | 51.1260793 | ZT12 | Present_Koike2012 | Absent_Koike2012 |
| ENSMUSG00000028826 | Tmem57 | 4 | 134358674 | 134409260 | - | 1.89E-05 | 0.00114238 | 5.2270208 | 4.68533069 | 7.5670924 | 6.83691614 | 7.5670924 | ZT12 | Present_Koike2012 | Present_Koike2012 |
| ENSMUSG00000024955 | Esrra | 19 | 6985467 | 6996303 | - | 1.89E-05 | 0.00114238 | 6.80572869 | 5.10141294 | 7.33762716 | 7.14928093 | 7.33762716 | ZT12 | Present_Koike2012 | Present_Koike2012 |
| ENSMUSG00000029287 | Tgfbr3 | 5 | 107535589 | 107718648 | - | 1.98E-05 | 0.00118511 | 2.02661842 | 2.60678586 | 2.64803441 | 2.11482356 | 2.64803441 | ZT12 | Present_Koike2012 | Absent_Koike2012 |
| ENSMUSG00000050323 | 2310030N02 | 4 | 10978192 | 11003352 | - | 2.10E-05 | 0.00124109 | 3.61407424 | 3.64798645 | 4.54492605 | 4.41411406 | 4.54492605 | ZT12 | Absent_Koike2012 | Absent_Koike2012 |
| ENSMUSG00000029426 | Scarb2 | 5 | 92872899 | 92934634 | - | 2.13E-05 | 0.00125515 | 56.211087 | 54.5022338 | 70.0164415 | 61.6450805 | 70.0164415 | ZT12 | Absent_Koike2012 | Absent_Koike2012 |
| ENSMUSG00000050627 | Gpd1l | 9 | 114808459 | 114843105 | - | 2.17E-05 | 0.00127759 | 4.42588695 | 4.33457868 | 6.44935725 | 5.45017796 | 6.44935725 | ZT12 | Present_Koike2012 | Present_Koike2012 |
| ENSMUSG00000028962 | Slc4a2 | 5 | 23929655 | 23946768 | + | 2.20E-05 | 0.00128653 | 3.0203652 | 2.92811118 | 3.88214892 | 3.10996169 | 3.88214892 | ZT12 | Absent_Koike2012 | Absent_Koike2012 |
| ENSMUSG00000062209 | Erbb4 | 1 | 68078760 | 69154633 | - | 2.25E-05 | 0.00130321 | 0.4035887 | 0.59828871 | 0.65529869 | 0.13355313 | 0.65529869 | ZT12 | Present_Koike2012 | Absent_Koike2012 |
| ENSMUSG00000021969 | Zdhhc20 | 14 | 58451539 | 58509099 | - | 2.25E-05 | 0.00130321 | 2.23656773 | 2.82636086 | 2.92201551 | 1.95129494 | 2.92201551 | ZT12 | Present_Koike2012 | Absent_Koike2012 |
| ENSMUSG00000037885 | Stk35 | 2 | 129626253 | 129658023 | + | 2.27E-05 | 0.00130838 | 1.30228669 | 1.29413946 | 2.27004084 | 1.62468574 | 2.27004084 | ZT12 | Present_Koike2012 | Present_Koike2012 |
| ENSMUSG00000053664 | Uck2 | 1 | 169153958 | 169154050 | - | 2.28E-05 | 0.00130889 | 1.9965575 | 1.27949525 | 2.608322 | 2.3453025 | 2.608322 | ZT12 | Absent_Koike2012 | Absent_Koike2012 |
| ENSMUSG00000041147 | Brca2 | 5 | 151325205 | 151372321 | + | 2.30E-05 | 0.00130891 | 0.24957775 | 0.397782 | 0.610768 | 0.35288275 | 0.610768 | ZT12 | Present_Koike2012 | Absent_Koike2012 |
| ENSMUSG00000010064 | Slc38a3 | 9 | 107552970 | 107569705 | - | 2.29E-05 | 0.00130891 | 142.474924 | 272.707532 | 308.057982 | 159.832843 | 308.057982 | ZT12 | Present_Koike2012 | Absent_Koike2012 |
| ENSMUSG00000040022 | Rab11fip2 | 19 | 59978787 | 60019557 | - | 2.35E-05 | 0.00132639 | 1.10145448 | 1.20089724 | 1.63504489 | 1.48245177 | 1.63504489 | ZT12 | Absent_Koike2012 | Absent_Koike2012 |
| ENSMUSG00000023846 | Riok2 | 17 | 17511296 | 17531862 | + | 2.36E-05 | 0.00132739 | 4.23629805 | 3.4350751 | 5.94521368 | 4.56532447 | 5.94521368 | ZT12 | Present_Koike2012 | Present_Koike2012 |
| ENSMUSG00000037847 | BC016495 | 19 | 18706440 | 18726684 | + | 2.37E-05 | 0.00132961 | 4.22258956 | 3.47624028 | 7.99467888 | 5.48715708 | 7.99467888 | ZT12 | Present_Koike2012 | Present_Koike2012 |
| ENSMUSG00000075511 | 1700001L05I | 15 | 83187990 | 83197707 | - | 2.41E-05 | 0.00134251 | 0.90040875 | 1.47815425 | 2.4521275 | 1.378819 | 2.4521275 | ZT12 | Present_Koike2012 | Absent_Koike2012 |
| ENSMUSG00000030555 | Ttc23 | 7 | 74792296 | 74871462 | + | 2.47E-05 | 0.0013711 | 2.39461929 | 1.7778248 | 3.7278566 | 3.45759879 | 3.7278566 | ZT12 | Present_Koike2012 | Present_Koike2012 |
| ENSMUSG00000030681 | Mvp | 7 | 134130374 | 134158135 | - | 2.50E-05 | 0.00138278 | 4.46392143 | 4.41128037 | 5.77834038 | 4.84848455 | 5.77834038 | ZT12 | Absent_Koike2012 | Absent_Koike2012 |
| ENSMUSG00000049044 | Rapgef4 | 2 | 71819272 | 72095531 | + | 2.51E-05 | 0.00138633 | 5.85640939 | 9.85834037 | 11.8813259 | 5.672809 | 11.8813259 | ZT12 | Present_Koike2012 | Present_Koike2012 |
| ENSMUSG00000038299 | Wdr36 | 18 | 32996879 | 33026074 | + | 2.61E-05 | 0.00142765 | 7.02466553 | 6.69175175 | 8.25447207 | 7.69660457 | 8.25447207 | ZT12 | Absent_Koike2012 | Present_Koike2012 |
| ENSMUSG00000024254 | Abcg8 | 17 | 85075642 | 85099673 | + | 2.62E-05 | 0.00142998 | 9.59557136 | 21.3492574 | 23.2678908 | 8.35790492 | 23.2678908 | ZT12 | Present_Koike2012 | Absent_Koike2012 |
| ENSMUSG00000028212 | Ccne2 | 4 | 11118498 | 11131926 | + | 2.71E-05 | 0.0014526 | 0.36464079 | 0.25596299 | 0.891258 | 0.55801125 | 0.891258 | ZT12 | Absent_Koike2012 | Absent_Koike2012 |
| ENSMUSG00000053931 | Cnn3 | 3 | 121129459 | 121161123 | + | 2.70E-05 | 0.0014526 | 37.4937163 | 38.20407 | 49.3494473 | 44.4774143 | 49.3494473 | ZT12 | Absent_Koike2012 | Absent_Koike2012 |
| ENSMUSG00000086265 | Marcksl1-ps4 | 13 | 4247981 | 4248271 | + | 2.77E-05 | 0.0014723 | 8.90537225 | 7.2445485 | 16.6501015 | 14.1122825 | 16.6501015 | ZT12 | Absent_Koike2012 | Absent_Koike2012 |
| ENSMUSG00000002733 | Plekha3 | 2 | 76513338 | 76534885 | + | 2.83E-05 | 0.00149573 | 4.81597525 | 4.97185716 | 6.26741827 | 5.24663252 | 6.26741827 | ZT12 | Absent_Koike2012 | Absent_Koike2012 |
| ENSMUSG00000007613 | Tgfbr1 | 4 | 47360094 | 47427803 | + | 2.88E-05 | 0.00150704 | 4.75905026 | 5.9366987 | 7.26461227 | 5.05211417 | 7.26461227 | ZT12 | Present_Koike2012 | Present_Koike2012 |
| ENSMUSG00000030967 | Zranb1 | 7 | 140122825 | 140178074 | + | 2.88E-05 | 0.00150704 | 4.69770712 | 6.05304518 | 6.63539099 | 4.4681189 | 6.63539099 | ZT12 | Present_Koike2012 | Absent_Koike2012 |
| ENSMUSG00000015942 | Gtf2ird2 | 5 | 134659889 | 134700225 | + | 2.92E-05 | 0.00151505 | 0.57665507 | 0.59907424 | 0.87129907 | 0.51239565 | 0.87129907 | ZT12 | Absent_Koike2012 | Absent_Koike2012 |
| ENSMUSG00000035372 | 1810055G02 | 19 | 3708333 | 3717881 | + | 3.00E-05 | 0.00154605 | 2.88356594 | 5.23830491 | 9.74123425 | 3.73008825 | 9.74123425 | ZT12 | Absent_Koike2012 | Present_Koike2012 |
| ENSMUSG00000021133 | 4933426M11 | 12 | 81891546 | 81981819 | + | 3.00E-05 | 0.00154768 | 4.84815232 | 5.89651572 | 7.19930276 | 5.01027474 | 7.19930276 | ZT12 | Absent_Koike2012 | Absent_Koike2012 |

| | | | | | | | | | | | | | | | |
|---|---|---|---|---|---|---|---|---|---|---|---|---|---|---|---|
| ENSMUSG00000039976 | Tbc1d16 | 11 | 119004359 | 119089813 | - | 3.05E-05 | 0.00156349 | 2.33009805 | 2.49283896 | 3.60366588 | 2.66040478 | 3.60366588 | ZT12 | Absent_Koike2012 | Absent_Koike2012 |
| ENSMUSG00000073700 | Klhl21 | 4 | 151382912 | 151391789 | + | 3.09E-05 | 0.00157374 | 4.69648575 | 6.41667625 | 6.5051985 | 3.6476105 | 6.5051985 | ZT12 | Absent_Koike2012 | Absent_Koike2012 |
| ENSMUSG00000027227 | Sord | 2 | 122060485 | 122091076 | + | 3.16E-05 | 0.00160506 | 404.0196 | 363.662331 | 528.984726 | 519.127251 | 528.984726 | ZT12 | Present_Koike2012 | Absent_Koike2012 |
| ENSMUSG00000048960 | Prex2 | 1 | 10983546 | 11293763 | + | 3.20E-05 | 0.00162212 | 1.59727125 | 1.666224 | 2.01462475 | 1.859721 | 2.01462475 | ZT12 | Present_Koike2012 | Absent_Koike2012 |
| ENSMUSG00000034867 | Ankrd27 | 7 | 36371266 | 36424256 | + | 3.27E-05 | 0.00164323 | 4.90513254 | 5.18906894 | 6.31835057 | 5.2750656 | 6.31835057 | ZT12 | Absent_Koike2012 | Absent_Koike2012 |
| ENSMUSG00000085214 | 0610005C13I | 7 | 52823165 | 52830697 | - | 3.37E-05 | 0.00168585 | 11.0463244 | 12.5394264 | 15.3412829 | 11.6925951 | 15.3412829 | ZT12 | Absent_Koike2012 | Absent_Koike2012 |
| ENSMUSG00000061313 | Ddhd2 | 8 | 26835796 | 26865068 | - | 3.51E-05 | 0.00173138 | 3.40220568 | 3.2208576 | 5.88358568 | 4.1315089 | 5.88358568 | ZT12 | Present_Koike2012 | Absent_Koike2012 |
| ENSMUSG00000020532 | Acaca | 11 | 83943174 | 84215153 | + | 3.50E-05 | 0.00173138 | 12.9534724 | 13.6299162 | 24.3615951 | 18.8321741 | 24.3615951 | ZT12 | Present_Koike2012 | Absent_Koike2012 |
| ENSMUSG00000024429 | Gnl1 | 17 | 36116796 | 36126407 | + | 3.50E-05 | 0.00173138 | 4.19197152 | 3.97190262 | 5.60519685 | 4.6247897 | 5.60519685 | ZT12 | Absent_Koike2012 | Absent_Koike2012 |
| ENSMUSG00000069565 | Dazap1 | 10 | 79724231 | 79751153 | + | 3.56E-05 | 0.00175151 | 5.11426502 | 6.1564481 | 6.48677915 | 5.40728057 | 6.48677915 | ZT12 | Absent_Koike2012 | Absent_Koike2012 |
| ENSMUSG00000025809 | Itgb1 | 8 | 131209554 | 131257100 | + | 3.61E-05 | 0.00177203 | 24.7129302 | 27.9674378 | 30.5908664 | 25.5561984 | 30.5908664 | ZT12 | Present_Koike2012 | Present_Koike2012 |
| ENSMUSG00000039703 | Nploc4 | 11 | 120241684 | 120299022 | - | 3.67E-05 | 0.00178344 | 13.2581296 | 13.5869839 | 15.8115078 | 12.7262159 | 15.8115078 | ZT12 | Present_Koike2012 | Absent_Koike2012 |
| ENSMUSG00000035171 | 1110059E24I | 19 | 21671803 | 21727281 | - | 3.67E-05 | 0.00178344 | 8.32898016 | 8.75477671 | 12.8024581 | 8.65411838 | 12.8024581 | ZT12 | Absent_Koike2012 | Present_Koike2012 |
| ENSMUSG00000039463 | Slc9a8 | 2 | 167247212 | 167302500 | + | 3.67E-05 | 0.00178344 | 2.02325207 | 2.38414383 | 2.5214329 | 1.83842279 | 2.5214329 | ZT12 | Absent_Koike2012 | Absent_Koike2012 |
| ENSMUSG00000020085 | Aifm2 | 10 | 61178011 | 61202008 | + | 3.72E-05 | 0.00179383 | 2.9678329 | 2.43968225 | 4.22217042 | 3.26318162 | 4.22217042 | ZT12 | Absent_Koike2012 | Absent_Koike2012 |
| ENSMUSG00000019944 | Rhobtb1 | 10 | 68614182 | 68754539 | + | 3.72E-05 | 0.00179383 | 7.23999069 | 4.61649766 | 8.53688697 | 7.56684633 | 8.53688697 | ZT12 | Absent_Koike2012 | Absent_Koike2012 |
| ENSMUSG00000070348 | Ccnd1 | 7 | 152115836 | 152125774 | - | 3.76E-05 | 0.00180602 | 6.05269243 | 9.17016486 | 15.0801201 | 5.10756358 | 15.0801201 | ZT12 | Present_Koike2012 | Absent_Koike2012 |
| ENSMUSG00000043439 | E130012A19I | 11 | 97488707 | 97491030 | - | 3.89E-05 | 0.00185522 | 0.45396175 | 0.4630145 | 1.0398065 | 0.902784 | 1.0398065 | ZT12 | Absent_Koike2012 | Absent_Koike2012 |
| ENSMUSG00000040584 | Abcb1a | 5 | 8660077 | 8748575 | + | 4.04E-05 | 0.00190165 | 0.41760785 | 0.50620951 | 1.29975293 | 0.65752832 | 1.29975293 | ZT12 | Present_Koike2012 | Absent_Koike2012 |
| ENSMUSG00000031701 | Dnaja2 | 8 | 88061539 | 88079170 | - | 4.18E-05 | 0.00194776 | 25.8493995 | 27.0145388 | 34.3057783 | 27.055438 | 34.3057783 | ZT12 | Present_Koike2012 | Present_Koike2012 |
| ENSMUSG00000026082 | Rev1 | 1 | 38109632 | 38186507 | - | 4.23E-05 | 0.0019537 | 1.98103725 | 2.175931 | 2.88823225 | 2.13313775 | 2.88823225 | ZT12 | Present_Koike2012 | Present_Koike2012 |
| ENSMUSG00000021583 | Erap1 | 13 | 74777320 | 74829983 | + | 4.34E-05 | 0.0019957 | 16.3766191 | 17.116295 | 19.7846143 | 18.0005263 | 19.7846143 | ZT12 | Absent_Koike2012 | Present_Koike2012 |
| ENSMUSG00000028953 | Abcf2 | 5 | 24071161 | 24083217 | - | 4.38E-05 | 0.00199897 | 12.4097665 | 10.2957731 | 12.6435877 | 12.2212135 | 12.6435877 | ZT12 | Absent_Koike2012 | Absent_Koike2012 |
| ENSMUSG00000033955 | Tnks1bp1 | 2 | 84888179 | 84913205 | + | 4.37E-05 | 0.00199897 | 4.49475051 | 4.59216699 | 6.48265762 | 6.19951862 | 6.48265762 | ZT12 | Absent_Koike2012 | Absent_Koike2012 |
| ENSMUSG00000005547 | Cyp2a5 | 7 | 27620324 | 27628567 | + | 4.48E-05 | 0.00202635 | 56.8873144 | 64.9639384 | 213.844402 | 182.418937 | 213.844402 | ZT12 | Present_Koike2012 | Present_Koike2012 |
| ENSMUSG00000036402 | Gng12 | 6 | 66846391 | 66971344 | + | 4.62E-05 | 0.00206788 | 6.48025697 | 7.55538578 | 7.94353718 | 6.726463 | 7.94353718 | ZT12 | Absent_Koike2012 | Absent_Koike2012 |
| ENSMUSG00000048445 | Ccdc57 | 11 | 120687843 | 120794186 | - | 4.62E-05 | 0.00206788 | 0.24623201 | 0.22950743 | 0.42439953 | 0.39892737 | 0.42439953 | ZT12 | Absent_Koike2012 | Absent_Koike2012 |
| ENSMUSG00000039509 | Nup133 | 8 | 126421023 | 126473165 | - | 4.66E-05 | 0.00207572 | 1.7410595 | 1.838156 | 1.94311425 | 1.43519375 | 1.94311425 | ZT12 | Absent_Koike2012 | Absent_Koike2012 |
| ENSMUSG00000043885 | Slc36a4 | 9 | 15514182 | 15546828 | + | 4.78E-05 | 0.00211747 | 1.7678839 | 2.16151028 | 2.37807124 | 1.73493146 | 2.37807124 | ZT12 | Absent_Koike2012 | Absent_Koike2012 |
| ENSMUSG00000086853 | Gm14720 | X | 7291459 | 7299226 | + | 4.83E-05 | 0.00212642 | 4.69355024 | 6.02606244 | 6.61833056 | 4.58782223 | 6.61833056 | ZT12 | Absent_Koike2012 | Absent_Koike2012 |
| ENSMUSG00000037119 | D15Ertd621e | 15 | 58247023 | 58289356 | + | 4.83E-05 | 0.00212642 | 14.5064403 | 16.5796862 | 16.6706523 | 13.564039 | 16.6706523 | ZT12 | Present_Koike2012 | Absent_Koike2012 |
| ENSMUSG00000026980 | Ly75 | 2 | 60130160 | 60221360 | - | 4.85E-05 | 0.00212642 | 0.59814944 | 0.7123856 | 0.98386448 | 0.72434362 | 0.98386448 | ZT12 | Absent_Koike2012 | Absent_Koike2012 |
| ENSMUSG00000024074 | Crim1 | 17 | 78599588 | 78775932 | + | 4.87E-05 | 0.00213129 | 2.661394 | 3.040776 | 3.907991 | 3.03414925 | 3.907991 | ZT12 | Absent_Koike2012 | Absent_Koike2012 |
| ENSMUSG00000089942 | Pira2 | 7 | 3788412 | 3796653 | - | 4.90E-05 | 0.00213726 | 0.53084776 | 0.53558162 | 0.71150057 | 0.68101646 | 0.71150057 | ZT12 | Absent_Koike2012 | Absent_Koike2012 |
| ENSMUSG00000021211 | Akr1c12 | 13 | 4267418 | 4278679 | - | 5.01E-05 | 0.0021599 | 44.1782295 | 45.508418 | 54.387577 | 51.426366 | 54.387577 | ZT12 | Present_Koike2012 | Present_Koike2012 |
| ENSMUSG00000021876 | Rnase4 | 14 | 51710752 | 51725826 | + | 4.98E-05 | 0.0021599 | 113.927109 | 198.841539 | 219.382222 | 141.379773 | 219.382222 | ZT12 | Present_Koike2012 | Present_Koike2012 |
| ENSMUSG00000026956 | Uap1l1 | 2 | 25215409 | 25221202 | - | 4.99E-05 | 0.0021599 | 0.6630133 | 0.79027394 | 0.92292022 | 0.63941363 | 0.92292022 | ZT12 | Absent_Koike2012 | Absent_Koike2012 |
| ENSMUSG00000036104 | Rab3gap1 | 1 | 129765350 | 129840445 | + | 5.16E-05 | 0.00221101 | 4.99370877 | 5.72716243 | 6.02586252 | 4.60892881 | 6.02586252 | ZT12 | Absent_Koike2012 | Absent_Koike2012 |
| ENSMUSG00000022477 | Aco2 | 15 | 81702739 | 81745563 | + | 5.26E-05 | 0.00224838 | 33.6483834 | 30.1737245 | 40.1127764 | 36.5622379 | 40.1127764 | ZT12 | Absent_Koike2012 | Present_Koike2012 |
| ENSMUSG00000058351 | 2010107H07 | 14 | 31902055 | 31942116 | - | 5.34E-05 | 0.00226684 | 3.92283052 | 4.91224538 | 6.65977471 | 5.38303812 | 6.65977471 | ZT12 | Absent_Koike2012 | Absent_Koike2012 |
| ENSMUSG00000059970 | Hspa2 | 12 | 77505163 | 77507921 | + | 5.39E-05 | 0.00227719 | 0.67034136 | 0.48372774 | 1.51821459 | 1.12169741 | 1.51821459 | ZT12 | Present_Koike2012 | Absent_Koike2012 |
| ENSMUSG00000044477 | Zfand3 | 17 | 30142032 | 30346964 | + | 5.41E-05 | 0.00227719 | 5.9968143 | 7.36826625 | 7.60271727 | 5.63323698 | 7.60271727 | ZT12 | Absent_Koike2012 | Absent_Koike2012 |
| ENSMUSG00000026475 | Rgs16 | 1 | 155587479 | 155592598 | + | 5.40E-05 | 0.00227719 | 0.1439955 | 3.47302575 | 8.1296875 | 0.269185 | 8.1296875 | ZT12 | Absent_Koike2012 | Absent_Koike2012 |
| ENSMUSG00000020412 | Ascc2 | 11 | 4537750 | 4585702 | + | 5.56E-05 | 0.00231861 | 3.43800447 | 3.38507369 | 3.81873887 | 3.54839121 | 3.81873887 | ZT12 | Absent_Koike2012 | Absent_Koike2012 |
| ENSMUSG00000025228 | Actr1a | 19 | 46451304 | 46470225 | - | 5.59E-05 | 0.00232321 | 16.7254683 | 14.984547 | 17.1039183 | 16.0977578 | 17.1039183 | ZT12 | Absent_Koike2012 | Absent_Koike2012 |
| ENSMUSG00000073792 | Alg6 | 4 | 99382355 | 99430151 | + | 5.63E-05 | 0.0023292 | 0.8062667 | 0.84404334 | 0.94224939 | 0.70065742 | 0.94224939 | ZT12 | Absent_Koike2012 | Absent_Koike2012 |
| ENSMUSG00000027679 | Dnajc19 | 3 | 33954942 | 33980243 | - | 5.78E-05 | 0.00236138 | 5.17505628 | 5.11932394 | 6.00641012 | 5.87870946 | 6.00641012 | ZT12 | Absent_Koike2012 | Absent_Koike2012 |
| ENSMUSG00000008090 | Fgfrl1 | 5 | 109123248 | 109135969 | + | 5.81E-05 | 0.00236811 | 4.42675805 | 6.20347334 | 8.12391466 | 4.50419195 | 8.12391466 | ZT12 | Absent_Koike2012 | Absent_Koike2012 |
| ENSMUSG00000021771 | Vdac2 | 14 | 22650491 | 22665101 | + | 5.89E-05 | 0.00239217 | 10.0094542 | 10.2775048 | 12.5587947 | 10.1310656 | 12.5587947 | ZT12 | Absent_Koike2012 | Absent_Koike2012 |
| ENSMUSG00000004567 | Mcoln1 | 8 | 3500457 | 3515232 | + | 6.08E-05 | 0.0024525 | 3.3324738 | 3.56220896 | 4.32572623 | 3.92512623 | 4.32572623 | ZT12 | Absent_Koike2012 | Absent_Koike2012 |
| ENSMUSG00000022389 | Tef | 15 | 81632851 | 81657291 | + | 6.13E-05 | 0.00246017 | 2.29668792 | 6.99430172 | 12.1030584 | 7.81730081 | 12.1030584 | ZT12 | Present_Koike2012 | Present_Koike2012 |
| ENSMUSG00000025278 | Flnb | 14 | 8650471 | 8784102 | + | 6.22E-05 | 0.00248712 | 12.6093007 | 10.6627355 | 16.0079637 | 13.3387098 | 16.0079637 | ZT12 | Present_Koike2012 | Present_Koike2012 |
| ENSMUSG00000025410 | Dctn2 | 10 | 126703442 | 126718862 | + | 6.40E-05 | 0.00252863 | 33.3748008 | 37.1663543 | 39.3688 | 30.1807333 | 39.3688 | ZT12 | Absent_Koike2012 | Absent_Koike2012 |
| ENSMUSG00000018362 | Kpna2 | 11 | 106849943 | 106860855 | - | 6.39E-05 | 0.00252863 | 13.2169378 | 12.1586853 | 15.1605518 | 11.6190252 | 15.1605518 | ZT12 | Absent_Koike2012 | Absent_Koike2012 |
| ENSMUSG00000030316 | 1500001M2( | 6 | 114954399 | 114987894 | - | 6.54E-05 | 0.00256769 | 0.27836698 | 0.31911385 | 0.35951126 | 0.24363844 | 0.35951126 | ZT12 | Absent_Koike2012 | Absent_Koike2012 |
| ENSMUSG00000023980 | Taf8 | 17 | 47624999 | 47639241 | - | 6.55E-05 | 0.00256769 | 1.60381961 | 1.99732832 | 2.02682524 | 1.63631449 | 2.02682524 | ZT12 | Absent_Koike2012 | Absent_Koike2012 |
| ENSMUSG00000027610 | Gss | 2 | 155388917 | 155418566 | - | 6.55E-05 | 0.00256769 | 4.8819924 | 6.31838934 | 7.33788943 | 5.0546044 | 7.33788943 | ZT12 | Absent_Koike2012 | Absent_Koike2012 |
| ENSMUSG00000032127 | Vps11 | 9 | 44156189 | 44169753 | - | 6.59E-05 | 0.00257837 | 6.63641725 | 7.02492975 | 7.41867725 | 5.87793675 | 7.41867725 | ZT12 | Absent_Koike2012 | Absent_Koike2012 |
| ENSMUSG00000017765 | Slc12a4 | 8 | 108467490 | 108489974 | - | 6.67E-05 | 0.002592 | 3.42996554 | 3.82103982 | 5.05148087 | 3.64807978 | 5.05148087 | ZT12 | Absent_Koike2012 | Absent_Koike2012 |
| ENSMUSG00000032114 | Slc37a4 | 9 | 44205182 | 44211046 | + | 6.67E-05 | 0.002592 | 28.6853827 | 23.4516967 | 47.9922645 | 40.439098 | 47.9922645 | ZT12 | Present_Koike2012 | Present_Koike2012 |
| ENSMUSG00000043648 | Pld6 | 11 | 59597395 | 59601147 | - | 6.81E-05 | 0.00262595 | 0.1473611 | 0.32084003 | 0.35015623 | 0.25095388 | 0.35015623 | ZT12 | Absent_Koike2012 | Absent_Koike2012 |
| ENSMUSG00000033379 | Atp6v0b | 4 | 117556931 | 117559938 | - | 6.86E-05 | 0.00263795 | 12.3560624 | 11.4515686 | 15.5470639 | 13.7454651 | 15.5470639 | ZT12 | Absent_Koike2012 | Absent_Koike2012 |
| ENSMUSG00000029221 | Slc30a9 | 5 | 67698194 | 67749804 | + | 6.95E-05 | 0.00266531 | 5.1018484 | 5.27319384 | 6.61047665 | 4.61640511 | 6.61047665 | ZT12 | Present_Koike2012 | Present_Koike2012 |
| ENSMUSG00000028164 | Manba | 3 | 135148575 | 135234368 | + | 6.98E-05 | 0.00266689 | 2.69091997 | 2.51312532 | 3.4213298 | 3.25302227 | 3.4213298 | ZT12 | Absent_Koike2012 | Absent_Koike2012 |
| ENSMUSG00000021687 | Scamp1 | 13 | 94970043 | 95055812 | - | 7.08E-05 | 0.0027028 | 7.5026396 | 9.4491761 | 11.590511 | 7.40368855 | 11.590511 | ZT12 | Present_Koike2012 | Absent_Koike2012 |
| ENSMUSG00000028655 | Mfsd2a | 4 | 122624093 | 122638431 | - | 7.20E-05 | 0.00272763 | 1.59982492 | 2.26383175 | 15.634638 | 3.06792644 | 15.634638 | ZT12 | Absent_Koike2012 | Present_Koike2012 |
| ENSMUSG00000035686 | Thrsp | 7 | 104561448 | 104566240 | - | 7.22E-05 | 0.00272763 | 142.94983 | 67.5751472 | 556.138255 | 487.878503 | 556.138255 | ZT12 | Present_Koike2012 | Present_Koike2012 |
| ENSMUSG00000037343 | Taf2 | 15 | 54846684 | 54903707 | - | 7.24E-05 | 0.00272789 | 3.37641568 | 3.47211249 | 4.57771037 | 3.20645548 | 4.57771037 | ZT12 | Present_Koike2012 | Present_Koike2012 |
| ENSMUSG00000027613 | Eif6 | 2 | 155645568 | 155652661 | - | 7.26E-05 | 0.00273256 | 9.22880085 | 8.80960675 | 11.0792526 | 10.2476393 | 11.0792526 | ZT12 | Absent_Koike2012 | Absent_Koike2012 |

| | | | | | | | | | | | | | | | |
|---|---|---|---|---|---|---|---|---|---|---|---|---|---|---|---|
| **ENSMUSG00000084129** | Hmgb1-ps1 | 11 | 40955265 | 40955898 | + | 7.40E-05 | 0.00278069 | 6.1106235 | 6.9148445 | 7.896454 | 6.71955175 | 7.896454 | ZT12 | Absent_Koike2012 | Absent_Koike2012 |
| **ENSMUSG00000029610** | Aimp2 | 5 | 144663573 | 144670716 | - | 7.55E-05 | 0.00281147 | 2.92980847 | 2.73736746 | 3.65845561 | 3.49481827 | 3.65845561 | ZT12 | Present_Koike2012 | Absent_Koike2012 |
| **ENSMUSG00000021072** | Tmx1 | 12 | 71554082 | 71569027 | + | 7.57E-05 | 0.00281476 | 10.1742193 | 10.2058965 | 12.1371982 | 10.7742403 | 12.1371982 | ZT12 | Absent_Koike2012 | Absent_Koike2012 |
| **ENSMUSG00000034795** | Ccdc122 | 14 | 77436579 | 77512064 | + | 7.62E-05 | 0.00282578 | 1.33781708 | 1.91220047 | 1.94489075 | 1.29808376 | 1.94489075 | ZT12 | Absent_Koike2012 | Absent_Koike2012 |
| **ENSMUSG00000021076** | Actr10 | 12 | 72038844 | 72065705 | + | 7.74E-05 | 0.0028464 | 6.34866125 | 6.78246075 | 6.85274375 | 5.9220235 | 6.85274375 | ZT12 | Absent_Koike2012 | Absent_Koike2012 |
| **ENSMUSG00000030556** | Lrrc28 | 7 | 74658297 | 74790122 | - | 7.85E-05 | 0.00287606 | 3.55741445 | 2.47520752 | 5.68844504 | 4.9018361 | 5.68844504 | ZT12 | Present_Koike2012 | Present_Koike2012 |
| **ENSMUSG00000032051** | Fdx1 | 9 | 51751412 | 51771661 | - | 7.94E-05 | 0.00290346 | 15.0653341 | 15.8132169 | 18.7550935 | 16.8392491 | 18.7550935 | ZT12 | Absent_Koike2012 | Present_Koike2012 |
| **ENSMUSG00000042699** | Dhx9 | 1 | 155302888 | 155334790 | - | 8.01E-05 | 0.00291727 | 5.5090415 | 6.52421425 | 8.261877 | 6.2779625 | 8.261877 | ZT12 | Present_Koike2012 | Absent_Koike2012 |
| **ENSMUSG00000043286** | Pnpla1 | 17 | 28995356 | 29027253 | + | 8.09E-05 | 0.00293109 | 0.11777896 | 0.15766766 | 0.17578758 | 0.10920978 | 0.17578758 | ZT12 | Absent_Koike2012 | Absent_Koike2012 |
| **ENSMUSG00000025487** | Psmd13 | 7 | 148067867 | 148084542 | + | 8.21E-05 | 0.00296885 | 8.23838952 | 7.60697189 | 9.07857399 | 8.3508463 | 9.07857399 | ZT12 | Absent_Koike2012 | Absent_Koike2012 |
| **ENSMUSG00000042834** | D0H4S114 | 18 | 33596673 | 33623683 | - | 8.49E-05 | 0.00303448 | 7.2639555 | 5.12519756 | 17.0167111 | 11.7805665 | 17.0167111 | ZT12 | Present_Koike2012 | Present_Koike2012 |
| **ENSMUSG00000029683** | Lmod2 | 6 | 24547771 | 24555414 | + | 8.76E-05 | 0.0031138 | 0.03143775 | 0.00611475 | 0.1489235 | 0.146658 | 0.1489235 | ZT12 | Absent_Koike2012 | Absent_Koike2012 |
| **ENSMUSG00000030660** | Pik3c2a | 7 | 123480790 | 123586972 | - | 8.78E-05 | 0.00311583 | 10.8589573 | 9.87908381 | 13.1598068 | 10.3263012 | 13.1598068 | ZT12 | Absent_Koike2012 | Present_Koike2012 |
| **ENSMUSG00000025485** | Ric8 | 7 | 148042856 | 148049629 | + | 8.83E-05 | 0.00312846 | 2.44672294 | 2.73266442 | 2.95003431 | 2.413882 | 2.95003431 | ZT12 | Absent_Koike2012 | Absent_Koike2012 |
| **ENSMUSG00000071637** | Cebpd | 16 | 15887379 | 15888559 | + | 8.93E-05 | 0.00315668 | 1.05294025 | 0.83375775 | 2.47945525 | 1.4953865 | 2.47945525 | ZT12 | Absent_Koike2012 | Absent_Koike2012 |
| **ENSMUSG00000027598** | Itch | 2 | 154959245 | 155052591 | + | 9.23E-05 | 0.00322582 | 6.11148199 | 6.98547179 | 8.30054125 | 6.68694653 | 8.30054125 | ZT12 | Present_Koike2012 | Absent_Koike2012 |
| **ENSMUSG00000038582** | Pptc7 | 5 | 122734374 | 122774290 | + | 9.38E-05 | 0.00327075 | 2.29769925 | 2.68227986 | 3.80752209 | 2.403215 | 3.80752209 | ZT12 | Present_Koike2012 | Absent_Koike2012 |
| **ENSMUSG00000026409** | Pfkfb2 | 1 | 132585620 | 132625830 | - | 9.54E-05 | 0.00330874 | 2.51277456 | 2.10332975 | 3.79885214 | 2.40529589 | 3.79885214 | ZT12 | Present_Koike2012 | Present_Koike2012 |
| **ENSMUSG00000024527** | Afg3l2 | 18 | 67564421 | 67608820 | - | 9.53E-05 | 0.00330874 | 13.6329899 | 15.9976284 | 18.0187643 | 14.4555253 | 18.0187643 | ZT12 | Absent_Koike2012 | Absent_Koike2012 |
| **ENSMUSG00000031431** | Tsc22d3 | X | 137074067 | 137135198 | - | 9.62E-05 | 0.00332302 | 11.6104667 | 5.75494023 | 18.8483352 | 15.2224049 | 18.8483352 | ZT12 | Present_Koike2012 | Present_Koike2012 |
| **ENSMUSG00000081534** | Slc48a1 | 15 | 97614786 | 97623123 | + | 9.62E-05 | 0.00332302 | 7.76659707 | 8.72681698 | 9.50229605 | 7.98594786 | 9.50229605 | ZT12 | Absent_Koike2012 | Absent_Koike2012 |
| **ENSMUSG00000010453** | 4632411B12I | 1 | 36392575 | 36426026 | - | 9.70E-05 | 0.00333663 | 5.75751135 | 6.96954617 | 7.28992532 | 5.40729898 | 7.28992532 | ZT12 | Present_Koike2012 | Absent_Koike2012 |
| **ENSMUSG00000086165** | Gm15690 | 5 | 120929851 | 120930912 | - | 9.70E-05 | 0.00333663 | 0.16205375 | 0.75553675 | 2.33038725 | 0.6111615 | 2.33038725 | ZT12 | Absent_Koike2012 | Absent_Koike2012 |
| **ENSMUSG00000005142** | Man2b1 | 8 | 87607168 | 87622638 | + | 9.89E-05 | 0.00337204 | 22.5224759 | 21.848573 | 24.4823827 | 24.4174757 | 24.4823827 | ZT12 | Absent_Koike2012 | Absent_Koike2012 |
| **ENSMUSG00000040253** | Gbp7 | 3 | 142193302 | 142213047 | + | 0.00010169 | 0.00345032 | 2.60753161 | 3.72948771 | 3.96691281 | 2.49892013 | 3.96691281 | ZT12 | Absent_Koike2012 | Absent_Koike2012 |
| **ENSMUSG00000038780** | Smurf1 | 5 | 145637364 | 145726716 | - | 0.00010211 | 0.00345989 | 1.86413749 | 2.30862282 | 2.64416471 | 2.14983161 | 2.64416471 | ZT12 | Absent_Koike2012 | Absent_Koike2012 |
| **ENSMUSG00000085001** | BB114351 | 2 | 72062745 | 72070359 | - | 0.00010578 | 0.00355752 | 0.7903735 | 1.60038375 | 1.61173 | 0.82630325 | 1.61173 | ZT12 | Absent_Koike2012 | Absent_Koike2012 |
| **ENSMUSG00000053769** | Lysmd1 | 3 | 94938010 | 94943440 | + | 0.0001063 | 0.00356866 | 3.030553 | 3.08688025 | 3.139448 | 2.66251575 | 3.139448 | ZT12 | Absent_Koike2012 | Absent_Koike2012 |
| **ENSMUSG00000049580** | Tsku | 7 | 105499178 | 105509838 | - | 0.00010701 | 0.00358304 | 6.61551609 | 2.3132843 | 18.392512 | 10.8708461 | 18.392512 | ZT12 | Present_Koike2012 | Present_Koike2012 |
| **ENSMUSG00000024352** | Spata24 | 18 | 35816344 | 35821840 | - | 0.00010754 | 0.00359589 | 0.83025966 | 0.8713815 | 1.05507376 | 0.62170763 | 1.05507376 | ZT12 | Absent_Koike2012 | Present_Koike2012 |
| **ENSMUSG00000073563** | Csnk1g3 | 18 | 54021767 | 54115338 | + | 0.00010998 | 0.00366792 | 4.75547635 | 4.63526632 | 6.53301261 | 5.12025727 | 6.53301261 | ZT12 | Present_Koike2012 | Present_Koike2012 |
| **ENSMUSG00000032803** | Cdv3 | 9 | 103255432 | 103268110 | - | 0.00011183 | 0.00369585 | 22.9891253 | 21.4571859 | 26.775492 | 23.1461178 | 26.775492 | ZT12 | Present_Koike2012 | Present_Koike2012 |
| **ENSMUSG00000024212** | Mllt1 | 17 | 57032035 | 57074836 | - | 0.00011205 | 0.00369816 | 3.096283 | 2.791271 | 3.53570925 | 3.08574475 | 3.53570925 | ZT12 | Absent_Koike2012 | Absent_Koike2012 |
| **ENSMUSG00000029635** | Cdk8 | 5 | 147042806 | 147114450 | + | 0.00011266 | 0.00371347 | 4.28225844 | 4.60342034 | 4.92460465 | 3.98776156 | 4.92460465 | ZT12 | Present_Koike2012 | Absent_Koike2012 |
| **ENSMUSG00000024425** | Ndfip1 | 18 | 38578629 | 38624053 | + | 0.0001128 | 0.00371347 | 90.1497193 | 101.042049 | 111.463659 | 103.18738 | 111.463659 | ZT12 | Absent_Koike2012 | Absent_Koike2012 |
| **ENSMUSG00000038280** | Ostm1 | 10 | 42303628 | 42422265 | + | 0.00011341 | 0.00372373 | 1.71974502 | 1.63784313 | 2.09651651 | 1.76247692 | 2.09651651 | ZT12 | Absent_Koike2012 | Absent_Koike2012 |
| **ENSMUSG00000044252** | Osbpl1a | 18 | 12913823 | 13100350 | - | 0.00011411 | 0.00373713 | 10.1097278 | 11.790865 | 12.3178892 | 9.85321563 | 12.3178892 | ZT12 | Absent_Koike2012 | Absent_Koike2012 |
| **ENSMUSG00000041220** | Elovl6 | 3 | 129235304 | 129341413 | + | 0.00011831 | 0.00383506 | 12.7414279 | 10.5130194 | 33.3221572 | 24.8612585 | 33.3221572 | ZT12 | Present_Koike2012 | Present_Koike2012 |
| **ENSMUSG00000004096** | Cwc15 | 9 | 14305061 | 14315021 | + | 0.00011801 | 0.00383506 | 26.2098365 | 26.3392226 | 27.6933876 | 24.4359651 | 27.6933876 | ZT12 | Absent_Koike2012 | Absent_Koike2012 |
| **ENSMUSG00000026028** | Trak2 | 1 | 58957293 | 59030274 | - | 0.00011824 | 0.00383506 | 3.2136346 | 3.29279941 | 4.0750259 | 2.98533663 | 4.0750259 | ZT12 | Present_Koike2012 | Absent_Koike2012 |
| **ENSMUSG00000039156** | Stim2 | 5 | 54389656 | 54512296 | + | 0.00012109 | 0.00390552 | 3.21270082 | 2.90514401 | 5.04170595 | 3.71549999 | 5.04170595 | ZT12 | Absent_Koike2012 | Present_Koike2012 |
| **ENSMUSG00000040430** | Pitpnc1 | 11 | 107069206 | 107332034 | - | 0.00012266 | 0.00394365 | 2.45385241 | 3.03355319 | 4.22817517 | 3.03172112 | 4.22817517 | ZT12 | Present_Koike2012 | Absent_Koike2012 |
| **ENSMUSG00000022177** | Haus4 | 14 | 55160622 | 55173198 | - | 0.00012251 | 0.00394365 | 4.04409425 | 4.67359625 | 5.4701875 | 4.00378 | 5.4701875 | ZT12 | Absent_Koike2012 | Absent_Koike2012 |
| **ENSMUSG00000000339** | Rtcd1 | 3 | 116191881 | 116211126 | - | 0.00012296 | 0.00394572 | 2.06940295 | 2.26080872 | 2.35297952 | 1.76708157 | 2.35297952 | ZT12 | Absent_Koike2012 | Present_Koike2012 |
| **ENSMUSG00000038172** | Ttc39b | 4 | 82866204 | 82970159 | - | 0.00012658 | 0.00402483 | 2.80750988 | 4.17717828 | 4.36727071 | 2.6634203 | 4.36727071 | ZT12 | Present_Koike2012 | Absent_Koike2012 |
| **ENSMUSG00000020044** | Timp3 | 10 | 85763117 | 85812251 | + | 0.00013171 | 0.00413257 | 2.49488733 | 3.19848004 | 4.68639896 | 3.79142014 | 4.68639896 | ZT12 | Present_Koike2012 | Present_Koike2012 |
| **ENSMUSG00000060121** | Sip1 | 12 | 60114380 | 60129455 | + | 0.00013403 | 0.00417457 | 3.50781161 | 3.53298561 | 4.64905687 | 3.87247959 | 4.64905687 | ZT12 | Absent_Koike2012 | Absent_Koike2012 |
| **ENSMUSG00000019877** | Serinc1 | 10 | 57235580 | 57252336 | - | 0.00013707 | 0.00423156 | 28.9589304 | 32.5108756 | 34.4034618 | 29.966652 | 34.4034618 | ZT12 | Absent_Koike2012 | Absent_Koike2012 |
| **ENSMUSG00000039512** | Uhrf1bp1 | 17 | 27993435 | 28036985 | + | 0.00013719 | 0.00423156 | 2.68311292 | 2.43885696 | 3.23337836 | 2.80765272 | 3.23337836 | ZT12 | Absent_Koike2012 | Present_Koike2012 |
| **ENSMUSG00000036825** | Ssx2ip | 3 | 146067606 | 146103103 | + | 0.00014011 | 0.00429533 | 2.16923036 | 1.68274005 | 2.45405435 | 2.3734422 | 2.45405435 | ZT12 | Present_Koike2012 | Absent_Koike2012 |
| **ENSMUSG00000038034** | Igsf8 | 1 | 174191772 | 174249972 | + | 0.00014294 | 0.00433658 | 0.95460759 | 1.1124747 | 1.66376459 | 0.95964437 | 1.66376459 | ZT12 | Absent_Koike2012 | Absent_Koike2012 |
| **ENSMUSG00000058542** | Gm15590 | 17 | 6131695 | 6132613 | - | 0.00014408 | 0.00435969 | 167.197553 | 147.63071 | 213.597424 | 191.538734 | 213.597424 | ZT12 | Absent_Koike2012 | Absent_Koike2012 |
| **ENSMUSG00000026781** | Acbd5 | 2 | 22923688 | 22970030 | + | 0.00014534 | 0.00438723 | 46.1921665 | 57.0135861 | 57.6353613 | 40.7052129 | 57.6353613 | ZT12 | Present_Koike2012 | Absent_Koike2012 |
| **ENSMUSG00000044768** | D1Ertd622e | 1 | 99540480 | 99558651 | - | 0.00014561 | 0.00438917 | 1.93968426 | 2.18526496 | 3.52990825 | 2.01730724 | 3.52990825 | ZT12 | Present_Koike2012 | Present_Koike2012 |
| **ENSMUSG00000019194** | Scn1b | 7 | 31901544 | 31911964 | - | 0.00014708 | 0.00441843 | 0.980583 | 0.99683025 | 1.284121 | 1.05741925 | 1.284121 | ZT12 | Absent_Koike2012 | Absent_Koike2012 |
| **ENSMUSG00000046230** | Vps13a | 19 | 16691253 | 16855417 | - | 0.00014759 | 0.0044235 | 3.44480942 | 2.87093455 | 5.03742891 | 4.9992544 | 5.03742891 | ZT12 | Present_Koike2012 | Present_Koike2012 |
| **ENSMUSG00000010048** | Ifrd2 | 9 | 107490049 | 107495369 | + | 0.00014891 | 0.0044371 | 10.0390518 | 9.91741625 | 12.9744508 | 11.8250245 | 12.9744508 | ZT12 | Absent_Koike2012 | Absent_Koike2012 |
| **ENSMUSG00000046096** | BC030336 | 7 | 127821134 | 127878368 | + | 0.00015176 | 0.00449242 | 2.93704325 | 3.034935 | 3.924984 | 2.892834 | 3.924984 | ZT12 | Present_Koike2012 | Absent_Koike2012 |
| **ENSMUSG00000028149** | Rap1gds1 | 3 | 138588870 | 138738163 | - | 0.00015231 | 0.00449565 | 5.97346591 | 5.50661993 | 6.7355586 | 5.97900467 | 6.7355586 | ZT12 | Absent_Koike2012 | Present_Koike2012 |
| **ENSMUSG00000019849** | Prep | 10 | 44787012 | 44878803 | + | 0.0001532 | 0.00451199 | 8.77399665 | 8.89800403 | 11.4734071 | 9.96639124 | 11.4734071 | ZT12 | Absent_Koike2012 | Absent_Koike2012 |
| **ENSMUSG00000032536** | Trak1 | 9 | 121206620 | 121384034 | + | 0.00015563 | 0.00456773 | 2.77697145 | 3.10875427 | 4.03144974 | 3.17359459 | 4.03144974 | ZT12 | Present_Koike2012 | Absent_Koike2012 |
| **ENSMUSG00000025509** | Pnpla2 | 7 | 148641097 | 148646642 | + | 0.0001603 | 0.00468256 | 4.4612968 | 6.73037102 | 6.96070294 | 4.49537795 | 6.96070294 | ZT12 | Present_Koike2012 | Absent_Koike2012 |
| **ENSMUSG00000019715** | Gle1 | 2 | 29790946 | 29815891 | + | 0.00016253 | 0.00472659 | 2.99699483 | 3.16980637 | 3.56169962 | 2.74524224 | 3.56169962 | ZT12 | Absent_Koike2012 | Absent_Koike2012 |
| **ENSMUSG00000078521** | 2610002D18 | 4 | 134066914 | 134079842 | + | 0.0001635 | 0.00473866 | 0 | 0.024627 | 0.13509025 | 0.088999 | 0.13509025 | ZT12 | Absent_Koike2012 | Absent_Koike2012 |
| **ENSMUSG00000062382** | Gm10116 | 13 | 74543448 | 74544765 | + | 0.00016429 | 0.00474776 | 768.061132 | 676.525901 | 933.214798 | 863.388165 | 933.214798 | ZT12 | Absent_Koike2012 | Absent_Koike2012 |
| **ENSMUSG00000032180** | Tmed1 | 9 | 21311824 | 21314631 | - | 0.00016518 | 0.0047548 | 5.65274091 | 4.90746559 | 7.0607419 | 5.43264686 | 7.0607419 | ZT12 | Absent_Koike2012 | Absent_Koike2012 |
| **ENSMUSG00000020988** | L2hgdh | 12 | 70791423 | 70825861 | - | 0.00016501 | 0.0047548 | 12.8948868 | 13.319188 | 16.8029898 | 12.8193175 | 16.8029898 | ZT12 | Present_Koike2012 | Absent_Koike2012 |
| **ENSMUSG00000022425** | Enpp2 | 15 | 54670453 | 54751701 | - | 0.00016512 | 0.0047548 | 12.4448225 | 19.029873 | 23.5925163 | 18.1379549 | 23.5925163 | ZT12 | Present_Koike2012 | Absent_Koike2012 |
| **ENSMUSG00000029260** | Ugt2b34 | 5 | 87318792 | 87335962 | - | 0.00016855 | 0.00483007 | 112.556157 | 98.7140881 | 131.991183 | 128.741317 | 131.991183 | ZT12 | Absent_Koike2012 | Absent_Koike2012 |

| | | | | | | | | | | | | | | | |
|---|---|---|---|---|---|---|---|---|---|---|---|---|---|---|---|
| ENSMUSG00000027387 | Zc3h8 | 2 | 128752004 | 128769813 | - | 0.00016895 | 0.00483065 | 1.00211407 | 0.66124095 | 1.44133444 | 1.10049918 | 1.44133444 | ZT12 | Present_Koike2012 | Present_Koike2012 |
| ENSMUSG00000024270 | Slc39a6 | 18 | 24738382 | 24762318 | - | 0.00016978 | 0.00484338 | 0.57060889 | 0.44077044 | 0.68352791 | 0.66578387 | 0.68352791 | ZT12 | Absent_Koike2012 | Absent_Koike2012 |
| ENSMUSG00000042572 | Ube2q1 | 3 | 89577531 | 89587919 | + | 0.00017024 | 0.00485115 | 7.1008425 | 7.28472375 | 7.719632 | 6.456146 | 7.719632 | ZT12 | Absent_Koike2012 | Absent_Koike2012 |
| ENSMUSG00000040675 | Mthfd1l | 10 | 6179460 | 6373423 | - | 0.0001721 | 0.00488232 | 0.28112133 | 0.29673405 | 1.86510594 | 0.61825678 | 1.86510594 | ZT12 | Absent_Koike2012 | Absent_Koike2012 |
| ENSMUSG00000022856 | Tmem41a | 16 | 21934400 | 21947617 | - | 0.00017276 | 0.00489549 | 2.77052774 | 2.42441508 | 3.81840389 | 3.44919106 | 3.81840389 | ZT12 | Absent_Koike2012 | Absent_Koike2012 |
| ENSMUSG00000042148 | Cox10 | 11 | 63776129 | 63892970 | - | 0.00017307 | 0.00489897 | 2.74093961 | 2.6299663 | 2.97329279 | 2.24223301 | 2.97329279 | ZT12 | Absent_Koike2012 | Absent_Koike2012 |
| ENSMUSG00000020810 | Cygb | 11 | 116506909 | 116515627 | - | 0.00017406 | 0.00491599 | 1.25762475 | 1.689139 | 2.24970625 | 1.457909 | 2.24970625 | ZT12 | Absent_Koike2012 | Absent_Koike2012 |
| ENSMUSG00000069763 | Tmem100 | 11 | 89891662 | 89897822 | + | 0.00017749 | 0.00499078 | 0.02327725 | 0.06565175 | 0.18073075 | 0.029842 | 0.18073075 | ZT12 | Absent_Koike2012 | Absent_Koike2012 |
| ENSMUSG00000068742 | Cry2 | 2 | 92243803 | 92274200 | - | 0.00017782 | 0.00499452 | 1.65727881 | 2.80011094 | 3.33569921 | 3.04877211 | 3.33569921 | ZT12 | Present_Koike2012 | Absent_Koike2012 |
| ENSMUSG00000024360 | Etf1 | 18 | 35062446 | 35091657 | - | 0.0001788 | 0.00501098 | 19.8555945 | 21.9364328 | 23.8052045 | 18.7657968 | 23.8052045 | ZT12 | Absent_Koike2012 | Absent_Koike2012 |
| ENSMUSG00000003849 | Nqo1 | 8 | 109912125 | 109927106 | - | 0.0001826 | 0.00506768 | 2.12477425 | 3.000792 | 3.95118275 | 2.513006 | 3.95118275 | ZT12 | Absent_Koike2012 | Absent_Koike2012 |
| ENSMUSG00000033715 | Akr1c14 | 13 | 4048257 | 4089668 | + | 0.00018325 | 0.00506768 | 112.393387 | 108.283937 | 179.669631 | 178.171892 | 179.669631 | ZT12 | Present_Koike2012 | Present_Koike2012 |
| ENSMUSG00000024740 | Ddb1 | 19 | 10680058 | 10704306 | + | 0.00018247 | 0.00506768 | 44.1449955 | 45.7767501 | 53.6768134 | 48.6097823 | 53.6768134 | ZT12 | Present_Koike2012 | Present_Koike2012 |
| ENSMUSG00000039157 | Fam102a | 2 | 32390852 | 32425276 | + | 0.00018201 | 0.00506768 | 1.87344543 | 1.36866831 | 2.77613067 | 2.70930476 | 2.77613067 | ZT12 | Present_Koike2012 | Absent_Koike2012 |
| ENSMUSG00000024767 | Otub1 | 19 | 7272692 | 7280806 | - | 0.00018875 | 0.00518695 | 9.06577492 | 9.61596651 | 11.2014635 | 9.33347871 | 11.2014635 | ZT12 | Absent_Koike2012 | Present_Koike2012 |
| ENSMUSG00000037536 | Fbxo34 | 14 | 48092236 | 48151637 | + | 0.00019145 | 0.00525545 | 3.32151902 | 4.40665918 | 4.80261119 | 3.63524325 | 4.80261119 | ZT12 | Present_Koike2012 | Absent_Koike2012 |
| ENSMUSG00000039456 | Morc3 | 16 | 93832366 | 93876317 | + | 0.00019906 | 0.0054293 | 2.87086925 | 2.80061975 | 4.77435175 | 3.6256935 | 4.77435175 | ZT12 | Present_Koike2012 | Present_Koike2012 |
| ENSMUSG00000018707 | Dync1h1 | 12 | 111839662 | 111905155 | + | 0.00020074 | 0.00545743 | 11.8372733 | 12.0340368 | 13.1124292 | 10.4909368 | 13.1124292 | ZT12 | Absent_Koike2012 | Absent_Koike2012 |
| ENSMUSG00000041040 | Fam117b | 1 | 59969850 | 60042190 | + | 0.00020492 | 0.00554748 | 1.53767144 | 1.63643461 | 1.94196007 | 1.57658132 | 1.94196007 | ZT12 | Absent_Koike2012 | Absent_Koike2012 |
| ENSMUSG00000061132 | Blnk | 19 | 41003417 | 41069025 | - | 0.00020556 | 0.00555306 | 0.1833506 | 0.28242573 | 0.50788967 | 0.25100439 | 0.50788967 | ZT12 | Absent_Koike2012 | Absent_Koike2012 |
| ENSMUSG00000025089 | Gfra1 | 19 | 58310094 | 58530399 | - | 0.00020604 | 0.00555989 | 20.8475113 | 21.146203 | 27.9656556 | 27.6514973 | 27.9656556 | ZT12 | Absent_Koike2012 | Absent_Koike2012 |
| ENSMUSG00000029201 | Ugdh | 5 | 65804441 | 65827185 | - | 0.00020641 | 0.00556405 | 10.0019184 | 10.6451427 | 15.46985 | 11.6632966 | 15.46985 | ZT12 | Present_Koike2012 | Absent_Koike2012 |
| ENSMUSG00000015214 | Mtmr1 | X | 68617935 | 68672371 | + | 0.00020794 | 0.00558593 | 5.6120031 | 6.26958444 | 6.67126796 | 5.40253458 | 6.67126796 | ZT12 | Absent_Koike2012 | Absent_Koike2012 |
| ENSMUSG00000039770 | Ypel5 | 17 | 73186044 | 73200535 | + | 0.00020832 | 0.00558593 | 7.22575434 | 9.63615997 | 11.2799183 | 8.97571277 | 11.2799183 | ZT12 | Present_Koike2012 | Absent_Koike2012 |
| ENSMUSG00000030447 | Cyfip1 | 7 | 63097115 | 63186070 | + | 0.00021049 | 0.00562045 | 4.89327241 | 5.28998736 | 5.45389125 | 4.95817548 | 5.45389125 | ZT12 | Absent_Koike2012 | Absent_Koike2012 |
| ENSMUSG00000044548 | Dact1 | 12 | 72410871 | 72421094 | + | 0.00021046 | 0.00562045 | 0.56244626 | 0.66796704 | 1.07697684 | 0.81270989 | 1.07697684 | ZT12 | Present_Koike2012 | Absent_Koike2012 |
| ENSMUSG00000039842 | Mcph1 | 8 | 18595131 | 18803189 | + | 0.00021529 | 0.00570295 | 0.43020701 | 0.47642004 | 0.58399812 | 0.36666668 | 0.58399812 | ZT12 | Absent_Koike2012 | Absent_Koike2012 |
| ENSMUSG00000042228 | Lyn | 4 | 3605262 | 3740269 | + | 0.00021781 | 0.00573766 | 6.26440253 | 6.61542591 | 7.51197792 | 6.81111102 | 7.51197792 | ZT12 | Absent_Koike2012 | Absent_Koike2012 |
| ENSMUSG00000020190 | Mknk2 | 10 | 80128077 | 80139038 | - | 0.00021759 | 0.00573766 | 7.08516495 | 9.38253847 | 18.3483189 | 15.561606 | 18.3483189 | ZT12 | Present_Koike2012 | Absent_Koike2012 |
| ENSMUSG00000057649 | Brd9 | 13 | 74075286 | 74098342 | + | 0.00021872 | 0.00575574 | 1.88327175 | 2.35428925 | 2.35455825 | 1.622531 | 2.35455825 | ZT12 | Absent_Koike2012 | Absent_Koike2012 |
| ENSMUSG00000015112 | Slc25a13 | 6 | 5991219 | 6167173 | - | 0.00021895 | 0.00575586 | 59.9447618 | 57.5725797 | 72.9000543 | 64.1718831 | 72.9000543 | ZT12 | Absent_Koike2012 | Absent_Koike2012 |
| ENSMUSG00000025199 | Chuk | 19 | 44147824 | 44181970 | - | 0.00022096 | 0.00579834 | 20.0047734 | 25.9095224 | 27.0751662 | 20.0714257 | 27.0751662 | ZT12 | Present_Koike2012 | Absent_Koike2012 |
| ENSMUSG00000035469 | Rcbtb1 | 14 | 59820046 | 59856102 | + | 0.0002228 | 0.00583292 | 5.09271724 | 5.29827722 | 6.04591811 | 4.93426005 | 6.04591811 | ZT12 | Absent_Koike2012 | Absent_Koike2012 |
| ENSMUSG00000045455 | Gm9797 | 10 | 11329021 | 11329633 | + | 0.00022499 | 0.00587213 | 6.84056625 | 6.590309 | 9.68151575 | 7.90675125 | 9.68151575 | ZT12 | Absent_Koike2012 | Absent_Koike2012 |
| ENSMUSG00000006715 | Gmnn | 13 | 24843714 | 24853792 | - | 0.0002271 | 0.00591169 | 1.83461983 | 2.33137902 | 3.36626843 | 2.05680338 | 3.36626843 | ZT12 | Absent_Koike2012 | Absent_Koike2012 |
| ENSMUSG00000004897 | Hdgf | 3 | 87710243 | 87720054 | + | 0.00022945 | 0.0059579 | 28.5913841 | 27.5233606 | 34.0420506 | 32.7832755 | 34.0420506 | ZT12 | Absent_Koike2012 | Present_Koike2012 |
| ENSMUSG00000027536 | Chmp4c | 3 | 10366907 | 10391005 | + | 0.0002302 | 0.00597125 | 0.380079 | 0.381715 | 0.587764 | 0.4011665 | 0.587764 | ZT12 | Absent_Koike2012 | Absent_Koike2012 |
| ENSMUSG00000030754 | Copb1 | 7 | 121359073 | 121398225 | - | 0.00023081 | 0.00597563 | 25.8178245 | 28.0480687 | 28.725905 | 24.433389 | 28.725905 | ZT12 | Absent_Koike2012 | Absent_Koike2012 |
| ENSMUSG00000084364 | Gm15801 | 19 | 40635027 | 40635548 | - | 0.00023083 | 0.00597563 | 0.48454275 | 0.400834 | 0.91145475 | 0.64942875 | 0.91145475 | ZT12 | Absent_Koike2012 | Absent_Koike2012 |
| ENSMUSG00000028015 | Ctso | 3 | 81736523 | 81760647 | + | 0.00023272 | 0.00601835 | 8.37659209 | 9.0822918 | 9.40015714 | 8.20417678 | 9.40015714 | ZT12 | Absent_Koike2012 | Absent_Koike2012 |
| ENSMUSG00000053664 | Uck2 | 1 | 169153958 | 169154050 | - | 0.00023596 | 0.00604067 | 1.9965575 | 1.27949525 | 2.608322 | 2.3453025 | 2.608322 | ZT12 | Absent_Koike2012 | Absent_Koike2012 |
| ENSMUSG00000054676 | 1600014C10I | 7 | 38968236 | 38982582 | + | 0.00023439 | 0.00604067 | 22.6965091 | 27.1548884 | 28.3719316 | 22.9422736 | 28.3719316 | ZT12 | Absent_Koike2012 | Absent_Koike2012 |
| ENSMUSG00000030972 | Acsm5 | 7 | 126669818 | 126687307 | + | 0.00023412 | 0.00604067 | 22.5020472 | 27.8249163 | 30.911502 | 21.6727964 | 30.911502 | ZT12 | Present_Koike2012 | Absent_Koike2012 |
| ENSMUSG00000021458 | 2010111I01R | 13 | 63066253 | 63427404 | + | 0.00023405 | 0.00604067 | 1.20192449 | 1.27161491 | 1.85334692 | 1.43845807 | 1.85334692 | ZT12 | Absent_Koike2012 | Present_Koike2012 |
| ENSMUSG00000024878 | Cbwd1 | 19 | 24994917 | 25036048 | - | 0.0002359 | 0.00604067 | 3.1368715 | 3.013322 | 3.52464125 | 2.47859275 | 3.52464125 | ZT12 | Absent_Koike2012 | Absent_Koike2012 |
| ENSMUSG00000020847 | Rph3al | 11 | 75644501 | 75751931 | - | 0.00023705 | 0.0060564 | 0.16788544 | 0.17466747 | 0.26594023 | 0.18036156 | 0.26594023 | ZT12 | Absent_Koike2012 | Absent_Koike2012 |
| ENSMUSG00000050936 | Hist2h2bb | 3 | 96073644 | 96082924 | + | 0.0002394 | 0.00608595 | 2.40639897 | 1.49233888 | 2.95542884 | 2.66837801 | 2.95542884 | ZT12 | Absent_Koike2012 | Absent_Koike2012 |
| ENSMUSG00000058761 | Rnf169 | 7 | 107068764 | 107128968 | - | 0.00023996 | 0.00609406 | 4.4418963 | 5.94927756 | 6.94277896 | 5.41884892 | 6.94277896 | ZT12 | Absent_Koike2012 | Absent_Koike2012 |
| ENSMUSG00000055538 | Zcchc24 | 14 | 26531126 | 26588342 | - | 0.0002422 | 0.00613248 | 11.3521202 | 13.5825893 | 14.666478 | 13.154626 | 14.666478 | ZT12 | Absent_Koike2012 | Absent_Koike2012 |
| ENSMUSG00000040158 | Tax1bp3 | 11 | 72990585 | 72996664 | + | 0.00024626 | 0.00620437 | 2.27100004 | 1.98081834 | 2.88834853 | 2.39097444 | 2.88834853 | ZT12 | Absent_Koike2012 | Absent_Koike2012 |
| ENSMUSG00000026634 | Angel2 | 1 | 192748991 | 192770841 | + | 0.00024723 | 0.00620437 | 2.18252925 | 2.5737745 | 3.57709765 | 3.04613985 | 3.57709765 | ZT12 | Absent_Koike2012 | Absent_Koike2012 |
| ENSMUSG00000027775 | Mfsd1 | 3 | 67386690 | 67408153 | + | 0.00024704 | 0.00620437 | 24.4650685 | 25.7106555 | 26.497504 | 21.9566375 | 26.497504 | ZT12 | Present_Koike2012 | Absent_Koike2012 |
| ENSMUSG00000020794 | Ube2g1 | 11 | 72420785 | 72499983 | + | 0.00024784 | 0.00621353 | 7.77530096 | 8.75435515 | 9.17272154 | 7.96457936 | 9.17272154 | ZT12 | Absent_Koike2012 | Present_Koike2012 |
| ENSMUSG00000034574 | Daam1 | 12 | 72932065 | 73093354 | + | 0.00025156 | 0.00627133 | 6.49998834 | 7.30420674 | 9.51169579 | 8.42965715 | 9.51169579 | ZT12 | Absent_Koike2012 | Absent_Koike2012 |
| ENSMUSG00000028793 | Rnf19b | 4 | 128735515 | 128763130 | + | 0.00025411 | 0.00631466 | 3.98491652 | 5.72606045 | 6.44112896 | 4.82219698 | 6.44112896 | ZT12 | Present_Koike2012 | Absent_Koike2012 |
| ENSMUSG00000022948 | Setd4 | 16 | 93583702 | 93604308 | + | 0.00025494 | 0.00631908 | 24.2624573 | 20.6926209 | 30.9916968 | 30.0667372 | 30.9916968 | ZT12 | Absent_Koike2012 | Absent_Koike2012 |
| ENSMUSG00000049672 | Zfp161 | 17 | 69732390 | 69740090 | + | 0.00025541 | 0.00631908 | 2.05083078 | 1.8932866 | 2.74823516 | 2.48372484 | 2.74823516 | ZT12 | Absent_Koike2012 | Absent_Koike2012 |
| ENSMUSG00000024622 | Hmgxb3 | 18 | 61290933 | 61336704 | - | 0.00025545 | 0.00631908 | 3.13843315 | 2.85133925 | 3.31162949 | 2.50994061 | 3.31162949 | ZT12 | Absent_Koike2012 | Absent_Koike2012 |
| ENSMUSG00000026839 | Upp2 | 2 | 58419709 | 58645382 | + | 0.00025649 | 0.00632457 | 14.3968498 | 30.3031019 | 55.9274199 | 31.3298588 | 55.9274199 | ZT12 | Present_Koike2012 | Absent_Koike2012 |
| ENSMUSG00000001891 | Ugp2 | 11 | 21221141 | 21271204 | - | 0.00025901 | 0.00634021 | 63.9887133 | 78.9435436 | 98.3679262 | 75.4905467 | 98.3679262 | ZT12 | Present_Koike2012 | Present_Koike2012 |
| ENSMUSG00000073725 | Lmbrd1 | 1 | 24685383 | 24823144 | + | 0.00026011 | 0.00635832 | 10.1622379 | 11.4118132 | 11.7569086 | 9.8048244 | 11.7569086 | ZT12 | Absent_Koike2012 | Absent_Koike2012 |
| ENSMUSG00000059937 | Gdap10 | 12 | 33506737 | 33511769 | + | 0.00026351 | 0.00640343 | 0.44937275 | 0.746077 | 1.7395735 | 0.80153325 | 1.7395735 | ZT12 | Absent_Koike2012 | Absent_Koike2012 |
| ENSMUSG00000000420 | Galnt1 | 18 | 24363845 | 24445319 | + | 0.00026535 | 0.00643078 | 7.80873188 | 5.1360706 | 8.29264408 | 8.00308484 | 8.29264408 | ZT12 | Absent_Koike2012 | Absent_Koike2012 |
| ENSMUSG00000071252 | 2210408I21R | 13 | 77274797 | 77752940 | + | 0.00026671 | 0.00645159 | 0.47742788 | 0.60441153 | 0.67742891 | 0.42395167 | 0.67742891 | ZT12 | Absent_Koike2012 | Present_Koike2012 |
| ENSMUSG00000056091 | St3gal5 | 6 | 72047607 | 72104564 | + | 0.00026732 | 0.00646012 | 7.04071273 | 8.42254187 | 17.3695375 | 12.5474627 | 17.3695375 | ZT12 | Present_Koike2012 | Present_Koike2012 |
| ENSMUSG00000082062 | Ftl2 | 4 | 125181532 | 125182083 | + | 0.00026869 | 0.00648444 | 171.800214 | 155.684157 | 218.654351 | 193.727668 | 218.654351 | ZT12 | Absent_Koike2012 | Absent_Koike2012 |
| ENSMUSG00000054434 | Tmem120b | 5 | 123518424 | 123567758 | + | 0.00027148 | 0.00653235 | 1.02859633 | 0.76896098 | 1.42899115 | 0.88586302 | 1.42899115 | ZT12 | Absent_Koike2012 | Absent_Koike2012 |
| ENSMUSG00000034557 | Zfyve9 | 4 | 108310071 | 108453403 | - | 0.00027364 | 0.00655679 | 4.15895769 | 4.28158902 | 5.16973999 | 4.99436151 | 5.16973999 | ZT12 | Present_Koike2012 | Present_Koike2012 |
| ENSMUSG00000029669 | Tspan12 | 6 | 21721395 | 21802515 | - | 0.00027425 | 0.00656523 | 44.6590259 | 52.9958677 | 58.5239744 | 43.1738151 | 58.5239744 | ZT12 | Present_Koike2012 | Absent_Koike2012 |

| | | | | | | | | | | | | | | | |
|---|---|---|---|---|---|---|---|---|---|---|---|---|---|---|---|
| ENSMUSG00000033557 | Fam20b | 1 | 158608676 | 158649132 | - | 0.00027577 | 0.00659134 | 2.47272956 | 2.31877031 | 2.94528159 | 2.62383684 | 2.94528159 | ZT12 | Absent_Koike2012 | Absent_Koike2012 |
| ENSMUSG00000067813 | Xkr9 | 1 | 13658852 | 13691804 | + | 0.00028234 | 0.00670837 | 6.78828827 | 6.91627903 | 9.44910391 | 7.30958918 | 9.44910391 | ZT12 | Absent_Koike2012 | Absent_Koike2012 |
| ENSMUSG00000028261 | Ndufaf4 | 4 | 24825230 | 24832148 | + | 0.00028408 | 0.00673308 | 1.8376293 | 1.24111893 | 1.8887354 | 1.79366811 | 1.8887354 | ZT12 | Absent_Koike2012 | Present_Koike2012 |
| ENSMUSG00000027599 | Armc1 | 3 | 19032146 | 19063065 | - | 0.00028631 | 0.00677239 | 12.068645 | 11.254328 | 13.2495065 | 12.2224555 | 13.2495065 | ZT12 | Present_Koike2012 | Absent_Koike2012 |
| ENSMUSG00000021576 | Pdcd6 | 13 | 74440569 | 74454774 | - | 0.00029113 | 0.0068596 | 24.384413 | 27.0744914 | 29.3842179 | 26.8444046 | 29.3842179 | ZT12 | Absent_Koike2012 | Absent_Koike2012 |
| ENSMUSG00000031696 | Vps35 | 8 | 87784296 | 87823412 | - | 0.00030024 | 0.00699001 | 24.3643008 | 25.35185 | 27.2602443 | 22.0028638 | 27.2602443 | ZT12 | Absent_Koike2012 | Absent_Koike2012 |
| ENSMUSG00000054474 | Thnsl2 | 6 | 71078160 | 71094340 | - | 0.00030631 | 0.00710525 | 13.2892515 | 14.3226191 | 14.8264934 | 11.0689115 | 14.8264934 | ZT12 | Absent_Koike2012 | Absent_Koike2012 |
| ENSMUSG00000001472 | Tcf25 | 8 | 125897724 | 125928073 | + | 0.00030658 | 0.00710525 | 17.8011593 | 18.4097227 | 20.7587522 | 18.8602766 | 20.7587522 | ZT12 | Present_Koike2012 | Absent_Koike2012 |
| ENSMUSG00000062563 | Cys1 | 12 | 25350703 | 25366678 | - | 0.00031248 | 0.00719922 | 0.21143818 | 0.2292418 | 0.45988187 | 0.42170338 | 0.45988187 | ZT12 | Absent_Koike2012 | Absent_Koike2012 |
| ENSMUSG00000072964 | Bhlhb9 | X | 132420392 | 132425620 | + | 0.00031304 | 0.00719922 | 0.31328951 | 0.38520552 | 0.65000056 | 0.29041831 | 0.65000056 | ZT12 | Absent_Koike2012 | Absent_Koike2012 |
| ENSMUSG00000032279 | Idh3a | 9 | 54434288 | 54452468 | + | 0.00031491 | 0.00721285 | 11.0437042 | 9.16748032 | 13.1246415 | 9.68105245 | 13.1246415 | ZT12 | Absent_Koike2012 | Present_Koike2012 |
| ENSMUSG00000045648 | Vwc2l | 1 | 70772289 | 70931971 | + | 0.00031823 | 0.0072735 | 0 | 0 | 0.01575656 | 0.00393411 | 0.01575656 | ZT12 | Absent_Koike2012 | Absent_Koike2012 |
| ENSMUSG00000037278 | Tmem97 | 11 | 78355319 | 78364279 | - | 0.0003179 | 0.0072735 | 38.050231 | 36.1643188 | 53.209852 | 41.3095678 | 53.209852 | ZT12 | Absent_Koike2012 | Absent_Koike2012 |
| ENSMUSG00000017664 | Slc35c2 | 2 | 165102054 | 165113369 | - | 0.00031841 | 0.0072735 | 2.76402268 | 2.87679962 | 3.27686501 | 2.38900997 | 3.27686501 | ZT12 | Present_Koike2012 | Absent_Koike2012 |
| ENSMUSG00000006304 | Arpc2 | 1 | 74283071 | 74314783 | + | 0.00031911 | 0.00728293 | 30.1903662 | 32.3870789 | 35.8248307 | 31.7891345 | 35.8248307 | ZT12 | Absent_Koike2012 | Absent_Koike2012 |
| ENSMUSG00000026663 | Atf6 | 1 | 172634588 | 172797902 | - | 0.0003219 | 0.00733338 | 9.42409125 | 11.1405913 | 13.192742 | 10.3587198 | 13.192742 | ZT12 | Present_Koike2012 | Absent_Koike2012 |
| ENSMUSG00000060376 | Bckdha | 7 | 26414871 | 26443780 | - | 0.00032576 | 0.00739484 | 73.2729798 | 83.617668 | 88.1660615 | 75.0863553 | 88.1660615 | ZT12 | Absent_Koike2012 | Absent_Koike2012 |
| ENSMUSG00000025813 | Homer2 | 7 | 88745367 | 88851811 | - | 0.00032708 | 0.00741811 | 2.21464011 | 1.78021407 | 3.54938058 | 2.68112713 | 3.54938058 | ZT12 | Present_Koike2012 | Present_Koike2012 |
| ENSMUSG00000026021 | Sumo1 | 1 | 59696278 | 59727678 | - | 0.00033018 | 0.00747567 | 10.3124704 | 10.0054783 | 13.1250297 | 11.1032165 | 13.1250297 | ZT12 | Absent_Koike2012 | Present_Koike2012 |
| ENSMUSG00000027513 | Pck1 | 2 | 172978549 | 172984774 | + | 0.00033187 | 0.00749997 | 152.833933 | 487.721052 | 739.948423 | 159.786707 | 739.948423 | ZT12 | Absent_Koike2012 | Absent_Koike2012 |
| ENSMUSG00000015013 | Trappc2l | 8 | 125135526 | 125139488 | + | 0.00033294 | 0.00750782 | 9.97362425 | 10.6200245 | 11.4478085 | 8.70628225 | 11.4478085 | ZT12 | Absent_Koike2012 | Absent_Koike2012 |
| ENSMUSG00000004394 | Tmed4 | 11 | 6170372 | 6174873 | - | 0.00033256 | 0.00750782 | 16.815948 | 16.7582244 | 19.1055602 | 17.6298504 | 19.1055602 | ZT12 | Absent_Koike2012 | Present_Koike2012 |
| ENSMUSG00000021868 | Ppif | 14 | 26513640 | 26519954 | + | 0.0003331 | 0.00750782 | 4.10313958 | 4.06304285 | 5.69556117 | 5.12154687 | 5.69556117 | ZT12 | Absent_Koike2012 | Absent_Koike2012 |
| ENSMUSG00000038683 | Pak1ip1 | 13 | 41096392 | 41108388 | + | 0.00033667 | 0.00753834 | 5.64672281 | 4.99690681 | 6.04551128 | 5.67627602 | 6.04551128 | ZT12 | Absent_Koike2012 | Absent_Koike2012 |
| ENSMUSG00000014867 | Surf4 | 2 | 26775560 | 26789448 | - | 0.00033666 | 0.00753834 | 61.7425928 | 56.5677694 | 64.93409 | 63.8576249 | 64.93409 | ZT12 | Absent_Koike2012 | Present_Koike2012 |
| ENSMUSG00000027787 | Nmd3 | 3 | 69525907 | 69552964 | + | 0.00034113 | 0.00759463 | 4.76409528 | 4.90809279 | 6.05150595 | 4.85164498 | 6.05150595 | ZT12 | Absent_Koike2012 | Present_Koike2012 |
| ENSMUSG00000034430 | Zxdc | 6 | 90319486 | 90353484 | + | 0.00034357 | 0.00764233 | 1.02535526 | 1.12387951 | 1.21153561 | 0.89116102 | 1.21153561 | ZT12 | Absent_Koike2012 | Present_Koike2012 |
| ENSMUSG00000019210 | Atp6v1e1 | 6 | 120745263 | 120772703 | - | 0.00034427 | 0.00765121 | 44.4712928 | 50.2177973 | 59.0956728 | 44.5510063 | 59.0956728 | ZT12 | Absent_Koike2012 | Absent_Koike2012 |
| ENSMUSG00000032776 | Mctp2 | 7 | 79222718 | 79451481 | - | 0.00035013 | 0.0077408 | 3.61947425 | 3.90788375 | 4.335573 | 3.3406375 | 4.335573 | ZT12 | Absent_Koike2012 | Present_Koike2012 |
| ENSMUSG00000023345 | Poc1a | 9 | 106183392 | 106252852 | + | 0.00035352 | 0.00778871 | 0.57389312 | 0.42690014 | 0.85191514 | 0.61728099 | 0.85191514 | ZT12 | Absent_Koike2012 | Absent_Koike2012 |
| ENSMUSG00000025612 | Bach1 | 16 | 87699190 | 87733591 | + | 0.00035491 | 0.00780587 | 2.92705013 | 4.62730985 | 7.86391267 | 3.861804 | 7.86391267 | ZT12 | Present_Koike2012 | Present_Koike2012 |
| ENSMUSG00000040667 | Nup88 | 11 | 70756560 | 70783475 | - | 0.00035591 | 0.00782082 | 3.43941365 | 3.61363171 | 3.97035414 | 3.25993199 | 3.97035414 | ZT12 | Present_Koike2012 | Present_Koike2012 |
| ENSMUSG00000048546 | Tob2 | 15 | 81678700 | 81688756 | - | 0.00036373 | 0.0079621 | 2.67288626 | 3.34871534 | 5.59424727 | 2.89782101 | 5.59424727 | ZT12 | Present_Koike2012 | Absent_Koike2012 |
| ENSMUSG00000024900 | Cpt1a | 19 | 3323301 | 3385732 | + | 0.00036469 | 0.00796587 | 48.7033443 | 79.3143168 | 87.1219063 | 48.626118 | 87.1219063 | ZT12 | Present_Koike2012 | Absent_Koike2012 |
| ENSMUSG00000027566 | Psma7 | 2 | 179771078 | 179777138 | - | 0.00036572 | 0.00797456 | 11.1838529 | 11.7842618 | 12.1708736 | 10.1367094 | 12.1708736 | ZT12 | Absent_Koike2012 | Absent_Koike2012 |
| ENSMUSG00000073639 | Rab18 | 18 | 6765198 | 6790320 | + | 0.00037104 | 0.00804923 | 18.9682265 | 22.2046205 | 23.2432475 | 21.3075023 | 23.2432475 | ZT12 | Absent_Koike2012 | Present_Koike2012 |
| ENSMUSG00000029273 | Sult1d1 | 5 | 87983670 | 87998052 | - | 0.00037348 | 0.0080887 | 25.5997554 | 33.2758417 | 43.7159143 | 38.8504132 | 43.7159143 | ZT12 | Present_Koike2012 | Absent_Koike2012 |
| ENSMUSG00000030515 | Tarsl2 | 7 | 72789784 | 72836977 | + | 0.00037349 | 0.0080887 | 3.73729094 | 3.93091383 | 4.39516701 | 3.79559327 | 4.39516701 | ZT12 | Absent_Koike2012 | Absent_Koike2012 |
| ENSMUSG00000045973 | Mcart1 | 4 | 45408795 | 45421633 | - | 0.00037744 | 0.00816036 | 4.41290267 | 4.68754584 | 8.02006336 | 4.12899132 | 8.02006336 | ZT12 | Present_Koike2012 | Absent_Koike2012 |
| ENSMUSG00000026358 | Rgs1 | 1 | 146091102 | 146096372 | - | 0.00038066 | 0.00822296 | 0.06190977 | 0.119195 | 0.27592074 | 0.10379111 | 0.27592074 | ZT12 | Absent_Koike2012 | Absent_Koike2012 |
| ENSMUSG00000022360 | Atad2 | 15 | 57925602 | 57966637 | - | 0.00038237 | 0.00824672 | 1.13410856 | 0.86660204 | 1.6577351 | 1.24640802 | 1.6577351 | ZT12 | Present_Koike2012 | Absent_Koike2012 |
| ENSMUSG00000028955 | Vamp3 | 4 | 150421409 | 150432072 | - | 0.00038539 | 0.00828957 | 2.94569931 | 3.47041099 | 4.06385304 | 3.29488176 | 4.06385304 | ZT12 | Present_Koike2012 | Absent_Koike2012 |
| ENSMUSG00000031629 | Mlf1ip | 8 | 47637409 | 47665361 | + | 0.00038579 | 0.00828957 | 0.04244984 | 0.03791269 | 0.14550312 | 0.02093309 | 0.14550312 | ZT12 | Absent_Koike2012 | Absent_Koike2012 |
| ENSMUSG00000032311 | Nrg4 | 9 | 55068029 | 55174651 | - | 0.00038602 | 0.00828957 | 0.91245939 | 1.12400801 | 2.19533476 | 1.38532992 | 2.19533476 | ZT12 | Present_Koike2012 | Present_Koike2012 |
| ENSMUSG00000021236 | Entpd5 | 12 | 85714807 | 85749979 | - | 0.00038508 | 0.00828957 | 24.2185259 | 21.2282672 | 31.4823784 | 24.4901805 | 31.4823784 | ZT12 | Absent_Koike2012 | Absent_Koike2012 |
| ENSMUSG00000021180 | Rps6ka5 | 12 | 101787988 | 101963238 | - | 0.00038698 | 0.00829284 | 0.25377725 | 0.39329889 | 0.54416056 | 0.42301172 | 0.54416056 | ZT12 | Absent_Koike2012 | Absent_Koike2012 |
| ENSMUSG00000070871 | Ccnyl1 | 1 | 64737029 | 64772218 | + | 0.00038715 | 0.00829284 | 1.18887241 | 1.24639094 | 1.6632689 | 1.30729646 | 1.6632689 | ZT12 | Absent_Koike2012 | Present_Koike2012 |
| ENSMUSG00000000915 | Hip1r | 5 | 124423637 | 124453220 | + | 0.00038911 | 0.00832783 | 4.15999429 | 3.46033802 | 5.40367571 | 4.00025993 | 5.40367571 | ZT12 | Absent_Koike2012 | Absent_Koike2012 |
| ENSMUSG00000032599 | Ip6k2 | 9 | 108686127 | 108706864 | + | 0.00039405 | 0.0084133 | 2.0662202 | 3.94596912 | 4.82305883 | 2.49724194 | 4.82305883 | ZT12 | Absent_Koike2012 | Absent_Koike2012 |
| ENSMUSG00000024769 | Cdc42bpg | 19 | 6306456 | 6325652 | + | 0.0003941 | 0.0084133 | 1.67506241 | 1.61207229 | 2.16009667 | 1.64338783 | 2.16009667 | ZT12 | Absent_Koike2012 | Absent_Koike2012 |
| ENSMUSG00000031608 | Galnt7 | 8 | 60002625 | 60131829 | - | 0.00039885 | 0.00848613 | 0.47856443 | 0.55261166 | 0.86794938 | 0.60998093 | 0.86794938 | ZT12 | Present_Koike2012 | Absent_Koike2012 |
| ENSMUSG00000029063 | Nadk | 4 | 154936487 | 154965110 | + | 0.00040199 | 0.0085174 | 29.5689806 | 26.9569679 | 36.2847714 | 34.6549182 | 36.2847714 | ZT12 | Absent_Koike2012 | Absent_Koike2012 |
| ENSMUSG00000016257 | Slmo2 | 2 | 174290568 | 174298582 | - | 0.00040158 | 0.0085174 | 11.953144 | 8.56767292 | 12.3619216 | 11.4664016 | 12.3619216 | ZT12 | Absent_Koike2012 | Absent_Koike2012 |
| ENSMUSG00000020538 | Srebf1 | 11 | 60012591 | 60036083 | - | 0.00040392 | 0.0085504 | 26.285003 | 21.9557032 | 47.778612 | 39.5483532 | 47.778612 | ZT12 | Present_Koike2012 | Absent_Koike2012 |
| ENSMUSG00000021707 | Dhfr | 13 | 93124738 | 93159008 | + | 0.00040422 | 0.0085504 | 4.26069425 | 5.25503025 | 5.885515 | 4.083951 | 5.885515 | ZT12 | Absent_Koike2012 | Absent_Koike2012 |
| ENSMUSG00000029629 | Phf14 | 6 | 11857809 | 12031197 | + | 0.00040597 | 0.00856735 | 2.32557148 | 2.8779143 | 3.26581182 | 2.40703833 | 3.26581182 | ZT12 | Absent_Koike2012 | Present_Koike2012 |
| ENSMUSG00000019948 | Actr6 | 10 | 89174718 | 89195040 | - | 0.00041588 | 0.00871667 | 4.0950525 | 3.550234 | 4.98035075 | 4.58920275 | 4.98035075 | ZT12 | Absent_Koike2012 | Present_Koike2012 |
| ENSMUSG00000036863 | Syde2 | 3 | 145650834 | 145684684 | + | 0.00041794 | 0.00873566 | 1.228142 | 1.0793165 | 2.118838 | 1.582642 | 2.118838 | ZT12 | Present_Koike2012 | Present_Koike2012 |
| ENSMUSG00000038312 | Edem2 | 2 | 155527413 | 155555211 | - | 0.00042051 | 0.0087567 | 11.3343291 | 10.5445736 | 13.3284403 | 12.7999878 | 13.3284403 | ZT12 | Absent_Koike2012 | Absent_Koike2012 |
| ENSMUSG00000057177 | Gsk3a | 7 | 26013277 | 26022870 | - | 0.00042273 | 0.0087886 | 6.79465054 | 7.2768249 | 7.67848932 | 6.3569894 | 7.67848932 | ZT12 | Absent_Koike2012 | Absent_Koike2012 |
| ENSMUSG00000026554 | Dcaf8 | 1 | 174078146 | 174126524 | + | 0.00042346 | 0.00879661 | 24.8533142 | 27.7460789 | 27.8338345 | 24.101347 | 27.8338345 | ZT12 | Present_Koike2012 | Absent_Koike2012 |
| ENSMUSG00000060096 | Amd-ps3 | 18 | 64608334 | 64609669 | - | 0.00042574 | 0.00880089 | 1.41652025 | 0.94668475 | 2.2101405 | 1.428927 | 2.2101405 | ZT12 | Absent_Koike2012 | Absent_Koike2012 |
| ENSMUSG00000040848 | Sft2d2 | 1 | 167104472 | 167124564 | - | 0.00042546 | 0.00880089 | 10.5812289 | 11.0420898 | 11.5258301 | 10.5663537 | 11.5258301 | ZT12 | Absent_Koike2012 | Absent_Koike2012 |
| ENSMUSG00000034189 | Hsdl1 | 8 | 122085879 | 122099100 | - | 0.00042664 | 0.00881002 | 2.50719425 | 2.40739475 | 2.96500175 | 2.74073375 | 2.96500175 | ZT12 | Absent_Koike2012 | Absent_Koike2012 |
| ENSMUSG00000032267 | Usp28 | 9 | 48793490 | 48850622 | + | 0.00042688 | 0.00881002 | 1.14293048 | 1.51684561 | 1.53644368 | 1.16994328 | 1.53644368 | ZT12 | Absent_Koike2012 | Absent_Koike2012 |
| ENSMUSG00000029321 | Slc10a6 | 5 | 104034730 | 104058422 | - | 0.000433 | 0.0089002 | 0.0026385 | 0.044359 | 0.08666425 | 0.0201335 | 0.08666425 | ZT12 | Absent_Koike2012 | Absent_Koike2012 |
| ENSMUSG00000029334 | Prkg2 | 5 | 99358792 | 99466370 | - | 0.00044866 | 0.009127 | 0.00912913 | 0.01673334 | 0.03382548 | 0.00575405 | 0.03382548 | ZT12 | Absent_Koike2012 | Absent_Koike2012 |
| ENSMUSG00000026437 | Cdk18 | 1 | 134010127 | 134036262 | - | 0.00045187 | 0.00918431 | 3.81735099 | 3.39560983 | 5.2411568 | 4.60869797 | 5.2411568 | ZT12 | Absent_Koike2012 | Absent_Koike2012 |
| ENSMUSG00000047843 | Bri3 | 5 | 145005306 | 145207626 | + | 0.00045228 | 0.00918523 | 4.50728477 | 5.58059329 | 8.18326937 | 5.55068159 | 8.18326937 | ZT12 | Absent_Koike2012 | Absent_Koike2012 |

| | | | | | | | | | | | | | | | |
|---|---|---|---|---|---|---|---|---|---|---|---|---|---|---|---|
| ENSMUSG00000049791 | Fzd4 | 7 | 96552876 | 96558620 | + | 0.00045716 | 0.009254 | 3.04960443 | 3.41598483 | 4.83888607 | 4.6241599 | 4.83888607 | ZT12 | Present_Koike2012 | Absent_Koike2012 |
| ENSMUSG00000067713 | Prkag1 | 15 | 98643229 | 98661939 | - | 0.00045764 | 0.009254 | 9.99757717 | 10.0757935 | 11.413403 | 9.96119475 | 11.413403 | ZT12 | Absent_Koike2012 | Absent_Koike2012 |
| ENSMUSG00000019820 | Utrn | 10 | 12101986 | 12589163 | - | 0.00046342 | 0.00935179 | 4.1530116 | 5.228332 | 5.30404828 | 4.64871041 | 5.30404828 | ZT12 | Absent_Koike2012 | Absent_Koike2012 |
| ENSMUSG00000027365 | Trpm7 | 2 | 126617301 | 126701966 | - | 0.00046509 | 0.00937059 | 3.30007289 | 4.53875569 | 4.65997106 | 3.46067844 | 4.65997106 | ZT12 | Present_Koike2012 | Present_Koike2012 |
| ENSMUSG00000037370 | Enpp1 | 10 | 24357720 | 24431965 | - | 0.00046674 | 0.00938905 | 1.71928171 | 1.9801588 | 2.38714002 | 1.96400739 | 2.38714002 | ZT12 | Present_Koike2012 | Absent_Koike2012 |
| ENSMUSG00000056952 | Tatdn2 | 6 | 113647044 | 113661063 | + | 0.00047419 | 0.00953128 | 5.91172053 | 5.37551238 | 7.15880959 | 6.72775327 | 7.15880959 | ZT12 | Absent_Koike2012 | Absent_Koike2012 |
| ENSMUSG00000030103 | Bhlhe40 | 6 | 108610623 | 108616919 | + | 0.00047725 | 0.00955915 | 11.8403746 | 15.1833017 | 26.1907569 | 22.2423489 | 26.1907569 | ZT12 | Present_Koike2012 | Present_Koike2012 |
| ENSMUSG00000024378 | Stard4 | 18 | 33359009 | 33373516 | - | 0.00047769 | 0.00955915 | 12.9492289 | 16.7889242 | 17.2723112 | 15.253147 | 17.2723112 | ZT12 | Absent_Koike2012 | Present_Koike2012 |
| ENSMUSG00000022456 | sep-03 | 15 | 82105365 | 82124872 | + | 0.000484 | 0.00965227 | 0.00372313 | 0.01090386 | 0.01915384 | 0.00558517 | 0.01915384 | ZT12 | Absent_Koike2012 | Absent_Koike2012 |
| ENSMUSG00000074886 | Grk6 | 13 | 55546695 | 55562286 | + | 0.00048893 | 0.00972769 | 3.77333113 | 4.04520133 | 4.65576257 | 4.03443541 | 4.65576257 | ZT12 | Absent_Koike2012 | Absent_Koike2012 |
| ENSMUSG00000028937 | Acot7 | 4 | 151552243 | 151645964 | + | 0.00048982 | 0.0097378 | 4.29159732 | 3.86222013 | 5.736489 | 5.47740255 | 5.736489 | ZT12 | Present_Koike2012 | Absent_Koike2012 |
| ENSMUSG00000070035 | Gm4885 | 7 | 79817704 | 79819164 | + | 0.00049109 | 0.00974794 | 0.7990915 | 0.910547 | 1.24853025 | 0.7767755 | 1.24853025 | ZT12 | Absent_Koike2012 | Absent_Koike2012 |
| ENSMUSG00000024870 | Rab1b | 19 | 5099207 | 5107019 | - | 0.00049171 | 0.00975258 | 26.5595843 | 26.9464053 | 29.1725678 | 24.0389433 | 29.1725678 | ZT12 | Absent_Koike2012 | Absent_Koike2012 |
| ENSMUSG00000020328 | Nudcd2 | 11 | 40547169 | 40553548 | + | 0.00049273 | 0.00976519 | 6.29616509 | 5.1716183 | 6.29616509 | 5.74740309 | 6.29616509 | ZT12 | Absent_Koike2012 | Absent_Koike2012 |
| ENSMUSG00000049550 | Clip1 | 5 | 124027804 | 124134627 | - | 0.00049636 | 0.00982173 | 3.11798043 | 3.21814546 | 4.53504522 | 3.18716197 | 4.53504522 | ZT12 | Absent_Koike2012 | Absent_Koike2012 |
| ENSMUSG00000024896 | Minpp1 | 19 | 32560259 | 32589860 | + | 0.00050138 | 0.00990584 | 9.95289525 | 11.5243598 | 12.1433895 | 9.90815825 | 12.1433895 | ZT12 | Absent_Koike2012 | Absent_Koike2012 |
| ENSMUSG00000060548 | Tnfrsf19 | 14 | 61582671 | 61665692 | - | 0.00050257 | 0.00992153 | 0.54051883 | 0.85925523 | 1.21694057 | 0.67402123 | 1.21694057 | ZT12 | Absent_Koike2012 | Absent_Koike2012 |
| ENSMUSG00000025532 | Crcp | 5 | 130505160 | 130536659 | + | 0.00050334 | 0.00992906 | 6.27097243 | 7.64033208 | 9.35334952 | 7.41540859 | 9.35334952 | ZT12 | Absent_Koike2012 | Absent_Koike2012 |
| ENSMUSG00000034024 | Cct2 | 10 | 116488054 | 116500870 | - | 0.00050619 | 0.00996988 | 40.1506493 | 37.7087478 | 43.6996195 | 41.192665 | 43.6996195 | ZT12 | Present_Koike2012 | Absent_Koike2012 |
| ENSMUSG00000028150 | Rorc | 3 | 94176716 | 94202198 | + | 5.27E-11 | 4.47E-07 | 9.36487115 | 3.08470401 | 9.16194497 | 19.0217653 | 19.0217653 | ZT18 | Present_Koike2012 | Present_Koike2012 |
| ENSMUSG00000024966 | Stip1 | 19 | 7095186 | 7114516 | - | 9.10E-11 | 5.79E-07 | 28.0985433 | 16.8007368 | 22.2288565 | 36.2329233 | 36.2329233 | ZT18 | Absent_Koike2012 | Present_Koike2012 |
| ENSMUSG00000082896 | Gm5844 | 3 | 17908806 | 17910996 | + | 3.60E-09 | 7.81E-06 | 3.9411565 | 1.78732175 | 1.983543 | 5.60344525 | 5.60344525 | ZT18 | Absent_Koike2012 | Absent_Koike2012 |
| ENSMUSG00000030357 | Fkbp4 | 6 | 128379753 | 128388695 | - | 4.13E-09 | 8.08E-06 | 50.2660738 | 30.6988963 | 35.6781592 | 61.3899836 | 61.3899836 | ZT18 | Present_Koike2012 | Present_Koike2012 |
| ENSMUSG00000007950 | Abhd8 | 8 | 73980599 | 73987556 | - | 1.20E-08 | 1.84E-05 | 2.1562925 | 1.178634 | 1.86087425 | 3.0329185 | 3.0329185 | ZT18 | Absent_Koike2012 | Absent_Koike2012 |
| ENSMUSG00000055866 | Per2 | 1 | 93312561 | 93355907 | - | 2.21E-08 | 2.95E-05 | 1.57462335 | 0.87842135 | 4.15582411 | 5.11541537 | 5.11541537 | ZT18 | Present_Koike2012 | Present_Koike2012 |
| ENSMUSG00000015656 | Hspa8 | 9 | 40609067 | 40613283 | + | 2.45E-08 | 3.09E-05 | 177.100101 | 78.5854634 | 152.389588 | 178.252291 | 178.252291 | ZT18 | Absent_Koike2012 | Absent_Koike2012 |
| ENSMUSG00000016028 | Celsr1 | 15 | 85729188 | 85864207 | - | 3.02E-08 | 3.50E-05 | 2.08100816 | 0.83441597 | 1.18757426 | 2.50799196 | 2.50799196 | ZT18 | Absent_Koike2012 | Absent_Koike2012 |
| ENSMUSG00000048878 | Hexim1 | 11 | 102977639 | 102981039 | + | 3.58E-08 | 3.80E-05 | 9.657663 | 5.82345325 | 7.940109 | 9.6902355 | 9.6902355 | ZT18 | Absent_Koike2012 | Absent_Koike2012 |
| ENSMUSG00000021037 | Ahsa1 | 12 | 88607696 | 88614947 | + | 3.53E-08 | 3.80E-05 | 44.5765475 | 34.2881938 | 36.892249 | 52.8229095 | 52.8229095 | ZT18 | Absent_Koike2012 | Absent_Koike2012 |
| ENSMUSG00000050677 | Ccdc96 | 5 | 36827237 | 36830820 | + | 3.83E-08 | 3.85E-05 | 0.06458275 | 0.0126715 | 0.057713 | 0.08651075 | 0.08651075 | ZT18 | Absent_Koike2012 | Absent_Koike2012 |
| ENSMUSG00000091282 | Gm17431 | 5 | 5781530 | 5783636 | + | 4.07E-08 | 3.85E-05 | 94.1698588 | 41.2234813 | 80.753008 | 94.2488558 | 94.2488558 | ZT18 | Absent_Koike2012 | Absent_Koike2012 |
| ENSMUSG00000020038 | Cry1 | 10 | 84594447 | 84647799 | - | 5.34E-08 | 4.38E-05 | 8.36009775 | 1.70469475 | 3.4407245 | 8.96204125 | 8.96204125 | ZT18 | Present_Koike2012 | Present_Koike2012 |
| ENSMUSG00000059814 | Gm8355 | 10 | 14465084 | 14467190 | - | 6.43E-08 | 4.67E-05 | 29.1238633 | 13.0734258 | 25.3210053 | 29.329724 | 29.329724 | ZT18 | Absent_Koike2012 | Absent_Koike2012 |
| ENSMUSG00000028405 | Aco1 | 4 | 40090114 | 40146042 | + | 8.90E-08 | 5.69E-05 | 47.3412138 | 39.4963158 | 48.1917405 | 52.1185618 | 52.1185618 | ZT18 | Absent_Koike2012 | Present_Koike2012 |
| ENSMUSG00000016239 | Lonrf3 | X | 33868348 | 33902336 | + | 1.44E-07 | 8.01E-05 | 2.00939844 | 0.43877643 | 1.37423586 | 2.06798032 | 2.06798032 | ZT18 | Present_Koike2012 | Present_Koike2012 |
| ENSMUSG00000011179 | Odc1 | 12 | 17551679 | 17558308 | + | 1.64E-07 | 8.01E-05 | 32.8182766 | 20.5267472 | 28.612338 | 41.8680963 | 41.8680963 | ZT18 | Present_Koike2012 | Absent_Koike2012 |
| ENSMUSG00000081058 | Hist2h3c2 | 3 | 96042040 | 96043050 | - | 1.93E-07 | 8.15E-05 | 1.54534425 | 0.745427 | 0.73123825 | 2.13984625 | 2.13984625 | ZT18 | Absent_Koike2012 | Absent_Koike2012 |
| ENSMUSG00000027488 | Snta1 | 2 | 154202049 | 154233835 | - | 2.07E-07 | 8.34E-05 | 3.03839408 | 2.47780622 | 3.69808211 | 4.17151999 | 4.17151999 | ZT18 | Absent_Koike2012 | Absent_Koike2012 |
| ENSMUSG00000023944 | Hsp90ab1 | 17 | 45704724 | 45710220 | - | 2.25E-07 | 8.42E-05 | 159.670125 | 119.269453 | 115.709515 | 196.881469 | 196.881469 | ZT18 | Absent_Koike2012 | Absent_Koike2012 |
| ENSMUSG00000034880 | Mrpl34 | 8 | 73988825 | 73989651 | + | 3.10E-07 | 0.00010108 | 26.2181025 | 19.173653 | 21.2985165 | 27.207663 | 27.207663 | ZT18 | Absent_Koike2012 | Absent_Koike2012 |
| ENSMUSG00000025869 | Nop16 | 13 | 54685546 | 54691451 | - | 3.78E-07 | 0.00011155 | 10.0331854 | 6.69455033 | 7.7585351 | 11.8031192 | 11.8031192 | ZT18 | Absent_Koike2012 | Absent_Koike2012 |
| ENSMUSG00000059248 | sep-09 | 11 | 117060975 | 117223639 | + | 4.53E-07 | 0.00012127 | 5.60928727 | 4.2816366 | 3.40320766 | 5.70285649 | 5.70285649 | ZT18 | Absent_Koike2012 | Absent_Koike2012 |
| ENSMUSG00000040181 | Fmo1 | 1 | 164759692 | 164796679 | - | 5.55E-07 | 0.00013341 | 25.4487204 | 18.1300598 | 25.3233613 | 31.9063838 | 31.9063838 | ZT18 | Present_Koike2012 | Present_Koike2012 |
| ENSMUSG00000025757 | Hspa4l | 3 | 40548417 | 40600025 | + | 5.68E-07 | 0.00013341 | 3.00340816 | 2.01691424 | 1.71422307 | 4.45454842 | 4.45454842 | ZT18 | Present_Koike2012 | Absent_Koike2012 |
| ENSMUSG00000030170 | Wnt5b | 6 | 119382549 | 119494336 | - | 6.26E-07 | 0.00014128 | 1.62158001 | 0.92553629 | 1.99533479 | 2.92503861 | 2.92503861 | ZT18 | Absent_Koike2012 | Present_Koike2012 |
| ENSMUSG00000026922 | Agpat2 | 2 | 26448577 | 26459937 | - | 6.27E-07 | 0.00014128 | 84.3706418 | 54.213245 | 57.2280098 | 85.8531339 | 85.8531339 | ZT18 | Present_Koike2012 | Absent_Koike2012 |
| ENSMUSG00000071653 | 1810009A15 | 19 | 8963393 | 8965230 | + | 6.58E-07 | 0.00014583 | 16.7189765 | 12.7782853 | 12.8839858 | 18.224308 | 18.224308 | ZT18 | Absent_Koike2012 | Absent_Koike2012 |
| ENSMUSG00000028766 | Alpl | 4 | 137297648 | 137352299 | - | 7.55E-07 | 0.00015625 | 2.15713107 | 1.62063848 | 3.12795408 | 3.82841979 | 3.82841979 | ZT18 | Absent_Koike2012 | Absent_Koike2012 |
| ENSMUSG00000021361 | Tmem14c | 13 | 41111619 | 41117944 | + | 7.88E-07 | 0.00016168 | 80.2635898 | 65.0001268 | 79.7298525 | 89.5406955 | 89.5406955 | ZT18 | Absent_Koike2012 | Absent_Koike2012 |
| ENSMUSG00000050777 | Tmem37 | 1 | 121963954 | 121970651 | - | 8.85E-07 | 0.00017017 | 31.9900673 | 22.3652085 | 31.1394748 | 39.073125 | 39.073125 | ZT18 | Absent_Koike2012 | Absent_Koike2012 |
| ENSMUSG00000025732 | Fam195a | 17 | 26000643 | 26005683 | - | 8.80E-07 | 0.00017017 | 15.9713765 | 10.9128035 | 16.76465 | 18.9787448 | 18.9787448 | ZT18 | Absent_Koike2012 | Absent_Koike2012 |
| ENSMUSG00000029657 | Hsph1 | 5 | 150416862 | 150438951 | - | 1.03E-06 | 0.00018718 | 6.48177661 | 1.73915697 | 4.07435424 | 7.68346531 | 7.68346531 | ZT18 | Present_Koike2012 | Present_Koike2012 |
| ENSMUSG00000005374 | Tbl2 | 5 | 135625527 | 135639743 | + | 1.35E-06 | 0.00021037 | 2.89041567 | 1.84859922 | 2.93483038 | 2.94836889 | 2.94836889 | ZT18 | Absent_Koike2012 | Absent_Koike2012 |
| ENSMUSG00000024150 | Mcfd2 | 17 | 87653998 | 87665275 | - | 1.46E-06 | 0.00022196 | 50.7325122 | 36.9144106 | 43.8994383 | 52.8697761 | 52.8697761 | ZT18 | Present_Koike2012 | Absent_Koike2012 |
| ENSMUSG00000048481 | Mypop | 7 | 19576594 | 19587115 | + | 1.69E-06 | 0.00024774 | 0.48017533 | 0.25712585 | 0.3112716 | 0.68019688 | 0.68019688 | ZT18 | Absent_Koike2012 | Absent_Koike2012 |
| ENSMUSG00000001416 | Cct3 | 3 | 88101038 | 88125689 | + | 1.67E-06 | 0.00024774 | 14.3942008 | 11.5111987 | 12.112283 | 14.5673321 | 14.5673321 | ZT18 | Absent_Koike2012 | Absent_Koike2012 |
| ENSMUSG00000020087 | Tysnd1 | 10 | 61158262 | 61165521 | + | 1.77E-06 | 0.0002547 | 13.817498 | 9.647824 | 11.7000395 | 13.948663 | 13.948663 | ZT18 | Absent_Koike2012 | Absent_Koike2012 |
| ENSMUSG00000027808 | Serp1 | 3 | 58325891 | 58329795 | - | 1.99E-06 | 0.00027709 | 118.677153 | 95.1120873 | 135.804783 | 155.206207 | 155.206207 | ZT18 | Absent_Koike2012 | Absent_Koike2012 |
| ENSMUSG00000025533 | Asl | 5 | 130487128 | 130501002 | - | 2.13E-06 | 0.00029178 | 68.1688149 | 47.2941607 | 81.8804393 | 111.176449 | 111.176449 | ZT18 | Absent_Koike2012 | Present_Koike2012 |
| ENSMUSG00000021270 | Hsp90aa1 | 12 | 111928815 | 111940938 | - | 2.69E-06 | 0.00034578 | 45.1380884 | 18.290506 | 21.6107738 | 64.6381237 | 64.6381237 | ZT18 | Present_Koike2012 | Present_Koike2012 |
| ENSMUSG00000039680 | Mrps6 | 16 | 92058515 | 92112472 | + | 3.72E-06 | 0.00042815 | 1.83037218 | 1.12356268 | 1.47811061 | 2.08817824 | 2.08817824 | ZT18 | Absent_Koike2012 | Present_Koike2012 |
| ENSMUSG00000041119 | Pde9a | 17 | 31523179 | 31613255 | + | 4.68E-06 | 0.0004819 | 5.03712701 | 4.05363328 | 2.48499235 | 5.34569195 | 5.34569195 | ZT18 | Present_Koike2012 | Present_Koike2012 |
| ENSMUSG00000051652 | Lrrc3 | 10 | 77360320 | 77365281 | - | 5.21E-06 | 0.00050374 | 16.4444963 | 14.131429 | 13.0369565 | 19.3859893 | 19.3859893 | ZT18 | Present_Koike2012 | Absent_Koike2012 |
| ENSMUSG00000036427 | Gpi1 | 7 | 34987141 | 35015308 | - | 5.19E-06 | 0.00050374 | 23.8398681 | 17.9693657 | 25.7382431 | 27.6029821 | 27.6029821 | ZT18 | Absent_Koike2012 | Absent_Koike2012 |
| ENSMUSG00000021484 | Lman2 | 13 | 55445194 | 55464144 | - | 5.22E-06 | 0.00050374 | 46.7997703 | 37.940995 | 40.89122 | 48.4834528 | 48.4834528 | ZT18 | Present_Koike2012 | Absent_Koike2012 |
| ENSMUSG00000004270 | Lpcat3 | 6 | 124613045 | 124654436 | - | 7.19E-06 | 0.00063372 | 11.0611474 | 14.9248207 | 13.6826294 | 14.9333468 | 14.9333468 | ZT18 | Present_Koike2012 | Present_Koike2012 |
| ENSMUSG00000025511 | Tspan4 | 7 | 148661139 | 148679326 | + | 7.24E-06 | 0.00063372 | 9.42556448 | 7.28469065 | 14.914729 | 18.281032 | 18.281032 | ZT18 | Absent_Koike2012 | Present_Koike2012 |
| ENSMUSG00000031832 | Taf1c | 8 | 122121875 | 122129122 | - | 8.06E-06 | 0.00068033 | 0.52205045 | 0.25853632 | 0.40720737 | 0.61175546 | 0.61175546 | ZT18 | Absent_Koike2012 | Absent_Koike2012 |
| ENSMUSG00000046794 | Ppp1r3b | 8 | 36438795 | 36451178 | + | 8.04E-06 | 0.00068033 | 59.8188255 | 14.8566039 | 58.4253991 | 79.0844267 | 79.0844267 | ZT18 | Present_Koike2012 | Present_Koike2012 |

| | | | | | | | | | | | | | | | |
|---|---|---|---|---|---|---|---|---|---|---|---|---|---|---|---|
| ENSMUSG00000090038 | Gm16573 | 14 | 57226098 | 57239350 | - | 8.34E-06 | 0.0006956 | 1.72052183 | 1.00062762 | 2.26684966 | 2.52120649 | 2.52120649 | ZT18 | Absent_Koike2012 | Absent_Koike2012 |
| ENSMUSG00000061479 | Snrpa | 7 | 27972024 | 27981290 | - | 8.59E-06 | 0.00070511 | 5.18150146 | 4.5651672 | 4.64806175 | 7.15247314 | 7.15247314 | ZT18 | Absent_Koike2012 | Absent_Koike2012 |
| ENSMUSG00000030835 | Nomo1 | 7 | 53289066 | 53339582 | + | 8.72E-06 | 0.00071011 | 8.89527 | 7.79118625 | 9.567218 | 10.0503075 | 10.0503075 | ZT18 | Absent_Koike2012 | Absent_Koike2012 |
| ENSMUSG00000018906 | P4ha2 | 11 | 53913597 | 53945167 | + | 9.31E-06 | 0.00073809 | 0.59005479 | 0.47277672 | 0.84104873 | 1.07771554 | 1.07771554 | ZT18 | Absent_Koike2012 | Absent_Koike2012 |
| ENSMUSG00000051615 | Rap2a | 14 | 120877666 | 120906416 | + | 1.00E-05 | 0.0007797 | 6.20787573 | 4.38803459 | 7.33537814 | 8.32604043 | 8.32604043 | ZT18 | Absent_Koike2012 | Absent_Koike2012 |
| ENSMUSG00000028344 | Invs | 4 | 48292632 | 48444826 | + | 1.02E-05 | 0.00079225 | 1.10110356 | 0.87504858 | 0.92662947 | 1.32008113 | 1.32008113 | ZT18 | Absent_Koike2012 | Present_Koike2012 |
| ENSMUSG00000028671 | Gale | 4 | 135519642 | 135524093 | + | 1.14E-05 | 0.00084346 | 7.31216379 | 4.33467418 | 3.51950581 | 7.8052303 | 7.8052303 | ZT18 | Present_Koike2012 | Absent_Koike2012 |
| ENSMUSG00000043067 | Dpy19l1 | 9 | 24216220 | 24307584 | - | 1.39E-05 | 0.00093788 | 10.207578 | 7.02142056 | 9.30952743 | 11.0983314 | 11.0983314 | ZT18 | Present_Koike2012 | Present_Koike2012 |
| ENSMUSG00000001440 | Kpnb1 | 11 | 97021028 | 97049195 | - | 1.43E-05 | 0.0009538 | 15.90527 | 13.230897 | 14.1992348 | 16.7296741 | 16.7296741 | ZT18 | Absent_Koike2012 | Absent_Koike2012 |
| ENSMUSG00000029556 | Hnf1a | 5 | 115398370 | 115421076 | - | 1.48E-05 | 0.00097445 | 4.20666125 | 4.1043405 | 4.01629776 | 5.078915 | 5.078915 | ZT18 | Absent_Koike2012 | Absent_Koike2012 |
| ENSMUSG00000024118 | 1600002H07 | 17 | 24351999 | 24357796 | - | 1.48E-05 | 0.00097445 | 17.23285 | 6.99529093 | 17.9443068 | 18.6240386 | 18.6240386 | ZT18 | Absent_Koike2012 | Absent_Koike2012 |
| ENSMUSG00000055912 | Tmem150a | 6 | 72305441 | 72309756 | + | 1.49E-05 | 0.00097915 | 32.9453952 | 26.5946397 | 26.1088174 | 37.9654315 | 37.9654315 | ZT18 | Absent_Koike2012 | Present_Koike2012 |
| ENSMUSG00000019055 | Plod1 | 4 | 147283862 | 147310876 | - | 1.65E-05 | 0.00105034 | 5.74554187 | 4.65848709 | 5.41645156 | 6.46066622 | 6.46066622 | ZT18 | Absent_Koike2012 | Absent_Koike2012 |
| ENSMUSG00000026203 | Dnajb2 | 1 | 75233040 | 75241379 | + | 1.73E-05 | 0.0010811 | 10.9641142 | 7.72246772 | 8.65485438 | 12.4407629 | 12.4407629 | ZT18 | Absent_Koike2012 | Absent_Koike2012 |
| ENSMUSG00000017204 | Gsdma | 11 | 98525665 | 98539022 | + | 1.82E-05 | 0.00112572 | 0 | 0 | 0 | 0.00761625 | 0.00761625 | ZT18 | Absent_Koike2012 | Absent_Koike2012 |
| ENSMUSG00000029560 | Snx8 | 5 | 140816258 | 140865216 | - | 1.87E-05 | 0.00113764 | 3.96627597 | 2.85957693 | 4.01181598 | 4.19894938 | 4.19894938 | ZT18 | Absent_Koike2012 | Absent_Koike2012 |
| ENSMUSG00000027933 | Ints3 | 3 | 90195310 | 90237558 | - | 1.92E-05 | 0.00115918 | 5.06445985 | 4.08501581 | 4.01614011 | 5.15313449 | 5.15313449 | ZT18 | Present_Koike2012 | Absent_Koike2012 |
| ENSMUSG00000025534 | Gusb | 5 | 130464881 | 130478919 | - | 1.94E-05 | 0.00116159 | 4.81686768 | 4.10388581 | 5.29495704 | 5.97184445 | 5.97184445 | ZT18 | Absent_Koike2012 | Absent_Koike2012 |
| ENSMUSG00000058624 | Gda | 19 | 21465797 | 21547935 | - | 1.95E-05 | 0.00116464 | 4.90165288 | 3.46716851 | 6.22398485 | 7.75375346 | 7.75375346 | ZT18 | Present_Koike2012 | Present_Koike2012 |
| ENSMUSG00000091898 | Tnnc1 | 14 | 31983021 | 32024915 | + | 2.04E-05 | 0.00121103 | 0.28616099 | 0.08896969 | 0.41378682 | 0.89048167 | 0.89048167 | ZT18 | Absent_Koike2012 | Absent_Koike2012 |
| ENSMUSG00000010936 | Vac14 | 8 | 113142503 | 113244296 | + | 2.25E-05 | 0.00130321 | 3.88914096 | 3.46791808 | 5.51141292 | 5.94994206 | 5.94994206 | ZT18 | Absent_Koike2012 | Present_Koike2012 |
| ENSMUSG00000023262 | Acy1 | 9 | 106335327 | 106340567 | - | 2.26E-05 | 0.00130321 | 11.8934433 | 12.0362275 | 15.9198685 | 16.128625 | 16.128625 | ZT18 | Absent_Koike2012 | Absent_Koike2012 |
| ENSMUSG00000034126 | Pomt2 | 12 | 88447816 | 88488852 | - | 2.25E-05 | 0.00130321 | 4.752895 | 4.365054 | 4.89285 | 5.26091925 | 5.26091925 | ZT18 | Absent_Koike2012 | Absent_Koike2012 |
| ENSMUSG00000025934 | Gsta3 | 1 | 21230670 | 21255442 | + | 2.36E-05 | 0.00132639 | 844.206791 | 598.336111 | 746.255981 | 948.421498 | 948.421498 | ZT18 | Present_Koike2012 | Present_Koike2012 |
| ENSMUSG00000074521 | Gm14327 | 2 | 177631801 | 177640489 | + | 2.38E-05 | 0.00133072 | 1.06570331 | 0.62206746 | 1.3398929 | 1.66635498 | 1.66635498 | ZT18 | Absent_Koike2012 | Absent_Koike2012 |
| ENSMUSG00000020496 | Rnf187 | 11 | 58745790 | 58752418 | - | 2.41E-05 | 0.00134251 | 25.8766413 | 22.492314 | 25.857926 | 27.4391195 | 27.4391195 | ZT18 | Absent_Koike2012 | Absent_Koike2012 |
| ENSMUSG00000017707 | Serinc3 | 2 | 163449008 | 163470867 | - | 2.52E-05 | 0.0013867 | 43.0293029 | 41.5047642 | 49.4003352 | 49.6768246 | 49.6768246 | ZT18 | Absent_Koike2012 | Absent_Koike2012 |
| ENSMUSG00000021504 | B4galt7 | 13 | 55701257 | 55711804 | + | 2.60E-05 | 0.00142529 | 2.96399936 | 2.03665417 | 3.04488206 | 3.14320791 | 3.14320791 | ZT18 | Absent_Koike2012 | Absent_Koike2012 |
| ENSMUSG00000028343 | Erp44 | 4 | 48206195 | 48292430 | - | 2.66E-05 | 0.00143856 | 14.4175732 | 13.2866366 | 15.4187647 | 15.8736898 | 15.8736898 | ZT18 | Absent_Koike2012 | Present_Koike2012 |
| ENSMUSG00000084936 | Gm11583 | 11 | 97022304 | 97022989 | + | 2.74E-05 | 0.00146576 | 5.58250775 | 4.33448275 | 4.3738165 | 6.4097195 | 6.4097195 | ZT18 | Absent_Koike2012 | Absent_Koike2012 |
| ENSMUSG00000035181 | Heatr5a | 12 | 52976860 | 53072308 | - | 2.76E-05 | 0.00147042 | 2.8242005 | 2.610323 | 2.92501625 | 3.240753 | 3.240753 | ZT18 | Absent_Koike2012 | Present_Koike2012 |
| ENSMUSG00000040471 | Ggt6 | 11 | 72249028 | 72252067 | + | 2.81E-05 | 0.00148973 | 3.77964319 | 2.88801041 | 3.09593666 | 4.53263312 | 4.53263312 | ZT18 | Absent_Koike2012 | Absent_Koike2012 |
| ENSMUSG00000036083 | Slc17a3 | 13 | 23931303 | 23952583 | + | 2.89E-05 | 0.00150775 | 4.94498571 | 3.41412681 | 3.30319204 | 5.69785314 | 5.69785314 | ZT18 | Present_Koike2012 | Absent_Koike2012 |
| ENSMUSG00000020964 | Sel1l | 12 | 93044483 | 93087597 | - | 2.90E-05 | 0.00151047 | 13.6465129 | 10.9005096 | 12.2220219 | 15.3637472 | 15.3637472 | ZT18 | Absent_Koike2012 | Present_Koike2012 |
| ENSMUSG00000031232 | Magt1 | X | 103163423 | 103207245 | - | 2.98E-05 | 0.00154335 | 16.4400003 | 14.1859767 | 16.9610523 | 20.3349257 | 20.3349257 | ZT18 | Absent_Koike2012 | Absent_Koike2012 |
| ENSMUSG00000035863 | Palm | 10 | 79256317 | 79283641 | + | 3.04E-05 | 0.00156095 | 0.57711225 | 0.33976774 | 0.48024903 | 0.75446837 | 0.75446837 | ZT18 | Absent_Koike2012 | Absent_Koike2012 |
| ENSMUSG00000027597 | Ahcy | 2 | 154885046 | 154900233 | - | 3.06E-05 | 0.00156349 | 427.807759 | 343.23015 | 454.305366 | 543.747999 | 543.747999 | ZT18 | Present_Koike2012 | Present_Koike2012 |
| ENSMUSG00000074277 | Phldb3 | 7 | 25396347 | 25414307 | + | 3.16E-05 | 0.00160506 | 1.962578 | 1.64245125 | 1.40530575 | 2.2427505 | 2.2427505 | ZT18 | Absent_Koike2012 | Absent_Koike2012 |
| ENSMUSG00000023272 | Creld2 | 15 | 88650076 | 88657111 | + | 3.45E-05 | 0.00171445 | 27.5136968 | 12.1452555 | 13.6530953 | 32.72303 | 32.72303 | ZT18 | Absent_Koike2012 | Absent_Koike2012 |
| ENSMUSG00000014226 | Cacybp | 1 | 162133056 | 162142908 | - | 3.61E-05 | 0.00177203 | 15.7343045 | 11.0116133 | 13.9992808 | 18.6699383 | 18.6699383 | ZT18 | Absent_Koike2012 | Absent_Koike2012 |
| ENSMUSG00000058056 | Palld | 8 | 63992043 | 64381487 | - | 3.66E-05 | 0.00178344 | 1.25827983 | 0.67510078 | 0.62327087 | 1.36262823 | 1.36262823 | ZT18 | Absent_Koike2012 | Present_Koike2012 |
| ENSMUSG00000022571 | Pycrl | 15 | 75746893 | 75751990 | - | 3.66E-05 | 0.00178344 | 10.1988183 | 8.024944 | 9.65763725 | 11.5518033 | 11.5518033 | ZT18 | Absent_Koike2012 | Absent_Koike2012 |
| ENSMUSG00000046388 | Gm9806 | 19 | 16855037 | 16855593 | + | 4.10E-05 | 0.00192747 | 0.63398275 | 0.59374025 | 1.097898 | 1.18533025 | 1.18533025 | ZT18 | Absent_Koike2012 | Absent_Koike2012 |
| ENSMUSG00000028410 | Dnaja1 | 4 | 40669183 | 40684182 | + | 4.15E-05 | 0.00194692 | 18.9148181 | 11.8501499 | 18.983265 | 20.5971483 | 20.5971483 | ZT18 | Present_Koike2012 | Present_Koike2012 |
| ENSMUSG00000062070 | Pgk1 | X | 103382439 | 103399038 | + | 4.23E-05 | 0.0019537 | 141.023274 | 108.810613 | 129.01962 | 142.794514 | 142.794514 | ZT18 | Absent_Koike2012 | Present_Koike2012 |
| ENSMUSG00000025130 | P4hb | 11 | 120421612 | 120434567 | - | 4.44E-05 | 0.00201826 | 379.869355 | 313.742715 | 363.332315 | 388.518949 | 388.518949 | ZT18 | Present_Koike2012 | Present_Koike2012 |
| ENSMUSG00000020097 | Sgpl1 | 10 | 60561390 | 60610451 | - | 4.54E-05 | 0.00205007 | 12.1033963 | 10.3566498 | 7.6317572 | 12.9122529 | 12.9122529 | ZT18 | Present_Koike2012 | Absent_Koike2012 |
| ENSMUSG00000025574 | Tk1 | 11 | 117676840 | 117687406 | - | 4.61E-05 | 0.00206788 | 1.68360764 | 1.44400652 | 2.29373968 | 2.48897826 | 2.48897826 | ZT18 | Absent_Koike2012 | Absent_Koike2012 |
| ENSMUSG00000006705 | Pknox1 | 17 | 31701694 | 31744625 | + | 4.65E-05 | 0.00207572 | 1.94333801 | 1.75249882 | 1.51910095 | 2.16608176 | 2.16608176 | ZT18 | Absent_Koike2012 | Absent_Koike2012 |
| ENSMUSG00000071644 | Eef1g | 19 | 9041531 | 9052969 | + | 4.81E-05 | 0.00212382 | 49.5281368 | 41.7938502 | 46.8068928 | 50.8979115 | 50.8979115 | ZT18 | Absent_Koike2012 | Absent_Koike2012 |
| ENSMUSG00000029186 | Pi4k2b | 5 | 53132826 | 53160583 | + | 4.84E-05 | 0.00212642 | 6.0859019 | 5.94929303 | 8.1763574 | 8.90977395 | 8.90977395 | ZT18 | Absent_Koike2012 | Absent_Koike2012 |
| ENSMUSG00000036181 | Hist1h1c | 13 | 23830676 | 23832235 | + | 4.85E-05 | 0.00212642 | 37.2440073 | 23.9672488 | 25.9520208 | 46.8968508 | 46.8968508 | ZT18 | Present_Koike2012 | Absent_Koike2012 |
| ENSMUSG00000020781 | Tsen54 | 11 | 115676038 | 115684408 | + | 4.88E-05 | 0.00213174 | 1.59237595 | 1.09138285 | 1.35311478 | 1.6668512 | 1.6668512 | ZT18 | Absent_Koike2012 | Absent_Koike2012 |
| ENSMUSG00000004951 | Hspb1 | 5 | 136363789 | 136365433 | + | 5.55E-05 | 0.00231861 | 13.6184917 | 6.02901652 | 5.20059501 | 21.3744594 | 21.3744594 | ZT18 | Absent_Koike2012 | Absent_Koike2012 |
| ENSMUSG00000001700 | Gramd3 | 18 | 56559991 | 56663446 | + | 5.54E-05 | 0.00231861 | 7.06935632 | 9.50961544 | 8.71825295 | 10.2608005 | 10.2608005 | ZT18 | Present_Koike2012 | Absent_Koike2012 |
| ENSMUSG00000037470 | Uggt1 | 1 | 36198375 | 36301150 | - | 5.60E-05 | 0.00232321 | 11.1827803 | 10.4585688 | 11.8267633 | 13.1272718 | 13.1272718 | ZT18 | Absent_Koike2012 | Present_Koike2012 |
| ENSMUSG00000036285 | Noa1 | 5 | 77723199 | 77739109 | - | 5.89E-05 | 0.00239217 | 1.64293622 | 1.17110218 | 1.62125961 | 1.68025605 | 1.68025605 | ZT18 | Absent_Koike2012 | Absent_Koike2012 |
| ENSMUSG00000032115 | Hyou1 | 9 | 44187573 | 44200452 | + | 5.91E-05 | 0.00239656 | 20.0024568 | 15.2054802 | 15.8599437 | 26.5106415 | 26.5106415 | ZT18 | Absent_Koike2012 | Absent_Koike2012 |
| ENSMUSG00000004460 | Dnajb11 | 16 | 22857918 | 22879707 | + | 6.45E-05 | 0.00254501 | 18.6799445 | 14.7693613 | 14.5639066 | 21.0471901 | 21.0471901 | ZT18 | Present_Koike2012 | Absent_Koike2012 |
| ENSMUSG00000030872 | Gga2 | 7 | 129130236 | 129164736 | - | 6.66E-05 | 0.002592 | 1.21755723 | 0.89267077 | 1.1697008 | 1.72738473 | 1.72738473 | ZT18 | Absent_Koike2012 | Absent_Koike2012 |
| ENSMUSG00000020823 | Sec14l1 | 11 | 116976482 | 117020582 | + | 6.64E-05 | 0.002592 | 1.55964353 | 1.10441122 | 1.44808357 | 1.57256164 | 1.57256164 | ZT18 | Absent_Koike2012 | Absent_Koike2012 |
| ENSMUSG00000013629 | Cad | 5 | 31357153 | 31380852 | + | 7.22E-05 | 0.00272763 | 1.70919437 | 1.25691301 | 1.73964849 | 2.31137868 | 2.31137868 | ZT18 | Absent_Koike2012 | Absent_Koike2012 |
| ENSMUSG00000020368 | Canx | 11 | 50107969 | 50139175 | - | 7.21E-05 | 0.00272763 | 117.808699 | 107.088702 | 116.492511 | 124.752771 | 124.752771 | ZT18 | Absent_Koike2012 | Absent_Koike2012 |
| ENSMUSG00000023963 | Cyp39a1 | 17 | 43804374 | 43888380 | + | 7.22E-05 | 0.00272763 | 6.50305462 | 4.79212615 | 8.35369132 | 9.50552014 | 9.50552014 | ZT18 | Absent_Koike2012 | Present_Koike2012 |
| ENSMUSG00000074207 | Adh1 | 3 | 137923955 | 137953662 | + | 7.48E-05 | 0.00280186 | 611.64526 | 455.08166 | 693.030811 | 738.797565 | 738.797565 | ZT18 | Present_Koike2012 | Present_Koike2012 |
| ENSMUSG00000038374 | Rbm8a | 3 | 96433851 | 96437702 | + | 7.69E-05 | 0.00283581 | 5.92538875 | 4.96848275 | 5.288907 | 6.265945 | 6.265945 | ZT18 | Absent_Koike2012 | Absent_Koike2012 |
| ENSMUSG00000026870 | D730039F16 | 2 | 34729916 | 34747652 | + | 8.01E-05 | 0.00291727 | 4.62528721 | 4.1666923 | 3.95045154 | 5.09214502 | 5.09214502 | ZT18 | Present_Koike2012 | Present_Koike2012 |
| ENSMUSG00000008604 | Ubqln4 | 3 | 88357638 | 88373647 | + | 8.46E-05 | 0.00302813 | 7.90475725 | 5.89443625 | 6.9945545 | 8.42390725 | 8.42390725 | ZT18 | Absent_Koike2012 | Absent_Koike2012 |
| ENSMUSG00000051391 | Ywhag | 5 | 136384279 | 136410486 | - | 8.55E-05 | 0.00305005 | 11.375742 | 8.97612063 | 12.666151 | 13.6164328 | 13.6164328 | ZT18 | Absent_Koike2012 | Present_Koike2012 |

| | | | | | | | | | | | | | | | |
|---|---|---|---|---|---|---|---|---|---|---|---|---|---|---|---|
| ENSMUSG00000025226 | Fbxl15 | 19 | 46402674 | 46404936 | + | 8.99E-05 | 0.00316614 | 1.64435625 | 1.1384145 | 1.335 | 1.9192815 | 1.9192815 | ZT18 | Absent_Koike2012 | Absent_Koike2012 |
| ENSMUSG00000032392 | Parp16 | 9 | 65062497 | 65087026 | + | 9.04E-05 | 0.00317281 | 3.30766055 | 2.75661443 | 3.80503196 | 4.71260897 | 4.71260897 | ZT18 | Present_Koike2012 | Absent_Koike2012 |
| ENSMUSG00000024856 | Cdk2ap2 | 19 | 4097182 | 4099019 | + | 9.18E-05 | 0.00321327 | 15.0368301 | 10.5744617 | 12.4442826 | 15.9974686 | 15.9974686 | ZT18 | Absent_Koike2012 | Absent_Koike2012 |
| ENSMUSG00000019935 | Slc17a8 | 10 | 89036765 | 89083998 | - | 9.29E-05 | 0.00324189 | 1.94746904 | 0.56295289 | 0.89303662 | 2.03656831 | 2.03656831 | ZT18 | Present_Koike2012 | Absent_Koike2012 |
| ENSMUSG00000034807 | Glt25d1 | 8 | 74134923 | 74148808 | + | 9.77E-05 | 0.00335182 | 14.7910595 | 12.5796353 | 14.2424125 | 15.8614618 | 15.8614618 | ZT18 | Absent_Koike2012 | Absent_Koike2012 |
| ENSMUSG00000039632 | Ccdc151 | 9 | 21794315 | 21807078 | - | 0.00010421 | 0.003517 | 0.59758084 | 0.31095178 | 0.46128475 | 0.92365084 | 0.92365084 | ZT18 | Absent_Koike2012 | Absent_Koike2012 |
| ENSMUSG00000007646 | Rad51c | 11 | 87190147 | 87218456 | - | 0.00010659 | 0.00357359 | 0.2428247 | 0.1087885 | 0.16852305 | 0.39512503 | 0.39512503 | ZT18 | Absent_Koike2012 | Absent_Koike2012 |
| ENSMUSG00000079495 | Gm11128 | 6 | 85758018 | 85770948 | - | 0.00010789 | 0.00360311 | 8.90253199 | 2.48179774 | 8.10882773 | 14.5554811 | 14.5554811 | ZT18 | Absent_Koike2012 | Absent_Koike2012 |
| ENSMUSG00000025153 | Fasn | 11 | 120667254 | 120685492 | - | 0.00011168 | 0.00369556 | 77.7398103 | 53.1337066 | 157.405558 | 157.852263 | 157.852263 | ZT18 | Absent_Koike2012 | Present_Koike2012 |
| ENSMUSG00000068221 | Pdxp | 15 | 78744349 | 78749947 | + | 0.00011359 | 0.00372491 | 0.17881875 | 0.0467615 | 0.38141225 | 0.55511825 | 0.55511825 | ZT18 | Absent_Koike2012 | Absent_Koike2012 |
| ENSMUSG00000024778 | Fas | 19 | 34365149 | 34402260 | + | 0.000126 | 0.00401286 | 4.84580565 | 4.49979009 | 4.33726877 | 5.77483683 | 5.77483683 | ZT18 | Present_Koike2012 | Absent_Koike2012 |
| ENSMUSG00000072949 | Acot1 | 12 | 85350448 | 85358621 | + | 0.00012746 | 0.00403041 | 8.86684495 | 4.05934313 | 4.47102174 | 9.70876048 | 9.70876048 | ZT18 | Present_Koike2012 | Absent_Koike2012 |
| ENSMUSG00000040283 | Btnl9 | 11 | 48979087 | 49000661 | - | 0.00012795 | 0.00403527 | 1.40351817 | 1.19300555 | 1.24501792 | 1.63986322 | 1.63986322 | ZT18 | Absent_Koike2012 | Absent_Koike2012 |
| ENSMUSG00000022951 | Rcan1 | 16 | 92392198 | 92466391 | - | 0.00013231 | 0.00414646 | 8.65517846 | 3.64152559 | 9.2424595 | 9.55061312 | 9.55061312 | ZT18 | Absent_Koike2012 | Present_Koike2012 |
| ENSMUSG00000035473 | Galm | 17 | 80526811 | 80584441 | + | 0.00013329 | 0.00416335 | 31.1245497 | 28.4492772 | 32.9080127 | 35.2672347 | 35.2672347 | ZT18 | Absent_Koike2012 | Absent_Koike2012 |
| ENSMUSG00000002633 | Shh | 5 | 28783355 | 28793641 | - | 0.00013357 | 0.00416525 | 0.04170725 | 0.01477425 | 0.01620325 | 0.08614325 | 0.08614325 | ZT18 | Absent_Koike2012 | Absent_Koike2012 |
| ENSMUSG00000006818 | Sod2 | 17 | 13200705 | 13210985 | + | 0.00013543 | 0.00419759 | 37.0050857 | 31.5758441 | 36.3304537 | 39.6353296 | 39.6353296 | ZT18 | Absent_Koike2012 | Absent_Koike2012 |
| ENSMUSG00000054455 | Vapb | 2 | 173563012 | 173609837 | + | 0.00014027 | 0.00429533 | 6.68617016 | 6.09713254 | 6.65534861 | 6.81091259 | 6.81091259 | ZT18 | Absent_Koike2012 | Absent_Koike2012 |
| ENSMUSG00000074210 | E130208F15I | 7 | 31106759 | 31108491 | + | 0.00014044 | 0.00429544 | 2.08000525 | 1.604638 | 2.02980525 | 2.2886405 | 2.2886405 | ZT18 | Absent_Koike2012 | Absent_Koike2012 |
| ENSMUSG00000024921 | Smarca2 | 19 | 26679605 | 26852811 | + | 0.00014229 | 0.00433658 | 7.3038244 | 7.48835184 | 10.3779293 | 10.403448 | 10.403448 | ZT18 | Present_Koike2012 | Present_Koike2012 |
| ENSMUSG00000021018 | Polr2h | 16 | 20717738 | 20722340 | + | 0.00014724 | 0.00441843 | 3.70056689 | 2.78576055 | 3.78077861 | 3.84613152 | 3.84613152 | ZT18 | Present_Koike2012 | Absent_Koike2012 |
| ENSMUSG00000048578 | Mlec | 5 | 115592990 | 115608188 | - | 0.00015097 | 0.00448262 | 18.938004 | 13.789572 | 18.1883332 | 21.2451655 | 21.2451655 | ZT18 | Absent_Koike2012 | Absent_Koike2012 |
| ENSMUSG00000034165 | Ccnd3 | 17 | 47642000 | 47736632 | + | 0.00015337 | 0.00451199 | 5.61482981 | 4.37923375 | 5.37396058 | 6.11931397 | 6.11931397 | ZT18 | Absent_Koike2012 | Absent_Koike2012 |
| ENSMUSG00000040613 | Apobec1 | 6 | 122527810 | 122552462 | - | 0.0001608 | 0.00468709 | 3.11111005 | 4.02700724 | 4.00742857 | 4.1366856 | 4.1366856 | ZT18 | Present_Koike2012 | Absent_Koike2012 |
| ENSMUSG00000041762 | Gpr155 | 2 | 73179563 | 73224629 | - | 0.00016437 | 0.00474776 | 5.50502944 | 4.39117936 | 5.67068404 | 6.0830825 | 6.0830825 | ZT18 | Absent_Koike2012 | Absent_Koike2012 |
| ENSMUSG00000025271 | Pfkfb1 | X | 147022772 | 147078421 | + | 0.00016675 | 0.00478918 | 4.67882716 | 2.86073935 | 4.79151343 | 6.37807486 | 6.37807486 | ZT18 | Present_Koike2012 | Present_Koike2012 |
| ENSMUSG00000018965 | Ywhah | 5 | 33361465 | 33370615 | + | 0.00016847 | 0.00483007 | 5.44792152 | 5.0183568 | 6.16508483 | 6.3898465 | 6.3898465 | ZT18 | Absent_Koike2012 | Absent_Koike2012 |
| ENSMUSG00000023942 | Slc29a1 | 17 | 45722149 | 45736555 | - | 0.00017387 | 0.00491594 | 20.9525878 | 15.4471541 | 18.7469896 | 24.4478224 | 24.4478224 | ZT18 | Absent_Koike2012 | Present_Koike2012 |
| ENSMUSG00000063428 | Ddo | 10 | 40349817 | 40401653 | + | 0.00017559 | 0.00494825 | 1.48907408 | 0.96851922 | 1.30286718 | 1.77993232 | 1.77993232 | ZT18 | Present_Koike2012 | Present_Koike2012 |
| ENSMUSG00000028032 | Papss1 | 3 | 131227732 | 131306634 | + | 0.00018341 | 0.00506768 | 5.00049825 | 4.322763 | 4.2160995 | 5.65083375 | 5.65083375 | ZT18 | Absent_Koike2012 | Absent_Koike2012 |
| ENSMUSG00000079003 | Samd1 | 8 | 86521571 | 86524285 | + | 0.00018142 | 0.00506768 | 2.60252975 | 1.71318 | 2.54522725 | 3.58953675 | 3.58953675 | ZT18 | Absent_Koike2012 | Absent_Koike2012 |
| ENSMUSG00000020534 | Shmt1 | 11 | 60601606 | 60625220 | - | 0.00018212 | 0.00506768 | 38.146867 | 32.1415709 | 35.3027821 | 44.8301361 | 44.8301361 | ZT18 | Absent_Koike2012 | Absent_Koike2012 |
| ENSMUSG00000034371 | Dak | 19 | 10662281 | 10679922 | - | 0.00018502 | 0.00510101 | 47.6593879 | 35.2956816 | 68.0062447 | 78.7400957 | 78.7400957 | ZT18 | Absent_Koike2012 | Present_Koike2012 |
| ENSMUSG00000031451 | Gas6 | 8 | 13465374 | 13494535 | - | 0.0002094 | 0.0056032 | 10.1324378 | 7.1142315 | 8.76974425 | 10.4854998 | 10.4854998 | ZT18 | Present_Koike2012 | Absent_Koike2012 |
| ENSMUSG00000062075 | Lmnb2 | 10 | 80363948 | 80380990 | - | 0.00021228 | 0.00565651 | 1.63054078 | 1.0362885 | 1.51396325 | 1.70931931 | 1.70931931 | ZT18 | Absent_Koike2012 | Absent_Koike2012 |
| ENSMUSG00000047061 | Gm9817 | 13 | 45174061 | 45176591 | + | 0.00021353 | 0.00566609 | 0.49586725 | 0.22610425 | 0.16588775 | 1.006996 | 1.006996 | ZT18 | Absent_Koike2012 | Absent_Koike2012 |
| ENSMUSG00000029370 | Rassf6 | 5 | 91032102 | 91069513 | - | 0.00022125 | 0.00579834 | 3.86513934 | 3.16253383 | 4.29639939 | 6.28679172 | 6.28679172 | ZT18 | Absent_Koike2012 | Present_Koike2012 |
| ENSMUSG00000060093 | Hist1h4a | 13 | 23852561 | 23853099 | - | 0.00023567 | 0.00604067 | 3.39331375 | 1.53533725 | 3.31195 | 3.57586075 | 3.57586075 | ZT18 | Absent_Koike2012 | Absent_Koike2012 |
| ENSMUSG00000023393 | Slc17a9 | 2 | 180459968 | 180476985 | + | 0.00023777 | 0.00606547 | 0.72661539 | 0.59375324 | 0.79413628 | 1.2087782 | 1.2087782 | ZT18 | Absent_Koike2012 | Absent_Koike2012 |
| ENSMUSG00000024169 | Ift140 | 17 | 25153036 | 25236440 | + | 0.00024307 | 0.00614841 | 0.90728742 | 0.75265953 | 0.85234566 | 1.08506231 | 1.08506231 | ZT18 | Absent_Koike2012 | Absent_Koike2012 |
| ENSMUSG00000025870 | Arl10 | 13 | 54676376 | 54682489 | + | 0.00024864 | 0.00621735 | 1.12337404 | 0.86948469 | 1.05870905 | 1.31707737 | 1.31707737 | ZT18 | Absent_Koike2012 | Absent_Koike2012 |
| ENSMUSG00000036585 | Fgf1 | 18 | 38998327 | 39089058 | - | 0.00024872 | 0.00621735 | 14.4537269 | 10.2829685 | 13.3800343 | 20.0600553 | 20.0600553 | ZT18 | Present_Koike2012 | Present_Koike2012 |
| ENSMUSG00000032724 | Abtb2 | 2 | 103406467 | 103558580 | + | 0.00025441 | 0.00631615 | 3.75432111 | 2.47623883 | 2.02433713 | 4.01497763 | 4.01497763 | ZT18 | Present_Koike2012 | Present_Koike2012 |
| ENSMUSG00000030533 | Unc45a | 7 | 87470178 | 87485891 | - | 0.00025886 | 0.00634021 | 2.28858376 | 2.06360736 | 2.04883754 | 2.51299606 | 2.51299606 | ZT18 | Absent_Koike2012 | Absent_Koike2012 |
| ENSMUSG00000003348 | Mob3a | 10 | 80148001 | 80164565 | - | 0.000262 | 0.00638864 | 1.17183725 | 1.047131 | 1.39006425 | 1.56646475 | 1.56646475 | ZT18 | Absent_Koike2012 | Absent_Koike2012 |
| ENSMUSG00000026434 | Nucks1 | 1 | 133807035 | 133832898 | + | 0.00026315 | 0.0064019 | 22.0564028 | 19.0154218 | 24.4122715 | 24.6878243 | 24.6878243 | ZT18 | Present_Koike2012 | Absent_Koike2012 |
| ENSMUSG00000001424 | Snd1 | 6 | 28425139 | 28885162 | + | 0.00026372 | 0.00640343 | 14.5043713 | 15.4652181 | 18.720333 | 19.1685527 | 19.1685527 | ZT18 | Absent_Koike2012 | Present_Koike2012 |
| ENSMUSG00000071650 | Ganab | 19 | 8972580 | 8991153 | + | 0.00026457 | 0.00641798 | 42.1257444 | 40.7032282 | 42.8534474 | 45.1879664 | 45.1879664 | ZT18 | Absent_Koike2012 | Absent_Koike2012 |
| ENSMUSG00000028307 | Aldob | 4 | 49548867 | 49562418 | - | 0.00026568 | 0.00643268 | 1095.51093 | 874.769278 | 1309.15539 | 1423.19539 | 1423.19539 | ZT18 | Present_Koike2012 | Present_Koike2012 |
| ENSMUSG00000026617 | Bpnt1 | 1 | 187156028 | 187181656 | + | 0.00027064 | 0.00652183 | 25.6219402 | 21.4818318 | 23.2111635 | 25.7703337 | 25.7703337 | ZT18 | Absent_Koike2012 | Present_Koike2012 |
| ENSMUSG00000035780 | Ugt2a3 | 5 | 87753997 | 87766220 | - | 0.00029354 | 0.00688697 | 121.935614 | 126.619538 | 149.609741 | 150.373799 | 150.373799 | ZT18 | Present_Koike2012 | Present_Koike2012 |
| ENSMUSG00000064264 | Zfp428 | 7 | 25292025 | 25300701 | + | 0.00029364 | 0.00688697 | 0.427949 | 0.2429879 | 0.33137196 | 0.6557183 | 0.6557183 | ZT18 | Absent_Koike2012 | Absent_Koike2012 |
| ENSMUSG00000031532 | Tmem66 | 8 | 35217617 | 35233899 | + | 0.00029489 | 0.00689712 | 48.971304 | 42.1289863 | 46.7766265 | 49.1231518 | 49.1231518 | ZT18 | Absent_Koike2012 | Absent_Koike2012 |
| ENSMUSG00000020917 | Acly | 11 | 100337667 | 100389314 | - | 0.00029446 | 0.00689712 | 72.3982667 | 51.8460701 | 83.5075213 | 112.016899 | 112.016899 | ZT18 | Present_Koike2012 | Present_Koike2012 |
| ENSMUSG00000046062 | Ppp1r15b | 1 | 135027720 | 135036377 | + | 0.00029485 | 0.00689712 | 17.2459678 | 14.4507561 | 18.3824108 | 19.437529 | 19.437529 | ZT18 | Absent_Koike2012 | Present_Koike2012 |
| ENSMUSG00000040842 | D4Ertd22e | 4 | 140668916 | 140695690 | - | 0.00030943 | 0.00715817 | 15.4036314 | 14.3191226 | 13.7374224 | 16.0398558 | 16.0398558 | ZT18 | Present_Koike2012 | Absent_Koike2012 |
| ENSMUSG00000067071 | Hes6 | 1 | 93308060 | 93310615 | - | 0.00031302 | 0.00719922 | 32.8284867 | 19.902434 | 38.5036492 | 39.5508314 | 39.5508314 | ZT18 | Absent_Koike2012 | Absent_Koike2012 |
| ENSMUSG00000035049 | Rrp12 | 19 | 41937342 | 41970643 | - | 0.00031306 | 0.00719922 | 2.618781 | 1.472633 | 2.437126 | 2.7286325 | 2.7286325 | ZT18 | Absent_Koike2012 | Absent_Koike2012 |
| ENSMUSG00000025823 | Pdia4 | 6 | 47746141 | 47763511 | - | 0.0003227 | 0.00734513 | 112.391012 | 90.3042009 | 76.8888584 | 115.022433 | 115.022433 | ZT18 | Present_Koike2012 | Absent_Koike2012 |
| ENSMUSG00000021591 | Glrx | 13 | 75977316 | 75987599 | + | 0.00033486 | 0.00753411 | 14.7229557 | 11.2675423 | 16.8837454 | 17.1822323 | 17.1822323 | ZT18 | Absent_Koike2012 | Absent_Koike2012 |
| ENSMUSG00000032437 | Stt3b | 9 | 115151725 | 115219539 | - | 0.00033735 | 0.00753834 | 23.6948178 | 18.9002033 | 20.3806338 | 23.8765978 | 23.8765978 | ZT18 | Absent_Koike2012 | Present_Koike2012 |
| ENSMUSG00000022131 | Gpr180 | 14 | 118536349 | 118563454 | + | 0.00033742 | 0.00753834 | 5.753118 | 5.10502275 | 6.435106 | 6.4607195 | 6.4607195 | ZT18 | Absent_Koike2012 | Absent_Koike2012 |
| ENSMUSG00000038403 | Hfe2 | 3 | 96329108 | 96333139 | + | 0.00034738 | 0.00770022 | 15.0312167 | 16.1746441 | 11.5094289 | 16.8820968 | 16.8820968 | ZT18 | Present_Koike2012 | Absent_Koike2012 |
| ENSMUSG00000004748 | Mtfp1 | 11 | 3991483 | 3995448 | - | 0.00034852 | 0.00771859 | 9.80223671 | 7.36667194 | 9.88310166 | 9.88737632 | 9.88737632 | ZT18 | Absent_Koike2012 | Absent_Koike2012 |
| ENSMUSG00000018189 | Uchl5 | 1 | 145624408 | 145654596 | + | 0.00034919 | 0.00772682 | 4.4703355 | 4.6786145 | 5.56989675 | 5.7546005 | 5.7546005 | ZT18 | Absent_Koike2012 | Absent_Koike2012 |
| ENSMUSG00000078889 | Gm14288 | 2 | 175788489 | 175800225 | - | 0.0003642 | 0.0079621 | 1.99181466 | 1.39209393 | 2.0106683 | 2.55030883 | 2.55030883 | ZT18 | Absent_Koike2012 | Absent_Koike2012 |
| ENSMUSG00000078884 | Gm14436 | 2 | 176131382 | 176141650 | + | 0.00036848 | 0.00801437 | 0.84935292 | 0.59603199 | 0.84671253 | 1.20893892 | 1.20893892 | ZT18 | Absent_Koike2012 | Absent_Koike2012 |
| ENSMUSG00000019738 | Polr2i | 7 | 31016967 | 31018409 | + | 0.00039787 | 0.00847948 | 7.10219424 | 5.23125425 | 6.58925121 | 7.10764974 | 7.10764974 | ZT18 | Present_Koike2012 | Absent_Koike2012 |
| ENSMUSG00000057068 | Fam47e | 5 | 92984095 | 93020305 | + | 0.00039825 | 0.00848052 | 4.81931895 | 2.79691277 | 6.35060153 | 6.84125104 | 6.84125104 | ZT18 | Present_Koike2012 | Absent_Koike2012 |

| | | | | | | | | | | | | | | | |
|---|---|---|---|---|---|---|---|---|---|---|---|---|---|---|---|
| ENSMUSG00000069008 | Gm5537 | 3 | 57906916 | 57908169 | - | 0.00040824 | 0.00860689 | 8.4133275 | 6.4112825 | 7.90606 | 9.16272 | 9.16272 | ZT18 | Absent_Koike2012 | Absent_Koike2012 |
| ENSMUSG00000021867 | Gm9746 | 14 | 27022841 | 27026557 | - | 0.00041352 | 0.00867783 | 16.567492 | 14.2151362 | 12.3003503 | 17.6979346 | 17.6979346 | ZT18 | Absent_Koike2012 | Absent_Koike2012 |
| ENSMUSG00000022234 | Cct5 | 15 | 31520555 | 31531602 | - | 0.00041365 | 0.00867783 | 34.9145717 | 29.7303847 | 33.3353569 | 35.2268817 | 35.2268817 | ZT18 | Absent_Koike2012 | Absent_Koike2012 |
| ENSMUSG00000043445 | Pgp | 17 | 24607418 | 24608536 | + | 0.00041905 | 0.00874055 | 7.85163425 | 5.75501375 | 7.91526025 | 8.079318 | 8.079318 | ZT18 | Absent_Koike2012 | Absent_Koike2012 |
| ENSMUSG00000028496 | Mllt3 | 4 | 87415829 | 87679268 | - | 0.00042249 | 0.0087886 | 0.23930932 | 0.18923452 | 0.25854295 | 0.29073633 | 0.29073633 | ZT18 | Absent_Koike2012 | Present_Koike2012 |
| ENSMUSG00000029326 | Enoph1 | 5 | 100469013 | 100497781 | + | 0.00042457 | 0.00880089 | 1.45490976 | 1.1771994 | 1.60085101 | 1.64508609 | 1.64508609 | ZT18 | Absent_Koike2012 | Absent_Koike2012 |
| ENSMUSG00000007659 | Bcl2l1 | 2 | 152593052 | 152657464 | - | 0.00042522 | 0.00880089 | 2.64883882 | 1.53472479 | 2.86327884 | 3.0222478 | 3.0222478 | ZT18 | Absent_Koike2012 | Absent_Koike2012 |
| ENSMUSG00000030104 | Edem1 | 6 | 108778635 | 108809350 | + | 0.00044564 | 0.00910852 | 38.2707103 | 31.4959758 | 32.7808363 | 38.6157333 | 38.6157333 | ZT18 | Absent_Koike2012 | Present_Koike2012 |
| ENSMUSG00000035198 | Tubg1 | 11 | 100981252 | 100987733 | + | 0.00044742 | 0.009127 | 7.19563088 | 5.65519784 | 6.59714292 | 8.1598447 | 8.1598447 | ZT18 | Absent_Koike2012 | Absent_Koike2012 |
| ENSMUSG00000028914 | Casp9 | 4 | 141349527 | 141371891 | + | 0.00045332 | 0.00919899 | 2.08858293 | 2.86911075 | 2.62681264 | 2.97726801 | 2.97726801 | ZT18 | Absent_Koike2012 | Absent_Koike2012 |
| ENSMUSG00000030284 | Creld1 | 6 | 113433291 | 113443332 | + | 0.00045613 | 0.00924143 | 4.88412193 | 3.69832195 | 3.86097085 | 5.0516098 | 5.0516098 | ZT18 | Absent_Koike2012 | Absent_Koike2012 |
| ENSMUSG00000035834 | Polr3g | 13 | 81812842 | 81850012 | - | 0.0004631 | 0.00935179 | 2.47867553 | 1.87808973 | 2.42801818 | 2.9748298 | 2.9748298 | ZT18 | Absent_Koike2012 | Absent_Koike2012 |
| ENSMUSG00000024978 | Gpam | 19 | 55142142 | 55201824 | - | 0.00046392 | 0.00935453 | 37.1333223 | 38.0612333 | 50.0085901 | 60.7085012 | 60.7085012 | ZT18 | Absent_Koike2012 | Absent_Koike2012 |
| ENSMUSG00000078864 | Gm14322 | 2 | 177493993 | 177505177 | + | 0.00047783 | 0.00955915 | 0.40372832 | 0.29683044 | 0.59029681 | 0.61962714 | 0.61962714 | ZT18 | Absent_Koike2012 | Absent_Koike2012 |
| ENSMUSG00000070705 | Eid2b | 7 | 29062725 | 29065148 | + | 0.00048374 | 0.00965227 | 0.75743425 | 0.64259625 | 0.55499875 | 0.823806 | 0.823806 | ZT18 | Absent_Koike2012 | Absent_Koike2012 |
| ENSMUSG00000007594 | Hapln4 | 8 | 72607428 | 72614761 | + | 0.00048566 | 0.00967024 | 1.35335325 | 0.783779 | 0.82371175 | 1.49851775 | 1.49851775 | ZT18 | Absent_Koike2012 | Absent_Koike2012 |
| ENSMUSG00000026603 | Smyd2 | 1 | 191704371 | 191746242 | - | 0.00049091 | 0.00974794 | 4.297904 | 3.336685 | 4.12658532 | 5.01924679 | 5.01924679 | ZT18 | Absent_Koike2012 | Present_Koike2012 |
| ENSMUSG00000020889 | Nr1d1 | 11 | 98629246 | 98636647 | - | 7.42E-13 | 1.89E-08 | 0.31459875 | 8.10482102 | 1.85358045 | 0.21143885 | 8.10482102 | ZT6 | Present_Koike2012 | Present_Koike2012 |
| ENSMUSG00000027346 | Gpcpd1 | 2 | 132354818 | 132413465 | - | 7.25E-09 | 1.32E-05 | 3.80441417 | 13.1689666 | 12.2796978 | 3.91292989 | 13.1689666 | ZT6 | Present_Koike2012 | Present_Koike2012 |
| ENSMUSG00000052407 | 4930473A06 | 4 | 83171449 | 83510635 | + | 2.55E-08 | 3.09E-05 | 0.62182551 | 0.88505699 | 0.65085142 | 0.4468301 | 0.88505699 | ZT6 | Absent_Koike2012 | Absent_Koike2012 |
| ENSMUSG00000032898 | Fbxo21 | 5 | 118426779 | 118460200 | + | 4.09E-08 | 3.85E-05 | 10.4907955 | 29.149968 | 13.968951 | 8.4008906 | 29.149968 | ZT6 | Present_Koike2012 | Absent_Koike2012 |
| ENSMUSG00000024785 | Rcl1 | 19 | 29175865 | 29218333 | + | 4.50E-08 | 3.95E-05 | 26.238682 | 50.4596271 | 36.1044761 | 24.8448145 | 50.4596271 | ZT6 | Present_Koike2012 | Absent_Koike2012 |
| ENSMUSG00000083250 | Gm5403 | X | 124593067 | 124595861 | - | 5.56E-08 | 4.42E-05 | 0.08803775 | 0.280903 | 0.13668325 | 0.071919 | 0.280903 | ZT6 | Absent_Koike2012 | Absent_Koike2012 |
| ENSMUSG00000009647 | Ccdc109a | 10 | 58909732 | 59079440 | - | 1.38E-07 | 7.96E-05 | 2.15848398 | 2.23147194 | 1.33179981 | 1.34238912 | 2.23147194 | ZT6 | Absent_Koike2012 | Absent_Koike2012 |
| ENSMUSG00000016487 | Ppfibp1 | 6 | 146837009 | 146980547 | + | 1.69E-07 | 8.01E-05 | 2.73619039 | 3.95237329 | 2.36404991 | 2.00132118 | 3.95237329 | ZT6 | Present_Koike2012 | Absent_Koike2012 |
| ENSMUSG00000028991 | Mtor | 4 | 147822720 | 147931792 | + | 1.93E-07 | 8.15E-05 | 4.66629343 | 6.36706961 | 5.97315045 | 3.94202462 | 6.36706961 | ZT6 | Present_Koike2012 | Absent_Koike2012 |
| ENSMUSG00000003623 | Crot | 5 | 8966033 | 8997324 | - | 2.34E-07 | 8.49E-05 | 96.0875177 | 108.178887 | 44.5297627 | 41.4489719 | 108.178887 | ZT6 | Present_Koike2012 | Present_Koike2012 |
| ENSMUSG00000071655 | Ubxn1 | 19 | 8946049 | 8950153 | + | 2.56E-07 | 9.05E-05 | 45.179721 | 48.4896021 | 30.0848793 | 27.7239569 | 48.4896021 | ZT6 | Present_Koike2012 | Present_Koike2012 |
| ENSMUSG00000050628 | Fam100b | 11 | 116295408 | 116300391 | + | 2.67E-07 | 9.30E-05 | 4.55137208 | 7.46994292 | 5.76884158 | 4.44231643 | 7.46994292 | ZT6 | Absent_Koike2012 | Absent_Koike2012 |
| ENSMUSG00000035864 | Syt1 | 10 | 107934706 | 108448038 | - | 2.72E-07 | 9.35E-05 | 0.63772935 | 1.15270205 | 1.03763024 | 0.33138373 | 1.15270205 | ZT6 | Present_Koike2012 | Absent_Koike2012 |
| ENSMUSG00000041143 | Tmco4 | 4 | 138528803 | 138615086 | + | 3.32E-07 | 0.0001057 | 2.29473666 | 3.57613099 | 3.37824579 | 2.22666081 | 3.57613099 | ZT6 | Absent_Koike2012 | Absent_Koike2012 |
| ENSMUSG00000025795 | Rassf3 | 10 | 120847406 | 120913306 | - | 3.45E-07 | 0.00010809 | 12.8950188 | 19.257481 | 15.6463575 | 11.3046963 | 19.257481 | ZT6 | Present_Koike2012 | Absent_Koike2012 |
| ENSMUSG00000022297 | Fzd6 | 15 | 38837879 | 38869732 | + | 5.45E-07 | 0.00013341 | 0.455198 | 1.2922155 | 0.85212125 | 0.6313165 | 1.2922155 | ZT6 | Absent_Koike2012 | Absent_Koike2012 |
| ENSMUSG00000028630 | Dyrk2 | 10 | 118296405 | 118305959 | - | 5.25E-07 | 0.00013341 | 6.8438265 | 9.8654995 | 5.99258775 | 6.35288475 | 9.8654995 | ZT6 | Absent_Koike2012 | Absent_Koike2012 |
| ENSMUSG00000020034 | Tcp11l2 | 10 | 84039371 | 84077104 | + | 7.96E-07 | 0.00016197 | 2.07203893 | 6.59104129 | 3.70890238 | 2.06502448 | 6.59104129 | ZT6 | Present_Koike2012 | Absent_Koike2012 |
| ENSMUSG00000037712 | Fermt2 | 14 | 46078467 | 46149793 | - | 8.28E-07 | 0.0001633 | 22.7948452 | 27.6024503 | 16.1757357 | 21.6985302 | 27.6024503 | ZT6 | Present_Koike2012 | Absent_Koike2012 |
| ENSMUSG00000014791 | Elmo3 | 8 | 107829501 | 107835300 | + | 9.00E-07 | 0.00017091 | 4.7122257 | 5.45652974 | 5.37345982 | 4.15140669 | 5.45652974 | ZT6 | Absent_Koike2012 | Absent_Koike2012 |
| ENSMUSG00000043962 | Thrap3 | 4 | 125841326 | 125880004 | - | 1.00E-06 | 0.00018515 | 5.52989885 | 6.59600423 | 5.23499888 | 3.67240845 | 6.59600423 | ZT6 | Absent_Koike2012 | Absent_Koike2012 |
| ENSMUSG00000027167 | Elp4 | 2 | 105537477 | 105744721 | - | 1.03E-06 | 0.00018718 | 2.36381082 | 2.50124811 | 1.51053324 | 1.54679946 | 2.50124811 | ZT6 | Absent_Koike2012 | Absent_Koike2012 |
| ENSMUSG00000030161 | Gabarapl1 | 6 | 129483218 | 129492349 | + | 1.07E-06 | 0.00019068 | 88.71451 | 143.608036 | 90.910154 | 61.7643853 | 143.608036 | ZT6 | Present_Koike2012 | Present_Koike2012 |
| ENSMUSG00000020472 | Zkscan17 | 11 | 59299022 | 59340253 | - | 1.16E-06 | 0.00019624 | 1.89311002 | 2.94801363 | 2.1666903 | 1.45903552 | 2.94801363 | ZT6 | Absent_Koike2012 | Absent_Koike2012 |
| ENSMUSG00000090394 | 4930523C07I | 1 | 161974515 | 162008559 | + | 1.14E-06 | 0.00019624 | 0.61757226 | 1.19352665 | 1.1525134 | 0.77600309 | 1.19352665 | ZT6 | Absent_Koike2012 | Absent_Koike2012 |
| ENSMUSG00000028354 | Fmn2 | 1 | 176431956 | 176752848 | + | 1.28E-06 | 0.0002057 | 0.049346 | 0.1067885 | 0.0296515 | 0.02214175 | 0.1067885 | ZT6 | Absent_Koike2012 | Absent_Koike2012 |
| ENSMUSG00000042308 | Setd1a | 7 | 134920184 | 134943636 | + | 1.39E-06 | 0.00021598 | 28.6381581 | 31.8922039 | 11.8640997 | 15.2850735 | 31.8922039 | ZT6 | Absent_Koike2012 | Absent_Koike2012 |
| ENSMUSG00000020919 | Stat5b | 11 | 100642045 | 100712038 | - | 1.78E-06 | 0.00025747 | 9.18785257 | 10.3222853 | 6.55911698 | 7.71094499 | 10.3222853 | ZT6 | Present_Koike2012 | Absent_Koike2012 |
| ENSMUSG00000022270 | Fam134b | 15 | 25773019 | 25903442 | + | 1.80E-06 | 0.00025798 | 5.52902028 | 12.4839625 | 5.62257842 | 5.90759259 | 12.4839625 | ZT6 | Absent_Koike2012 | Absent_Koike2012 |
| ENSMUSG00000037003 | Tenc1 | 15 | 101930909 | 101946832 | + | 2.81E-06 | 0.00035563 | 3.02347606 | 6.19059371 | 4.50032076 | 3.15804413 | 6.19059371 | ZT6 | Present_Koike2012 | Present_Koike2012 |
| ENSMUSG00000024908 | Ppp6r3 | 19 | 3454933 | 3575749 | - | 2.96E-06 | 0.00036935 | 13.2485238 | 16.026809 | 14.6464063 | 12.792851 | 16.026809 | ZT6 | Absent_Koike2012 | Absent_Koike2012 |
| ENSMUSG00000027997 | Casp6 | 3 | 129604343 | 129617021 | + | 3.31E-06 | 0.0003969 | 4.07002745 | 7.5635958 | 4.20043895 | 2.58140205 | 7.5635958 | ZT6 | Present_Koike2012 | Absent_Koike2012 |
| ENSMUSG00000026888 | Grb14 | 2 | 64750533 | 64863044 | - | 3.49E-06 | 0.00041108 | 4.20351954 | 5.16511674 | 3.72833013 | 3.23909503 | 5.16511674 | ZT6 | Absent_Koike2012 | Absent_Koike2012 |
| ENSMUSG00000029512 | Ulk1 | 5 | 111213507 | 111239116 | - | 3.91E-06 | 0.00044378 | 3.218233 | 6.33425275 | 5.9346225 | 2.789423 | 6.33425275 | ZT6 | Present_Koike2012 | Absent_Koike2012 |
| ENSMUSG00000037622 | Wdtc1 | 4 | 132848374 | 132909396 | - | 4.02E-06 | 0.00045035 | 10.0927524 | 12.2040348 | 8.76970145 | 7.40216389 | 12.2040348 | ZT6 | Present_Koike2012 | Absent_Koike2012 |
| ENSMUSG00000020614 | Fam20a | 11 | 109531063 | 109583593 | - | 4.19E-06 | 0.00045771 | 4.9366071 | 5.4813862 | 3.16127158 | 3.97688401 | 5.4813862 | ZT6 | Present_Koike2012 | Absent_Koike2012 |
| ENSMUSG00000040272 | Accs | 2 | 93673624 | 93690100 | - | 4.23E-06 | 0.00045951 | 0.15191237 | 0.25303026 | 0.16873746 | 0.12776339 | 0.25303026 | ZT6 | Absent_Koike2012 | Absent_Koike2012 |
| ENSMUSG00000056820 | Tsnax | 8 | 127536897 | 127558092 | + | 4.30E-06 | 0.00046155 | 8.62005075 | 9.80033875 | 9.72575 | 7.89386675 | 9.80033875 | ZT6 | Absent_Koike2012 | Absent_Koike2012 |
| ENSMUSG00000032066 | Bco2 | 9 | 50341192 | 50363286 | - | 4.39E-06 | 0.0004674 | 6.64078123 | 7.54880951 | 4.74051063 | 5.93218384 | 7.54880951 | ZT6 | Present_Koike2012 | Absent_Koike2012 |
| ENSMUSG00000026662 | Sephs1 | 2 | 4802610 | 4831603 | + | 4.52E-06 | 0.00047761 | 3.30488262 | 4.01865114 | 3.98435037 | 3.15835975 | 4.01865114 | ZT6 | Present_Koike2012 | Absent_Koike2012 |
| ENSMUSG00000027340 | Slc23a2 | 2 | 131878232 | 131970844 | - | 4.64E-06 | 0.0004817 | 1.51398429 | 2.5441846 | 1.69944863 | 1.3813989 | 2.5441846 | ZT6 | Absent_Koike2012 | Absent_Koike2012 |
| ENSMUSG00000029592 | Usp30 | 5 | 114550342 | 114573515 | + | 4.78E-06 | 0.00048619 | 3.172655 | 3.6896205 | 3.49416925 | 2.77332525 | 3.6896205 | ZT6 | Absent_Koike2012 | Absent_Koike2012 |
| ENSMUSG00000058655 | Eif4b | 15 | 101904204 | 101927603 | + | 4.76E-06 | 0.00048619 | 42.1285059 | 50.1701801 | 48.7301082 | 33.1053283 | 50.1701801 | ZT6 | Absent_Koike2012 | Present_Koike2012 |
| ENSMUSG00000025188 | Hps1 | 19 | 42829595 | 42854468 | - | 4.80E-06 | 0.00048619 | 0.93171853 | 0.93865079 | 0.65361731 | 0.5727862 | 0.93865079 | ZT6 | Absent_Koike2012 | Absent_Koike2012 |
| ENSMUSG00000027809 | Etfdh | 3 | 79407710 | 79432785 | - | 4.81E-06 | 0.00048619 | 28.1396892 | 28.5748608 | 23.4100495 | 20.5554718 | 28.5748608 | ZT6 | Present_Koike2012 | Absent_Koike2012 |
| ENSMUSG00000020381 | 3010026O09 | 11 | 49987946 | 50013617 | + | 4.88E-06 | 0.00048968 | 0.7044541 | 1.41120813 | 0.3949457 | 0.50014423 | 1.41120813 | ZT6 | Present_Koike2012 | Absent_Koike2012 |
| ENSMUSG00000040123 | Zmym5 | 14 | 57409425 | 57430553 | - | 4.96E-06 | 0.00048968 | 2.47534775 | 3.73098019 | 3.13493103 | 2.22612022 | 3.73098019 | ZT6 | Present_Koike2012 | Absent_Koike2012 |
| ENSMUSG00000032715 | Trib3 | 2 | 152163158 | 152169768 | - | 4.96E-06 | 0.00048968 | 0.45201348 | 3.23278689 | 0.94597269 | 0.36568921 | 3.23278689 | ZT6 | Absent_Koike2012 | Absent_Koike2012 |
| ENSMUSG00000037204 | 9430023L20I | 15 | 101114732 | 101121365 | + | 5.03E-06 | 0.00049371 | 12.349844 | 12.7173782 | 8.56007976 | 11.2258954 | 12.7173782 | ZT6 | Present_Koike2012 | Absent_Koike2012 |
| ENSMUSG00000028029 | Aimp1 | 3 | 132323462 | 132346843 | - | 5.30E-06 | 0.00050565 | 11.6184608 | 13.331545 | 12.998437 | 10.0297725 | 13.331545 | ZT6 | Absent_Koike2012 | Absent_Koike2012 |
| ENSMUSG00000042446 | Zmym4 | 4 | 126539069 | 126645379 | - | 5.30E-06 | 0.00050565 | 1.16290592 | 1.56572611 | 1.33813585 | 1.0925996 | 1.56572611 | ZT6 | Absent_Koike2012 | Absent_Koike2012 |
| ENSMUSG00000045827 | Serpinb9 | 13 | 33095119 | 33109826 | + | 5.31E-06 | 0.00050565 | 2.4910814 | 3.03941125 | 1.98547452 | 1.43962861 | 3.03941125 | ZT6 | Present_Koike2012 | Absent_Koike2012 |

| | | | | | | | | | | | | | | | |
|---|---|---|---|---|---|---|---|---|---|---|---|---|---|---|---|
| ENSMUSG00000040964 | Arhgef10l | 4 | 140070400 | 140221927 | - | 5.33E-06 | 0.0005064 | 2.69672385 | 4.26859211 | 2.52894774 | 2.27539592 | 4.26859211 | ZT6 | Absent_Koike2012 | Present_Koike2012 |
| ENSMUSG00000034154 | Ino80 | 2 | 119198778 | 119303423 | - | 5.59E-06 | 0.00052649 | 1.86134341 | 2.39613908 | 2.30652718 | 1.71487478 | 2.39613908 | ZT6 | Absent_Koike2012 | Absent_Koike2012 |
| ENSMUSG00000029763 | Exoc4 | 6 | 33199085 | 33923979 | + | 5.64E-06 | 0.00052982 | 1.63532932 | 1.77822561 | 1.62222136 | 1.17454989 | 1.77822561 | ZT6 | Absent_Koike2012 | Absent_Koike2012 |
| ENSMUSG00000035212 | Leprot | 4 | 101320323 | 101331969 | + | 5.92E-06 | 0.00055166 | 4.4207645 | 5.84243658 | 5.24102529 | 3.91840989 | 5.84243658 | ZT6 | Absent_Koike2012 | Absent_Koike2012 |
| ENSMUSG00000025764 | Phf17 | 3 | 41359656 | 41420784 | + | 5.96E-06 | 0.00055396 | 5.37158729 | 9.11410039 | 7.28072574 | 4.51884359 | 9.11410039 | ZT6 | Present_Koike2012 | Absent_Koike2012 |
| ENSMUSG00000030551 | Nr2f2 | 7 | 77496832 | 77511621 | - | 5.99E-06 | 0.00055462 | 3.2033442 | 3.57947907 | 2.56603036 | 2.33083594 | 3.57947907 | ZT6 | Present_Koike2012 | Absent_Koike2012 |
| ENSMUSG00000042328 | Hps4 | 5 | 112772103 | 112807434 | + | 6.69E-06 | 0.00060569 | 1.4421706 | 1.99140996 | 1.28315667 | 1.47524546 | 1.99140996 | ZT6 | Absent_Koike2012 | Absent_Koike2012 |
| ENSMUSG00000067889 | Spnb3 | 19 | 4711208 | 4752353 | + | 7.09E-06 | 0.00063268 | 9.28372852 | 13.9303375 | 10.2684668 | 9.12118817 | 13.9303375 | ZT6 | Absent_Koike2012 | Absent_Koike2012 |
| ENSMUSG00000028194 | Ddah1 | 3 | 145421639 | 145557241 | + | 7.25E-06 | 0.00063372 | 22.7594449 | 26.0506391 | 21.7155557 | 17.7874364 | 26.0506391 | ZT6 | Present_Koike2012 | Present_Koike2012 |
| ENSMUSG00000026278 | Bok | 1 | 95582271 | 95592933 | + | 8.12E-06 | 0.00068033 | 3.14546453 | 3.5323358 | 1.73532413 | 2.50897136 | 3.5323358 | ZT6 | Absent_Koike2012 | Absent_Koike2012 |
| ENSMUSG00000027810 | Eif2a | 3 | 58329743 | 58361423 | + | 8.50E-06 | 0.0007025 | 3.54465231 | 4.48171429 | 4.36135044 | 3.17094882 | 4.48171429 | ZT6 | Absent_Koike2012 | Absent_Koike2012 |
| ENSMUSG00000022350 | E430025E21I | 15 | 59163553 | 59205722 | - | 8.91E-06 | 0.00071292 | 6.1576308 | 7.02126626 | 6.57829118 | 5.49920957 | 7.02126626 | ZT6 | Present_Koike2012 | Absent_Koike2012 |
| ENSMUSG00000089917 | Uckl1 | 2 | 181303854 | 181319597 | - | 1.04E-05 | 0.00079946 | 1.91986529 | 2.36013585 | 2.27207446 | 1.73775517 | 2.36013585 | ZT6 | Absent_Koike2012 | Absent_Koike2012 |
| ENSMUSG00000045733 | Sprn | 7 | 147336527 | 147340776 | - | 1.05E-05 | 0.0007996 | 0.18838875 | 0.38913325 | 0.36553875 | 0.2039605 | 0.38913325 | ZT6 | Absent_Koike2012 | Absent_Koike2012 |
| ENSMUSG00000042688 | Mapk6 | 9 | 75216869 | 75257812 | - | 1.06E-05 | 0.00080251 | 2.70751572 | 3.54608646 | 3.39971128 | 2.74353408 | 3.54608646 | ZT6 | Present_Koike2012 | Present_Koike2012 |
| ENSMUSG00000042500 | Eif2c4 | 4 | 126166785 | 126210716 | - | 1.08E-05 | 0.00081049 | 0.39236702 | 0.61083174 | 0.43932806 | 0.38254528 | 0.61083174 | ZT6 | Absent_Koike2012 | Absent_Koike2012 |
| ENSMUSG00000042032 | Mat2b | 11 | 40492816 | 40508705 | - | 1.09E-05 | 0.00081541 | 3.73996776 | 4.95054532 | 4.81884353 | 3.90605782 | 4.95054532 | ZT6 | Present_Koike2012 | Absent_Koike2012 |
| ENSMUSG00000022770 | Dlg1 | 16 | 31663529 | 31873436 | + | 1.09E-05 | 0.00081541 | 3.04053685 | 3.83253032 | 3.34218577 | 2.66839667 | 3.83253032 | ZT6 | Absent_Koike2012 | Absent_Koike2012 |
| ENSMUSG00000040725 | Hnrnpul1 | 7 | 26506517 | 26539749 | - | 1.10E-05 | 0.00081583 | 11.6480981 | 13.2830229 | 12.3558135 | 10.6450874 | 13.2830229 | ZT6 | Absent_Koike2012 | Absent_Koike2012 |
| ENSMUSG00000009646 | Pla2g12b | 10 | 58866408 | 58884724 | + | 1.11E-05 | 0.00082536 | 25.290051 | 25.873865 | 16.99086 | 19.5852625 | 25.873865 | ZT6 | Present_Koike2012 | Absent_Koike2012 |
| ENSMUSG00000048581 | E130311K13I | 3 | 63718606 | 63733394 | - | 1.21E-05 | 0.00087818 | 1.50367965 | 1.79853799 | 0.95246203 | 1.03999354 | 1.79853799 | ZT6 | Present_Koike2012 | Absent_Koike2012 |
| ENSMUSG00000015165 | Hnrnpl | 7 | 29593560 | 29607285 | + | 1.23E-05 | 0.00087967 | 7.08722294 | 8.98316605 | 7.62835167 | 5.47948186 | 8.98316605 | ZT6 | Present_Koike2012 | Absent_Koike2012 |
| ENSMUSG00000028423 | Nfx1 | 4 | 40917939 | 40973026 | + | 1.26E-05 | 0.00089369 | 3.70631009 | 4.59076043 | 4.38729603 | 3.0558758 | 4.59076043 | ZT6 | Present_Koike2012 | Absent_Koike2012 |
| ENSMUSG00000038371 | Sbf2 | 7 | 117451527 | 117758436 | - | 1.27E-05 | 0.0009016 | 2.34696676 | 2.81487769 | 2.29991044 | 1.92227463 | 2.81487769 | ZT6 | Present_Koike2012 | Absent_Koike2012 |
| ENSMUSG00000020653 | Klf11 | 12 | 25336236 | 25347639 | + | 1.33E-05 | 0.00092324 | 0.90757541 | 3.09500528 | 1.82715199 | 1.14768123 | 3.09500528 | ZT6 | Absent_Koike2012 | Absent_Koike2012 |
| ENSMUSG00000025017 | Pik3ap1 | 19 | 41346863 | 41459586 | - | 1.33E-05 | 0.00092324 | 8.01806253 | 12.0053799 | 11.4802632 | 6.42279882 | 12.0053799 | ZT6 | Present_Koike2012 | Absent_Koike2012 |
| ENSMUSG00000002031 | Ift46 | 9 | 44575991 | 44600797 | + | 1.37E-05 | 0.00093173 | 1.91418017 | 2.40045076 | 2.00480868 | 1.69264941 | 2.40045076 | ZT6 | Present_Koike2012 | Absent_Koike2012 |
| ENSMUSG00000059586 | Nsmce2 | 15 | 59205753 | 59433238 | + | 1.36E-05 | 0.00093173 | 1.53375077 | 1.79169759 | 1.72773058 | 1.1739683 | 1.79169759 | ZT6 | Absent_Koike2012 | Absent_Koike2012 |
| ENSMUSG00000062604 | Srpk2 | 5 | 23009174 | 23182266 | - | 1.40E-05 | 0.00094281 | 1.80665503 | 2.40975488 | 2.24468436 | 1.85862095 | 2.40975488 | ZT6 | Absent_Koike2012 | Absent_Koike2012 |
| ENSMUSG00000020661 | Dnmt3a | 12 | 3806007 | 3914443 | + | 1.45E-05 | 0.00096032 | 0.82120031 | 1.18181165 | 0.94593133 | 0.77140009 | 1.18181165 | ZT6 | Present_Koike2012 | Absent_Koike2012 |
| ENSMUSG00000042851 | Zc3h6 | 2 | 128793138 | 128844299 | + | 1.50E-05 | 0.0009807 | 0.1845434 | 0.46702039 | 0.26003541 | 0.18497867 | 0.46702039 | ZT6 | Absent_Koike2012 | Absent_Koike2012 |
| ENSMUSG00000034412 | Tbc1d10a | 11 | 4086792 | 4115508 | + | 1.51E-05 | 0.00098148 | 1.96317941 | 3.01239766 | 2.23206533 | 1.57190962 | 3.01239766 | ZT6 | Absent_Koike2012 | Absent_Koike2012 |
| ENSMUSG00000010755 | Cars | 7 | 150743135 | 150785995 | - | 1.52E-05 | 0.00098448 | 3.9955189 | 5.74496045 | 4.82191571 | 3.28586732 | 5.74496045 | ZT6 | Present_Koike2012 | Absent_Koike2012 |
| ENSMUSG00000035248 | Zcchc6 | 13 | 59873240 | 59924508 | - | 1.58E-05 | 0.0010187 | 12.1132051 | 15.398217 | 13.8408445 | 10.9784559 | 15.398217 | ZT6 | Present_Koike2012 | Absent_Koike2012 |
| ENSMUSG00000050931 | Sgms2 | 3 | 131021903 | 131047858 | - | 1.64E-05 | 0.00105034 | 5.59522941 | 8.9090695 | 6.07056545 | 5.44002178 | 8.9090695 | ZT6 | Present_Koike2012 | Absent_Koike2012 |
| ENSMUSG00000041961 | Znrf3 | 11 | 5176327 | 5344850 | - | 1.70E-05 | 0.00106931 | 0.98143476 | 1.36301928 | 1.21997862 | 0.79675372 | 1.36301928 | ZT6 | Absent_Koike2012 | Absent_Koike2012 |
| ENSMUSG00000064138 | Fam172a | 13 | 77847951 | 78305499 | + | 1.70E-05 | 0.00106931 | 1.26870345 | 1.47924689 | 1.25711634 | 0.98550711 | 1.47924689 | ZT6 | Absent_Koike2012 | Absent_Koike2012 |
| ENSMUSG00000037921 | Ddx60 | 8 | 64406884 | 64516498 | + | 1.75E-05 | 0.00109087 | 0.73234433 | 1.4859193 | 0.95304957 | 0.64376165 | 1.4859193 | ZT6 | Absent_Koike2012 | Absent_Koike2012 |
| ENSMUSG00000018166 | Erbb3 | 10 | 128004579 | 128026708 | - | 1.83E-05 | 0.00112572 | 13.470424 | 28.4486618 | 16.2098575 | 18.0114143 | 28.4486618 | ZT6 | Present_Koike2012 | Present_Koike2012 |
| ENSMUSG00000093577 | RP24-201C14 | 3 | 96043316 | 96045742 | + | 1.83E-05 | 0.00112572 | 0.10727525 | 0.246422 | 0.125333 | 0.09316425 | 0.246422 | ZT6 | Absent_Koike2012 | Absent_Koike2012 |
| ENSMUSG00000020608 | Smc6 | 12 | 11272692 | 11326591 | + | 1.86E-05 | 0.00113764 | 3.7763125 | 4.711199 | 4.48114625 | 3.46952725 | 4.711199 | ZT6 | Absent_Koike2012 | Present_Koike2012 |
| ENSMUSG00000001151 | Pcnt | 10 | 75813199 | 75905657 | - | 2.01E-05 | 0.00119655 | 1.19886151 | 1.65686894 | 1.44717042 | 1.09836839 | 1.65686894 | ZT6 | Absent_Koike2012 | Absent_Koike2012 |
| ENSMUSG00000031534 | AI316807 | 8 | 23573087 | 23587351 | - | 2.04E-05 | 0.00121103 | 4.99397266 | 6.90875395 | 5.53734995 | 4.79240468 | 6.90875395 | ZT6 | Absent_Koike2012 | Absent_Koike2012 |
| ENSMUSG00000057054 | Inca1 | 11 | 70501863 | 70513657 | - | 2.13E-05 | 0.00125622 | 4.42448367 | 6.22587467 | 3.08415089 | 2.23528896 | 6.22587467 | ZT6 | Absent_Koike2012 | Absent_Koike2012 |
| ENSMUSG00000079045 | Gm17566 | 1 | 191994175 | 192041155 | + | 2.20E-05 | 0.00128653 | 2.4873035 | 2.87150024 | 1.87495512 | 1.6892704 | 2.87150024 | ZT6 | Absent_Koike2012 | Absent_Koike2012 |
| ENSMUSG00000019659 | Ccdc12 | 9 | 110559007 | 110614094 | + | 2.30E-05 | 0.00130891 | 11.5979978 | 11.7911843 | 9.21725275 | 8.66977825 | 11.7911843 | ZT6 | Absent_Koike2012 | Absent_Koike2012 |
| ENSMUSG00000029470 | P2rx4 | 5 | 123157593 | 123179747 | + | 2.35E-05 | 0.00132639 | 6.65150743 | 10.1280094 | 9.97856505 | 5.97336043 | 10.1280094 | ZT6 | Present_Koike2012 | Absent_Koike2012 |
| ENSMUSG00000031844 | Hsd17b2 | 8 | 120225846 | 120282929 | + | 2.35E-05 | 0.00132639 | 76.7515202 | 85.7644106 | 50.274822 | 50.5183063 | 85.7644106 | ZT6 | Present_Koike2012 | Absent_Koike2012 |
| ENSMUSG00000024325 | Ring1 | 17 | 34157737 | 34161625 | - | 2.43E-05 | 0.00135087 | 1.87112711 | 2.2721419 | 1.8750588 | 1.84984253 | 2.2721419 | ZT6 | Absent_Koike2012 | Absent_Koike2012 |
| ENSMUSG00000060477 | Irak2 | 6 | 113588461 | 113645003 | + | 2.52E-05 | 0.0013867 | 2.12146212 | 2.29878536 | 1.35560971 | 2.00567487 | 2.29878536 | ZT6 | Absent_Koike2012 | Absent_Koike2012 |
| ENSMUSG00000057342 | Sphk2 | 7 | 52964837 | 52973372 | - | 2.60E-05 | 0.00142527 | 5.05169061 | 9.80513513 | 7.54139341 | 6.38419972 | 9.80513513 | ZT6 | Absent_Koike2012 | Absent_Koike2012 |
| ENSMUSG00000039934 | Pion | 5 | 20692085 | 20797519 | + | 2.66E-05 | 0.00143856 | 6.609357 | 9.8420865 | 8.36040075 | 6.2053535 | 9.8420865 | ZT6 | Present_Koike2012 | Absent_Koike2012 |
| ENSMUSG00000032812 | Arap1 | 7 | 108496581 | 108561100 | + | 2.66E-05 | 0.00143856 | 4.64316429 | 5.37358994 | 3.83633285 | 4.40340703 | 5.37358994 | ZT6 | Absent_Koike2012 | Absent_Koike2012 |
| ENSMUSG00000004085 | B230120H23 | 2 | 72123694 | 72280667 | + | 2.67E-05 | 0.00143856 | 1.35288787 | 1.68187797 | 1.54767878 | 1.2183002 | 1.68187797 | ZT6 | Absent_Koike2012 | Absent_Koike2012 |
| ENSMUSG00000032560 | Dnajc13 | 9 | 104053927 | 104165260 | - | 2.74E-05 | 0.00146576 | 5.38165825 | 5.86051656 | 5.60258848 | 4.92170423 | 5.86051656 | ZT6 | Absent_Koike2012 | Absent_Koike2012 |
| ENSMUSG00000046169 | Adamts6 | 13 | 105077953 | 105284843 | + | 2.86E-05 | 0.00150551 | 0.07182635 | 0.250229 | 0.19020987 | 0.08802136 | 0.250229 | ZT6 | Absent_Koike2012 | Absent_Koike2012 |
| ENSMUSG00000001123 | Lgals9 | 11 | 78756476 | 78798448 | - | 2.86E-05 | 0.00150551 | 67.1356994 | 70.9654653 | 48.5379277 | 42.9222349 | 70.9654653 | ZT6 | Present_Koike2012 | Present_Koike2012 |
| ENSMUSG00000030788 | Rnf141 | 7 | 117943946 | 117987971 | - | 2.98E-05 | 0.00154335 | 8.47025605 | 8.69531284 | 6.56975454 | 6.77916448 | 8.69531284 | ZT6 | Absent_Koike2012 | Absent_Koike2012 |
| ENSMUSG00000034621 | Gpatch8 | 11 | 102337229 | 102417706 | - | 3.22E-05 | 0.00162856 | 1.44702395 | 1.8837652 | 1.73298143 | 1.50759111 | 1.8837652 | ZT6 | Absent_Koike2012 | Absent_Koike2012 |
| ENSMUSG00000004364 | Cul3 | 1 | 80261498 | 80337055 | - | 3.25E-05 | 0.00163775 | 9.05555701 | 10.8266184 | 10.3192486 | 8.35741402 | 10.8266184 | ZT6 | Absent_Koike2012 | Absent_Koike2012 |
| ENSMUSG00000026878 | Rab14 | 2 | 35035725 | 35056640 | - | 3.24E-05 | 0.00163775 | 11.5015255 | 12.3354287 | 10.3978071 | 10.8962562 | 12.3354287 | ZT6 | Absent_Koike2012 | Absent_Koike2012 |
| ENSMUSG00000030083 | Abtb1 | 6 | 88785909 | 88791929 | - | 3.28E-05 | 0.00164696 | 6.49760636 | 8.13681159 | 7.12403002 | 5.58368854 | 8.13681159 | ZT6 | Present_Koike2012 | Present_Koike2012 |
| ENSMUSG00000078429 | Ctdsp2 | 10 | 126415773 | 126437031 | + | 3.33E-05 | 0.00166944 | 4.58422682 | 7.81559672 | 6.13656741 | 4.69746272 | 7.81559672 | ZT6 | Absent_Koike2012 | Absent_Koike2012 |
| ENSMUSG00000056133 | Gm9992 | 17 | 7568061 | 7589654 | - | 3.39E-05 | 0.00169023 | 0.60676775 | 1.3026045 | 0.72062775 | 0.2275835 | 1.3026045 | ZT6 | Absent_Koike2012 | Absent_Koike2012 |
| ENSMUSG00000023079 | Gtf2ird1 | 5 | 134833539 | 134932581 | - | 3.40E-05 | 0.00169254 | 3.0972811 | 3.43351408 | 2.33509222 | 3.13358071 | 3.43351408 | ZT6 | Absent_Koike2012 | Absent_Koike2012 |
| ENSMUSG00000002996 | Hbp1 | 12 | 32611119 | 32635400 | - | 3.51E-05 | 0.00173138 | 8.20140785 | 10.3191993 | 8.78364185 | 7.46199945 | 10.3191993 | ZT6 | Absent_Koike2012 | Absent_Koike2012 |
| ENSMUSG00000049115 | Agtr1a | 13 | 30428310 | 30474736 | + | 3.88E-05 | 0.00185522 | 14.2810002 | 19.2919583 | 11.4255719 | 10.9453411 | 19.2919583 | ZT6 | Present_Koike2012 | Absent_Koike2012 |
| ENSMUSG00000039166 | Akap7 | 10 | 24888896 | 25027676 | - | 3.93E-05 | 0.00187128 | 0.92209689 | 1.10718915 | 0.98312332 | 0.7324441 | 1.10718915 | ZT6 | Absent_Koike2012 | Absent_Koike2012 |
| ENSMUSG00000022999 | Lmbr1l | 15 | 98734352 | 98748662 | - | 3.96E-05 | 0.00187915 | 0.96929869 | 1.55523396 | 1.55470915 | 1.21432239 | 1.55523396 | ZT6 | Absent_Koike2012 | Absent_Koike2012 |

| | | | | | | | | | | | | | | | | |
|---|---|---|---|---|---|---|---|---|---|---|---|---|---|---|---|---|
| ENSMUSG00000027465 | Tbc1d20 | | 2 | 152119564 | 152139732 | + | 4.00E-05 | 0.00188973 | 4.7383929 | 5.50564752 | 5.41081992 | 4.00368979 | 5.50564752 | ZT6 | Absent_Koike2012 | Absent_Koike2012 |
| ENSMUSG00000055817 | Mta3 | | 17 | 84105503 | 84220856 | + | 4.11E-05 | 0.00192747 | 1.13834606 | 1.40177689 | 1.20480519 | 0.95212539 | 1.40177689 | ZT6 | Absent_Koike2012 | Absent_Koike2012 |
| ENSMUSG00000074063 | Osgin1 | | 8 | 121958024 | 121970156 | + | 4.18E-05 | 0.00194776 | 11.5327381 | 14.7546956 | 5.45401929 | 7.95860692 | 14.7546956 | ZT6 | Present_Koike2012 | Present_Koike2012 |
| ENSMUSG00000024070 | Prkd3 | | 17 | 79348745 | 79420156 | - | 4.20E-05 | 0.00194776 | 14.4226359 | 18.768448 | 11.3744748 | 8.92857471 | 18.768448 | ZT6 | Present_Koike2012 | Absent_Koike2012 |
| ENSMUSG00000086593 | Gm16548 | | 1 | 166078722 | 166082348 | - | 4.20E-05 | 0.00194776 | 2.45198081 | 4.06202896 | 2.84974967 | 2.28185178 | 4.06202896 | ZT6 | Absent_Koike2012 | Absent_Koike2012 |
| ENSMUSG00000081763 | Gm9435 | X | | 14532224 | 14533100 | - | 4.39E-05 | 0.00199897 | 0.00598975 | 0.0652255 | 0.00752425 | 0.00809675 | 0.0652255 | ZT6 | Absent_Koike2012 | Absent_Koike2012 |
| ENSMUSG00000000346 | Dazap2 | | 15 | 100446069 | 100451184 | + | 4.36E-05 | 0.00199897 | 31.9613493 | 35.552046 | 29.72866 | 26.3565785 | 35.552046 | ZT6 | Present_Koike2012 | Present_Koike2012 |
| ENSMUSG00000024975 | Pdcd4 | | 19 | 53966721 | 54004351 | + | 4.54E-05 | 0.00205007 | 24.9567315 | 39.1924178 | 22.07525 | 26.5243142 | 39.1924178 | ZT6 | Present_Koike2012 | Absent_Koike2012 |
| ENSMUSG00000022620 | Arsa | | 15 | 89302907 | 89307856 | - | 4.59E-05 | 0.00206788 | 4.75415466 | 5.74283021 | 5.45034873 | 4.02000335 | 5.74283021 | ZT6 | Present_Koike2012 | Present_Koike2012 |
| ENSMUSG00000046792 | Zfp787 | | 7 | 6083091 | 6107573 | - | 4.69E-05 | 0.00208769 | 2.89378317 | 2.97528408 | 2.82815666 | 2.51238026 | 2.97528408 | ZT6 | Present_Koike2012 | Absent_Koike2012 |
| ENSMUSG00000021144 | Mta1 | | 12 | 114336489 | 114375417 | + | 4.76E-05 | 0.00211341 | 3.06140467 | 3.65501681 | 3.08090479 | 2.60721059 | 3.65501681 | ZT6 | Absent_Koike2012 | Absent_Koike2012 |
| ENSMUSG00000002546 | Golga2 | | 2 | 32142904 | 32163441 | + | 4.80E-05 | 0.00212382 | 4.67563226 | 5.16444956 | 4.90549758 | 4.15294958 | 5.16444956 | ZT6 | Absent_Koike2012 | Absent_Koike2012 |
| ENSMUSG00000073678 | Pgap1 | | 1 | 54529844 | 54532810 | - | 4.94E-05 | 0.00214822 | 1.015013 | 2.024808 | 1.8984785 | 1.13982275 | 2.024808 | ZT6 | Present_Koike2012 | Absent_Koike2012 |
| ENSMUSG00000051375 | Pcdh1 | | 18 | 38345569 | 38371707 | - | 4.98E-05 | 0.0021599 | 4.54571425 | 5.61303668 | 3.59419289 | 3.3645214 | 5.61303668 | ZT6 | Absent_Koike2012 | Absent_Koike2012 |
| ENSMUSG00000022314 | Rad21 | | 15 | 51794151 | 51823306 | - | 5.02E-05 | 0.00216327 | 7.86997025 | 9.623015 | 8.8897075 | 7.130764 | 9.623015 | ZT6 | Absent_Koike2012 | Absent_Koike2012 |
| ENSMUSG00000004508 | Gab2 | | 7 | 104230261 | 104457450 | + | 5.11E-05 | 0.0021977 | 0.41998525 | 0.580502 | 0.56028875 | 0.4587485 | 0.580502 | ZT6 | Absent_Koike2012 | Absent_Koike2012 |
| ENSMUSG00000015890 | Amdhd1 | | 10 | 92986083 | 93002778 | - | 5.15E-05 | 0.00220948 | 68.5106925 | 82.125496 | 48.310545 | 35.8958868 | 82.125496 | ZT6 | Absent_Koike2012 | Present_Koike2012 |
| ENSMUSG00000040234 | Tm7sf3 | | 6 | 146550874 | 146591346 | - | 5.32E-05 | 0.00226662 | 15.0998858 | 17.0401007 | 15.8803368 | 14.216842 | 17.0401007 | ZT6 | Absent_Koike2012 | Absent_Koike2012 |
| ENSMUSG00000063785 | Utp14a | X | | 45610039 | 45635630 | + | 5.35E-05 | 0.00226942 | 2.14006676 | 2.78716468 | 2.02741495 | 1.55881371 | 2.78716468 | ZT6 | Absent_Koike2012 | Absent_Koike2012 |
| ENSMUSG00000050390 | C77080 | | 4 | 128896822 | 128938648 | - | 5.38E-05 | 0.00227674 | 3.4273516 | 5.28860799 | 4.7305511 | 2.90745318 | 5.28860799 | ZT6 | Present_Koike2012 | Absent_Koike2012 |
| ENSMUSG00000022180 | Slc7a8 | | 14 | 55341053 | 55400723 | - | 5.42E-05 | 0.00227719 | 0.72579425 | 0.73766275 | 0.59896975 | 0.45952325 | 0.73766275 | ZT6 | Absent_Koike2012 | Absent_Koike2012 |
| ENSMUSG00000027048 | Abcb11 | | 2 | 69076339 | 69180671 | - | 5.42E-05 | 0.00227719 | 140.276098 | 149.718641 | 100.31689 | 81.7613044 | 149.718641 | ZT6 | Present_Koike2012 | Present_Koike2012 |
| ENSMUSG00000040511 | Pvr | | 7 | 20488927 | 20506509 | - | 5.65E-05 | 0.00233392 | 0.67233743 | 1.06775494 | 0.9388782 | 0.59470379 | 1.06775494 | ZT6 | Absent_Koike2012 | Present_Koike2012 |
| ENSMUSG00000032485 | Scap | | 9 | 110235792 | 110287450 | + | 5.70E-05 | 0.00235078 | 19.2279366 | 22.6946118 | 17.8170802 | 14.4233846 | 22.6946118 | ZT6 | Absent_Koike2012 | Absent_Koike2012 |
| ENSMUSG00000036822 | Topors | | 4 | 40206634 | 40216883 | - | 5.73E-05 | 0.00235429 | 6.84118375 | 8.0247595 | 7.89196575 | 6.1082005 | 8.0247595 | ZT6 | Absent_Koike2012 | Absent_Koike2012 |
| ENSMUSG00000032743 | D430042O09 | | 7 | 132851390 | 133018311 | + | 5.78E-05 | 0.00236138 | 2.13472592 | 2.34246094 | 1.58559929 | 1.66033523 | 2.34246094 | ZT6 | Absent_Koike2012 | Absent_Koike2012 |
| ENSMUSG00000032612 | Usp4 | | 9 | 108250162 | 108294860 | + | 5.76E-05 | 0.00236138 | 13.9750867 | 15.4661549 | 14.7069161 | 13.0740206 | 15.4661549 | ZT6 | Absent_Koike2012 | Absent_Koike2012 |
| ENSMUSG00000021357 | Exoc2 | | 13 | 30905788 | 31065916 | - | 6.02E-05 | 0.00243461 | 2.60241831 | 3.01949535 | 2.98056186 | 2.45435 | 3.01949535 | ZT6 | Absent_Koike2012 | Absent_Koike2012 |
| ENSMUSG00000040195 | Tmem194 | | 10 | 127103957 | 127138105 | + | 6.12E-05 | 0.00246004 | 0.52412593 | 0.77457524 | 0.62765007 | 0.48939435 | 0.77457524 | ZT6 | Absent_Koike2012 | Absent_Koike2012 |
| ENSMUSG00000034101 | Ctnnd1 | | 2 | 84440228 | 84490922 | - | 6.12E-05 | 0.00246004 | 10.0921901 | 11.0788109 | 8.89018624 | 8.95339402 | 11.0788109 | ZT6 | Absent_Koike2012 | Absent_Koike2012 |
| ENSMUSG00000024331 | Dsc2 | | 18 | 20189134 | 20218055 | - | 6.22E-05 | 0.00248712 | 4.06370461 | 7.02280372 | 6.96580521 | 4.99049647 | 7.02280372 | ZT6 | Absent_Koike2012 | Absent_Koike2012 |
| ENSMUSG00000041135 | Ripk2 | | 4 | 16049880 | 16090794 | - | 6.31E-05 | 0.00250194 | 1.19337967 | 2.01092109 | 1.99217139 | 1.27928929 | 2.01092109 | ZT6 | Present_Koike2012 | Absent_Koike2012 |
| ENSMUSG00000046711 | Hmga1 | | 17 | 27693565 | 27700619 | + | 6.55E-05 | 0.00256769 | 1.71445248 | 2.53050021 | 2.49593041 | 1.48505042 | 2.53050021 | ZT6 | Absent_Koike2012 | Absent_Koike2012 |
| ENSMUSG00000069114 | Zbtb10 | | 3 | 9250602 | 9285333 | + | 6.62E-05 | 0.00258684 | 0.5555393 | 0.82453574 | 0.66861962 | 0.48356691 | 0.82453574 | ZT6 | Absent_Koike2012 | Absent_Koike2012 |
| ENSMUSG00000026768 | Itga8 | | 2 | 12028259 | 12223547 | - | 6.72E-05 | 0.00260122 | 1.2412482 | 1.62323875 | 1.16723102 | 0.7406778 | 1.62323875 | ZT6 | Present_Koike2012 | Absent_Koike2012 |
| ENSMUSG00000026970 | Rbms1 | | 2 | 60588250 | 60801261 | - | 6.71E-05 | 0.00260122 | 3.3010534 | 5.00271355 | 4.35517921 | 2.86313307 | 5.00271355 | ZT6 | Present_Koike2012 | Absent_Koike2012 |
| ENSMUSG00000067049 | Unc93a | | 17 | 13301483 | 13324657 | - | 6.73E-05 | 0.00260157 | 0.18263586 | 0.39829409 | 0.20135715 | 0.0502442 | 0.39829409 | ZT6 | Absent_Koike2012 | Absent_Koike2012 |
| ENSMUSG00000074782 | 4833422C13 | | 13 | 91841270 | 91881477 | - | 6.97E-05 | 0.00266689 | 0.13358227 | 0.20631443 | 0.16211 | 0.11832074 | 0.20631443 | ZT6 | Absent_Koike2012 | Absent_Koike2012 |
| ENSMUSG00000035671 | Zswim4 | | 8 | 86734577 | 86760954 | - | 7.12E-05 | 0.00271416 | 1.242748 | 1.5692265 | 1.25478875 | 1.082794 | 1.5692265 | ZT6 | Absent_Koike2012 | Absent_Koike2012 |
| ENSMUSG00000039145 | Camk1d | | 2 | 5214503 | 5635561 | - | 7.48E-05 | 0.00280186 | 8.52737292 | 9.27876458 | 4.0659322 | 4.80211807 | 9.27876458 | ZT6 | Present_Koike2012 | Present_Koike2012 |
| ENSMUSG00000060679 | Mrps9 | | 1 | 42908070 | 42962527 | + | 7.50E-05 | 0.00280626 | 4.1686803 | 5.06467367 | 4.79469765 | 3.77370038 | 5.06467367 | ZT6 | Absent_Koike2012 | Present_Koike2012 |
| ENSMUSG00000024059 | Clip4 | | 17 | 72119042 | 72207691 | + | 7.54E-05 | 0.00281147 | 0.08257475 | 0.12661106 | 0.08436785 | 0.06595136 | 0.12661106 | ZT6 | Absent_Koike2012 | Absent_Koike2012 |
| ENSMUSG00000032410 | Xrn1 | | 9 | 95855179 | 95953446 | + | 7.73E-05 | 0.00284639 | 1.6786695 | 2.23692776 | 1.93673164 | 1.60363259 | 2.23692776 | ZT6 | Absent_Koike2012 | Absent_Koike2012 |
| ENSMUSG00000030805 | Stx4a | | 7 | 134967808 | 134992533 | + | 7.76E-05 | 0.00285016 | 2.30085772 | 2.6797272 | 1.75313785 | 1.99553607 | 2.6797272 | ZT6 | Absent_Koike2012 | Absent_Koike2012 |
| ENSMUSG00000021198 | Unc79 | | 12 | 104187069 | 104422208 | + | 8.05E-05 | 0.00292791 | 0 | 0.00557311 | 0.00099995 | 0 | 0.00557311 | ZT6 | Absent_Koike2012 | Absent_Koike2012 |
| ENSMUSG00000037686 | Aspg | | 12 | 113344894 | 113365784 | + | 8.07E-05 | 0.0029296 | 18.4391308 | 24.3893466 | 19.5079572 | 14.5124113 | 24.3893466 | ZT6 | Present_Koike2012 | Absent_Koike2012 |
| ENSMUSG00000020605 | Hs1bp3 | | 12 | 8320239 | 8350630 | + | 8.10E-05 | 0.00293109 | 2.29467175 | 2.66177175 | 1.93105225 | 2.0617105 | 2.66177175 | ZT6 | Absent_Koike2012 | Absent_Koike2012 |
| ENSMUSG00000092241 | Gm20522 | | 17 | 35259683 | 35263385 | - | 8.25E-05 | 0.00297926 | 0.2385445 | 0.571856 | 0.317149 | 0.302294 | 0.571856 | ZT6 | Absent_Koike2012 | Absent_Koike2012 |
| ENSMUSG00000042323 | Pbrm1 | | 14 | 31832324 | 31934778 | + | 8.32E-05 | 0.00299777 | 5.07000523 | 6.34063854 | 5.73056381 | 4.69530379 | 6.34063854 | ZT6 | Absent_Koike2012 | Absent_Koike2012 |
| ENSMUSG00000037656 | Slc20a2 | | 8 | 23587172 | 23680079 | + | 8.39E-05 | 0.0030199 | 11.3905203 | 14.1081568 | 12.1730708 | 10.490243 | 14.1081568 | ZT6 | Absent_Koike2012 | Absent_Koike2012 |
| ENSMUSG00000025925 | Terf1 | | 1 | 15795739 | 15833551 | + | 8.41E-05 | 0.00302215 | 1.99456846 | 2.82131096 | 2.40331076 | 1.83165986 | 2.82131096 | ZT6 | Absent_Koike2012 | Absent_Koike2012 |
| ENSMUSG00000021546 | Hnrnpk | | 13 | 58492503 | 58504704 | - | 8.43E-05 | 0.00302505 | 22.8667888 | 26.8091439 | 23.116891 | 17.0412884 | 26.8091439 | ZT6 | Absent_Koike2012 | Present_Koike2012 |
| ENSMUSG00000057143 | Trim12c | | 7 | 111487268 | 111501876 | - | 8.46E-05 | 0.00302813 | 1.77218687 | 2.43081002 | 1.59416244 | 1.49987963 | 2.43081002 | ZT6 | Absent_Koike2012 | Absent_Koike2012 |
| ENSMUSG00000052609 | Plekhg3 | | 12 | 77634547 | 77680026 | + | 8.73E-05 | 0.00311266 | 3.16116294 | 4.35581247 | 4.30309065 | 3.36069345 | 4.35581247 | ZT6 | Absent_Koike2012 | Absent_Koike2012 |
| ENSMUSG00000084957 | Bbip1 | | 19 | 54003757 | 54019117 | - | 8.97E-05 | 0.00316614 | 7.47693904 | 11.3006829 | 6.66019463 | 8.24903673 | 11.3006829 | ZT6 | Absent_Koike2012 | Absent_Koike2012 |
| ENSMUSG00000034480 | Diap2 | X | | 126284278 | 127000370 | + | 9.04E-05 | 0.00317281 | 1.03896019 | 1.35968743 | 1.33548457 | 0.85410461 | 1.35968743 | ZT6 | Present_Koike2012 | Present_Koike2012 |
| ENSMUSG00000021843 | Ktn1 | | 14 | 48203644 | 48356238 | + | 9.16E-05 | 0.0032101 | 2.92999362 | 3.97192148 | 3.35122196 | 2.52343788 | 3.97192148 | ZT6 | Present_Koike2012 | Absent_Koike2012 |
| ENSMUSG00000030256 | Bhlhe41 | | 6 | 145806763 | 145814078 | - | 9.54E-05 | 0.00330874 | 0.04632486 | 0.32587953 | 0.20097773 | 0.06458014 | 0.32587953 | ZT6 | Absent_Koike2012 | Absent_Koike2012 |
| ENSMUSG00000023045 | Soat2 | | 15 | 101980957 | 101993905 | + | 9.79E-05 | 0.00335182 | 8.39676484 | 11.219204 | 11.1315803 | 7.27732549 | 11.219204 | ZT6 | Absent_Koike2012 | Absent_Koike2012 |
| ENSMUSG00000029328 | Hnrpdl | | 5 | 100462596 | 100468683 | - | 9.88E-05 | 0.00337204 | 3.56787811 | 5.36374925 | 4.94767589 | 3.38617143 | 5.36374925 | ZT6 | Absent_Koike2012 | Absent_Koike2012 |
| ENSMUSG00000024014 | Pim1 | | 17 | 29627757 | 29632930 | + | 0.00010115 | 0.00343665 | 2.54924362 | 6.30093895 | 3.08535041 | 3.18436723 | 6.30093895 | ZT6 | Absent_Koike2012 | Absent_Koike2012 |
| ENSMUSG00000023055 | Calcoco1 | | 15 | 102537210 | 102552609 | - | 0.00010335 | 0.00349266 | 2.50990284 | 3.44648275 | 2.68899394 | 2.10667561 | 3.44648275 | ZT6 | Absent_Koike2012 | Absent_Koike2012 |
| ENSMUSG00000025384 | 2310003H01 | | 11 | 120230369 | 120240078 | - | 0.00010502 | 0.00353957 | 0.78458376 | 1.00280841 | 0.92043279 | 0.60609013 | 1.00280841 | ZT6 | Absent_Koike2012 | Absent_Koike2012 |
| ENSMUSG00000086024 | Gm16639 | | 15 | 54784503 | 54796197 | + | 0.00010583 | 0.00355752 | 0.30884825 | 0.64347775 | 0.393519 | 0.19570875 | 0.64347775 | ZT6 | Absent_Koike2012 | Absent_Koike2012 |
| ENSMUSG00000041236 | Vps41 | | 13 | 18809161 | 18958678 | + | 0.00011012 | 0.00366792 | 13.1007085 | 15.8779598 | 14.8125275 | 11.7426793 | 15.8779598 | ZT6 | Present_Koike2012 | Absent_Koike2012 |
| ENSMUSG00000054469 | Lclat1 | | 17 | 73457325 | 73592708 | + | 0.00011043 | 0.00367341 | 5.57810378 | 6.82543167 | 6.66695617 | 5.0385843 | 6.82543167 | ZT6 | Present_Koike2012 | Absent_Koike2012 |
| ENSMUSG00000004626 | Stxbp2 | | 8 | 3630955 | 3643644 | + | 0.00011139 | 0.00369556 | 3.22328063 | 3.67407617 | 3.41464501 | 2.78135532 | 3.67407617 | ZT6 | Absent_Koike2012 | Absent_Koike2012 |
| ENSMUSG00000026142 | Rhbdd1 | | 1 | 82313027 | 82441941 | + | 0.00011162 | 0.00369556 | 4.00271393 | 4.84230267 | 4.43453525 | 3.4602682 | 4.84230267 | ZT6 | Absent_Koike2012 | Absent_Koike2012 |
| ENSMUSG00000041560 | Gltscr2 | | 7 | 16523187 | 16531457 | - | 0.00011327 | 0.00372373 | 24.7944185 | 29.4441678 | 28.994474 | 20.3191865 | 29.4441678 | ZT6 | Absent_Koike2012 | Absent_Koike2012 |

| | | | | | | | | | | | | | | | |
|---|---|---|---|---|---|---|---|---|---|---|---|---|---|---|---|
| ENSMUSG00000017009 | Sdc4 | 2 | 164249747 | 164269387 | - | 0.0001143 | 0.00373853 | 119.65413 | 159.861982 | 146.783059 | 112.28157 | 159.861982 | ZT6 | Absent_Koike2012 | Absent_Koike2012 |
| ENSMUSG00000050310 | Rictor | 15 | 6658381 | 6750398 | + | 0.00011493 | 0.00374465 | 2.62675451 | 4.15916038 | 3.34411725 | 2.4697568 | 4.15916038 | ZT6 | Absent_Koike2012 | Absent_Koike2012 |
| ENSMUSG00000054519 | Zfp867 | 11 | 59274699 | 59285976 | - | 0.00011774 | 0.00383131 | 0.77294748 | 1.60124676 | 1.24249294 | 0.95917432 | 1.60124676 | ZT6 | Absent_Koike2012 | Absent_Koike2012 |
| ENSMUSG00000069495 | Epc2 | 2 | 49307006 | 49407468 | + | 0.0001199 | 0.00388178 | 2.15695361 | 3.06372853 | 2.57803427 | 2.07103236 | 3.06372853 | ZT6 | Absent_Koike2012 | Absent_Koike2012 |
| ENSMUSG00000026977 | March7 | 2 | 60047944 | 60088733 | + | 0.00012274 | 0.00394365 | 1.32523316 | 2.30263869 | 1.89541063 | 1.40215381 | 2.30263869 | ZT6 | Absent_Koike2012 | Absent_Koike2012 |
| ENSMUSG00000024427 | Spry4 | 18 | 38745922 | 38760922 | - | 0.00012397 | 0.00396601 | 0.98115025 | 1.73701475 | 1.159378 | 0.72887275 | 1.73701475 | ZT6 | Absent_Koike2012 | Absent_Koike2012 |
| ENSMUSG00000024773 | Atg2a | 19 | 6241668 | 6262335 | + | 0.00012484 | 0.00398088 | 4.63365956 | 6.23403039 | 5.68790691 | 4.113119 | 6.23403039 | ZT6 | Absent_Koike2012 | Absent_Koike2012 |
| ENSMUSG00000033182 | Kbtbd12 | 6 | 88497716 | 88577439 | - | 0.00012701 | 0.00402483 | 0.11269903 | 0.26100405 | 0.15493131 | 0.12101025 | 0.26100405 | ZT6 | Absent_Koike2012 | Absent_Koike2012 |
| ENSMUSG00000081788 | Gm5898 | 7 | 70687365 | 70689435 | + | 0.00012693 | 0.00402483 | 1.91518714 | 2.149403 | 1.96352375 | 1.24028675 | 2.149403 | ZT6 | Absent_Koike2012 | Absent_Koike2012 |
| ENSMUSG00000031842 | Pde4c | 8 | 73247619 | 73275085 | + | 0.00012692 | 0.00402483 | 1.11177545 | 1.50223377 | 1.17038948 | 0.84035399 | 1.50223377 | ZT6 | Absent_Koike2012 | Absent_Koike2012 |
| ENSMUSG00000021990 | Spata13 | 14 | 61252838 | 61383393 | + | 0.0001275 | 0.00403041 | 1.87165237 | 3.10810566 | 2.43059692 | 1.82087155 | 3.10810566 | ZT6 | Absent_Koike2012 | Absent_Koike2012 |
| ENSMUSG00000028134 | Ptbp2 | 3 | 119421660 | 119486306 | - | 0.00012797 | 0.00403527 | 1.93943175 | 2.365124 | 2.292825 | 1.4395725 | 2.365124 | ZT6 | Present_Koike2012 | Absent_Koike2012 |
| ENSMUSG00000028329 | Xpa | 4 | 46168219 | 46209183 | - | 0.00013022 | 0.00409613 | 2.45829773 | 3.18537673 | 2.50324953 | 2.03231104 | 3.18537673 | ZT6 | Absent_Koike2012 | Absent_Koike2012 |
| ENSMUSG00000038975 | Rabggtb | 3 | 153570253 | 153575930 | - | 0.00013158 | 0.00413257 | 12.3725755 | 13.3677783 | 11.837718 | 8.48158649 | 13.3677783 | ZT6 | Absent_Koike2012 | Absent_Koike2012 |
| ENSMUSG00000047797 | Gjb1 | X | 98571717 | 98580968 | + | 0.00013269 | 0.00415325 | 243.585316 | 261.188053 | 221.295341 | 195.827079 | 261.188053 | ZT6 | Present_Koike2012 | Present_Koike2012 |
| ENSMUSG00000062901 | Klhl24 | 16 | 20097627 | 20127817 | + | 0.00013334 | 0.00416335 | 8.00910806 | 16.6710881 | 12.8048898 | 8.24172075 | 16.6710881 | ZT6 | Absent_Koike2012 | Absent_Koike2012 |
| ENSMUSG00000024544 | D18Ertd653e | 18 | 68092911 | 68415203 | + | 0.00013427 | 0.00417691 | 1.37033982 | 1.83485578 | 1.48743775 | 0.99063065 | 1.83485578 | ZT6 | Present_Koike2012 | Absent_Koike2012 |
| ENSMUSG00000086463 | Gm16898 | 2 | 169459146 | 169714003 | - | 0.00013461 | 0.00418245 | 1.54088311 | 1.94915601 | 1.58326253 | 1.44986823 | 1.94915601 | ZT6 | Absent_Koike2012 | Absent_Koike2012 |
| ENSMUSG00000026896 | Ifih1 | 2 | 62433855 | 62484312 | - | 0.00013491 | 0.00418661 | 3.68921921 | 4.96052494 | 4.9180006 | 3.50596765 | 4.96052494 | ZT6 | Absent_Koike2012 | Absent_Koike2012 |
| ENSMUSG00000021703 | Serinc5 | 13 | 93381093 | 93481899 | + | 0.00013609 | 0.0042078 | 8.912593 | 10.4321988 | 8.7160815 | 7.998943 | 10.4321988 | ZT6 | Present_Koike2012 | Absent_Koike2012 |
| ENSMUSG00000022244 | Amacr | 15 | 10911511 | 10926379 | + | 0.00013976 | 0.00429002 | 37.1590328 | 43.7419738 | 35.0662738 | 31.3544623 | 43.7419738 | ZT6 | Present_Koike2012 | Absent_Koike2012 |
| ENSMUSG00000006281 | Tep1 | 14 | 51443737 | 51490235 | - | 0.00014075 | 0.00429961 | 1.80776975 | 1.90674425 | 1.710533 | 1.451083 | 1.90674425 | ZT6 | Absent_Koike2012 | Absent_Koike2012 |
| ENSMUSG00000056493 | Foxk1 | 5 | 142877451 | 142937968 | + | 0.00014258 | 0.00433658 | 1.483947 | 1.87615775 | 1.42773875 | 1.19231875 | 1.87615775 | ZT6 | Absent_Koike2012 | Absent_Koike2012 |
| ENSMUSG00000000881 | Dlg3 | X | 97963061 | 98013749 | + | 0.00014298 | 0.00433658 | 1.86710944 | 2.43745222 | 1.90728424 | 2.10408755 | 2.43745222 | ZT6 | Absent_Koike2012 | Absent_Koike2012 |
| ENSMUSG00000034308 | Sdr42e1 | 8 | 120185299 | 120197589 | - | 0.00014377 | 0.00435542 | 23.8479017 | 27.5651949 | 15.773208 | 18.7462507 | 27.5651949 | ZT6 | Absent_Koike2012 | Absent_Koike2012 |
| ENSMUSG00000019518 | Ap4m1 | 5 | 138613230 | 138619936 | + | 0.00014453 | 0.00436795 | 1.12867615 | 1.56961592 | 1.30836464 | 0.92221643 | 1.56961592 | ZT6 | Absent_Koike2012 | Absent_Koike2012 |
| ENSMUSG00000020604 | Arsg | 11 | 109334688 | 109434644 | + | 0.00014592 | 0.00438917 | 2.99999766 | 3.4564471 | 1.40815654 | 1.83638656 | 3.4564471 | ZT6 | Present_Koike2012 | Present_Koike2012 |
| ENSMUSG00000024969 | Mark2 | 19 | 7349886 | 7416350 | - | 0.00014828 | 0.0044235 | 0.81558747 | 0.90913854 | 0.80178949 | 0.67160289 | 0.90913854 | ZT6 | Absent_Koike2012 | Absent_Koike2012 |
| ENSMUSG00000068749 | Psma5 | 3 | 108059844 | 108082892 | + | 0.00014777 | 0.0044235 | 30.276377 | 32.3560553 | 30.5769803 | 25.2478248 | 32.3560553 | ZT6 | Absent_Koike2012 | Absent_Koike2012 |
| ENSMUSG00000091264 | BC024659 | 13 | 41345229 | 41371943 | + | 0.0001498 | 0.00445843 | 4.363584 | 6.2197545 | 4.81244525 | 4.29921225 | 6.2197545 | ZT6 | Absent_Koike2012 | Absent_Koike2012 |
| ENSMUSG00000020525 | Ppm1d | 11 | 85124746 | 85160568 | + | 0.00015129 | 0.00448711 | 1.10071165 | 1.39849272 | 1.33064523 | 1.13756175 | 1.39849272 | ZT6 | Absent_Koike2012 | Absent_Koike2012 |
| ENSMUSG00000029022 | Miip | 4 | 147234887 | 147242925 | - | 0.00015214 | 0.00449565 | 1.53900778 | 2.03721387 | 1.87979926 | 1.39301018 | 2.03721387 | ZT6 | Absent_Koike2012 | Absent_Koike2012 |
| ENSMUSG00000048856 | Slc25a47 | 12 | 110089339 | 110095024 | + | 0.00015246 | 0.00449565 | 63.4066898 | 181.359406 | 158.501045 | 65.8350773 | 181.359406 | ZT6 | Present_Koike2012 | Absent_Koike2012 |
| ENSMUSG00000018076 | Med13l | 5 | 119010728 | 119215444 | + | 0.00015651 | 0.0045885 | 1.51800576 | 1.94240385 | 1.5533893 | 1.2812109 | 1.94240385 | ZT6 | Absent_Koike2012 | Absent_Koike2012 |
| ENSMUSG00000044600 | 9130011J15R | 8 | 75089087 | 75094987 | - | 0.00015863 | 0.00464517 | 10.2739822 | 11.8103102 | 11.6658633 | 9.22236224 | 11.8103102 | ZT6 | Present_Koike2012 | Absent_Koike2012 |
| ENSMUSG00000087579 | 1500017E21R | 19 | 36688542 | 36780131 | - | 0.00016046 | 0.00468256 | 7.73512317 | 9.04045186 | 7.46372962 | 4.46442871 | 9.04045186 | ZT6 | Present_Koike2012 | Present_Koike2012 |
| ENSMUSG00000026944 | Abca2 | 2 | 25284223 | 25304060 | + | 0.00016035 | 0.00468256 | 5.41633538 | 5.78269304 | 4.34300093 | 4.01336556 | 5.78269304 | ZT6 | Absent_Koike2012 | Absent_Koike2012 |
| ENSMUSG00000021432 | Slc35b3 | 13 | 39024009 | 39052744 | - | 0.00016198 | 0.004716 | 2.42290621 | 3.14536746 | 2.44781039 | 2.16769605 | 3.14536746 | ZT6 | Present_Koike2012 | Absent_Koike2012 |
| ENSMUSG00000020903 | Stx8 | 11 | 67779695 | 68020650 | + | 0.00016281 | 0.00472961 | 1.02515264 | 1.1417422 | 1.05256563 | 0.87160638 | 1.1417422 | ZT6 | Absent_Koike2012 | Absent_Koike2012 |
| ENSMUSG00000038520 | Tbc1d17 | 7 | 52096271 | 52104449 | - | 0.00016382 | 0.00474245 | 4.34899608 | 5.56384781 | 4.35921297 | 3.33835378 | 5.56384781 | ZT6 | Present_Koike2012 | Absent_Koike2012 |
| ENSMUSG00000028057 | Rit1 | 3 | 88520760 | 88534971 | + | 0.00017044 | 0.00485142 | 1.37166666 | 1.74215137 | 1.26948681 | 1.34477757 | 1.74215137 | ZT6 | Absent_Koike2012 | Absent_Koike2012 |
| ENSMUSG00000028821 | Syf2 | 4 | 134486813 | 134493463 | + | 0.00017133 | 0.00487125 | 9.6935159 | 10.9646888 | 10.8047047 | 8.99150827 | 10.9646888 | ZT6 | Absent_Koike2012 | Absent_Koike2012 |
| ENSMUSG00000030470 | Csrp3 | 7 | 56085768 | 56103400 | - | 0.00017204 | 0.00488232 | 12.1556 | 12.2395516 | 6.46431852 | 5.62315637 | 12.2395516 | ZT6 | Absent_Koike2012 | Absent_Koike2012 |
| ENSMUSG00000033712 | 2610301G19 | 14 | 70537984 | 70553618 | - | 0.00017508 | 0.00493921 | 4.0262835 | 4.37008825 | 3.62492175 | 3.654712 | 4.37008825 | ZT6 | Absent_Koike2012 | Absent_Koike2012 |
| ENSMUSG00000032633 | Flcn | 11 | 59604910 | 59623518 | - | 0.00017957 | 0.00502682 | 2.4848611 | 4.81323676 | 3.80040222 | 2.29235225 | 4.81323676 | ZT6 | Present_Koike2012 | Absent_Koike2012 |
| ENSMUSG00000025607 | Copg2 | 6 | 30697554 | 30846794 | - | 0.00018288 | 0.00506768 | 6.38843489 | 6.87901875 | 6.11524909 | 5.31823645 | 6.87901875 | ZT6 | Present_Koike2012 | Absent_Koike2012 |
| ENSMUSG00000055720 | Ubl7 | 9 | 57758793 | 57777775 | + | 0.00018188 | 0.00506768 | 7.03911867 | 8.21687489 | 7.22675904 | 7.00011592 | 8.21687489 | ZT6 | Absent_Koike2012 | Absent_Koike2012 |
| ENSMUSG00000078249 | Hmga1-rs1 | 11 | 120624108 | 120625733 | + | 0.00018329 | 0.00506768 | 0.75242167 | 1.13549592 | 1.00822042 | 0.64198085 | 1.13549592 | ZT6 | Absent_Koike2012 | Absent_Koike2012 |
| ENSMUSG00000059669 | Taf1b | 12 | 25183446 | 25243436 | + | 0.00018299 | 0.00506768 | 1.2968745 | 1.92026375 | 1.39641425 | 1.1545175 | 1.92026375 | ZT6 | Absent_Koike2012 | Absent_Koike2012 |
| ENSMUSG00000060798 | Intu | 3 | 40430208 | 40508696 | + | 0.00018448 | 0.00509169 | 0.60639325 | 0.7153716 | 0.57582995 | 0.4693444 | 0.7153716 | ZT6 | Absent_Koike2012 | Absent_Koike2012 |
| ENSMUSG00000087161 | Gm16845 | 9 | 21875476 | 21890566 | + | 0.00018551 | 0.00510896 | 0.2225725 | 0.3180165 | 0.27389075 | 0.1417715 | 0.3180165 | ZT6 | Absent_Koike2012 | Absent_Koike2012 |
| ENSMUSG00000021013 | Ttc8 | 12 | 100158784 | 100221448 | + | 0.00018592 | 0.0051148 | 1.08602177 | 1.17956594 | 1.01183418 | 0.86256556 | 1.17956594 | ZT6 | Absent_Koike2012 | Absent_Koike2012 |
| ENSMUSG00000045691 | Thtpa | 14 | 55713672 | 55717823 | + | 0.00019277 | 0.00528597 | 6.41582725 | 6.727374 | 5.19529125 | 6.16340325 | 6.727374 | ZT6 | Absent_Koike2012 | Absent_Koike2012 |
| ENSMUSG00000041777 | Cir1 | 2 | 73121162 | 73150758 | - | 0.00019604 | 0.00535835 | 2.64066102 | 3.93228033 | 3.02658779 | 2.69648521 | 3.93228033 | ZT6 | Absent_Koike2012 | Absent_Koike2012 |
| ENSMUSG00000060961 | Slc4a4 | 5 | 89315843 | 89668678 | + | 0.00019797 | 0.00540529 | 4.92380207 | 5.87070833 | 3.96828509 | 2.73793302 | 5.87070833 | ZT6 | Present_Koike2012 | Absent_Koike2012 |
| ENSMUSG00000040525 | Cblc | 7 | 20364230 | 20382158 | - | 0.00019988 | 0.00543983 | 3.44782796 | 4.25644803 | 3.72122106 | 3.06256171 | 4.25644803 | ZT6 | Absent_Koike2012 | Absent_Koike2012 |
| ENSMUSG00000024983 | Vti1a | 19 | 55390817 | 55701797 | + | 0.0002019 | 0.00548319 | 0.94722319 | 1.22256099 | 1.08002149 | 0.7838845 | 1.22256099 | ZT6 | Absent_Koike2012 | Absent_Koike2012 |
| ENSMUSG00000024759 | Atl3 | 19 | 7568530 | 7613098 | + | 0.00020394 | 0.00553256 | 12.1513846 | 14.0241353 | 13.5173381 | 11.3079977 | 14.0241353 | ZT6 | Present_Koike2012 | Present_Koike2012 |
| ENSMUSG00000038764 | Ptpn3 | 4 | 57203713 | 57314709 | - | 0.00020434 | 0.00553751 | 3.8572005 | 5.16176088 | 5.0185006 | 4.55320613 | 5.16176088 | ZT6 | Present_Koike2012 | Absent_Koike2012 |
| ENSMUSG00000026843 | Fubp3 | 2 | 31428171 | 31473046 | + | 0.00020517 | 0.00554821 | 2.4283032 | 3.10116086 | 2.78025579 | 2.14684398 | 3.10116086 | ZT6 | Absent_Koike2012 | Absent_Koike2012 |
| ENSMUSG00000028191 | Bcl10 | 3 | 145587322 | 145597247 | + | 0.00021309 | 0.00566609 | 3.89642218 | 5.02986796 | 4.28555211 | 3.66488507 | 5.02986796 | ZT6 | Absent_Koike2012 | Absent_Koike2012 |
| ENSMUSG00000047866 | Lonp2 | 8 | 89147942 | 89247772 | + | 0.00021537 | 0.00570295 | 40.6164063 | 43.8543442 | 35.7611507 | 29.9651429 | 43.8543442 | ZT6 | Present_Koike2012 | Absent_Koike2012 |
| ENSMUSG00000037608 | Bclaf1 | 10 | 20032275 | 20062307 | + | 0.00021576 | 0.00570742 | 6.22801528 | 8.2179635 | 6.16176988 | 3.947093 | 8.2179635 | ZT6 | Present_Koike2012 | Present_Koike2012 |
| ENSMUSG00000029110 | Rnf4 | 5 | 34688360 | 34696083 | + | 0.00021754 | 0.00573766 | 21.9494962 | 24.4101245 | 21.6853902 | 18.2668745 | 24.4101245 | ZT6 | Present_Koike2012 | Absent_Koike2012 |
| ENSMUSG00000052331 | Ankrd44 | 1 | 54702184 | 54983231 | - | 0.00022107 | 0.00579834 | 0.7488135 | 0.898097 | 0.74238375 | 0.65679425 | 0.898097 | ZT6 | Absent_Koike2012 | Absent_Koike2012 |
| ENSMUSG00000049232 | Tigd2 | 6 | 59158864 | 59162027 | + | 0.0002272 | 0.00591169 | 3.009977 | 4.16472175 | 3.611956 | 3.010637 | 4.16472175 | ZT6 | Absent_Koike2012 | Absent_Koike2012 |
| ENSMUSG00000040746 | Rnf167 | 11 | 70460737 | 70464923 | + | 0.00022877 | 0.00594648 | 2.85437101 | 3.58048331 | 2.70956589 | 2.59842475 | 3.58048331 | ZT6 | Present_Koike2012 | Present_Koike2012 |
| ENSMUSG00000087413 | Gm11266 | 4 | 82153920 | 82193196 | + | 0.00023567 | 0.00604067 | 0.55385475 | 0.92877175 | 0.36153925 | 0.760773 | 0.92877175 | ZT6 | Absent_Koike2012 | Absent_Koike2012 |
| ENSMUSG00000031930 | Wwp2 | 8 | 109960298 | 110082494 | + | 0.0002357 | 0.00604067 | 3.33889944 | 4.23684355 | 3.14589577 | 2.80668683 | 4.23684355 | ZT6 | Absent_Koike2012 | Absent_Koike2012 |

| | | | | | | | | | | | | | | | |
|---|---|---|---|---|---|---|---|---|---|---|---|---|---|---|---|
| **ENSMUSG00000053333** | Dis3l2 | 1 | 88600379 | 88946672 | + | 0.00023493 | 0.00604067 | 3.39104287 | 3.99928223 | 3.98502616 | 2.78202604 | 3.99928223 | ZT6 | Present_Koike2012 | Absent_Koike2012 |
| **ENSMUSG00000076520** | Igkv1-110 | 6 | 68220521 | 68221261 | + | 0.00023661 | 0.00605121 | 0 | 0.20363725 | 0.0667585 | 0 | 0.20363725 | ZT6 | Absent_Koike2012 | Absent_Koike2012 |
| **ENSMUSG00000005362** | Crbn | 6 | 106728243 | 106750068 | - | 0.00023788 | 0.00606547 | 4.23144984 | 5.2352071 | 4.7586324 | 3.88833496 | 5.2352071 | ZT6 | Present_Koike2012 | Absent_Koike2012 |
| **ENSMUSG00000024246** | Thumpd2 | 17 | 81425667 | 81464425 | - | 0.00023872 | 0.0060746 | 0.70063564 | 0.89415604 | 0.60426768 | 0.58519516 | 0.89415604 | ZT6 | Absent_Koike2012 | Absent_Koike2012 |
| **ENSMUSG00000028402** | Mpdz | 4 | 80924404 | 81088719 | - | 0.00023866 | 0.0060746 | 1.59600722 | 2.09377864 | 1.70371181 | 1.48979905 | 2.09377864 | ZT6 | Present_Koike2012 | Absent_Koike2012 |
| **ENSMUSG00000034484** | Snx2 | 18 | 53336019 | 53380514 | + | 0.00024061 | 0.00610441 | 15.6769913 | 18.3007638 | 17.9432018 | 14.4168345 | 18.3007638 | ZT6 | Present_Koike2012 | Absent_Koike2012 |
| **ENSMUSG00000031666** | Rbl2 | 8 | 93593956 | 93647743 | + | 0.00024367 | 0.00615142 | 3.9425595 | 4.93533867 | 4.31290321 | 3.70465416 | 4.93533867 | ZT6 | Present_Koike2012 | Absent_Koike2012 |
| **ENSMUSG00000031198** | Fundc2 | X | 72627738 | 72641798 | + | 0.00024711 | 0.00620437 | 1.40126816 | 2.07922542 | 1.90169822 | 1.57548196 | 2.07922542 | ZT6 | Absent_Koike2012 | Absent_Koike2012 |
| **ENSMUSG00000028936** | Rpl22 | 4 | 151699851 | 151708180 | + | 0.0002513 | 0.00627133 | 14.528319 | 15.2132741 | 14.7616483 | 12.2473853 | 15.2132741 | ZT6 | Absent_Koike2012 | Absent_Koike2012 |
| **ENSMUSG00000070315** | 4930581F22I | 9 | 34924313 | 34938507 | + | 0.00025162 | 0.00627133 | 0.77798092 | 1.31279047 | 1.06518893 | 0.7501254 | 1.31279047 | ZT6 | Absent_Koike2012 | Absent_Koike2012 |
| **ENSMUSG00000040557** | Wbscr27 | 5 | 135408238 | 135418507 | + | 0.00025214 | 0.00627809 | 0.77989629 | 1.05291526 | 0.80782763 | 0.64121579 | 1.05291526 | ZT6 | Absent_Koike2012 | Present_Koike2012 |
| **ENSMUSG00000055493** | Epm2a | 10 | 11063202 | 11179442 | + | 0.00025293 | 0.00629148 | 1.59303075 | 1.9611735 | 1.70259575 | 1.36417253 | 1.9611735 | ZT6 | Absent_Koike2012 | Absent_Koike2012 |
| **ENSMUSG00000073678** | Pgap1 | 1 | 54529844 | 54532810 | - | 0.00025589 | 0.00631908 | 1.015013 | 2.024808 | 1.8984785 | 1.13982275 | 2.024808 | ZT6 | Present_Koike2012 | Absent_Koike2012 |
| **ENSMUSG00000021820** | Camk2g | 14 | 21554097 | 21613310 | - | 0.00025877 | 0.00634021 | 1.15175971 | 1.72798585 | 1.39209601 | 1.13832313 | 1.72798585 | ZT6 | Absent_Koike2012 | Absent_Koike2012 |
| **ENSMUSG00000026273** | Mterfd2 | 1 | 95195788 | 95202492 | - | 0.00025891 | 0.00634021 | 1.850426 | 2.03260943 | 1.54027846 | 1.52500922 | 2.03260943 | ZT6 | Present_Koike2012 | Absent_Koike2012 |
| **ENSMUSG00000031503** | Col4a2 | 8 | 11312805 | 11449287 | + | 0.00026233 | 0.00638864 | 3.8259826 | 4.5285986 | 4.03004315 | 2.61291846 | 4.5285986 | ZT6 | Absent_Koike2012 | Absent_Koike2012 |
| **ENSMUSG00000029613** | Eif2ak1 | 5 | 144578657 | 144665125 | + | 0.00026884 | 0.00648444 | 4.35621666 | 4.69231613 | 4.30476478 | 3.88487135 | 4.69231613 | ZT6 | Absent_Koike2012 | Absent_Koike2012 |
| **ENSMUSG00000030849** | Fgfr2 | 7 | 137305965 | 140315033 | - | 0.00027185 | 0.00653235 | 2.24759604 | 3.29709333 | 2.52527602 | 1.8990211 | 3.29709333 | ZT6 | Present_Koike2012 | Absent_Koike2012 |
| **ENSMUSG00000057440** | Mpp7 | 18 | 7347960 | 7626861 | - | 0.00027182 | 0.00653235 | 1.13039587 | 1.84845392 | 1.43768716 | 0.99556042 | 1.84845392 | ZT6 | Absent_Koike2012 | Absent_Koike2012 |
| **ENSMUSG00000027890** | Gstm4 | 3 | 107843326 | 107847812 | - | 0.00027278 | 0.006546 | 14.3235052 | 17.0937735 | 13.9639193 | 11.047699 | 17.0937735 | ZT6 | Present_Koike2012 | Absent_Koike2012 |
| **ENSMUSG00000030509** | Asb7 | 7 | 73789451 | 73834482 | - | 0.00027586 | 0.00659134 | 1.2394799 | 1.67785557 | 1.67508851 | 1.37267783 | 1.67785557 | ZT6 | Absent_Koike2012 | Absent_Koike2012 |
| **ENSMUSG00000060470** | Gpr97 | 8 | 97541592 | 97569147 | + | 0.00027996 | 0.00666426 | 1.53565325 | 1.63573475 | 1.29771525 | 0.82747175 | 1.63573475 | ZT6 | Present_Koike2012 | Absent_Koike2012 |
| **ENSMUSG00000020130** | Tbc1d15 | 10 | 114634933 | 114688549 | - | 0.00028032 | 0.00666666 | 3.73433861 | 6.03286231 | 5.80624921 | 3.78168592 | 6.03286231 | ZT6 | Present_Koike2012 | Absent_Koike2012 |
| **ENSMUSG00000020166** | Cnot2 | 10 | 115922217 | 116018567 | - | 0.00028367 | 0.00673308 | 5.23846107 | 5.97595243 | 5.25080105 | 4.76369117 | 5.97595243 | ZT6 | Present_Koike2012 | Present_Koike2012 |
| **ENSMUSG00000000127** | Fer | 17 | 64213364 | 64488843 | + | 0.00028849 | 0.00681634 | 1.3034569 | 1.64563312 | 1.46752806 | 1.32178242 | 1.64563312 | ZT6 | Absent_Koike2012 | Absent_Koike2012 |
| **ENSMUSG00000024493** | Lars | 18 | 42362004 | 42421781 | - | 0.00028975 | 0.00683972 | 8.14820972 | 9.70733696 | 9.26176565 | 7.38380187 | 9.70733696 | ZT6 | Absent_Koike2012 | Absent_Koike2012 |
| **ENSMUSG00000031337** | Mtm1 | X | 68463942 | 68568866 | + | 0.00029111 | 0.0068596 | 0.72952342 | 1.00574625 | 0.91474531 | 0.72454419 | 1.00574625 | ZT6 | Absent_Koike2012 | Absent_Koike2012 |
| **ENSMUSG00000028128** | F3 | 3 | 121426455 | 121437970 | + | 0.00029361 | 0.00688697 | 0.94444775 | 1.68247175 | 1.337947 | 1.09295475 | 1.68247175 | ZT6 | Absent_Koike2012 | Absent_Koike2012 |
| **ENSMUSG00000031955** | Bcar1 | 8 | 114234367 | 114267748 | - | 0.0002958 | 0.00691144 | 5.01606141 | 6.92257674 | 4.74127765 | 4.58827836 | 6.92257674 | ZT6 | Absent_Koike2012 | Absent_Koike2012 |
| **ENSMUSG00000074578** | 1500012F01I | 2 | 166888434 | 166891362 | + | 0.00029605 | 0.00691144 | 1.33516972 | 1.73071861 | 1.57658334 | 1.03189508 | 1.73071861 | ZT6 | Absent_Koike2012 | Absent_Koike2012 |
| **ENSMUSG00000027068** | Dhrs9 | 2 | 69218502 | 69242590 | + | 0.00029808 | 0.00695247 | 1.44185325 | 1.6506685 | 0.25089625 | 1.02989675 | 1.6506685 | ZT6 | Present_Koike2012 | Absent_Koike2012 |
| **ENSMUSG00000029229** | Chic2 | 5 | 75397051 | 75440799 | - | 0.00029961 | 0.00698193 | 1.13162756 | 1.69298291 | 1.55365632 | 1.12926487 | 1.69298291 | ZT6 | Absent_Koike2012 | Absent_Koike2012 |
| **ENSMUSG00000020064** | Herc4 | 10 | 62706659 | 62780622 | + | 0.0003058 | 0.00710525 | 11.2467523 | 13.8585505 | 12.319089 | 11.230093 | 13.8585505 | ZT6 | Absent_Koike2012 | Present_Koike2012 |
| **ENSMUSG00000039349** | C130074G19 | 1 | 186695805 | 186707097 | - | 0.00030609 | 0.00710525 | 32.6153126 | 36.2109338 | 26.3066635 | 28.5143178 | 36.2109338 | ZT6 | Present_Koike2012 | Absent_Koike2012 |
| **ENSMUSG00000031295** | Phka2 | X | 156940098 | 157036810 | + | 0.00030895 | 0.00715359 | 2.77102727 | 3.1701256 | 2.89860207 | 2.29912936 | 3.1701256 | ZT6 | Absent_Koike2012 | Absent_Koike2012 |
| **ENSMUSG00000007564** | Ppp2r1a | 17 | 21082275 | 21102880 | + | 0.00031199 | 0.00719922 | 10.4288243 | 10.8177279 | 9.80193665 | 8.85601298 | 10.8177279 | ZT6 | Absent_Koike2012 | Absent_Koike2012 |
| **ENSMUSG00000035757** | 1300018J18F | 15 | 88919514 | 88930770 | + | 0.00031375 | 0.00720585 | 8.09196786 | 9.66054019 | 9.14305523 | 7.66681585 | 9.66054019 | ZT6 | Present_Koike2012 | Absent_Koike2012 |
| **ENSMUSG00000085772** | D630024D03 | 11 | 31690113 | 31724524 | - | 0.0003146 | 0.00721228 | 0.19178161 | 0.39026929 | 0.21453444 | 0.08984074 | 0.39026929 | ZT6 | Absent_Koike2012 | Absent_Koike2012 |
| **ENSMUSG00000027412** | Lpin3 | 2 | 160706406 | 160731738 | + | 0.00031435 | 0.00721228 | 0.91480294 | 1.2806839 | 0.96043486 | 0.70660742 | 1.2806839 | ZT6 | Absent_Koike2012 | Absent_Koike2012 |
| **ENSMUSG00000022833** | Ccdc14 | 16 | 34690702 | 34725280 | + | 0.00032038 | 0.00730529 | 0.12038925 | 0.23168175 | 0.2255925 | 0.14347525 | 0.23168175 | ZT6 | Absent_Koike2012 | Absent_Koike2012 |
| **ENSMUSG00000028580** | Pum1 | 4 | 130219236 | 130337479 | + | 0.00032501 | 0.00738758 | 5.56228459 | 7.32221277 | 6.02463098 | 5.54097562 | 7.32221277 | ZT6 | Absent_Koike2012 | Absent_Koike2012 |
| **ENSMUSG00000028381** | Ugcg | 4 | 59207328 | 59235705 | + | 0.0003302 | 0.00747567 | 2.70967287 | 4.62065958 | 3.55842515 | 2.45525692 | 4.62065958 | ZT6 | Present_Koike2012 | Absent_Koike2012 |
| **ENSMUSG00000032870** | Smap2 | 4 | 120640922 | 120689852 | - | 0.00033072 | 0.00748065 | 4.4474606 | 5.02716628 | 3.85706629 | 4.04651979 | 5.02716628 | ZT6 | Absent_Koike2012 | Absent_Koike2012 |
| **ENSMUSG00000052837** | Junb | 8 | 87500811 | 87502617 | - | 0.00033636 | 0.00753834 | 2.69285725 | 9.98846875 | 4.92630075 | 3.667986 | 9.98846875 | ZT6 | Absent_Koike2012 | Absent_Koike2012 |
| **ENSMUSG00000086841** | 2410006H16 | 11 | 62416379 | 62418309 | + | 0.0003372 | 0.00753834 | 7.23726975 | 11.2826318 | 9.31167675 | 5.81027275 | 11.2826318 | ZT6 | Absent_Koike2012 | Absent_Koike2012 |
| **ENSMUSG00000050199** | Lgr4 | 2 | 109757804 | 109854414 | + | 0.00033771 | 0.00753834 | 11.617749 | 16.421407 | 16.0376397 | 13.6465599 | 16.421407 | ZT6 | Absent_Koike2012 | Absent_Koike2012 |
| **ENSMUSG00000039981** | Zc3h12d | 10 | 7552268 | 7590195 | + | 0.0003387 | 0.00755373 | 0.558034 | 0.99407475 | 0.391222 | 0.53617025 | 0.99407475 | ZT6 | Absent_Koike2012 | Absent_Koike2012 |
| **ENSMUSG00000066440** | Zfyve26 | 12 | 80333334 | 80397269 | - | 0.00034086 | 0.00759463 | 2.847475 | 3.7919985 | 2.83967125 | 2.66132775 | 3.7919985 | ZT6 | Absent_Koike2012 | Absent_Koike2012 |
| **ENSMUSG00000030180** | Kdm5a | 6 | 120314117 | 120394592 | + | 0.00034591 | 0.00768087 | 2.15209464 | 2.78598489 | 2.47104892 | 1.7220857 | 2.78598489 | ZT6 | Absent_Koike2012 | Absent_Koike2012 |
| **ENSMUSG00000091144** | Gm6907 | 14 | 59999671 | 60014218 | - | 0.00034668 | 0.00769137 | 1.84807452 | 2.42288204 | 2.23675827 | 1.63178825 | 2.42288204 | ZT6 | Absent_Koike2012 | Absent_Koike2012 |
| **ENSMUSG00000047394** | Odf3b | 15 | 89207881 | 89209685 | - | 0.00035114 | 0.00775653 | 1.933768 | 2.017403 | 1.12563 | 1.17442875 | 2.017403 | ZT6 | Absent_Koike2012 | Absent_Koike2012 |
| **ENSMUSG00000034042** | Gpbp1l1 | 4 | 116230263 | 116266505 | + | 0.00035245 | 0.00777201 | 10.8605985 | 12.7670492 | 12.1098266 | 10.1198911 | 12.7670492 | ZT6 | Absent_Koike2012 | Absent_Koike2012 |
| **ENSMUSG00000021022** | Ppp2r3c | 12 | 56381801 | 56403987 | - | 0.00035233 | 0.00777201 | 3.301693 | 3.36446925 | 2.94657575 | 2.64834775 | 3.36446925 | ZT6 | Absent_Koike2012 | Absent_Koike2012 |
| **ENSMUSG00000089716** | Gm6264 | 1 | 85156339 | 85190835 | + | 0.0003549 | 0.00780587 | 1.25051722 | 1.63844366 | 1.51825418 | 1.17689351 | 1.63844366 | ZT6 | Absent_Koike2012 | Absent_Koike2012 |
| **ENSMUSG00000038224** | Serpinf2 | 11 | 75245234 | 75253093 | - | 0.00037025 | 0.0080459 | 220.558764 | 269.891848 | 189.440349 | 155.658477 | 269.891848 | ZT6 | Present_Koike2012 | Present_Koike2012 |
| **ENSMUSG00000045193** | Cirbp | 10 | 79628730 | 79635531 | + | 0.0003709 | 0.00804923 | 1.4182789 | 3.28022577 | 2.15166536 | 1.40674474 | 3.28022577 | ZT6 | Absent_Koike2012 | Absent_Koike2012 |
| **ENSMUSG00000051853** | Arf3 | 15 | 98568060 | 98593645 | - | 0.0003746 | 0.0081058 | 7.03002752 | 8.61803476 | 8.57917528 | 6.33097717 | 8.61803476 | ZT6 | Present_Koike2012 | Absent_Koike2012 |
| **ENSMUSG00000033392** | Clasp2 | 9 | 113650591 | 113828815 | + | 0.00038482 | 0.00828957 | 1.81683156 | 2.10909262 | 2.01889834 | 1.69750879 | 2.10909262 | ZT6 | Present_Koike2012 | Present_Koike2012 |
| **ENSMUSG00000027530** | Fabp12 | 3 | 10244209 | 10301183 | - | 0.00038675 | 0.00829284 | 0.03574021 | 0.08757708 | 0.01857537 | 0.01019927 | 0.08757708 | ZT6 | Absent_Koike2012 | Absent_Koike2012 |
| **ENSMUSG00000039637** | Coro7 | 16 | 4626133 | 4679777 | - | 0.00039344 | 0.00841327 | 1.02457724 | 1.14395235 | 1.13211099 | 0.86789371 | 1.14395235 | ZT6 | Absent_Koike2012 | Present_Koike2012 |
| **ENSMUSG00000038206** | Fbxo8 | 8 | 59029887 | 59071190 | + | 0.00039651 | 0.00845762 | 9.99177178 | 11.3680981 | 8.98095341 | 9.02861541 | 11.3680981 | ZT6 | Present_Koike2012 | Absent_Koike2012 |
| **ENSMUSG00000028670** | Lypla2 | 4 | 135524139 | 135528541 | - | 0.00039925 | 0.00848773 | 7.09320295 | 7.3543862 | 6.5295628 | 6.02225092 | 7.3543862 | ZT6 | Absent_Koike2012 | Absent_Koike2012 |
| **ENSMUSG00000044022** | Pcdhb21 | 18 | 37673306 | 37677967 | + | 0.00039991 | 0.00849457 | 0.02283325 | 0.0501595 | 0.027304 | 0.042236 | 0.0501595 | ZT6 | Absent_Koike2012 | Absent_Koike2012 |
| **ENSMUSG00000032369** | Plscr1 | 9 | 92144895 | 92166813 | + | 0.00040182 | 0.0085174 | 4.92981125 | 6.80733775 | 5.91315525 | 4.44640825 | 6.80733775 | ZT6 | Absent_Koike2012 | Absent_Koike2012 |
| **ENSMUSG00000031928** | Mre11a | 9 | 14589090 | 14641567 | + | 0.00040603 | 0.00856735 | 1.05751775 | 1.12625516 | 1.09647755 | 0.93094379 | 1.12625516 | ZT6 | Absent_Koike2012 | Absent_Koike2012 |
| **ENSMUSG00000033628** | Pik3c3 | 18 | 30432401 | 30507780 | + | 0.00041062 | 0.00863704 | 3.18748536 | 3.8909862 | 3.53572645 | 3.09555249 | 3.8909862 | ZT6 | Absent_Koike2012 | Absent_Koike2012 |
| **ENSMUSG00000002227** | Mov10 | 3 | 104597754 | 104621481 | - | 0.00041619 | 0.00871667 | 3.48741843 | 4.84620243 | 4.317241 | 3.29568319 | 4.84620243 | ZT6 | Absent_Koike2012 | Absent_Koike2012 |
| **ENSMUSG00000025920** | Stau2 | 1 | 16218755 | 16510193 | - | 0.00041813 | 0.00873566 | 0.45967564 | 0.53303476 | 0.49479269 | 0.35577477 | 0.53303476 | ZT6 | Absent_Koike2012 | Absent_Koike2012 |
| **ENSMUSG00000031586** | Rbpms | 8 | 34893116 | 35040313 | - | 0.00041998 | 0.00875277 | 3.73649744 | 4.72969879 | 3.0954695 | 3.36744189 | 4.72969879 | ZT6 | Absent_Koike2012 | Absent_Koike2012 |

| | | | | | | | | | | | | | | | |
|---|---|---|---|---|---|---|---|---|---|---|---|---|---|---|---|
| **ENSMUSG00000070697** | Utp3 | 5 | 88983508 | 88985108 | + | 0.00042485 | 0.00880089 | 11.213015 | 11.4854495 | 9.35171175 | 8.7107945 | 11.4854495 | ZT6 | Absent_Koike2012 | Absent_Koike2012 |
| **ENSMUSG00000053470** | Kdm3a | 6 | 71538968 | 71582899 | - | 0.0004249 | 0.00880089 | 2.54544254 | 3.66617682 | 2.78476854 | 2.24853218 | 3.66617682 | ZT6 | Absent_Koike2012 | Absent_Koike2012 |
| **ENSMUSG00000059552** | Trp53 | 11 | 69393861 | 69405375 | + | 0.0004276 | 0.00881784 | 1.29039338 | 1.43331293 | 1.08667062 | 0.98599713 | 1.43331293 | ZT6 | Absent_Koike2012 | Absent_Koike2012 |
| **ENSMUSG00000027455** | Nsfl1c | 2 | 151319918 | 151337150 | + | 0.00043699 | 0.00897507 | 5.25693719 | 5.72814809 | 5.31254521 | 4.74651465 | 5.72814809 | ZT6 | Absent_Koike2012 | Absent_Koike2012 |
| **ENSMUSG00000032492** | Pth1r | 9 | 110624592 | 110649649 | - | 0.00043857 | 0.00900033 | 1.8690987 | 2.65299691 | 2.2859305 | 1.78485116 | 2.65299691 | ZT6 | Absent_Koike2012 | Absent_Koike2012 |
| **ENSMUSG00000032754** | Slc24a6 | 5 | 120961177 | 120984033 | + | 0.00044027 | 0.00902786 | 3.27456938 | 7.44068005 | 4.16723429 | 3.40601249 | 7.44068005 | ZT6 | Present_Koike2012 | Absent_Koike2012 |
| **ENSMUSG00000034765** | Dusp5 | 19 | 53603599 | 53616921 | + | 0.00044603 | 0.00910931 | 0.11617925 | 0.2206995 | 0.187991 | 0.0975425 | 0.2206995 | ZT6 | Absent_Koike2012 | Absent_Koike2012 |
| **ENSMUSG00000030309** | Caprin2 | 6 | 148791014 | 148844759 | - | 0.00044869 | 0.009127 | 0.09701271 | 0.13426311 | 0.07155044 | 0.04729748 | 0.13426311 | ZT6 | Absent_Koike2012 | Absent_Koike2012 |
| **ENSMUSG00000032502** | Stac | 9 | 111463941 | 111592852 | - | 0.00044789 | 0.009127 | 0.00390577 | 0.01408625 | 0 | 0.00202025 | 0.01408625 | ZT6 | Absent_Koike2012 | Absent_Koike2012 |
| **ENSMUSG00000003345** | Csnk1g2 | 10 | 80085525 | 80103516 | + | 0.0004486 | 0.009127 | 10.0006299 | 11.6501414 | 11.5489284 | 9.56754169 | 11.6501414 | ZT6 | Absent_Koike2012 | Absent_Koike2012 |
| **ENSMUSG00000022358** | Fbxo32 | 15 | 58011634 | 58046433 | - | 0.00046614 | 0.00938438 | 0.374786 | 1.07500025 | 0.64324925 | 0.4624025 | 1.07500025 | ZT6 | Absent_Koike2012 | Absent_Koike2012 |
| **ENSMUSG00000024155** | 4930528F23I | 17 | 24941327 | 24976732 | + | 0.00047732 | 0.00955915 | 1.85572343 | 3.25090956 | 2.12450783 | 1.39131528 | 3.25090956 | ZT6 | Absent_Koike2012 | Absent_Koike2012 |
| **ENSMUSG00000037872** | Darc | 1 | 175262017 | 175263634 | - | 0.00048237 | 0.00963498 | 0.3904885 | 0.5689405 | 0.23909525 | 0.31307675 | 0.5689405 | ZT6 | Absent_Koike2012 | Absent_Koike2012 |
| **ENSMUSG00000024130** | Abca3 | 17 | 24488895 | 24547146 | + | 0.00049993 | 0.00988469 | 7.73929161 | 7.75326716 | 6.87028754 | 6.57126665 | 7.75326716 | ZT6 | Present_Koike2012 | Absent_Koike2012 |
| **ENSMUSG00000021704** | Mtx3 | 13 | 93614742 | 93628185 | + | 0.00050743 | 0.0099866 | 1.84254225 | 2.78191125 | 1.78026575 | 1.338281 | 2.78191125 | ZT6 | Absent_Koike2012 | Absent_Koike2012 |

Table S4. **Promoter Capture HiC statistics.** HiCUP summary results for independent P-CHiC replicates and significant interactions detected with CHiCAGO.

| Sample | Total number of read pairs | Unique paired alignments | Total number of valid read pairs | % Valid pairs | % Unique valid pairs | % cis reads | % trans reads | Significant interactions from CHiCAGO (merged replicates) |
|---|---|---|---|---|---|---|---|---|
| P-CHiC ZT0 Pool A | 283,175,427 | 184,542,315 | 114,469,166 | 61.45 | 94.25 | 82.9 | 17.05 | 146,262 |
| P-CHiC ZT0 Pool B | 203,761,238 | 146,963,871 | 128,573,361 | 87.5 | 92.5 | 83 | 17 | |
| P-CHiC ZT0 Pool C | 248,116,780 | 186,680,809 | 163,782,589 | 87.7 | 91.1 | 79.2 | 20.8 | |
| P-CHiC ZT6 Pool A | 292,975,816 | 193,087,427 | 109,497,418 | 56.1 | 95.4 | 81.65 | 18.35 | 151,529 |
| P-CHiC ZT6 Pool B | 186,899,728 | 134,195,086 | 118,017,852 | 87.9 | 92.1 | 82 | 18 | |
| P-CHiC ZT6 Pool C | 263,245,280 | 192,761,953 | 176,879,721 | 91.8 | 94.3 | 80.6 | 19.4 | |
| P-CHiC ZT12 Pool A | 302,434,305 | 206,794,563 | 107,682,158 | 51.65 | 96.65 | 82.65 | 17.3 | 151,347 |
| P-CHiC ZT12 Pool B | 161,699,345 | 118,998,859 | 106,620,086 | 89.6 | 93.3 | 82.4 | 17.6 | |
| P-CHiC ZT12 Pool C | 246,045,695 | 176,211,831 | 159,632,206 | 90.6 | 93.1 | 80.3 | 19.7 | |
| P-CHiC ZT18 Pool A | 277,577,887 | 190,448,729 | 107,796,060 | 55.7 | 96.3 | 82.5 | 17.5 | 154,775 |
| P-CHiC ZT18 Pool B | 157,884,747 | 116,064,583 | 105,024,339 | 90.5 | 93.1 | 81.9 | 18.1 | |
| P-CHiC ZT18 Pool C | 236,881,106 | 173,794,906 | 159,098,792 | 91.5 | 90.2 | 79.4 | 20.6 | |

Table S5. **Transcription factor DNA binding motif enrichment analysis using MEME.** This table contains the enrichment results using dynamic and stable interactions of promoters associated with circadian genes from all time point ZT0, 6, 12 and 18 separately.

**ZT0 dynamic**

| MOTIF_INDEX | MOTIF_SOUF | MOTIF_ID | ALT_ID | CONSENSUS | WIDTH | SITES | E-VALUE | E-VALUE_SO | MOST_SIMILAR_MC | MOST_SIMILAR_MO | URL |
|---|---|---|---|---|---|---|---|---|---|---|---|
| 1 | MEME | GSCKGGAGF | MEME-1 | GSCKGGAGF | 29 | 101 | 1.5e-182 | MEME | db/JASPAR/JASPAR | **MA0471.1 (E2F6)** | http://jaspar2018.genereg.net/matrix/MA0471.1 |
| 2 | MEME | YGAGTTCGA | MEME-2 | YGAGTTCGA | 29 | 54 | 1.0e-163 | MEME | db/JASPAR/JASPAR | **MA0505.1 (Nr5a2)** | http://jaspar2018.genereg.net/matrix/MA0505.1 |
| 3 | MEME | TRCRCMKKC | MEME-3 | TRCRCMKKC | 30 | 18 | 7.2e-115 | MEME | | | |
| 4 | DREME | RTGGCTCA | DREME-1 | RTGGCTCA | 8 | 56 | 1.0e-007 | DREME | db/JASPAR/JASPAR | **MA0099.3 (FOS::JUN** | http://jaspar2018.genereg.net/matrix/MA0099.3 |
| 5 | DREME | TAHATA | DREME-2 | TAHATA | 6 | 191 | 4.5e-006 | DREME | | | |
| 6 | DREME | ACHCACA | DREME-4 | ACHCACA | 7 | 148 | 7.4e-005 | DREME | | | |
| 7 | DREME | GGCTGK | DREME-3 | GGCTGK | 6 | 220 | 1.6e-004 | DREME | | | |
| 8 | DREME | GAGGWTC | DREME-5 | GAGGWTC | 7 | 52 | 1.0e-003 | DREME | | | |
| 9 | DREME | GAACTCR | DREME-6 | GAACTCR | 7 | 68 | 1.9e-003 | DREME | db/JASPAR/JASPAR | **MA0693.2 (VDR)** | http://jaspar2018.genereg.net/matrix/MA0693.2 |
| 10 | DREME | GCAARCA | DREME-7 | GCAARCA | 7 | 61 | 5.6e-003 | DREME | | | |
| 11 | DREME | CCGGGASG | DREME-8 | CCGGGASG | 8 | 22 | 7.0e-003 | DREME | db/MOUSE/uniprobe | **UP00022_2 (Zfp740_s** | http://the_brain.bwh.harvard.edu/uniprobe/details2.php?id=00022 |
| 12 | DREME | CGGCATCK | DREME-9 | CGGCATCK | 8 | 22 | 7.0e-003 | DREME | | | |
| 13 | DREME | GGAAGMA | DREME-10 | GGAAGMA | 7 | 85 | 1.0e-002 | DREME | | | |
| 14 | DREME | KCTGGGA | DREME-11 | KCTGGGA | 7 | 79 | 3.8e-002 | DREME | db/JASPAR/JASPAR | **MA0144.2 (STAT3)** | http://jaspar2018.genereg.net/matrix/MA0144.2 |
| 15 | DREME | ATAAWTC | DREME-12 | ATAAWTC | 7 | 42 | 4.2e-002 | DREME | | | |

### MEME-ChIP (Motif Analysis of Large Nucleotide Datasets): Version 5.0.3 released on Sun Dec 02 18:41:45 2018 -0800
### The format of this file is described at http://meme-suite.org//doc/meme-chip-output-format.html
### meme-chip -oc . -time 300 -ccut 100 -order 1 -db db/EUKARYOTE/jolma2013.meme -db db/JASPAR/JASPAR2018_CORE_vertebrates_non-redundant.meme -db db/MOUSE/uniprobe_mouse.meme -meme-mod zoops -meme-minw 6 -meme-maxw 30 -meme-nmotifs 3 -meme-searchsize 1000(

**ZT6 static**

| MOTIF_INDEX | MOTIF_SOUF | MOTIF_ID | ALT_ID | CONSENSUS | WIDTH | SITES | E-VALUE | E-VALUE_SO | MOST_SIMILAR_MC | MOST_SIMILAR_MO | URL |
|---|---|---|---|---|---|---|---|---|---|---|---|
| 1 | MEME | GKRRGYKSV | MEME-2 | GKRRGYKSV | 30 | 91 | 1.8e-103 | MEME | db/JASPAR/JASPAR | **MA0505.1 (Nr5a2)** | http://jaspar2018.genereg.net/matrix/MA0505.1 |
| 2 | MEME | GRGSWGGG | MEME-3 | GRGSWGGG | 21 | 309 | 1.0e-088 | MEME | db/JASPAR/JASPAR | **MA0528.1 (ZNF263)** | http://jaspar2018.genereg.net/matrix/MA0528.1 |
| 3 | MEME | TTTKTTTTTD | MEME-1 | TTTKTTTTTD | 21 | 316 | 6.7e-074 | MEME | db/JASPAR/JASPAR | **MA1125.1 (ZNF384)** | http://jaspar2018.genereg.net/matrix/MA1125.1 |
| 4 | db/JASPAR/JA | MA0508.2 | PRDM1 | TCACTTTCW | 10 | 10 | 1.3e-020 | CENTRIMO | | | http://jaspar2018.genereg.net/matrix/MA0508.2 |
| 5 | db/EUKARYO | NR3C1_DBD | | DRGWACAYD | 17 | 10 | 2.6e-019 | CENTRIMO | | | http://floresta.eead.csic.es/footprintdb/index.php?db=HumanTF:1.0&motif=NR3C1_DBD |
| 6 | db/EUKARYO | NR3C2_DBD | | KRGWACAYW | 17 | 10 | 2.6e-019 | CENTRIMO | | | http://floresta.eead.csic.es/footprintdb/index.php?db=HumanTF:1.0&motif=NR3C2_DBD |
| 7 | db/JASPAR/JA | MA0113.3 | NR3C1 | DRGWACAYD | 17 | 10 | 2.6e-019 | CENTRIMO | | | http://jaspar2018.genereg.net/matrix/MA0113.3 |
| 8 | db/JASPAR/JA | MA0727.1 | NR3C2 | KRGWACAYW | 17 | 10 | 2.6e-019 | CENTRIMO | | | http://jaspar2018.genereg.net/matrix/MA0727.1 |
| 9 | db/MOUSE/ur | UP00122_1 | Tgif1_2342.2 | NABWWTGA | 17 | 10 | 4.5e-015 | CENTRIMO | | | http://the_brain.bwh.harvard.edu/uniprobe/details2.php?id=00122 |
| 10 | db/MOUSE/ur | UP00186_1 | Meis1_2335.1 | WANBASCTG | 16 | 10 | 5.0e-015 | CENTRIMO | | | http://the_brain.bwh.harvard.edu/uniprobe/details2.php?id=00186 |
| 11 | db/MOUSE/ur | UP00226_1 | Mrg1_2246.2 | WANBASCTG | 16 | 10 | 5.0e-015 | CENTRIMO | | | http://the_brain.bwh.harvard.edu/uniprobe/details2.php?id=00226 |
| 12 | db/MOUSE/ur | UP00210_1 | Mrg2_2302.1 | WANBASCTG | 16 | 10 | 5.0e-015 | CENTRIMO | | | http://the_brain.bwh.harvard.edu/uniprobe/details2.php?id=00210 |
| 13 | db/MOUSE/ur | UP00203_1 | Pknox1_2364. | WRNSASCTG | 16 | 10 | 5.0e-015 | CENTRIMO | | | http://the_brain.bwh.harvard.edu/uniprobe/details2.php?id=00203 |
| 14 | db/MOUSE/ur | UP00205_1 | Pknox2_3077. | AARSRSCTG | 16 | 10 | 5.0e-015 | CENTRIMO | | | http://the_brain.bwh.harvard.edu/uniprobe/details2.php?id=00205 |
| 15 | db/MOUSE/ur | UP00258_1 | Tgif2_3451.1 | AANBAGCTG | 16 | 10 | 5.0e-015 | CENTRIMO | | | http://the_brain.bwh.harvard.edu/uniprobe/details2.php?id=00258 |
| 16 | db/JASPAR/JA | MA0499.1 | Myod1 | NSCAGCTGY | 13 | 10 | 5.6e-015 | CENTRIMO | | | http://jaspar2018.genereg.net/matrix/MA0499.1 |
| 17 | db/EUKARYO | SNAI2_DBD | | DRCAGGTGE | 9 | 10 | 6.9e-015 | CENTRIMO | | | http://floresta.eead.csic.es/footprintdb/index.php?db=HumanTF:1.0&motif=SNAI2_DBD |
| 18 | db/JASPAR/JA | MA0745.1 | SNAI2 | DRCAGGTGE | 9 | 10 | 6.9e-015 | CENTRIMO | | | http://jaspar2018.genereg.net/matrix/MA0745.1 |
| 19 | db/EUKARYO | Ascl2_DBD | | ARCAGCTGC | 10 | 10 | 7.6e-015 | CENTRIMO | | | http://floresta.eead.csic.es/footprintdb/index.php?db=HumanTF:1.0&motif=Ascl2_DBD |
| 20 | db/JASPAR/JA | MA0816.1 | Ascl2 | ARCAGCTGC | 10 | 10 | 7.6e-015 | CENTRIMO | | | http://jaspar2018.genereg.net/matrix/MA0816.1 |
| 21 | db/EUKARYO | HOXA13_DBD_2 | | BCTCGTAAA | 11 | 10 | 3.2e-014 | CENTRIMO | | | http://floresta.eead.csic.es/footprintdb/index.php?db=HumanTF:1.0&motif=HOXA13_DBD_2 |
| 22 | db/EUKARYO | HOXB13_DBD_2 | | SCTCGTAAA | 11 | 10 | 3.2e-014 | CENTRIMO | | | http://floresta.eead.csic.es/footprintdb/index.php?db=HumanTF:1.0&motif=HOXB13_DBD_2 |
| 23 | db/EUKARYO | HOXC13_DBD_2 | | BCTCGTAAA | 11 | 10 | 3.2e-014 | CENTRIMO | | | http://floresta.eead.csic.es/footprintdb/index.php?db=HumanTF:1.0&motif=HOXC13_DBD_2 |
| 24 | db/EUKARYO | HOXD13_DBD_2 | | GCTCGTAAA | 11 | 10 | 3.2e-014 | CENTRIMO | | | http://floresta.eead.csic.es/footprintdb/index.php?db=HumanTF:1.0&motif=HOXD13_DBD_2 |
| 25 | db/JASPAR/JA | MA0907.1 | HOXC13 | BCTCGTAAA | 11 | 10 | 3.2e-014 | CENTRIMO | | | http://jaspar2018.genereg.net/matrix/MA0907.1 |
| 26 | db/JASPAR/JA | MA0485.1 | Hoxc9 | RGCCATAAAT | 13 | 10 | 3.8e-014 | CENTRIMO | | | http://jaspar2018.genereg.net/matrix/MA0485.1 |
| 27 | db/EUKARYO | CREB3L1_full_2 | | NTGCCACGT | 14 | 10 | 6.8e-014 | CENTRIMO | | | http://floresta.eead.csic.es/footprintdb/index.php?db=HumanTF:1.0&motif=CREB3L1_full_2 |
| 28 | db/JASPAR/JA | MA0161.2 | NFIC | NNCTTGGCA | 11 | 10 | 1.0e-013 | CENTRIMO | | | http://jaspar2018.genereg.net/matrix/MA0161.2 |
| 29 | db/EUKARYO | SOX8_DBD_2 | | ATGAATTKCA | 13 | 10 | 4.4e-012 | CENTRIMO | | | http://floresta.eead.csic.es/footprintdb/index.php?db=HumanTF:1.0&motif=SOX8_DBD_2 |
| 30 | db/MOUSE/ur | UP00076_2 | Rfxdc2_secon | BBRCDTRGA | 17 | 10 | 4.9e-012 | CENTRIMO | | | http://the_brain.bwh.harvard.edu/uniprobe/details2.php?id=00076 |
| 31 | db/MOUSE/ur | UP00056_2 | Rfx4_seconda | WNBCHTRGW | 15 | 10 | 1.2e-011 | CENTRIMO | | | http://the_brain.bwh.harvard.edu/uniprobe/details2.php?id=00056 |
| 32 | db/JASPAR/JA | MA0258.2 | ESR2 | AGGTCASVN | 15 | 10 | 1.4e-011 | CENTRIMO | | | http://jaspar2018.genereg.net/matrix/MA0258.2 |
| 33 | db/JASPAR/JA | MA0504.1 | NR2C2 | VGRGGTCAR | 15 | 10 | 3.1e-011 | CENTRIMO | | | http://jaspar2018.genereg.net/matrix/MA0504.1 |
| 34 | db/JASPAR/JA | MA0059.1 | MAX::MYC | RASCACGTG | 11 | 10 | 6.7e-011 | CENTRIMO | | | http://jaspar2018.genereg.net/matrix/MA0059.1 |
| 35 | db/MOUSE/ur | UP00057_1 | Zic2_primary | NMCCCCCG | 15 | 10 | 2.5e-010 | CENTRIMO | | | http://the_brain.bwh.harvard.edu/uniprobe/details2.php?id=00057 |
| 36 | db/MOUSE/ur | UP00006_1 | Zic3_primary | BMCCCCCG | 15 | 10 | 2.5e-010 | CENTRIMO | | | http://the_brain.bwh.harvard.edu/uniprobe/details2.php?id=00006 |
| 37 | db/JASPAR/JA | MA0138.2 | REST | TTCAGCACC | 21 | 10 | 2.5e-010 | CENTRIMO | | | http://jaspar2018.genereg.net/matrix/MA0138.2 |

| | | | | | | | | | |
|---|---|---|---|---|---|---|---|---|---|
| 38 | db/JASPAR/J/ | MA0062.2 | Gabpa | CCGGAAGTG | 11 | 10 | 1.2e-009 | CENTRIMO | http://jaspar2018.genereg.net/matrix/MA0062.2 |
| 39 | db/JASPAR/J/ | MA0076.2 | ELK4 | BCRCTTCCG | 11 | 10 | 1.2e-009 | CENTRIMO | http://jaspar2018.genereg.net/matrix/MA0076.2 |
| 40 | db/JASPAR/J/ | MA0750.2 | ZBTB7A | NVCCGGAAG | 13 | 10 | 1.2e-009 | CENTRIMO | http://jaspar2018.genereg.net/matrix/MA0750.2 |
| 41 | db/EUKARYO | ELK1_DBD_1 | | ACCGGAAGT | 10 | 10 | 1.3e-009 | CENTRIMO | http://floresta.eead.csic.es/footprintdb/index.php?db=HumanTF:1.0&motif=ELK1_DBD_1 |
| 42 | db/EUKARYO | ELK1_DBD_2 | | ACCGGAAGT | 10 | 10 | 1.3e-009 | CENTRIMO | http://floresta.eead.csic.es/footprintdb/index.php?db=HumanTF:1.0&motif=ELK1_DBD_2 |
| 43 | db/EUKARYO | ELK1_full_1 | | ACCGGAAGT | 10 | 10 | 1.3e-009 | CENTRIMO | http://floresta.eead.csic.es/footprintdb/index.php?db=HumanTF:1.0&motif=ELK1_full_1 |
| 44 | db/EUKARYO | ELK3_DBD | | ACCGGAAGT | 10 | 10 | 1.3e-009 | CENTRIMO | http://floresta.eead.csic.es/footprintdb/index.php?db=HumanTF:1.0&motif=ELK3_DBD |
| 45 | db/EUKARYO | ELK4_DBD | | ACCGGAART | 10 | 10 | 1.3e-009 | CENTRIMO | http://floresta.eead.csic.es/footprintdb/index.php?db=HumanTF:1.0&motif=ELK4_DBD |
| 46 | db/EUKARYO | ERF_DBD | | ACCGGAAGT | 10 | 10 | 1.3e-009 | CENTRIMO | http://floresta.eead.csic.es/footprintdb/index.php?db=HumanTF:1.0&motif=ERF_DBD |
| 47 | db/EUKARYO | ERG_DBD_1 | | ACCGGAAGT | 10 | 10 | 1.3e-009 | CENTRIMO | http://floresta.eead.csic.es/footprintdb/index.php?db=HumanTF:1.0&motif=ERG_DBD_1 |
| 48 | db/EUKARYO | ERG_full_1 | | ACCGGAART | 10 | 10 | 1.3e-009 | CENTRIMO | http://floresta.eead.csic.es/footprintdb/index.php?db=HumanTF:1.0&motif=ERG_full_1 |
| 49 | db/EUKARYO | ETS1_DBD_1 | | ACCGGAART | 10 | 10 | 1.3e-009 | CENTRIMO | http://floresta.eead.csic.es/footprintdb/index.php?db=HumanTF:1.0&motif=ETS1_DBD_1 |
| 50 | db/EUKARYO | ETS1_full_1 | | ACCGGAARY | 10 | 10 | 1.3e-009 | CENTRIMO | http://floresta.eead.csic.es/footprintdb/index.php?db=HumanTF:1.0&motif=ETS1_full_1 |
| 51 | db/EUKARYO | ETV1_DBD | | ACCGGAAGT | 10 | 10 | 1.3e-009 | CENTRIMO | http://floresta.eead.csic.es/footprintdb/index.php?db=HumanTF:1.0&motif=ETV1_DBD |
| 52 | db/EUKARYO | ETV3_DBD | | ACCGGAAGT | 10 | 10 | 1.3e-009 | CENTRIMO | http://floresta.eead.csic.es/footprintdb/index.php?db=HumanTF:1.0&motif=ETV3_DBD |
| 53 | db/EUKARYO | ETV4_DBD | | ACCGGAAGT | 10 | 10 | 1.3e-009 | CENTRIMO | http://floresta.eead.csic.es/footprintdb/index.php?db=HumanTF:1.0&motif=ETV4_DBD |
| 54 | db/EUKARYO | ETV5_DBD | | ACCGGAWGY | 10 | 10 | 1.3e-009 | CENTRIMO | http://floresta.eead.csic.es/footprintdb/index.php?db=HumanTF:1.0&motif=ETV5_DBD |
| 55 | db/EUKARYO | ETV6_full_2 | | MSCGGAAGT | 10 | 10 | 1.3e-009 | CENTRIMO | http://floresta.eead.csic.es/footprintdb/index.php?db=HumanTF:1.0&motif=ETV6_full_2 |
| 56 | db/EUKARYO | Elk3_DBD | | ACCGGAAGT | 10 | 10 | 1.3e-009 | CENTRIMO | http://floresta.eead.csic.es/footprintdb/index.php?db=HumanTF:1.0&motif=Elk3_DBD |
| 57 | db/EUKARYO | FEV_DBD | | ACCGGAAGT | 10 | 10 | 1.3e-009 | CENTRIMO | http://floresta.eead.csic.es/footprintdb/index.php?db=HumanTF:1.0&motif=FEV_DBD |
| 58 | db/EUKARYO | FLI1_DBD_1 | | ACCGGAART | 10 | 10 | 1.3e-009 | CENTRIMO | http://floresta.eead.csic.es/footprintdb/index.php?db=HumanTF:1.0&motif=FLI1_DBD_1 |
| 59 | db/EUKARYO | FLI1_full_1 | | ACCGGAART | 10 | 10 | 1.3e-009 | CENTRIMO | http://floresta.eead.csic.es/footprintdb/index.php?db=HumanTF:1.0&motif=FLI1_full_1 |
| 60 | db/EUKARYO | GABPA_full | | ACCGGAAGT | 10 | 10 | 1.3e-009 | CENTRIMO | http://floresta.eead.csic.es/footprintdb/index.php?db=HumanTF:1.0&motif=GABPA_full |
| 61 | db/JASPAR/J/ | MA0645.1 | ETV6 | MSCGGAAGT | 10 | 10 | 1.3e-009 | CENTRIMO | http://jaspar2018.genereg.net/matrix/MA0645.1 |
| 62 | db/JASPAR/J/ | MA0475.2 | FLI1 | ACCGGAART | 10 | 10 | 1.3e-009 | CENTRIMO | http://jaspar2018.genereg.net/matrix/MA0475.2 |
| 63 | db/JASPAR/J/ | MA0759.1 | ELK3 | ACCGGAAGT | 10 | 10 | 1.3e-009 | CENTRIMO | http://jaspar2018.genereg.net/matrix/MA0759.1 |
| 64 | db/JASPAR/J/ | MA0760.1 | ERF | ACCGGAAGT | 10 | 10 | 1.3e-009 | CENTRIMO | http://jaspar2018.genereg.net/matrix/MA0760.1 |
| 65 | db/JASPAR/J/ | MA0474.2 | ERG | ACCGGAART | 10 | 10 | 1.3e-009 | CENTRIMO | http://jaspar2018.genereg.net/matrix/MA0474.2 |
| 66 | db/JASPAR/J/ | MA0098.3 | ETS1 | ACCGGAART | 10 | 10 | 1.3e-009 | CENTRIMO | http://jaspar2018.genereg.net/matrix/MA0098.3 |
| 67 | db/JASPAR/J/ | MA0761.1 | ETV1 | ACCGGAAGT | 10 | 10 | 1.3e-009 | CENTRIMO | http://jaspar2018.genereg.net/matrix/MA0761.1 |
| 68 | db/JASPAR/J/ | MA0763.1 | ETV3 | ACCGGAAGT | 10 | 10 | 1.3e-009 | CENTRIMO | http://jaspar2018.genereg.net/matrix/MA0763.1 |
| 69 | db/JASPAR/J/ | MA0764.1 | ETV4 | ACCGGAAGT | 10 | 10 | 1.3e-009 | CENTRIMO | http://jaspar2018.genereg.net/matrix/MA0764.1 |
| 70 | db/JASPAR/J/ | MA0765.1 | ETV5 | ACCGGAWGY | 10 | 10 | 1.3e-009 | CENTRIMO | http://jaspar2018.genereg.net/matrix/MA0765.1 |
| 71 | db/JASPAR/J/ | MA0156.2 | FEV | ACCGGAAGT | 10 | 10 | 1.3e-009 | CENTRIMO | http://jaspar2018.genereg.net/matrix/MA0156.2 |
| 72 | db/JASPAR/J/ | MA0028.2 | ELK1 | ACCGGAAGT | 10 | 10 | 1.3e-009 | CENTRIMO | http://jaspar2018.genereg.net/matrix/MA0028.2 |
| 73 | db/EUKARYO | ETV2_DBD | | AACCGGAAA | 11 | 10 | 1.3e-009 | CENTRIMO | http://floresta.eead.csic.es/footprintdb/index.php?db=HumanTF:1.0&motif=ETV2_DBD |
| 74 | db/EUKARYO | SPDEF_DBD_1 | | AMCCGGATG | 11 | 10 | 1.3e-009 | CENTRIMO | http://floresta.eead.csic.es/footprintdb/index.php?db=HumanTF:1.0&motif=SPDEF_DBD_1 |
| 75 | db/EUKARYO | SPDEF_full_1 | | AMCCGGATG | 11 | 10 | 1.3e-009 | CENTRIMO | http://floresta.eead.csic.es/footprintdb/index.php?db=HumanTF:1.0&motif=SPDEF_full_1 |
| 76 | db/JASPAR/J/ | MA0686.1 | SPDEF | AMCCGGATG | 11 | 10 | 1.3e-009 | CENTRIMO | http://jaspar2018.genereg.net/matrix/MA0686.1 |
| 77 | db/JASPAR/J/ | MA0762.1 | ETV2 | AACCGGAAA | 11 | 10 | 1.3e-009 | CENTRIMO | http://jaspar2018.genereg.net/matrix/MA0762.1 |
| 78 | db/MOUSE/ur | UP00013_1 | Gabpa_primar | MNWWACCG | 17 | 10 | 1.3e-009 | CENTRIMO | http://the_brain.bwh.harvard.edu/uniprobe/details2.php?id=00013 |
| 79 | db/EUKARYO | ELF1_DBD | | AACCCGGAA | 12 | 10 | 1.3e-009 | CENTRIMO | http://floresta.eead.csic.es/footprintdb/index.php?db=HumanTF:1.0&motif=ELF1_DBD |
| 80 | db/EUKARYO | ELF1_full | | AACCCGGAA | 12 | 10 | 1.3e-009 | CENTRIMO | http://floresta.eead.csic.es/footprintdb/index.php?db=HumanTF:1.0&motif=ELF1_full |
| 81 | db/EUKARYO | ELF4_full | | AACCCGGAA | 12 | 10 | 1.3e-009 | CENTRIMO | http://floresta.eead.csic.es/footprintdb/index.php?db=HumanTF:1.0&motif=ELF4_full |
| 82 | db/JASPAR/J/ | MA0641.1 | ELF4 | AACCCGGAA | 12 | 10 | 1.3e-009 | CENTRIMO | http://jaspar2018.genereg.net/matrix/MA0641.1 |
| 83 | db/JASPAR/J/ | MA0473.2 | ELF1 | AACCCGGAA | 12 | 10 | 1.3e-009 | CENTRIMO | http://jaspar2018.genereg.net/matrix/MA0473.2 |
| 84 | db/JASPAR/J/ | MA1106.1 | HIF1A | SBACGTGCN | 10 | 10 | 3.4e-009 | CENTRIMO | http://jaspar2018.genereg.net/matrix/MA1106.1 |
| 85 | db/MOUSE/ur | UP00198_1 | Cphx_3484.1 | NTGATBVMA | 14 | 10 | 4.0e-009 | CENTRIMO | http://the_brain.bwh.harvard.edu/uniprobe/details2.php?id=00198 |
| 86 | db/EUKARYO | PAX2_DBD | | SGTCACGCV | 18 | 10 | 5.0e-009 | CENTRIMO | http://floresta.eead.csic.es/footprintdb/index.php?db=HumanTF:1.0&motif=PAX2_DBD |
| 87 | db/JASPAR/J/ | MA1114.1 | PBX3 | NNNTGAGTG | 17 | 10 | 1.8e-008 | CENTRIMO | http://jaspar2018.genereg.net/matrix/MA1114.1 |
| 88 | db/MOUSE/ur | UP00069_2 | Sox1_second | NYRTAATTGT | 15 | 10 | 4.8e-008 | CENTRIMO | http://the_brain.bwh.harvard.edu/uniprobe/details2.php?id=00069 |
| 89 | db/JASPAR/J/ | MA0069.1 | Pax6 | TTCACGCWT | 14 | 10 | 1.1e-007 | CENTRIMO | http://jaspar2018.genereg.net/matrix/MA0069.1 |
| 90 | db/EUKARYO | E2F1_DBD_3 | | WWTTGGCG | 14 | 10 | 4.6e-007 | CENTRIMO | http://floresta.eead.csic.es/footprintdb/index.php?db=HumanTF:1.0&motif=E2F1_DBD_3 |
| 91 | db/JASPAR/J/ | MA0629.1 | Rhox11 | ANKNCGCTG | 17 | 10 | 7.0e-007 | CENTRIMO | http://jaspar2018.genereg.net/matrix/MA0629.1 |
| 92 | db/MOUSE/ur | UP00193_1 | Rhox11_1765 | ANKNCGCTG | 17 | 10 | 7.0e-007 | CENTRIMO | http://the_brain.bwh.harvard.edu/uniprobe/details2.php?id=00193 |
| 93 | DREME | GAGRCAGR | DREME-1 | GAGRCAGR | 8 | 109 | 8.2e-007 | DREME | |
| 94 | db/MOUSE/ur | UP00024_1 | Glis2_primary | YWHNGACC | 16 | 10 | 1.0e-006 | CENTRIMO | http://the_brain.bwh.harvard.edu/uniprobe/details2.php?id=00024 |
| 95 | db/MOUSE/ur | UP00026_1 | Zscan4_prima | DHNATGTGC | 17 | 10 | 1.3e-006 | CENTRIMO | http://the_brain.bwh.harvard.edu/uniprobe/details2.php?id=00026 |
| 96 | db/EUKARYO | ETS1_DBD_2 | | ACCGGAWGY | 18 | 10 | 3.2e-006 | CENTRIMO | http://floresta.eead.csic.es/footprintdb/index.php?db=HumanTF:1.0&motif=ETS1_DBD_2 |
| 97 | db/EUKARYO | NRF1_full | | YGCGCATGC | 12 | 10 | 3.5e-006 | CENTRIMO | http://floresta.eead.csic.es/footprintdb/index.php?db=HumanTF:1.0&motif=NRF1_full |
| 98 | db/JASPAR/J/ | MA1118.1 | SIX1 | GWAACCTGA | 11 | 10 | 4.0e-006 | CENTRIMO | http://jaspar2018.genereg.net/matrix/MA1118.1 |

| | | | | | | | | | | | |
|---|---|---|---|---|---|---|---|---|---|---|---|
| 99 | db/JASPAR/J | MA1119.1 | SIX2 | NNCTGAAAC | 16 | 10 | 4.1e-006 | CENTRIMO | | | http://jaspar2018.genereg.net/matrix/MA1119.1 |
| 100 | db/EUKARYO | RUNX2_DBD_2 | | WRACCGCA | 18 | 10 | 4.4e-006 | CENTRIMO | | | http://floresta.eead.csic.es/footprintdb/index.php?db=HumanTF:1.0&motif=RUNX2_DBD_2 |
| 101 | DREME | AAAYAWA | DREME-4 | AAAYAWA | 7 | 217 | 1.2e-005 | DREME | db/EUKARYOTE/joln | **Foxc1_DBD_1** | http://floresta.eead.csic.es/footprintdb/index.php?db=HumanTF:1.0&motif=Foxc1_DBD_1 |
| 102 | db/MOUSE/ur | UP00043_2 | Bcl6b_second | HNNBCCGC | 16 | 10 | 1.7e-005 | CENTRIMO | | | http://the_brain.bwh.harvard.edu/uniprobe/details2.php?id=00043 |
| 103 | db/MOUSE/ur | UP00033_2 | Zfp410_secon | NYNYYCCGC | 17 | 10 | 1.7e-005 | CENTRIMO | | | http://the_brain.bwh.harvard.edu/uniprobe/details2.php?id=00033 |
| 104 | db/JASPAR/J | MA0471.1 | E2F6 | RGGCGGGA | 11 | 10 | 1.9e-005 | CENTRIMO | | | http://jaspar2018.genereg.net/matrix/MA0471.1 |
| 105 | db/JASPAR/J | MA0470.1 | E2F4 | GGGCGGGA | 11 | 10 | 1.9e-005 | CENTRIMO | | | http://jaspar2018.genereg.net/matrix/MA0470.1 |
| 106 | db/JASPAR/J | MA1122.1 | TFDP1 | VSGCGGGA | 11 | 10 | 1.9e-005 | CENTRIMO | | | http://jaspar2018.genereg.net/matrix/MA1122.1 |
| 107 | db/JASPAR/J | MA0071.1 | RORA | AWMWAGGT | 10 | 10 | 2.1e-005 | CENTRIMO | | | http://jaspar2018.genereg.net/matrix/MA0071.1 |
| 108 | db/EUKARYO | TFAP2A_DBD_2 | | HGCCYSAG | 11 | 10 | 2.9e-005 | CENTRIMO | | | http://floresta.eead.csic.es/footprintdb/index.php?db=HumanTF:1.0&motif=TFAP2A_DBD_2 |
| 109 | db/EUKARYO | TFAP2A_DBD_4 | | HGCCTSAG | 11 | 10 | 2.9e-005 | CENTRIMO | | | http://floresta.eead.csic.es/footprintdb/index.php?db=HumanTF:1.0&motif=TFAP2A_DBD_4 |
| 110 | db/EUKARYO | TFAP2C_DBD_2 | | HGCCTSAG | 11 | 10 | 2.9e-005 | CENTRIMO | | | http://floresta.eead.csic.es/footprintdb/index.php?db=HumanTF:1.0&motif=TFAP2C_DBD_2 |
| 111 | db/EUKARYO | TFAP2C_full_3 | | HGCCTSAG | 11 | 10 | 2.9e-005 | CENTRIMO | | | http://floresta.eead.csic.es/footprintdb/index.php?db=HumanTF:1.0&motif=TFAP2C_full_3 |
| 112 | db/EUKARYO | Tcfap2a_DBD_2 | | HGCCYSAG | 11 | 10 | 2.9e-005 | CENTRIMO | | | http://floresta.eead.csic.es/footprintdb/index.php?db=HumanTF:1.0&motif=Tcfap2a_DBD_2 |
| 113 | db/JASPAR/J | MA0003.3 | TFAP2A | HGCCYSAG | 11 | 10 | 2.9e-005 | CENTRIMO | | | http://jaspar2018.genereg.net/matrix/MA0003.3 |
| 114 | db/JASPAR/J | MA0814.1 | TFAP2C(var.2 | HGCCTSAG | 11 | 10 | 2.9e-005 | CENTRIMO | | | http://jaspar2018.genereg.net/matrix/MA0814.1 |
| 115 | db/MOUSE/ur | UP00015_2 | Ehf_secondar | WWVDABTT | 16 | 10 | 3.8e-005 | CENTRIMO | | | http://the_brain.bwh.harvard.edu/uniprobe/details2.php?id=00015 |
| 116 | db/JASPAR/J | MA0137.3 | STAT1 | TTTCYRGGA | 11 | 10 | 4.3e-005 | CENTRIMO | | | http://jaspar2018.genereg.net/matrix/MA0137.3 |
| 117 | db/EUKARYO | TFCP2_full_2 | | ACCGGTTYA | 16 | 10 | 4.7e-005 | CENTRIMO | | | http://floresta.eead.csic.es/footprintdb/index.php?db=HumanTF:1.0&motif=TFCP2_full_2 |
| 118 | db/MOUSE/ur | UP00085_2 | Sfpi1_second | HHAWTTMV | 14 | 10 | 4.9e-005 | CENTRIMO | | | http://the_brain.bwh.harvard.edu/uniprobe/details2.php?id=00085 |
| 119 | db/MOUSE/ur | UP00075_1 | Sox15_primar | WNNDGAAC | 17 | 10 | 6.6e-005 | CENTRIMO | | | http://the_brain.bwh.harvard.edu/uniprobe/details2.php?id=00075 |
| 120 | db/EUKARYO | GLI2_DBD_2 | | GCGACCAC | 12 | 10 | 6.6e-005 | CENTRIMO | | | http://floresta.eead.csic.es/footprintdb/index.php?db=HumanTF:1.0&motif=GLI2_DBD_2 |
| 121 | db/JASPAR/J | MA0734.1 | GLI2 | GCGACCAC | 12 | 10 | 6.6e-005 | CENTRIMO | | | http://jaspar2018.genereg.net/matrix/MA0734.1 |
| 122 | db/MOUSE/ur | UP00071_2 | Sox21_secon | NVYSWATTG | 17 | 10 | 6.7e-005 | CENTRIMO | | | http://the_brain.bwh.harvard.edu/uniprobe/details2.php?id=00071 |
| 123 | db/EUKARYO | TEAD1_full_2 | | RCATTCCDN | 17 | 10 | 7.1e-005 | CENTRIMO | | | http://floresta.eead.csic.es/footprintdb/index.php?db=HumanTF:1.0&motif=TEAD1_full_2 |
| 124 | db/EUKARYO | POU2F3_DBD_2 | | ATGMATATK | 12 | 10 | 8.5e-005 | CENTRIMO | | | http://floresta.eead.csic.es/footprintdb/index.php?db=HumanTF:1.0&motif=POU2F3_DBD_2 |
| 125 | db/EUKARYO | POU5F1P1_DBD_2 | | ATGMATATK | 12 | 10 | 8.5e-005 | CENTRIMO | | | http://floresta.eead.csic.es/footprintdb/index.php?db=HumanTF:1.0&motif=POU5F1P1_DBD_2 |
| 126 | db/EUKARYO | POU2F1_DBD_2 | | HATGMATAT | 14 | 10 | 8.5e-005 | CENTRIMO | | | http://floresta.eead.csic.es/footprintdb/index.php?db=HumanTF:1.0&motif=POU2F1_DBD_2 |
| 127 | db/EUKARYO | POU2F2_DBD_2 | | HWTRMATAT | 14 | 10 | 8.5e-005 | CENTRIMO | | | http://floresta.eead.csic.es/footprintdb/index.php?db=HumanTF:1.0&motif=POU2F2_DBD_2 |
| 128 | db/EUKARYO | Pou2f2_DBD_1 | | YWTGAATAT | 14 | 10 | 8.5e-005 | CENTRIMO | | | http://floresta.eead.csic.es/footprintdb/index.php?db=HumanTF:1.0&motif=Pou2f2_DBD_1 |
| 129 | db/MOUSE/ur | UP00093_2 | Klf7_secondar | VRNBATACG | 17 | 10 | 1.3e-004 | CENTRIMO | | | http://the_brain.bwh.harvard.edu/uniprobe/details2.php?id=00093 |
| 130 | db/EUKARYO | HSFY2_DBD_2 | | TTTCGAACG | 9 | 10 | 2.0e-004 | CENTRIMO | | | http://floresta.eead.csic.es/footprintdb/index.php?db=HumanTF:1.0&motif=HSFY2_DBD_2 |
| 131 | DREME | TACATR | DREME-2 | TACATR | 6 | 180 | 2.0e-004 | DREME | db/MOUSE/uniprobe_ | **UP00082_1 (Zfp187_** | http://the_brain.bwh.harvard.edu/uniprobe/details2.php?id=00082 |
| 132 | db/EUKARYO | HSFY2_DBD_1 | | TTCGAAHV | 15 | 10 | 2.1e-004 | CENTRIMO | | | http://floresta.eead.csic.es/footprintdb/index.php?db=HumanTF:1.0&motif=HSFY2_DBD_1 |
| 133 | db/EUKARYO | HSFY2_DBD_3 | | TTCGAAHS | 15 | 10 | 2.1e-004 | CENTRIMO | | | http://floresta.eead.csic.es/footprintdb/index.php?db=HumanTF:1.0&motif=HSFY2_DBD_3 |
| 134 | db/MOUSE/ur | UP00066_2 | Hnf4a_second | KKYHAAAGT | 16 | 10 | 2.2e-004 | CENTRIMO | | | http://the_brain.bwh.harvard.edu/uniprobe/details2.php?id=00066 |
| 135 | db/JASPAR/J | MA0604.1 | Atf1 | RTGACGTA | 8 | 10 | 2.6e-004 | CENTRIMO | | | http://jaspar2018.genereg.net/matrix/MA0604.1 |
| 136 | db/JASPAR/J | MA0609.1 | Crem | NATGACGTM | 10 | 10 | 2.6e-004 | CENTRIMO | | | http://jaspar2018.genereg.net/matrix/MA0609.1 |
| 137 | db/JASPAR/J | MA0608.1 | Creb3l2 | GCCACGTG | 9 | 10 | 2.6e-004 | CENTRIMO | | | http://jaspar2018.genereg.net/matrix/MA0608.1 |
| 138 | db/EUKARYO | MLXIPL_full | | RTCACGTGA | 10 | 10 | 2.6e-004 | CENTRIMO | | | http://floresta.eead.csic.es/footprintdb/index.php?db=HumanTF:1.0&motif=MLXIPL_full |
| 139 | db/JASPAR/J | MA0664.1 | MLXIPL | RTCACGTGA | 10 | 10 | 2.6e-004 | CENTRIMO | | | http://jaspar2018.genereg.net/matrix/MA0664.1 |
| 140 | db/EUKARYO | CREB3_full_2 | | VTGCCACGT | 14 | 10 | 2.6e-004 | CENTRIMO | | | http://floresta.eead.csic.es/footprintdb/index.php?db=HumanTF:1.0&motif=CREB3_full_2 |
| 141 | db/JASPAR/J | MA0638.1 | CREB3 | VTGCCACGT | 14 | 10 | 2.6e-004 | CENTRIMO | | | http://jaspar2018.genereg.net/matrix/MA0638.1 |
| 142 | db/EUKARYO | VSX1_full | | DGCYAATT | 11 | 10 | 2.6e-004 | CENTRIMO | | | http://floresta.eead.csic.es/footprintdb/index.php?db=HumanTF:1.0&motif=VSX1_full |
| 143 | db/MOUSE/ur | UP00256_2 | Lhx6_3432.1 | WMNVCTAAT | 17 | 10 | 2.6e-004 | CENTRIMO | | | http://the_brain.bwh.harvard.edu/uniprobe/details2.php?id=00256 |
| 144 | db/EUKARYO | BARX1_DBD_2 | | VCMATTAV | 8 | 10 | 2.6e-004 | CENTRIMO | | | http://floresta.eead.csic.es/footprintdb/index.php?db=HumanTF:1.0&motif=BARX1_DBD_2 |
| 145 | db/EUKARYO | BSX_DBD | | SYVATTAV | 8 | 10 | 2.6e-004 | CENTRIMO | | | http://floresta.eead.csic.es/footprintdb/index.php?db=HumanTF:1.0&motif=BSX_DBD |
| 146 | db/EUKARYO | DLX2_DBD | | NYAATTAN | 8 | 10 | 2.6e-004 | CENTRIMO | | | http://floresta.eead.csic.es/footprintdb/index.php?db=HumanTF:1.0&motif=DLX2_DBD |
| 147 | db/EUKARYO | EN1_full_1 | | SYAATTAV | 8 | 10 | 2.6e-004 | CENTRIMO | | | http://floresta.eead.csic.es/footprintdb/index.php?db=HumanTF:1.0&motif=EN1_full_1 |
| 148 | db/EUKARYO | ISX_DBD_2 | | CYAATTAV | 8 | 10 | 2.6e-004 | CENTRIMO | | | http://floresta.eead.csic.es/footprintdb/index.php?db=HumanTF:1.0&motif=ISX_DBD_2 |
| 149 | db/EUKARYO | ISX_full | | YYAATTAR | 8 | 10 | 2.6e-004 | CENTRIMO | | | http://floresta.eead.csic.es/footprintdb/index.php?db=HumanTF:1.0&motif=ISX_full |
| 150 | db/EUKARYO | LHX9_DBD_1 | | BYAATTAR | 8 | 10 | 2.6e-004 | CENTRIMO | | | http://floresta.eead.csic.es/footprintdb/index.php?db=HumanTF:1.0&motif=LHX9_DBD_1 |
| 151 | db/EUKARYO | MSX1_DBD_2 | | SCAATTAV | 8 | 10 | 2.6e-004 | CENTRIMO | | | http://floresta.eead.csic.es/footprintdb/index.php?db=HumanTF:1.0&motif=MSX1_DBD_2 |
| 152 | db/EUKARYO | MSX1_full | | SCAATTAV | 8 | 10 | 2.6e-004 | CENTRIMO | | | http://floresta.eead.csic.es/footprintdb/index.php?db=HumanTF:1.0&motif=MSX1_full |
| 153 | db/EUKARYO | MSX2_DBD_2 | | CCAATTAV | 8 | 10 | 2.6e-004 | CENTRIMO | | | http://floresta.eead.csic.es/footprintdb/index.php?db=HumanTF:1.0&motif=MSX2_DBD_2 |
| 154 | db/EUKARYO | Msx3_DBD_2 | | SCAATTAN | 8 | 10 | 2.6e-004 | CENTRIMO | | | http://floresta.eead.csic.es/footprintdb/index.php?db=HumanTF:1.0&motif=Msx3_DBD_2 |
| 155 | db/EUKARYO | PRRX1_DBD | | YYAATTAR | 8 | 10 | 2.6e-004 | CENTRIMO | | | http://floresta.eead.csic.es/footprintdb/index.php?db=HumanTF:1.0&motif=PRRX1_DBD |
| 156 | db/EUKARYO | PRRX1_full_1 | | YYAATTAN | 8 | 10 | 2.6e-004 | CENTRIMO | | | http://floresta.eead.csic.es/footprintdb/index.php?db=HumanTF:1.0&motif=PRRX1_full_1 |
| 157 | db/EUKARYO | PRRX2_full | | YYAATTAN | 8 | 10 | 2.6e-004 | CENTRIMO | | | http://floresta.eead.csic.es/footprintdb/index.php?db=HumanTF:1.0&motif=PRRX2_full |
| 158 | db/EUKARYO | RAXL1_DBD | | YYAATTAV | 8 | 10 | 2.6e-004 | CENTRIMO | | | http://floresta.eead.csic.es/footprintdb/index.php?db=HumanTF:1.0&motif=RAXL1_DBD |
| 159 | db/EUKARYO | SHOX2_DBD | | YYAATTAR | 8 | 10 | 2.6e-004 | CENTRIMO | | | http://floresta.eead.csic.es/footprintdb/index.php?db=HumanTF:1.0&motif=SHOX2_DBD |

| | | | | | | | | | | |
|---|---|---|---|---|---|---|---|---|---|---|
| 160 | db/EUKARYO | SHOX_DBD | | YYAATTAR | 8 | 10 | 2.6e-004 | CENTRIMO | | http://floresta.eead.csic.es/footprintdb/index.php?db=HumanTF:1.0&motif=SHOX_DBD |
| 161 | db/EUKARYO | Shox2_DBD | | YYAATTAR | 8 | 10 | 2.6e-004 | CENTRIMO | | http://floresta.eead.csic.es/footprintdb/index.php?db=HumanTF:1.0&motif=Shox2_DBD |
| 162 | db/EUKARYO | Vsx1_DBD | | YYAATTAN | 8 | 10 | 2.6e-004 | CENTRIMO | | http://floresta.eead.csic.es/footprintdb/index.php?db=HumanTF:1.0&motif=Vsx1_DBD |
| 163 | db/JASPAR/J/ | MA0630.1 | SHOX | HTAATTRR | 8 | 10 | 2.6e-004 | CENTRIMO | | http://jaspar2018.genereg.net/matrix/MA0630.1 |
| 164 | db/JASPAR/J/ | MA0027.2 | EN1 | SYAATTAV | 8 | 10 | 2.6e-004 | CENTRIMO | | http://jaspar2018.genereg.net/matrix/MA0027.2 |
| 165 | db/JASPAR/J/ | MA0654.1 | ISX | YYAATTAR | 8 | 10 | 2.6e-004 | CENTRIMO | | http://jaspar2018.genereg.net/matrix/MA0654.1 |
| 166 | db/JASPAR/J/ | MA0666.1 | MSX1 | SCAATTAV | 8 | 10 | 2.6e-004 | CENTRIMO | | http://jaspar2018.genereg.net/matrix/MA0666.1 |
| 167 | db/JASPAR/J/ | MA0075.2 | Prrx2 | YYAATTAN | 8 | 10 | 2.6e-004 | CENTRIMO | | http://jaspar2018.genereg.net/matrix/MA0075.2 |
| 168 | db/JASPAR/J/ | MA0701.1 | LHX9 | BYAATTAR | 8 | 10 | 2.6e-004 | CENTRIMO | | http://jaspar2018.genereg.net/matrix/MA0701.1 |
| 169 | db/JASPAR/J/ | MA0708.1 | MSX2 | CCAATTAV | 8 | 10 | 2.6e-004 | CENTRIMO | | http://jaspar2018.genereg.net/matrix/MA0708.1 |
| 170 | db/JASPAR/J/ | MA0709.1 | Msx3 | SCAATTAN | 8 | 10 | 2.6e-004 | CENTRIMO | | http://jaspar2018.genereg.net/matrix/MA0709.1 |
| 171 | db/JASPAR/J/ | MA0716.1 | PRRX1 | YYAATTAR | 8 | 10 | 2.6e-004 | CENTRIMO | | http://jaspar2018.genereg.net/matrix/MA0716.1 |
| 172 | db/JASPAR/J/ | MA0717.1 | RAX2 | YYAATTAV | 8 | 10 | 2.6e-004 | CENTRIMO | | http://jaspar2018.genereg.net/matrix/MA0717.1 |
| 173 | db/JASPAR/J/ | MA0720.1 | Shox2 | YYAATTAR | 8 | 10 | 2.6e-004 | CENTRIMO | | http://jaspar2018.genereg.net/matrix/MA0720.1 |
| 174 | db/JASPAR/J/ | MA0875.1 | BARX1 | VCMATTAV | 8 | 10 | 2.6e-004 | CENTRIMO | | http://jaspar2018.genereg.net/matrix/MA0875.1 |
| 175 | db/JASPAR/J/ | MA0876.1 | BSX | SYVATTAV | 8 | 10 | 2.6e-004 | CENTRIMO | | http://jaspar2018.genereg.net/matrix/MA0876.1 |
| 176 | db/EUKARYO | ALX3_DBD | | RCYAATTAVV | 10 | 10 | 2.7e-004 | CENTRIMO | | http://floresta.eead.csic.es/footprintdb/index.php?db=HumanTF:1.0&motif=ALX3_DBD |
| 177 | db/EUKARYO | ALX3_full_1 | | YYYAATTANV | 10 | 10 | 2.7e-004 | CENTRIMO | | http://floresta.eead.csic.es/footprintdb/index.php?db=HumanTF:1.0&motif=ALX3_full_1 |
| 178 | db/EUKARYO | Alx1_DBD_1 | | YYTAATTAVV | 10 | 10 | 2.7e-004 | CENTRIMO | | http://floresta.eead.csic.es/footprintdb/index.php?db=HumanTF:1.0&motif=Alx1_DBD_1 |
| 179 | db/EUKARYO | EMX1_DBD_1 | | NBTAATTAVS | 10 | 10 | 2.7e-004 | CENTRIMO | | http://floresta.eead.csic.es/footprintdb/index.php?db=HumanTF:1.0&motif=EMX1_DBD_1 |
| 180 | db/EUKARYO | EN1_DBD_1 | | VSYAATTAVB | 10 | 10 | 2.7e-004 | CENTRIMO | | http://floresta.eead.csic.es/footprintdb/index.php?db=HumanTF:1.0&motif=EN1_DBD_1 |
| 181 | db/EUKARYO | EN2_full | | VBYAATTAVS | 10 | 10 | 2.7e-004 | CENTRIMO | | http://floresta.eead.csic.es/footprintdb/index.php?db=HumanTF:1.0&motif=EN2_full |
| 182 | db/EUKARYO | ESX1_DBD | | NYYAATTANC | 10 | 10 | 2.7e-004 | CENTRIMO | | http://floresta.eead.csic.es/footprintdb/index.php?db=HumanTF:1.0&motif=ESX1_DBD |
| 183 | db/EUKARYO | ESX1_full | | NYYAATTAVM | 10 | 10 | 2.7e-004 | CENTRIMO | | http://floresta.eead.csic.es/footprintdb/index.php?db=HumanTF:1.0&motif=ESX1_full |
| 184 | db/EUKARYO | GBX1_DBD | | VCYAATTASY | 10 | 10 | 2.7e-004 | CENTRIMO | | http://floresta.eead.csic.es/footprintdb/index.php?db=HumanTF:1.0&motif=GBX1_DBD |
| 185 | db/EUKARYO | GBX2_DBD_1 | | RSYAATTARN | 10 | 10 | 2.7e-004 | CENTRIMO | | http://floresta.eead.csic.es/footprintdb/index.php?db=HumanTF:1.0&motif=GBX2_DBD_1 |
| 186 | db/EUKARYO | GBX2_full | | NSYAATTANV | 10 | 10 | 2.7e-004 | CENTRIMO | | http://floresta.eead.csic.es/footprintdb/index.php?db=HumanTF:1.0&motif=GBX2_full |
| 187 | db/EUKARYO | Gbx1_DBD | | NYYAATTAVN | 10 | 10 | 2.7e-004 | CENTRIMO | | http://floresta.eead.csic.es/footprintdb/index.php?db=HumanTF:1.0&motif=Gbx1_DBD |
| 188 | db/EUKARYO | HESX1_DBD_1 | | GHTAATTRRI | 10 | 10 | 2.7e-004 | CENTRIMO | | http://floresta.eead.csic.es/footprintdb/index.php?db=HumanTF:1.0&motif=HESX1_DBD_1 |
| 189 | db/EUKARYO | LBX2_DBD_2 | | NCYAATTARN | 10 | 10 | 2.7e-004 | CENTRIMO | | http://floresta.eead.csic.es/footprintdb/index.php?db=HumanTF:1.0&motif=LBX2_DBD_2 |
| 190 | db/EUKARYO | LHX2_DBD_1 | | VCTAATTARB | 10 | 10 | 2.7e-004 | CENTRIMO | | http://floresta.eead.csic.es/footprintdb/index.php?db=HumanTF:1.0&motif=LHX2_DBD_1 |
| 191 | db/EUKARYO | MIXL1_full | | NBYAATTAVN | 10 | 10 | 2.7e-004 | CENTRIMO | | http://floresta.eead.csic.es/footprintdb/index.php?db=HumanTF:1.0&motif=MIXL1_full |
| 192 | db/JASPAR/J/ | MA0634.1 | ALX3 | YYYAATTANV | 10 | 10 | 2.7e-004 | CENTRIMO | | http://jaspar2018.genereg.net/matrix/MA0634.1 |
| 193 | db/JASPAR/J/ | MA0642.1 | EN2 | VBYAATTAVS | 10 | 10 | 2.7e-004 | CENTRIMO | | http://jaspar2018.genereg.net/matrix/MA0642.1 |
| 194 | db/JASPAR/J/ | MA0644.1 | ESX1 | NYYAATTAVM | 10 | 10 | 2.7e-004 | CENTRIMO | | http://jaspar2018.genereg.net/matrix/MA0644.1 |
| 195 | db/JASPAR/J/ | MA0662.1 | MIXL1 | NBYAATTAVN | 10 | 10 | 2.7e-004 | CENTRIMO | | http://jaspar2018.genereg.net/matrix/MA0662.1 |
| 196 | db/JASPAR/J/ | MA0699.1 | LBX2 | NCYAATTARN | 10 | 10 | 2.7e-004 | CENTRIMO | | http://jaspar2018.genereg.net/matrix/MA0699.1 |
| 197 | db/JASPAR/J/ | MA0700.1 | LHX2 | VCTAATTARB | 10 | 10 | 2.7e-004 | CENTRIMO | | http://jaspar2018.genereg.net/matrix/MA0700.1 |
| 198 | db/JASPAR/J/ | MA0889.1 | GBX1 | VCYAATTASY | 10 | 10 | 2.7e-004 | CENTRIMO | | http://jaspar2018.genereg.net/matrix/MA0889.1 |
| 199 | db/JASPAR/J/ | MA0890.1 | GBX2 | RSYAATTARN | 10 | 10 | 2.7e-004 | CENTRIMO | | http://jaspar2018.genereg.net/matrix/MA0890.1 |
| 200 | db/JASPAR/J/ | MA0894.1 | HESX1 | GHTAATTRRI | 10 | 10 | 2.7e-004 | CENTRIMO | | http://jaspar2018.genereg.net/matrix/MA0894.1 |
| 201 | db/EUKARYO | XBP1_DBD_2 | | WDKGMCACI | 14 | 10 | 2.7e-004 | CENTRIMO | | http://floresta.eead.csic.es/footprintdb/index.php?db=HumanTF:1.0&motif=XBP1_DBD_2 |
| 202 | db/JASPAR/J/ | MA0844.1 | XBP1 | WDKGMCACI | 14 | 10 | 2.7e-004 | CENTRIMO | | http://jaspar2018.genereg.net/matrix/MA0844.1 |
| 203 | db/MOUSE/ur | UP00243_1 | Isx_3445.1 | HNNVCYAAT | 16 | 10 | 2.7e-004 | CENTRIMO | | http://the_brain.bwh.harvard.edu/uniprobe/details2.php?id=00243 |
| 204 | DREME | RAACTCA | DREME-3 | RAACTCA | 7 | 102 | 3.1e-004 | DREME | db/JASPAR/JASPAR **MA0693.2 (VDR)** | http://jaspar2018.genereg.net/matrix/MA0693.2 |
| 205 | db/MOUSE/ur | UP00048_2 | Rara_seconda | MKMGYSGGI | 16 | 10 | 4.0e-004 | CENTRIMO | | http://the_brain.bwh.harvard.edu/uniprobe/details2.php?id=00048 |
| 206 | db/EUKARYO | ZNF524_full_1 | | ACCCTYGAA | 12 | 10 | 4.2e-004 | CENTRIMO | | http://floresta.eead.csic.es/footprintdb/index.php?db=HumanTF:1.0&motif=ZNF524_full_1 |
| 207 | db/MOUSE/ur | UP00220_1 | Nkx1-1_3856. | WGMVCYAAT | 17 | 10 | 4.3e-004 | CENTRIMO | | http://the_brain.bwh.harvard.edu/uniprobe/details2.php?id=00220 |
| 208 | db/EUKARYO | GLI2_DBD_1 | | GACCACCCA | 14 | 10 | 4.4e-004 | CENTRIMO | | http://floresta.eead.csic.es/footprintdb/index.php?db=HumanTF:1.0&motif=GLI2_DBD_1 |
| 209 | db/MOUSE/ur | UP00096_2 | Sox13_second | DWDHTGGG | 17 | 10 | 4.6e-004 | CENTRIMO | | http://the_brain.bwh.harvard.edu/uniprobe/details2.php?id=00096 |
| 210 | db/JASPAR/J/ | MA0883.1 | Dmbx1 | WDRWNMGG | 17 | 10 | 5.0e-004 | CENTRIMO | | http://jaspar2018.genereg.net/matrix/MA0883.1 |
| 211 | db/MOUSE/ur | UP00111_1 | Dmbx1_2277. | WDRWHMGG | 17 | 10 | 5.0e-004 | CENTRIMO | | http://the_brain.bwh.harvard.edu/uniprobe/details2.php?id=00111 |
| 212 | db/EUKARYO | RHOXF1_DBD_2 | | NTRATCCN | 8 | 10 | 5.0e-004 | CENTRIMO | | http://floresta.eead.csic.es/footprintdb/index.php?db=HumanTF:1.0&motif=RHOXF1_DBD_2 |
| 213 | db/JASPAR/J/ | MA0719.1 | RHOXF1 | NTRATCCN | 8 | 10 | 5.0e-004 | CENTRIMO | | http://jaspar2018.genereg.net/matrix/MA0719.1 |
| 214 | db/EUKARYO | DPRX_DBD_2 | | VRGATAATC | 11 | 10 | 5.1e-004 | CENTRIMO | | http://floresta.eead.csic.es/footprintdb/index.php?db=HumanTF:1.0&motif=DPRX_DBD_2 |
| 215 | db/EUKARYO | POU2F1_DBD_1 | | AWTATGCWA | 12 | 10 | 5.4e-004 | CENTRIMO | | http://floresta.eead.csic.es/footprintdb/index.php?db=HumanTF:1.0&motif=POU2F1_DBD_1 |
| 216 | db/JASPAR/J/ | MA0785.1 | POU2F1 | AWTATGCWA | 12 | 10 | 5.4e-004 | CENTRIMO | | http://jaspar2018.genereg.net/matrix/MA0785.1 |
| 217 | db/EUKARYO | POU2F3_DBD_1 | | TATGCWAAT | 9 | 10 | 5.4e-004 | CENTRIMO | | http://floresta.eead.csic.es/footprintdb/index.php?db=HumanTF:1.0&motif=POU2F3_DBD_1 |
| 218 | db/EUKARYO | POU3F4_DBD_1 | | TATGCWAAT | 9 | 10 | 5.4e-004 | CENTRIMO | | http://floresta.eead.csic.es/footprintdb/index.php?db=HumanTF:1.0&motif=POU3F4_DBD_1 |
| 219 | db/EUKARYO | POU5F1P1_DBD_1 | | TATGCWAAT | 9 | 10 | 5.4e-004 | CENTRIMO | | http://floresta.eead.csic.es/footprintdb/index.php?db=HumanTF:1.0&motif=POU5F1P1_DBD_1 |
| 220 | db/EUKARYO | Pou2f2_DBD_2 | | TATGCAAAT | 9 | 10 | 5.4e-004 | CENTRIMO | | http://floresta.eead.csic.es/footprintdb/index.php?db=HumanTF:1.0&motif=Pou2f2_DBD_2 |

| | | | | | | | | | | | |
|---|---|---|---|---|---|---|---|---|---|---|---|
| 221 | db/JASPAR/J | MA0789.1 | POU3F4 | TATGCWAAT | 9 | 10 | 5.4e-004 | CENTRIMO | | | http://jaspar2018.genereg.net/matrix/MA0789.1 |
| 222 | db/JASPAR/J | MA0792.1 | POU5F1B | TATGCWAAT | 9 | 10 | 5.4e-004 | CENTRIMO | | | http://jaspar2018.genereg.net/matrix/MA0792.1 |
| 223 | db/EUKARYO | POU2F2_DBD_1 | | DTATGCWAA | 11 | 10 | 5.5e-004 | CENTRIMO | | | http://floresta.eead.csic.es/footprintdb/index.php?db=HumanTF:1.0&motif=POU2F2_DBD_1 |
| 224 | db/JASPAR/J | MA1115.1 | POU5F1 | WHATGCAAA | 11 | 10 | 5.5e-004 | CENTRIMO | | | http://jaspar2018.genereg.net/matrix/MA1115.1 |
| 225 | db/EUKARYO | POU3F3_DBD_1 | | WWTATGCW | 13 | 10 | 5.5e-004 | CENTRIMO | | | http://floresta.eead.csic.es/footprintdb/index.php?db=HumanTF:1.0&motif=POU3F3_DBD_1 |
| 226 | db/JASPAR/J | MA0788.1 | POU3F3 | WWTATGCW | 13 | 10 | 5.5e-004 | CENTRIMO | | | http://jaspar2018.genereg.net/matrix/MA0788.1 |
| 227 | db/EUKARYO | POU3F1_DBD_1 | | WTATGCWA | 12 | 10 | 5.5e-004 | CENTRIMO | | | http://floresta.eead.csic.es/footprintdb/index.php?db=HumanTF:1.0&motif=POU3F1_DBD_1 |
| 228 | db/EUKARYO | POU3F2_DBD_2 | | WTATGCWA | 12 | 10 | 5.5e-004 | CENTRIMO | | | http://floresta.eead.csic.es/footprintdb/index.php?db=HumanTF:1.0&motif=POU3F2_DBD_2 |
| 229 | db/JASPAR/J | MA0786.1 | POU3F1 | WTATGCWA | 12 | 10 | 5.5e-004 | CENTRIMO | | | http://jaspar2018.genereg.net/matrix/MA0786.1 |
| 230 | db/JASPAR/J | MA0787.1 | POU3F2 | WTATGCWA | 12 | 10 | 5.5e-004 | CENTRIMO | | | http://jaspar2018.genereg.net/matrix/MA0787.1 |
| 231 | db/EUKARYO | POU1F1_DBD_2 | | AWTATGCWA | 14 | 10 | 5.5e-004 | CENTRIMO | | | http://floresta.eead.csic.es/footprintdb/index.php?db=HumanTF:1.0&motif=POU1F1_DBD_2 |
| 232 | db/JASPAR/J | MA0784.1 | POU1F1 | AWTATGCWA | 14 | 10 | 5.5e-004 | CENTRIMO | | | http://jaspar2018.genereg.net/matrix/MA0784.1 |
| 233 | db/JASPAR/J | MA0627.1 | Pou2f3 | THKTATGCA | 16 | 10 | 5.5e-004 | CENTRIMO | | | http://jaspar2018.genereg.net/matrix/MA0627.1 |
| 234 | db/MOUSE/ur | UP00191_1 | Pou2f2_3748. | BHDTATGCA | 16 | 10 | 5.5e-004 | CENTRIMO | | | http://the_brain.bwh.harvard.edu/uniprobe/details2.php?id=00191 |
| 235 | db/MOUSE/ur | UP00179_1 | Pou2f3_3986. | THKTATGCA | 16 | 10 | 5.5e-004 | CENTRIMO | | | http://the_brain.bwh.harvard.edu/uniprobe/details2.php?id=00179 |
| 236 | db/JASPAR/J | MA0507.1 | POU2F2 | YDNATTTGC | 13 | 10 | 5.6e-004 | CENTRIMO | | | http://jaspar2018.genereg.net/matrix/MA0507.1 |
| 237 | db/EUKARYO | NFATC1_full_1 | | AATGGAAAV | 20 | 10 | 6.1e-004 | CENTRIMO | | | http://floresta.eead.csic.es/footprintdb/index.php?db=HumanTF:1.0&motif=NFATC1_full_1 |
| 238 | db/EUKARYO | HSF1_DBD | | TTCTAGAAY | 13 | 10 | 6.6e-004 | CENTRIMO | | | http://floresta.eead.csic.es/footprintdb/index.php?db=HumanTF:1.0&motif=HSF1_DBD |
| 239 | db/EUKARYO | HSF1_full | | TTCTAGAAY | 13 | 10 | 6.6e-004 | CENTRIMO | | | http://floresta.eead.csic.es/footprintdb/index.php?db=HumanTF:1.0&motif=HSF1_full |
| 240 | db/EUKARYO | HSF2_DBD | | TTCTAGAAY | 13 | 10 | 6.6e-004 | CENTRIMO | | | http://floresta.eead.csic.es/footprintdb/index.php?db=HumanTF:1.0&motif=HSF2_DBD |
| 241 | db/EUKARYO | HSF4_DBD | | TTCTAGAAB | 13 | 10 | 6.6e-004 | CENTRIMO | | | http://floresta.eead.csic.es/footprintdb/index.php?db=HumanTF:1.0&motif=HSF4_DBD |
| 242 | db/JASPAR/J | MA0486.2 | HSF1 | TTCTAGAAY | 13 | 10 | 6.6e-004 | CENTRIMO | | | http://jaspar2018.genereg.net/matrix/MA0486.2 |
| 243 | db/JASPAR/J | MA0770.1 | HSF2 | TTCTAGAAY | 13 | 10 | 6.6e-004 | CENTRIMO | | | http://jaspar2018.genereg.net/matrix/MA0770.1 |
| 244 | db/JASPAR/J | MA0771.1 | HSF4 | TTCTAGAAB | 13 | 10 | 6.6e-004 | CENTRIMO | | | http://jaspar2018.genereg.net/matrix/MA0771.1 |
| 245 | db/EUKARYO | ETV6_full_1 | | CCGGAASC | 15 | 10 | 6.8e-004 | CENTRIMO | | | http://floresta.eead.csic.es/footprintdb/index.php?db=HumanTF:1.0&motif=ETV6_full_1 |
| 246 | DREME | GGATTAAA | DREME-5 | GGATTAAA | 8 | 29 | 8.7e-004 | DREME | db/EUKARYOTE/joln | **PITX1_full_1** | http://floresta.eead.csic.es/footprintdb/index.php?db=HumanTF:1.0&motif=PITX1_full_1 |
| 247 | DREME | GAGCCAYC | DREME-6 | GAGCCAYC | 8 | 43 | 1.1e-003 | DREME | db/JASPAR/JASPAR | **MA0099.3 (FOS::JUN** | http://jaspar2018.genereg.net/matrix/MA0099.3 |
| 248 | db/MOUSE/ur | UP00076_1 | Rfxdc2_prima | NYVCHTAGC | 15 | 10 | 1.3e-003 | CENTRIMO | | | http://the_brain.bwh.harvard.edu/uniprobe/details2.php?id=00076 |
| 249 | db/MOUSE/ur | UP00036_2 | Myf6_second | VNNRACAGV | 15 | 10 | 1.4e-003 | CENTRIMO | | | http://the_brain.bwh.harvard.edu/uniprobe/details2.php?id=00036 |
| 250 | db/MOUSE/ur | UP00014_1 | Sox17_primar | MDAAACAAT | 15 | 10 | 1.8e-003 | CENTRIMO | | | http://the_brain.bwh.harvard.edu/uniprobe/details2.php?id=00014 |
| 251 | db/MOUSE/ur | UP00069_1 | Sox1_primary | HWDYAATTC | 16 | 10 | 1.9e-003 | CENTRIMO | | | http://the_brain.bwh.harvard.edu/uniprobe/details2.php?id=00069 |
| 252 | db/MOUSE/ur | UP00135_1 | Hoxc12_3480 | YHRGGTCGT | 17 | 10 | 1.9e-003 | CENTRIMO | | | http://the_brain.bwh.harvard.edu/uniprobe/details2.php?id=00135 |
| 253 | db/MOUSE/ur | UP00177_1 | Hoxd12_3481 | HWNVGTCGT | 17 | 10 | 1.9e-003 | CENTRIMO | | | http://the_brain.bwh.harvard.edu/uniprobe/details2.php?id=00177 |
| 254 | db/EUKARYO | EVX1_DBD | | VVTAATTABS | 10 | 10 | 2.6e-003 | CENTRIMO | | | http://floresta.eead.csic.es/footprintdb/index.php?db=HumanTF:1.0&motif=EVX1_DBD |
| 255 | db/EUKARYO | EVX2_DBD | | VVTAATTAVB | 10 | 10 | 2.6e-003 | CENTRIMO | | | http://floresta.eead.csic.es/footprintdb/index.php?db=HumanTF:1.0&motif=EVX2_DBD |
| 256 | db/EUKARYO | HOXA2_DBD | | NSTMATTAV | 10 | 10 | 2.6e-003 | CENTRIMO | | | http://floresta.eead.csic.es/footprintdb/index.php?db=HumanTF:1.0&motif=HOXA2_DBD |
| 257 | db/EUKARYO | HOXB3_DBD | | NNYMATTAR | 10 | 10 | 2.6e-003 | CENTRIMO | | | http://floresta.eead.csic.es/footprintdb/index.php?db=HumanTF:1.0&motif=HOXB3_DBD |
| 258 | db/JASPAR/J | MA0612.1 | EMX1 | VYTAATKASE | 10 | 10 | 2.6e-003 | CENTRIMO | | | http://jaspar2018.genereg.net/matrix/MA0612.1 |
| 259 | db/JASPAR/J | MA0887.1 | EVX1 | VVTAATTABS | 10 | 10 | 2.6e-003 | CENTRIMO | | | http://jaspar2018.genereg.net/matrix/MA0887.1 |
| 260 | db/JASPAR/J | MA0888.1 | EVX2 | VVTAATTAVB | 10 | 10 | 2.6e-003 | CENTRIMO | | | http://jaspar2018.genereg.net/matrix/MA0888.1 |
| 261 | db/JASPAR/J | MA0900.1 | HOXA2 | NSTMATTAV | 10 | 10 | 2.6e-003 | CENTRIMO | | | http://jaspar2018.genereg.net/matrix/MA0900.1 |
| 262 | db/JASPAR/J | MA0903.1 | HOXB3 | NNYMATTAR | 10 | 10 | 2.6e-003 | CENTRIMO | | | http://jaspar2018.genereg.net/matrix/MA0903.1 |
| 263 | DREME | CCAKCC | DREME-7 | CCAKCC | 6 | 263 | 3.4e-003 | DREME | | | |
| 264 | db/EUKARYO | FOXO6_DBD_1 | | GTAAACATG | 14 | 10 | 4.0e-003 | CENTRIMO | | | http://floresta.eead.csic.es/footprintdb/index.php?db=HumanTF:1.0&motif=FOXO6_DBD_1 |
| 265 | db/EUKARYO | GRHL1_DBD_1 | | AACCGGTYT | 17 | 10 | 4.2e-003 | CENTRIMO | | | http://floresta.eead.csic.es/footprintdb/index.php?db=HumanTF:1.0&motif=GRHL1_DBD_1 |
| 266 | db/EUKARYO | MAFF_DBD | | WTGCTGAST | 15 | 10 | 5.4e-003 | CENTRIMO | | | http://floresta.eead.csic.es/footprintdb/index.php?db=HumanTF:1.0&motif=MAFF_DBD |
| 267 | db/EUKARYO | MAFK_full_2 | | DTGCTGAST | 15 | 10 | 5.4e-003 | CENTRIMO | | | http://floresta.eead.csic.es/footprintdb/index.php?db=HumanTF:1.0&motif=MAFK_full_2 |
| 268 | db/EUKARYO | Mafb_DBD_2 | | WNTGCTGAS | 17 | 10 | 5.4e-003 | CENTRIMO | | | http://floresta.eead.csic.es/footprintdb/index.php?db=HumanTF:1.0&motif=Mafb_DBD_2 |
| 269 | db/JASPAR/J | MA0496.2 | MAFK | WWWTGCTG | 19 | 10 | 5.5e-003 | CENTRIMO | | | http://jaspar2018.genereg.net/matrix/MA0496.2 |
| 270 | db/EUKARYO | MAFG_full | | WWWNTGCT | 21 | 10 | 5.5e-003 | CENTRIMO | | | http://floresta.eead.csic.es/footprintdb/index.php?db=HumanTF:1.0&motif=MAFG_full |
| 271 | db/EUKARYO | MAFK_DBD_2 | | DWWNTGCT | 21 | 10 | 5.5e-003 | CENTRIMO | | | http://floresta.eead.csic.es/footprintdb/index.php?db=HumanTF:1.0&motif=MAFK_DBD_2 |
| 272 | db/JASPAR/J | MA0659.1 | MAFG | WWWNTGCT | 21 | 10 | 5.5e-003 | CENTRIMO | | | http://jaspar2018.genereg.net/matrix/MA0659.1 |
| 273 | db/JASPAR/J | MA0495.2 | MAFF | DWWTTGCT | 21 | 10 | 5.5e-003 | CENTRIMO | | | http://jaspar2018.genereg.net/matrix/MA0495.2 |
| 274 | db/EUKARYO | RARG_full_3 | | RAGGTCAHB | 17 | 10 | 5.9e-003 | CENTRIMO | | | http://floresta.eead.csic.es/footprintdb/index.php?db=HumanTF:1.0&motif=RARG_full_3 |
| 275 | db/EUKARYO | RARA_DBD_1 | | AAAGGTCAT | 18 | 10 | 5.9e-003 | CENTRIMO | | | http://floresta.eead.csic.es/footprintdb/index.php?db=HumanTF:1.0&motif=RARA_DBD_1 |
| 276 | db/EUKARYO | RARA_full_2 | | ARRGGTCAH | 18 | 10 | 5.9e-003 | CENTRIMO | | | http://floresta.eead.csic.es/footprintdb/index.php?db=HumanTF:1.0&motif=RARA_full_2 |
| 277 | db/JASPAR/J | MA0155.1 | INSM1 | TGYCWGGG | 12 | 10 | 6.5e-003 | CENTRIMO | | | http://jaspar2018.genereg.net/matrix/MA0155.1 |
| 278 | db/EUKARYO | SMAD3_DBD | | YGTCTAGAC | 10 | 10 | 7.1e-003 | CENTRIMO | | | http://floresta.eead.csic.es/footprintdb/index.php?db=HumanTF:1.0&motif=SMAD3_DBD |
| 279 | db/JASPAR/J | MA0795.1 | SMAD3 | YGTCTAGAC | 10 | 10 | 7.1e-003 | CENTRIMO | | | http://jaspar2018.genereg.net/matrix/MA0795.1 |
| 280 | db/JASPAR/J | MA0259.1 | ARNT::HIF1A | VBACGTGC | 8 | 10 | 7.5e-003 | CENTRIMO | | | http://jaspar2018.genereg.net/matrix/MA0259.1 |
| 281 | DREME | AAGAKCCT | DREME-8 | AAGAKCCT | 8 | 39 | 7.6e-003 | DREME | | | |

| | | | | | | | | | | | |
|---|---|---|---|---|---|---|---|---|---|---|---|
| 282 | db/EUKARYO | TBX2_full_1 | | GGTGTGARA | 18 | 10 | 9.7e-003 | CENTRIMO | | | http://floresta.eead.csic.es/footprintdb/index.php?db=HumanTF:1.0&motif=TBX2_full_1 |
| 283 | db/EUKARYO | HEY2_DBD | | GRCACGTGC | 10 | 10 | 1.0e-002 | CENTRIMO | | | http://floresta.eead.csic.es/footprintdb/index.php?db=HumanTF:1.0&motif=HEY2_DBD |
| 284 | db/JASPAR/J | MA0632.1 | Tcfl5 | NBCDCGHG | 10 | 10 | 1.0e-002 | CENTRIMO | | | http://jaspar2018.genereg.net/matrix/MA0632.1 |
| 285 | db/EUKARYO | E2F1_DBD_1 | | WWTGGCGC | 12 | 10 | 1.0e-002 | CENTRIMO | | | http://floresta.eead.csic.es/footprintdb/index.php?db=HumanTF:1.0&motif=E2F1_DBD_1 |
| 286 | db/MOUSE/ur | UP00017_1 | Nkx3-1_prima | HWDANCCAC | 17 | 10 | 1.2e-002 | CENTRIMO | | | http://the_brain.bwh.harvard.edu/uniprobe/details2.php?id=00017 |
| 287 | db/JASPAR/J | MA0514.1 | Sox3 | CCTTTGTYY | 10 | 10 | 1.4e-002 | CENTRIMO | | | http://jaspar2018.genereg.net/matrix/MA0514.1 |
| 288 | db/MOUSE/ur | UP00101_2 | Sox12_secon | VVNBASACA | 16 | 10 | 1.4e-002 | CENTRIMO | | | http://the_brain.bwh.harvard.edu/uniprobe/details2.php?id=00101 |
| 289 | db/EUKARYO | Tcfap2a_DBD_1 | | TGCCCYVRG | 12 | 10 | 1.8e-002 | CENTRIMO | | | http://floresta.eead.csic.es/footprintdb/index.php?db=HumanTF:1.0&motif=Tcfap2a_DBD_1 |
| 290 | db/JASPAR/J | MA0626.1 | Npas2 | NSCACGTGT | 10 | 10 | 1.9e-002 | CENTRIMO | | | http://jaspar2018.genereg.net/matrix/MA0626.1 |
| 291 | db/EUKARYO | HEY1_DBD | | GRCACGTGC | 10 | 10 | 2.0e-002 | CENTRIMO | | | http://floresta.eead.csic.es/footprintdb/index.php?db=HumanTF:1.0&motif=HEY1_DBD |
| 292 | db/EUKARYO | HEY2_full | | GRCACGTGY | 10 | 10 | 2.0e-002 | CENTRIMO | | | http://floresta.eead.csic.es/footprintdb/index.php?db=HumanTF:1.0&motif=HEY2_full |
| 293 | db/JASPAR/J | MA0649.1 | HEY2 | GRCACGTGY | 10 | 10 | 2.0e-002 | CENTRIMO | | | http://jaspar2018.genereg.net/matrix/MA0649.1 |
| 294 | db/JASPAR/J | MA0823.1 | HEY1 | GRCACGTGC | 10 | 10 | 2.0e-002 | CENTRIMO | | | http://jaspar2018.genereg.net/matrix/MA0823.1 |
| 295 | db/EUKARYO | MTF1_DBD | | TTTGCACAC | 14 | 10 | 2.0e-002 | CENTRIMO | | | http://floresta.eead.csic.es/footprintdb/index.php?db=HumanTF:1.0&motif=MTF1_DBD |
| 296 | db/JASPAR/J | MA0863.1 | MTF1 | TTTGCACAC | 14 | 10 | 2.0e-002 | CENTRIMO | | | http://jaspar2018.genereg.net/matrix/MA0863.1 |
| 297 | db/MOUSE/ur | UP00050_2 | Bhlhb2_secon | YRYSNHTMC | 23 | 10 | 2.0e-002 | CENTRIMO | | | http://the_brain.bwh.harvard.edu/uniprobe/details2.php?id=00050 |
| 298 | db/MOUSE/ur | UP00097_1 | Mtf1_primary | DBRYCGTGT | 16 | 10 | 2.0e-002 | CENTRIMO | | | http://the_brain.bwh.harvard.edu/uniprobe/details2.php?id=00097 |
| 299 | db/MOUSE/ur | UP00065_1 | Zfp161_prima | KGGCGCGC | 16 | 10 | 2.1e-002 | CENTRIMO | | | http://the_brain.bwh.harvard.edu/uniprobe/details2.php?id=00065 |
| 300 | db/EUKARYO | Rhox11_DBD | | YGCTGTWAH | 9 | 10 | 2.6e-002 | CENTRIMO | | | http://floresta.eead.csic.es/footprintdb/index.php?db=HumanTF:1.0&motif=Rhox11_DBD |
| 301 | db/EUKARYO | HMBOX1_DBD | | MYTAGTTAM | 10 | 10 | 2.7e-002 | CENTRIMO | | | http://floresta.eead.csic.es/footprintdb/index.php?db=HumanTF:1.0&motif=HMBOX1_DBD |
| 302 | db/JASPAR/J | MA0895.1 | HMBOX1 | MYTAGTTAM | 10 | 10 | 2.7e-002 | CENTRIMO | | | http://jaspar2018.genereg.net/matrix/MA0895.1 |
| 303 | db/JASPAR/J | MA1099.1 | Hes1 | SVCACGYGH | 10 | 10 | 3.1e-002 | CENTRIMO | | | http://jaspar2018.genereg.net/matrix/MA1099.1 |
| 304 | db/JASPAR/J | MA0147.3 | MYC | NNCCACGTG | 12 | 10 | 3.1e-002 | CENTRIMO | | | http://jaspar2018.genereg.net/matrix/MA0147.3 |
| 305 | db/JASPAR/J | MA1108.1 | MXI1 | NVVCCACGT | 13 | 10 | 3.1e-002 | CENTRIMO | | | http://jaspar2018.genereg.net/matrix/MA1108.1 |
| 306 | db/EUKARYO | Egr3_DBD | | HACGCCCAC | 15 | 10 | 3.2e-002 | CENTRIMO | | | http://floresta.eead.csic.es/footprintdb/index.php?db=HumanTF:1.0&motif=Egr3_DBD |
| 307 | db/EUKARYO | EGR1_DBD | | HMCGCCCM | 14 | 10 | 3.2e-002 | CENTRIMO | | | http://floresta.eead.csic.es/footprintdb/index.php?db=HumanTF:1.0&motif=EGR1_DBD |
| 308 | db/EUKARYO | EGR1_full | | HACGCCCAC | 14 | 10 | 3.2e-002 | CENTRIMO | | | http://floresta.eead.csic.es/footprintdb/index.php?db=HumanTF:1.0&motif=EGR1_full |
| 309 | db/JASPAR/J | MA0162.3 | EGR1 | HACGCCCAC | 14 | 10 | 3.2e-002 | CENTRIMO | | | http://jaspar2018.genereg.net/matrix/MA0162.3 |
| 310 | db/EUKARYO | EGR4_DBD_2 | | HHACGCCCA | 16 | 10 | 3.2e-002 | CENTRIMO | | | http://floresta.eead.csic.es/footprintdb/index.php?db=HumanTF:1.0&motif=EGR4_DBD_2 |
| 311 | db/EUKARYO | EGR2_DBD | | MCGCCCACC | 11 | 10 | 3.2e-002 | CENTRIMO | | | http://floresta.eead.csic.es/footprintdb/index.php?db=HumanTF:1.0&motif=EGR2_DBD |
| 312 | db/JASPAR/J | MA0472.2 | EGR2 | MCGCCCACC | 11 | 10 | 3.2e-002 | CENTRIMO | | | http://jaspar2018.genereg.net/matrix/MA0472.2 |
| 313 | db/EUKARYO | EGR2_full | | NHMCGCCCA | 15 | 10 | 3.2e-002 | CENTRIMO | | | http://floresta.eead.csic.es/footprintdb/index.php?db=HumanTF:1.0&motif=EGR2_full |
| 314 | db/EUKARYO | EGR3_DBD | | HHMCGCCCA | 15 | 10 | 3.2e-002 | CENTRIMO | | | http://floresta.eead.csic.es/footprintdb/index.php?db=HumanTF:1.0&motif=EGR3_DBD |
| 315 | db/JASPAR/J | MA0732.1 | EGR3 | HHMCGCCCA | 15 | 10 | 3.2e-002 | CENTRIMO | | | http://jaspar2018.genereg.net/matrix/MA0732.1 |
| 316 | db/MOUSE/ur | UP00084_2 | Gmeb1_secor | KGRBCRACG | 16 | 10 | 3.5e-002 | CENTRIMO | | | http://the_brain.bwh.harvard.edu/uniprobe/details2.php?id=00084 |
| 317 | db/JASPAR/J | MA0506.1 | NRF1 | GCGCVTGCG | 11 | 10 | 3.5e-002 | CENTRIMO | | | http://jaspar2018.genereg.net/matrix/MA0506.1 |
| 318 | db/MOUSE/ur | UP00088_1 | Plagl1_primar | BNGGGGGS | 16 | 10 | 3.6e-002 | CENTRIMO | | | http://the_brain.bwh.harvard.edu/uniprobe/details2.php?id=00088 |
| 319 | db/EUKARYO | SRF_full | | TGMCCATAT | 16 | 10 | 4.4e-002 | CENTRIMO | | | http://floresta.eead.csic.es/footprintdb/index.php?db=HumanTF:1.0&motif=SRF_full |
| 320 | db/JASPAR/J | MA0083.3 | SRF | TGMCCATAT | 16 | 10 | 4.4e-002 | CENTRIMO | | | http://jaspar2018.genereg.net/matrix/MA0083.3 |
| 321 | db/JASPAR/J | MA0109.1 | HLTF | NHMCWTDK | 10 | 10 | 4.5e-002 | CENTRIMO | | | http://jaspar2018.genereg.net/matrix/MA0109.1 |

### MEME-ChIP (Motif Analysis of Large Nucleotide Datasets): Version 5.0.3 released on Sun Dec 02 18:41:45 2018 -0800
### The format of this file is described at http://meme-suite.org//doc/meme-chip-output-format.html
### meme-chip -oc . -time 300 -ccut 100 -order 1 -db db/EUKARYOTE/jolma2013.meme -db db/JASPAR/JASPAR2018_CORE_vertebrates_non-redundant.meme -db db/MOUSE/uniprobe_mouse.meme -meme-mod zoops -meme-minw 6 -meme-maxw 30 -meme-nmotifs 3 -meme-searchsize 1000(

**ZT6 dynamic**

| MOTIF_INDEX | MOTIF_SOUF | MOTIF_ID | ALT_ID | CONSENSUS | WIDTH | SITES | E-VALUE | E-VALUE_SO | MOST_SIMILAR_MC | **MOST_SIMILAR_MO** | URL |
|---|---|---|---|---|---|---|---|---|---|---|---|
| 1 | MEME | GTGTGTRTG | MEME-2 | GTGTGTRTG | 29 | 61 | 8.6e-116 | MEME | | | |
| 2 | MEME | GAGTTCMAG | MEME-1 | GAGTTCMAG | 21 | 60 | 2.9e-103 | MEME | db/JASPAR/JASPAR | **MA0505.1 (Nr5a2)** | http://jaspar2018.genereg.net/matrix/MA0505.1 |
| 3 | MEME | CCDCCTGCC | MEME-3 | CCDCCTGCC | 30 | 26 | 7.3e-085 | MEME | db/MOUSE/uniprobe | **UP00087_2 (Tcfap2c_** | http://the_brain.bwh.harvard.edu/uniprobe/details2.php?id=00087 |
| 4 | DREME | AKAAA | DREME-1 | AKAAA | 5 | 1377 | 2.1e-007 | DREME | | | |
| 5 | DREME | ACACRS | DREME-3 | ACACRS | 6 | 556 | 6.4e-007 | DREME | db/MOUSE/uniprobe | **UP00042_2 (Gm397_** | http://the_brain.bwh.harvard.edu/uniprobe/details2.php?id=00042 |
| 6 | DREME | CCAGSM | DREME-2 | CCAGSM | 6 | 714 | 9.3e-007 | DREME | db/JASPAR/JASPAR | **MA1121.1 (TEAD2)** | http://jaspar2018.genereg.net/matrix/MA1121.1 |
| 7 | DREME | CTGYCTC | DREME-4 | CTGYCTC | 7 | 166 | 1.7e-004 | DREME | | | |
| 8 | DREME | CTGAGTTC | DREME-5 | CTGAGTTC | 8 | 31 | 2.3e-002 | DREME | | | |
| 9 | DREME | GAAGRCA | DREME-6 | GAAGRCA | 7 | 92 | 2.7e-002 | DREME | | | |
| 10 | DREME | GCCACCAY | DREME-7 | GCCACCAY | 8 | 50 | 4.4e-002 | DREME | | | |
| 11 | DREME | GCCATCTY | DREME-8 | GCCATCTY | 8 | 37 | 4.4e-002 | DREME | db/JASPAR/JASPAR | **MA0095.2 (YY1)** | http://jaspar2018.genereg.net/matrix/MA0095.2 |

### MEME-ChIP (Motif Analysis of Large Nucleotide Datasets): Version 5.0.3 released on Sun Dec 02 18:41:45 2018 -0800
### The format of this file is described at http://meme-suite.org//doc/meme-chip-output-format.html
### meme-chip -oc . -time 300 -ccut 100 -order 1 -db db/EUKARYOTE/jolma2013.meme -db db/JASPAR/JASPAR2018_CORE_vertebrates_non-redundant.meme -db db/MOUSE/uniprobe_mouse.meme -meme-mod zoops -meme-minw 6 -meme-maxw 30 -meme-nmotifs 3 -meme-searchsize 1000(

**ZT6 static**

| MOTIF_INDEX | MOTIF_SOUF | MOTIF_ID | ALT_ID | CONSENSUS | WIDTH | SITES | E-VALUE | E-VALUE_SO | MOST_SIMILAR_MC | **MOST_SIMILAR_MO** | URL |
|---|---|---|---|---|---|---|---|---|---|---|---|
| 1 | MEME | TTTDTTTDTV | MEME-1 | TTTDTTTDTV | 30 | 357 | 8.0e-131 | MEME | db/JASPAR/JASPAR | **MA1125.1 (ZNF384)** | http://jaspar2018.genereg.net/matrix/MA1125.1 |
| 2 | MEME | CAGCCTGGT | MEME-2 | CAGCCTGGT | 30 | 20 | 3.9e-126 | MEME | | | |
| 3 | MEME | CKCCSNSNS | MEME-3 | CKCCSNSNS | 30 | 140 | 4.7e-083 | MEME | db/MOUSE/uniprobe | **UP00021_1 (Zfp281_I** | http://the_brain.bwh.harvard.edu/uniprobe/details2.php?id=00021 |
| 4 | db/EUKARYO | RHOXF1_DBD_1 | | GGATWAKCC | 9 | 12 | 8.0e-024 | CENTRIMO | | | http://floresta.eead.csic.es/footprintdb/index.php?db=HumanTF:1.0&motif=RHOXF1_DBD_1 |
| 5 | db/EUKARYO | RHOXF1_full_1 | | GGMTWATCC | 9 | 12 | 8.0e-024 | CENTRIMO | | | http://floresta.eead.csic.es/footprintdb/index.php?db=HumanTF:1.0&motif=RHOXF1_full_1 |
| 6 | db/EUKARYO | DPRX_DBD_2 | | VRGATAATCC | 11 | 12 | 8.0e-024 | CENTRIMO | | | http://floresta.eead.csic.es/footprintdb/index.php?db=HumanTF:1.0&motif=DPRX_DBD_2 |
| 7 | db/MOUSE/ur | UP00265_1 | Pitx3_3497.2 | VGGGGGATT | 16 | 12 | 1.8e-023 | CENTRIMO | | | http://the_brain.bwh.harvard.edu/uniprobe/details2.php?id=00265 |
| 8 | db/EUKARYO | MGA_DBD_3 | | GGTGTGAAW | 18 | 12 | 1.8e-023 | CENTRIMO | | | http://floresta.eead.csic.es/footprintdb/index.php?db=HumanTF:1.0&motif=MGA_DBD_3 |
| 9 | db/EUKARYO | TBX2_full_1 | | GGTGTGARA | 18 | 12 | 1.8e-023 | CENTRIMO | | | http://floresta.eead.csic.es/footprintdb/index.php?db=HumanTF:1.0&motif=TBX2_full_1 |
| 10 | db/EUKARYO | TBX4_DBD_2 | | AGGTGTGAD | 20 | 12 | 1.8e-023 | CENTRIMO | | | http://floresta.eead.csic.es/footprintdb/index.php?db=HumanTF:1.0&motif=TBX4_DBD_2 |
| 11 | db/EUKARYO | TBX5_DBD_2 | | AGGTGTKAN | 20 | 12 | 1.8e-023 | CENTRIMO | | | http://floresta.eead.csic.es/footprintdb/index.php?db=HumanTF:1.0&motif=TBX5_DBD_2 |
| 12 | db/MOUSE/ur | UP00176_1 | Crx_3485.1 | YBWWGGGG | 16 | 12 | 2.6e-022 | CENTRIMO | | | http://the_brain.bwh.harvard.edu/uniprobe/details2.php?id=00176 |
| 13 | db/JASPAR/J/ | MA0468.1 | DUX4 | TAAYYYAATC | 11 | 12 | 4.9e-020 | CENTRIMO | | | http://jaspar2018.genereg.net/matrix/MA0468.1 |
| 14 | db/EUKARYO | EOMES_DBD_1 | | RAGGTGTGA | 13 | 12 | 4.3e-018 | CENTRIMO | | | http://floresta.eead.csic.es/footprintdb/index.php?db=HumanTF:1.0&motif=EOMES_DBD_1 |
| 15 | db/JASPAR/J/ | MA0800.1 | EOMES | RAGGTGTGA | 13 | 12 | 4.3e-018 | CENTRIMO | | | http://jaspar2018.genereg.net/matrix/MA0800.1 |
| 16 | db/EUKARYO | TBR1_DBD | | AGGTGTGAA | 10 | 12 | 5.5e-018 | CENTRIMO | | | http://floresta.eead.csic.es/footprintdb/index.php?db=HumanTF:1.0&motif=TBR1_DBD |
| 17 | db/JASPAR/J/ | MA0802.1 | TBR1 | AGGTGTGAA | 10 | 12 | 5.5e-018 | CENTRIMO | | | http://jaspar2018.genereg.net/matrix/MA0802.1 |
| 18 | db/EUKARYO | MGA_DBD_1 | | AGGTGTGA | 8 | 12 | 9.2e-018 | CENTRIMO | | | http://floresta.eead.csic.es/footprintdb/index.php?db=HumanTF:1.0&motif=MGA_DBD_1 |
| 19 | db/EUKARYO | TBX1_DBD_3 | | AGGTGTGA | 8 | 12 | 9.2e-018 | CENTRIMO | | | http://floresta.eead.csic.es/footprintdb/index.php?db=HumanTF:1.0&motif=TBX1_DBD_3 |
| 20 | db/JASPAR/J/ | MA0801.1 | MGA | AGGTGTGA | 8 | 12 | 9.2e-018 | CENTRIMO | | | http://jaspar2018.genereg.net/matrix/MA0801.1 |
| 21 | db/JASPAR/J/ | MA0805.1 | TBX1 | AGGTGTGA | 8 | 12 | 9.2e-018 | CENTRIMO | | | http://jaspar2018.genereg.net/matrix/MA0805.1 |
| 22 | db/JASPAR/J/ | MA0690.1 | TBX21 | AAGGTGTGA | 10 | 12 | 9.2e-018 | CENTRIMO | | | http://jaspar2018.genereg.net/matrix/MA0690.1 |
| 23 | db/MOUSE/ur | UP00170_1 | Isl2_3430.1 | MHMAAYYMA | 16 | 12 | 9.1e-017 | CENTRIMO | | | http://the_brain.bwh.harvard.edu/uniprobe/details2.php?id=00170 |
| 24 | db/JASPAR/J/ | MA0896.1 | Hmx1 | VSVAGCAATT | 17 | 12 | 5.1e-016 | CENTRIMO | | | http://jaspar2018.genereg.net/matrix/MA0896.1 |
| 25 | db/EUKARYO | HINFP1_full_1 | | CARCGTCCG | 12 | 12 | 3.1e-015 | CENTRIMO | | | http://floresta.eead.csic.es/footprintdb/index.php?db=HumanTF:1.0&motif=HINFP1_full_1 |
| 26 | db/JASPAR/J/ | MA0131.2 | HINFP | CARCGTCCG | 12 | 12 | 3.1e-015 | CENTRIMO | | | http://jaspar2018.genereg.net/matrix/MA0131.2 |
| 27 | db/MOUSE/ur | UP00211_1 | Pou3f3_3235. | WANDTATGC | 17 | 12 | 3.3e-013 | CENTRIMO | | | http://the_brain.bwh.harvard.edu/uniprobe/details2.php?id=00211 |
| 28 | db/JASPAR/J/ | MA0119.1 | NFIC::TLX1 | TGGCASSRW | 14 | 12 | 3.5e-012 | CENTRIMO | | | http://jaspar2018.genereg.net/matrix/MA0119.1 |
| 29 | db/EUKARYO | NFIA_full_1 | | TTGGCAHND | 15 | 12 | 3.8e-012 | CENTRIMO | | | http://floresta.eead.csic.es/footprintdb/index.php?db=HumanTF:1.0&motif=NFIA_full_1 |
| 30 | db/EUKARYO | NFIB_full | | TTGGCAHND | 15 | 12 | 3.8e-012 | CENTRIMO | | | http://floresta.eead.csic.es/footprintdb/index.php?db=HumanTF:1.0&motif=NFIB_full |
| 31 | db/EUKARYO | NFIX_full_4 | | YTGGCAHND | 15 | 12 | 3.8e-012 | CENTRIMO | | | http://floresta.eead.csic.es/footprintdb/index.php?db=HumanTF:1.0&motif=NFIX_full_4 |
| 32 | db/JASPAR/J/ | MA0481.2 | FOXP1 | NDGTAAACA | 12 | 12 | 4.2e-012 | CENTRIMO | | | http://jaspar2018.genereg.net/matrix/MA0481.2 |
| 33 | db/JASPAR/J/ | MA0040.1 | Foxq1 | HATTGTTTAT | 11 | 12 | 4.5e-012 | CENTRIMO | | | http://jaspar2018.genereg.net/matrix/MA0040.1 |
| 34 | db/EUKARYO | NR2E1_full_2 | | AAGTCAAWA | 14 | 12 | 1.3e-011 | CENTRIMO | | | http://floresta.eead.csic.es/footprintdb/index.php?db=HumanTF:1.0&motif=NR2E1_full_2 |
| 35 | db/EUKARYO | Nr2e1_DBD_2 | | AAGTCAADA | 14 | 12 | 1.3e-011 | CENTRIMO | | | http://floresta.eead.csic.es/footprintdb/index.php?db=HumanTF:1.0&motif=Nr2e1_DBD_2 |
| 36 | db/EUKARYO | HEY1_DBD | | GRCACGTGC | 10 | 12 | 6.2e-011 | CENTRIMO | | | http://floresta.eead.csic.es/footprintdb/index.php?db=HumanTF:1.0&motif=HEY1_DBD |
| 37 | db/EUKARYO | TFE3_DBD | | VYCACGTGA | 10 | 12 | 6.2e-011 | CENTRIMO | | | http://floresta.eead.csic.es/footprintdb/index.php?db=HumanTF:1.0&motif=TFE3_DBD |
| 38 | db/JASPAR/J/ | MA1099.1 | Hes1 | SVCACGYGH | 10 | 12 | 6.2e-011 | CENTRIMO | | | http://jaspar2018.genereg.net/matrix/MA1099.1 |
| 39 | db/JASPAR/J/ | MA0823.1 | HEY1 | GRCACGTGC | 10 | 12 | 6.2e-011 | CENTRIMO | | | http://jaspar2018.genereg.net/matrix/MA0823.1 |
| 40 | db/JASPAR/J/ | MA0104.4 | MYCN | VVCCACGTG | 12 | 12 | 6.3e-011 | CENTRIMO | | | http://jaspar2018.genereg.net/matrix/MA0104.4 |
| 41 | db/EUKARYO | ERG_DBD_2 | | ACCGGAWAT | 14 | 12 | 8.2e-011 | CENTRIMO | | | http://floresta.eead.csic.es/footprintdb/index.php?db=HumanTF:1.0&motif=ERG_DBD_2 |
| 42 | db/EUKARYO | ERG_full_2 | | ACCGGAWAT | 14 | 12 | 8.2e-011 | CENTRIMO | | | http://floresta.eead.csic.es/footprintdb/index.php?db=HumanTF:1.0&motif=ERG_full_2 |
| 43 | db/EUKARYO | FLI1_DBD_2 | | ACCGGAWAT | 14 | 12 | 8.2e-011 | CENTRIMO | | | http://floresta.eead.csic.es/footprintdb/index.php?db=HumanTF:1.0&motif=FLI1_DBD_2 |
| 44 | db/EUKARYO | FLI1_full_2 | | ACCGGAAAT | 14 | 12 | 8.2e-011 | CENTRIMO | | | http://floresta.eead.csic.es/footprintdb/index.php?db=HumanTF:1.0&motif=FLI1_full_2 |
| 45 | db/MOUSE/ur | UP00038_2 | Spdef_second | DWNNACATC | 16 | 12 | 1.1e-010 | CENTRIMO | | | http://the_brain.bwh.harvard.edu/uniprobe/details2.php?id=00038 |
| 46 | db/EUKARYO | OTX1_DBD_1 | | BNTAATCCG/ | 15 | 12 | 1.3e-010 | CENTRIMO | | | http://floresta.eead.csic.es/footprintdb/index.php?db=HumanTF:1.0&motif=OTX1_DBD_1 |
| 47 | db/EUKARYO | OTX2_DBD_1 | | DHTAATCCG/ | 15 | 12 | 1.3e-010 | CENTRIMO | | | http://floresta.eead.csic.es/footprintdb/index.php?db=HumanTF:1.0&motif=OTX2_DBD_1 |
| 48 | db/EUKARYO | Otx1_DBD_1 | | NHTAATCCG/ | 15 | 12 | 1.3e-010 | CENTRIMO | | | http://floresta.eead.csic.es/footprintdb/index.php?db=HumanTF:1.0&motif=Otx1_DBD_1 |
| 49 | DREME | AKGCWGG | DREME-1 | AKGCWGG | 7 | 262 | 2.6e-010 | DREME | | | |
| 50 | DREME | ASACAS | DREME-2 | ASACAS | 6 | 891 | 2.1e-009 | DREME | | | |
| 51 | db/EUKARYO | EN1_full_2 | | TAATTRSNYA | 14 | 12 | 9.3e-009 | CENTRIMO | | | http://floresta.eead.csic.es/footprintdb/index.php?db=HumanTF:1.0&motif=EN1_full_2 |
| 52 | db/EUKARYO | ZNF410_DBD | | KMCATCCCA | 17 | 12 | 1.3e-008 | CENTRIMO | | | http://floresta.eead.csic.es/footprintdb/index.php?db=HumanTF:1.0&motif=ZNF410_DBD |

| | | | | | | | | | |
|---|---|---|---|---|---|---|---|---|---|
| 53 | db/JASPAR/JA | MA0752.1 | ZNF410 | KMCATCCCA | 17 | 12 | 1.3e-008 | CENTRIMO | http://jaspar2018.genereg.net/matrix/MA0752.1 |
| 54 | db/EUKARYO | Hoxd9_DBD_1 | | SCCATWAAA | 9 | 12 | 1.4e-008 | CENTRIMO | http://floresta.eead.csic.es/footprintdb/index.php?db=HumanTF:1.0&motif=Hoxd9_DBD_1 |
| 55 | db/EUKARYO | ZNF143_DBD | | TWCCCAYAA | 16 | 12 | 1.6e-008 | CENTRIMO | http://floresta.eead.csic.es/footprintdb/index.php?db=HumanTF:1.0&motif=ZNF143_DBD |
| 56 | db/JASPAR/JA | MA0088.2 | ZNF143 | TWCCCAYAA | 16 | 12 | 1.6e-008 | CENTRIMO | http://jaspar2018.genereg.net/matrix/MA0088.2 |
| 57 | db/MOUSE/ur | UP00192_1 | Six1_0935.2 | RATRGGGTA | 17 | 12 | 1.8e-008 | CENTRIMO | http://the_brain.bwh.harvard.edu/uniprobe/details2.php?id=00192 |
| 58 | db/MOUSE/ur | UP00063_2 | Hoxa3_secon | AAAANCBAT | 14 | 12 | 1.9e-008 | CENTRIMO | http://the_brain.bwh.harvard.edu/uniprobe/details2.php?id=00063 |
| 59 | db/EUKARYO | LHX9_DBD_2 | | TAATTGCYAA | 13 | 12 | 2.3e-008 | CENTRIMO | http://floresta.eead.csic.es/footprintdb/index.php?db=HumanTF:1.0&motif=LHX9_DBD_2 |
| 60 | db/EUKARYO | TBX21_DBD_3 | | TMACACCTH | 19 | 12 | 2.5e-008 | CENTRIMO | http://floresta.eead.csic.es/footprintdb/index.php?db=HumanTF:1.0&motif=TBX21_DBD_3 |
| 61 | db/EUKARYO | POU6F2_DBD_1 | | ASCTMATTAV | 10 | 12 | 2.8e-008 | CENTRIMO | http://floresta.eead.csic.es/footprintdb/index.php?db=HumanTF:1.0&motif=POU6F2_DBD_1 |
| 62 | db/JASPAR/JA | MA0793.1 | POU6F2 | ASCTMATTAV | 10 | 12 | 2.8e-008 | CENTRIMO | http://jaspar2018.genereg.net/matrix/MA0793.1 |
| 63 | db/MOUSE/ur | UP00146_1 | Pou6f1_1731. | VHNSWTAAT | 17 | 12 | 2.9e-008 | CENTRIMO | http://the_brain.bwh.harvard.edu/uniprobe/details2.php?id=00146 |
| 64 | db/MOUSE/ur | UP00146_2 | Pou6f1_3733. | VHHVATAATG | 17 | 12 | 2.9e-008 | CENTRIMO | http://the_brain.bwh.harvard.edu/uniprobe/details2.php?id=00146 |
| 65 | db/EUKARYO | NKX6-1_DBD | | NTMATTAA | 8 | 12 | 3.0e-008 | CENTRIMO | http://floresta.eead.csic.es/footprintdb/index.php?db=HumanTF:1.0&motif=NKX6-1_DBD |
| 66 | db/EUKARYO | NKX6-2_DBD | | NYMATTAA | 8 | 12 | 3.0e-008 | CENTRIMO | http://floresta.eead.csic.es/footprintdb/index.php?db=HumanTF:1.0&motif=NKX6-2_DBD |
| 67 | db/EUKARYO | NKX6-2_full | | NYMATTAA | 8 | 12 | 3.0e-008 | CENTRIMO | http://floresta.eead.csic.es/footprintdb/index.php?db=HumanTF:1.0&motif=NKX6-2_full |
| 68 | db/JASPAR/JA | MA0618.1 | LBX1 | TTAATTAG | 8 | 12 | 3.0e-008 | CENTRIMO | http://jaspar2018.genereg.net/matrix/MA0618.1 |
| 69 | db/JASPAR/JA | MA0675.1 | NKX6-2 | NYMATTAA | 8 | 12 | 3.0e-008 | CENTRIMO | http://jaspar2018.genereg.net/matrix/MA0675.1 |
| 70 | db/EUKARYO | GSX1_DBD | | NCYMATTAVN | 10 | 12 | 3.0e-008 | CENTRIMO | http://floresta.eead.csic.es/footprintdb/index.php?db=HumanTF:1.0&motif=GSX1_DBD |
| 71 | db/JASPAR/JA | MA0892.1 | GSX1 | NCYMATTAVN | 10 | 12 | 3.0e-008 | CENTRIMO | http://jaspar2018.genereg.net/matrix/MA0892.1 |
| 72 | db/MOUSE/ur | UP00254_1 | Pou2f1_3081. | DTDHRHTAA | 16 | 12 | 3.3e-008 | CENTRIMO | http://the_brain.bwh.harvard.edu/uniprobe/details2.php?id=00254 |
| 73 | db/MOUSE/ur | UP00054_2 | Tcf7_seconda | MBKTATTANA | 15 | 12 | 3.7e-008 | CENTRIMO | http://the_brain.bwh.harvard.edu/uniprobe/details2.php?id=00054 |
| 74 | db/EUKARYO | MEOX2_DBD_3 | | STMATCATCA | 14 | 12 | 3.9e-008 | CENTRIMO | http://floresta.eead.csic.es/footprintdb/index.php?db=HumanTF:1.0&motif=MEOX2_DBD_3 |
| 75 | db/MOUSE/ur | UP00023_2 | Sox30_secon | TMNBATTATA | 16 | 12 | 3.9e-008 | CENTRIMO | http://the_brain.bwh.harvard.edu/uniprobe/details2.php?id=00023 |
| 76 | db/MOUSE/ur | UP00069_2 | Sox1_second | NYRTAATTGT | 15 | 12 | 4.4e-008 | CENTRIMO | http://the_brain.bwh.harvard.edu/uniprobe/details2.php?id=00069 |
| 77 | db/MOUSE/ur | UP00219_2 | Cutl1_3494.2 | TARTGATRAT | 15 | 12 | 4.4e-008 | CENTRIMO | http://the_brain.bwh.harvard.edu/uniprobe/details2.php?id=00219 |
| 78 | db/EUKARYO | MEOX2_DBD_2 | | KTAATTACSS | 17 | 12 | 4.4e-008 | CENTRIMO | http://floresta.eead.csic.es/footprintdb/index.php?db=HumanTF:1.0&motif=MEOX2_DBD_2 |
| 79 | db/MOUSE/ur | UP00091_1 | Sox5_primary | WWDRGAAC | 16 | 12 | 4.6e-008 | CENTRIMO | http://the_brain.bwh.harvard.edu/uniprobe/details2.php?id=00091 |
| 80 | db/MOUSE/ur | UP00071_1 | Sox21_primar | HHHWATTATA | 16 | 12 | 4.9e-008 | CENTRIMO | http://the_brain.bwh.harvard.edu/uniprobe/details2.php?id=00071 |
| 81 | db/MOUSE/ur | UP00101_1 | Sox12_primar | THATTGTTMT | 14 | 12 | 5.3e-008 | CENTRIMO | http://the_brain.bwh.harvard.edu/uniprobe/details2.php?id=00101 |
| 82 | db/MOUSE/ur | UP00051_1 | Sox8_primary | NDATBWATT | 17 | 12 | 5.5e-008 | CENTRIMO | http://the_brain.bwh.harvard.edu/uniprobe/details2.php?id=00051 |
| 83 | db/EUKARYO | FOXD2_DBD_1 | | HRHWAATAT | 14 | 12 | 5.7e-008 | CENTRIMO | http://floresta.eead.csic.es/footprintdb/index.php?db=HumanTF:1.0&motif=FOXD2_DBD_1 |
| 84 | db/EUKARYO | BARHL2_DBD_3 | | TAAWYGNYV | 16 | 12 | 5.7e-008 | CENTRIMO | http://floresta.eead.csic.es/footprintdb/index.php?db=HumanTF:1.0&motif=BARHL2_DBD_3 |
| 85 | db/EUKARYO | BARHL2_full_3 | | TAAWYGNYS | 16 | 12 | 5.7e-008 | CENTRIMO | http://floresta.eead.csic.es/footprintdb/index.php?db=HumanTF:1.0&motif=BARHL2_full_3 |
| 86 | db/EUKARYO | Barhl1_DBD_3 | | TAAWYGBYS | 16 | 12 | 5.7e-008 | CENTRIMO | http://floresta.eead.csic.es/footprintdb/index.php?db=HumanTF:1.0&motif=Barhl1_DBD_3 |
| 87 | db/MOUSE/ur | UP00217_1 | Hoxa10_2318 | DMKGYAATA | 16 | 12 | 5.7e-008 | CENTRIMO | http://the_brain.bwh.harvard.edu/uniprobe/details2.php?id=00217 |
| 88 | db/EUKARYO | PDX1_DBD_1 | | SYAATTARBR | 18 | 12 | 5.8e-008 | CENTRIMO | http://floresta.eead.csic.es/footprintdb/index.php?db=HumanTF:1.0&motif=PDX1_DBD_1 |
| 89 | db/MOUSE/ur | UP00004_1 | Sox14_primar | DHNAATTATA | 16 | 12 | 6.2e-008 | CENTRIMO | http://the_brain.bwh.harvard.edu/uniprobe/details2.php?id=00004 |
| 90 | db/MOUSE/ur | UP00016_1 | Sry_primary | NHNWATTATA | 16 | 12 | 6.2e-008 | CENTRIMO | http://the_brain.bwh.harvard.edu/uniprobe/details2.php?id=00016 |
| 91 | db/MOUSE/ur | UP00034_1 | Sox7_primary | HNKDWWDA | 22 | 12 | 6.8e-008 | CENTRIMO | http://the_brain.bwh.harvard.edu/uniprobe/details2.php?id=00034 |
| 92 | db/MOUSE/ur | UP00037_1 | Zfp105_prima | HDYAAAHAAI | 15 | 12 | 6.9e-008 | CENTRIMO | http://the_brain.bwh.harvard.edu/uniprobe/details2.php?id=00037 |
| 93 | db/EUKARYO | POU1F1_DBD_1 | | NTYATGMATA | 17 | 12 | 7.0e-008 | CENTRIMO | http://floresta.eead.csic.es/footprintdb/index.php?db=HumanTF:1.0&motif=POU1F1_DBD_1 |
| 94 | db/MOUSE/ur | UP00075_1 | Sox15_primar | WNNDGAACA | 17 | 12 | 7.0e-008 | CENTRIMO | http://the_brain.bwh.harvard.edu/uniprobe/details2.php?id=00075 |
| 95 | db/MOUSE/ur | UP00158_1 | Pou1f1_3818. | DVNTAATTAA | 17 | 12 | 7.0e-008 | CENTRIMO | http://the_brain.bwh.harvard.edu/uniprobe/details2.php?id=00158 |
| 96 | db/MOUSE/ur | UP00105_1 | Pou3f4_3773. | DVWTAATTA | 17 | 12 | 7.0e-008 | CENTRIMO | http://the_brain.bwh.harvard.edu/uniprobe/details2.php?id=00105 |
| 97 | db/EUKARYO | POU3F3_DBD_3 | | WTRAATAWK | 12 | 12 | 7.1e-008 | CENTRIMO | http://floresta.eead.csic.es/footprintdb/index.php?db=HumanTF:1.0&motif=POU3F3_DBD_3 |
| 98 | db/EUKARYO | HNF1B_full_1 | | GTTAATNATT | 13 | 12 | 7.4e-008 | CENTRIMO | http://floresta.eead.csic.es/footprintdb/index.php?db=HumanTF:1.0&motif=HNF1B_full_1 |
| 99 | db/JASPAR/JA | MA0153.2 | HNF1B | GTTAATNATT | 13 | 12 | 7.4e-008 | CENTRIMO | http://jaspar2018.genereg.net/matrix/MA0153.2 |
| 100 | db/EUKARYO | HNF1A_full | | DRTTAATNAT | 15 | 12 | 7.5e-008 | CENTRIMO | http://floresta.eead.csic.es/footprintdb/index.php?db=HumanTF:1.0&motif=HNF1A_full |
| 101 | db/EUKARYO | HNF1B_full_2 | | NRTTAATNAT | 15 | 12 | 7.5e-008 | CENTRIMO | http://floresta.eead.csic.es/footprintdb/index.php?db=HumanTF:1.0&motif=HNF1B_full_2 |
| 102 | db/JASPAR/JA | MA0046.2 | HNF1A | DRTTAATNAT | 15 | 12 | 7.5e-008 | CENTRIMO | http://jaspar2018.genereg.net/matrix/MA0046.2 |
| 103 | db/EUKARYO | BARX1_DBD_1 | | CVATTAAAWA | 17 | 12 | 7.5e-008 | CENTRIMO | http://floresta.eead.csic.es/footprintdb/index.php?db=HumanTF:1.0&motif=BARX1_DBD_1 |
| 104 | db/MOUSE/ur | UP00255_1 | Dbx1_3486.1 | YARTWAATTA | 17 | 12 | 7.5e-008 | CENTRIMO | http://the_brain.bwh.harvard.edu/uniprobe/details2.php?id=00255 |
| 105 | db/MOUSE/ur | UP00129_1 | Pou3f1_3819. | DVNTAATTAA | 17 | 12 | 7.5e-008 | CENTRIMO | http://the_brain.bwh.harvard.edu/uniprobe/details2.php?id=00129 |
| 106 | db/EUKARYO | POU4F1_DBD | | ATGMATAATT | 14 | 12 | 7.7e-008 | CENTRIMO | http://floresta.eead.csic.es/footprintdb/index.php?db=HumanTF:1.0&motif=POU4F1_DBD |
| 107 | db/JASPAR/JA | MA0790.1 | POU4F1 | ATGMATAATT | 14 | 12 | 7.7e-008 | CENTRIMO | http://jaspar2018.genereg.net/matrix/MA0790.1 |
| 108 | db/MOUSE/ur | UP00024_2 | Glis2_second | HDTATTAWTA | 14 | 12 | 7.7e-008 | CENTRIMO | http://the_brain.bwh.harvard.edu/uniprobe/details2.php?id=00024 |
| 109 | db/MOUSE/ur | UP00218_1 | Dbx2_3487.1 | BDNHATTAAT | 16 | 12 | 7.8e-008 | CENTRIMO | http://the_brain.bwh.harvard.edu/uniprobe/details2.php?id=00218 |
| 110 | db/MOUSE/ur | UP00118_1 | Pou4f3_2791. | HVTTATTAAT | 16 | 12 | 7.8e-008 | CENTRIMO | http://the_brain.bwh.harvard.edu/uniprobe/details2.php?id=00118 |
| 111 | db/EUKARYO | MSX1_DBD_1 | | SCAATTAAAW | 18 | 12 | 7.8e-008 | CENTRIMO | http://floresta.eead.csic.es/footprintdb/index.php?db=HumanTF:1.0&motif=MSX1_DBD_1 |
| 112 | db/EUKARYO | MSX2_DBD_1 | | GCAATTAAAW | 18 | 12 | 7.8e-008 | CENTRIMO | http://floresta.eead.csic.es/footprintdb/index.php?db=HumanTF:1.0&motif=MSX2_DBD_1 |
| 113 | db/MOUSE/ur | UP00209_2 | Cart1_1275.1 | BVMNTTAATT | 17 | 12 | 8.1e-008 | CENTRIMO | http://the_brain.bwh.harvard.edu/uniprobe/details2.php?id=00209 |

| | | | | | | | | | |
|---|---|---|---|---|---|---|---|---|---|
| 114 | db/MOUSE/ur | UP00196_1 | Hoxa4_3426.1 | DDTTATTAAT | 17 | 12 | 8.1e-008 | CENTRIMO | http://the_brain.bwh.harvard.edu/uniprobe/details2.php?id=00196 |
| 115 | db/MOUSE/ur | UP00144_1 | Hoxb4_2627.1 | CNNRTTAATT | 17 | 12 | 8.1e-008 | CENTRIMO | http://the_brain.bwh.harvard.edu/uniprobe/details2.php?id=00144 |
| 116 | db/MOUSE/ur | UP00113_1 | Hoxc4_3491.1 | CRMRTTAAT | 17 | 12 | 8.1e-008 | CENTRIMO | http://the_brain.bwh.harvard.edu/uniprobe/details2.php?id=00113 |
| 117 | db/MOUSE/ur | UP00252_1 | Hoxc5_2630.2 | CGMRTTAAT | 17 | 12 | 8.1e-008 | CENTRIMO | http://the_brain.bwh.harvard.edu/uniprobe/details2.php?id=00252 |
| 118 | db/MOUSE/ur | UP00262_1 | Lhx1_2240.2 | VGWATTAAT | 17 | 12 | 8.1e-008 | CENTRIMO | http://the_brain.bwh.harvard.edu/uniprobe/details2.php?id=00262 |
| 119 | db/MOUSE/ur | UP00188_1 | Lmx1a_2238.2 | CGWWTTAAT | 17 | 12 | 8.1e-008 | CENTRIMO | http://the_brain.bwh.harvard.edu/uniprobe/details2.php?id=00188 |
| 120 | db/MOUSE/ur | UP00237_1 | Otp_3496.1 | VVYWRTTAA | 17 | 12 | 8.1e-008 | CENTRIMO | http://the_brain.bwh.harvard.edu/uniprobe/details2.php?id=00237 |
| 121 | db/MOUSE/ur | UP00248_1 | Pax7_3783.1 | MSHNYTAAT | 17 | 12 | 8.1e-008 | CENTRIMO | http://the_brain.bwh.harvard.edu/uniprobe/details2.php?id=00248 |
| 122 | db/MOUSE/ur | UP00128_1 | Pou3f2_2824. | SNWAATTAAT | 17 | 12 | 8.1e-008 | CENTRIMO | http://the_brain.bwh.harvard.edu/uniprobe/details2.php?id=00128 |
| 123 | db/MOUSE/ur | UP00172_1 | Prop1_3949.1 | VGVRTTAATT | 17 | 12 | 8.1e-008 | CENTRIMO | http://the_brain.bwh.harvard.edu/uniprobe/details2.php?id=00172 |
| 124 | db/MOUSE/ur | UP00141_1 | Vsx1_1728.1 | CSARTTAATT | 17 | 12 | 8.1e-008 | CENTRIMO | http://the_brain.bwh.harvard.edu/uniprobe/details2.php?id=00141 |
| 125 | db/EUKARYO | Lhx4_DBD | | YTAATTAR | 8 | 12 | 8.2e-008 | CENTRIMO | http://floresta.eead.csic.es/footprintdb/index.php?db=HumanTF:1.0&motif=Lhx4_DBD |
| 126 | db/JASPAR/J/ | MA0704.1 | Lhx4 | YTAATTAR | 8 | 12 | 8.2e-008 | CENTRIMO | http://jaspar2018.genereg.net/matrix/MA0704.1 |
| 127 | db/EUKARYO | TEF_DBD | | BRTTACRTA | 12 | 12 | 8.3e-008 | CENTRIMO | http://floresta.eead.csic.es/footprintdb/index.php?db=HumanTF:1.0&motif=TEF_DBD |
| 128 | db/EUKARYO | TEF_FL | | NRTTACRTA | 12 | 12 | 8.3e-008 | CENTRIMO | http://floresta.eead.csic.es/footprintdb/index.php?db=HumanTF:1.0&motif=TEF_FL |
| 129 | db/JASPAR/J/ | MA0843.1 | TEF | BRTTACRTA | 12 | 12 | 8.3e-008 | CENTRIMO | http://jaspar2018.genereg.net/matrix/MA0843.1 |
| 130 | db/EUKARYO | EMX1_DBD_2 | | YAATTARYTA | 14 | 12 | 8.3e-008 | CENTRIMO | http://floresta.eead.csic.es/footprintdb/index.php?db=HumanTF:1.0&motif=EMX1_DBD_2 |
| 131 | db/EUKARYO | EMX2_DBD_2 | | TAATTARYTA | 14 | 12 | 8.3e-008 | CENTRIMO | http://floresta.eead.csic.es/footprintdb/index.php?db=HumanTF:1.0&motif=EMX2_DBD_2 |
| 132 | db/EUKARYO | POU4F3_DBD | | RTGMATWAT | 16 | 12 | 8.4e-008 | CENTRIMO | http://floresta.eead.csic.es/footprintdb/index.php?db=HumanTF:1.0&motif=POU4F3_DBD |
| 133 | db/JASPAR/J/ | MA0791.1 | POU4F3 | RTGMATWAT | 16 | 12 | 8.4e-008 | CENTRIMO | http://jaspar2018.genereg.net/matrix/MA0791.1 |
| 134 | db/MOUSE/ur | UP00078_1 | Arid3a_primar | SNNHTTAATT | 17 | 12 | 8.7e-008 | CENTRIMO | http://the_brain.bwh.harvard.edu/uniprobe/details2.php?id=00078 |
| 135 | db/MOUSE/ur | UP00130_1 | Lhx3_3431.1 | VNAATTAATT | 17 | 12 | 8.7e-008 | CENTRIMO | http://the_brain.bwh.harvard.edu/uniprobe/details2.php?id=00130 |
| 136 | db/MOUSE/ur | UP00212_1 | Lhx5_2279.1 | VNAATTAATT | 17 | 12 | 8.7e-008 | CENTRIMO | http://the_brain.bwh.harvard.edu/uniprobe/details2.php?id=00212 |
| 137 | db/MOUSE/ur | UP00169_1 | Lmx1b_3433.2 | VDWWWTTA | 17 | 12 | 8.7e-008 | CENTRIMO | http://the_brain.bwh.harvard.edu/uniprobe/details2.php?id=00169 |
| 138 | db/MOUSE/ur | UP00200_1 | Nkx6-1_2825. | GVWAATTAA | 17 | 12 | 8.7e-008 | CENTRIMO | http://the_brain.bwh.harvard.edu/uniprobe/details2.php?id=00200 |
| 139 | db/MOUSE/ur | UP00149_1 | Phox2b_3948 | VVNAATTAAT | 17 | 12 | 8.7e-008 | CENTRIMO | http://the_brain.bwh.harvard.edu/uniprobe/details2.php?id=00149 |
| 140 | db/MOUSE/ur | UP00142_1 | Uncx4.1_2281 | VNTAATTAAT | 17 | 12 | 8.7e-008 | CENTRIMO | http://the_brain.bwh.harvard.edu/uniprobe/details2.php?id=00142 |
| 141 | db/MOUSE/ur | UP00029_1 | Tbp_primary | NYHWTWTAT | 16 | 12 | 9.0e-008 | CENTRIMO | http://the_brain.bwh.harvard.edu/uniprobe/details2.php?id=00029 |
| 142 | db/MOUSE/ur | UP00234_1 | Msx1_3031.2 | NRMHAYTAA | 16 | 12 | 9.0e-008 | CENTRIMO | http://the_brain.bwh.harvard.edu/uniprobe/details2.php?id=00234 |
| 143 | db/MOUSE/ur | UP00171_1 | Msx3_3206.1 | VAHAACYAAT | 16 | 12 | 9.0e-008 | CENTRIMO | http://the_brain.bwh.harvard.edu/uniprobe/details2.php?id=00171 |
| 144 | db/MOUSE/ur | UP00221_1 | Phox2a_3947 | NNNHAYTAAT | 16 | 12 | 9.0e-008 | CENTRIMO | http://the_brain.bwh.harvard.edu/uniprobe/details2.php?id=00221 |
| 145 | db/MOUSE/ur | UP00106_1 | Vax2_3500.1 | NDNCRCTAA | 16 | 12 | 9.0e-008 | CENTRIMO | http://the_brain.bwh.harvard.edu/uniprobe/details2.php?id=00106 |
| 146 | db/JASPAR/J/ | MA0025.1 | NFIL3 | TTAYGTAAYN | 11 | 12 | 9.2e-008 | CENTRIMO | http://jaspar2018.genereg.net/matrix/MA0025.1 |
| 147 | db/JASPAR/J/ | MA0601.1 | Arid3b | ATATTAATWA | 11 | 12 | 9.2e-008 | CENTRIMO | http://jaspar2018.genereg.net/matrix/MA0601.1 |
| 148 | db/EUKARYO | CART1_DBD | | NTAATYYAAT | 13 | 12 | 9.3e-008 | CENTRIMO | http://floresta.eead.csic.es/footprintdb/index.php?db=HumanTF:1.0&motif=CART1_DBD |
| 149 | db/MOUSE/ur | UP00225_1 | Hlx1_2350.1 | CMATAATTAA | 15 | 12 | 9.3e-008 | CENTRIMO | http://the_brain.bwh.harvard.edu/uniprobe/details2.php?id=00225 |
| 150 | db/JASPAR/J/ | MA0910.1 | Hoxd8 | TAADTAATTA | 17 | 12 | 9.4e-008 | CENTRIMO | http://jaspar2018.genereg.net/matrix/MA0910.1 |
| 151 | db/MOUSE/ur | UP00168_1 | Hoxd8_2644.1 | TAADTAATTA | 17 | 12 | 9.4e-008 | CENTRIMO | http://the_brain.bwh.harvard.edu/uniprobe/details2.php?id=00168 |
| 152 | db/MOUSE/ur | UP00244_1 | Tlx2_3498.2 | KAATYAATTA | 17 | 12 | 9.4e-008 | CENTRIMO | http://the_brain.bwh.harvard.edu/uniprobe/details2.php?id=00244 |
| 153 | db/EUKARYO | POU4F2_DBD | | RTGMATAATT | 16 | 12 | 1.0e-007 | CENTRIMO | http://floresta.eead.csic.es/footprintdb/index.php?db=HumanTF:1.0&motif=POU4F2_DBD |
| 154 | db/JASPAR/J/ | MA0898.1 | Hmx3 | ASAAGCAAT | 17 | 12 | 1.1e-007 | CENTRIMO | http://jaspar2018.genereg.net/matrix/MA0898.1 |
| 155 | db/MOUSE/ur | UP00157_1 | Hmx3_3490.2 | ASAAGCAAT | 17 | 12 | 1.1e-007 | CENTRIMO | http://the_brain.bwh.harvard.edu/uniprobe/details2.php?id=00157 |
| 156 | DREME | GAGTTCMA | DREME-3 | GAGTTCMA | 8 | 62 | 1.1e-007 | DREME | |
| 157 | db/EUKARYO | HOMEZ_DBD | | AAAACGATT | 12 | 12 | 1.2e-007 | CENTRIMO | http://floresta.eead.csic.es/footprintdb/index.php?db=HumanTF:1.0&motif=HOMEZ_DBD |
| 158 | db/EUKARYO | ESRRA_DBD_6 | | CAAGGTCAH | 19 | 12 | 1.4e-007 | CENTRIMO | http://floresta.eead.csic.es/footprintdb/index.php?db=HumanTF:1.0&motif=ESRRA_DBD_6 |
| 159 | db/JASPAR/J/ | MA0065.2 | Pparg::Rxra | STRGGGCAR | 15 | 12 | 1.5e-007 | CENTRIMO | http://jaspar2018.genereg.net/matrix/MA0065.2 |
| 160 | db/EUKARYO | Irx3_DBD | | YWACATGAM | 12 | 12 | 1.7e-007 | CENTRIMO | http://floresta.eead.csic.es/footprintdb/index.php?db=HumanTF:1.0&motif=Irx3_DBD |
| 161 | db/EUKARYO | NR2F1_DBD_1 | | RRGGTCAAA | 15 | 12 | 3.5e-007 | CENTRIMO | http://floresta.eead.csic.es/footprintdb/index.php?db=HumanTF:1.0&motif=NR2F1_DBD_1 |
| 162 | db/EUKARYO | HNF4A_full_2 | | GGGGTCAAA | 14 | 12 | 3.6e-007 | CENTRIMO | http://floresta.eead.csic.es/footprintdb/index.php?db=HumanTF:1.0&motif=HNF4A_full_2 |
| 163 | db/EUKARYO | NR2F6_full | | RRGGTCAAA | 14 | 12 | 3.6e-007 | CENTRIMO | http://floresta.eead.csic.es/footprintdb/index.php?db=HumanTF:1.0&motif=NR2F6_full |
| 164 | db/EUKARYO | NR2F1_DBD_2 | | RRGGTCAAR | 16 | 12 | 3.6e-007 | CENTRIMO | http://floresta.eead.csic.es/footprintdb/index.php?db=HumanTF:1.0&motif=NR2F1_DBD_2 |
| 165 | db/EUKARYO | RARA_full_1 | | ARGGTCAAA | 15 | 12 | 3.7e-007 | CENTRIMO | http://floresta.eead.csic.es/footprintdb/index.php?db=HumanTF:1.0&motif=RARA_full_1 |
| 166 | db/EUKARYO | SPDEF_full_2 | | GTGGTCCCG | 15 | 12 | 5.1e-007 | CENTRIMO | http://floresta.eead.csic.es/footprintdb/index.php?db=HumanTF:1.0&motif=SPDEF_full_2 |
| 167 | db/EUKARYO | ZNF524_full_2 | | CTCGAACCC | 14 | 12 | 5.6e-007 | CENTRIMO | http://floresta.eead.csic.es/footprintdb/index.php?db=HumanTF:1.0&motif=ZNF524_full_2 |
| 168 | db/JASPAR/J/ | MA0597.1 | THAP1 | YTGCCCDBA | 9 | 12 | 6.9e-007 | CENTRIMO | http://jaspar2018.genereg.net/matrix/MA0597.1 |
| 169 | db/EUKARYO | NR2F1_DBD_3 | | CARAGGTCA | 13 | 12 | 1.4e-006 | CENTRIMO | http://floresta.eead.csic.es/footprintdb/index.php?db=HumanTF:1.0&motif=NR2F1_DBD_3 |
| 170 | db/JASPAR/J/ | MA0017.2 | NR2F1 | CARAGGTCA | 13 | 12 | 1.4e-006 | CENTRIMO | http://jaspar2018.genereg.net/matrix/MA0017.2 |
| 171 | db/MOUSE/ur | UP00079_2 | Esrra_second | VKNNAGGGC | 17 | 12 | 1.5e-006 | CENTRIMO | http://the_brain.bwh.harvard.edu/uniprobe/details2.php?id=00079 |
| 172 | db/JASPAR/J/ | MA1112.1 | NR4A1 | NAAAGGTCA | 10 | 12 | 1.5e-006 | CENTRIMO | http://jaspar2018.genereg.net/matrix/MA1112.1 |
| 173 | db/MOUSE/ur | UP00009_1 | Nr2f2_primary | NHBNAAAGG | 16 | 12 | 1.5e-006 | CENTRIMO | http://the_brain.bwh.harvard.edu/uniprobe/details2.php?id=00009 |
| 174 | db/MOUSE/ur | UP00048_1 | Rara_primary | NHBMAAAGG | 16 | 12 | 1.5e-006 | CENTRIMO | http://the_brain.bwh.harvard.edu/uniprobe/details2.php?id=00048 |

| | | | | | | | | | | |
|---|---|---|---|---|---|---|---|---|---|---|
| 175 | db/JASPAR/J/ | MA0071.1 | RORA | AWMWAGGT | 10 | 12 | 1.6e-006 | CENTRIMO | | http://jaspar2018.genereg.net/matrix/MA0071.1 |
| 176 | DREME | AAAAHAAA | DREME-4 | AAAAHAAA | 8 | 187 | 3.2e-006 | DREME | db/JASPAR/JASPAR **MA1125.1 (ZNF384)** | http://jaspar2018.genereg.net/matrix/MA1125.1 |
| 177 | db/JASPAR/J/ | MA1120.1 | SOX13 | DVACAATGG | 11 | 12 | 4.4e-006 | CENTRIMO | | http://jaspar2018.genereg.net/matrix/MA1120.1 |
| 178 | db/JASPAR/J/ | MA0143.3 | Sox2 | CCWTTGTY | 8 | 12 | 4.5e-006 | CENTRIMO | | http://jaspar2018.genereg.net/matrix/MA0143.3 |
| 179 | db/EUKARYO | SOX9_DBD | | DAACAATRG | 9 | 12 | 4.6e-006 | CENTRIMO | | http://floresta.eead.csic.es/footprintdb/index.php?db=HumanTF:1.0&motif=SOX9_DBD |
| 180 | db/JASPAR/J/ | MA0077.1 | SOX9 | CYATTGTTY | 9 | 12 | 4.6e-006 | CENTRIMO | | http://jaspar2018.genereg.net/matrix/MA0077.1 |
| 181 | db/JASPAR/J/ | MA0442.2 | SOX10 | NDAACAAAG | 11 | 12 | 4.6e-006 | CENTRIMO | | http://jaspar2018.genereg.net/matrix/MA0442.2 |
| 182 | db/MOUSE/ur | UP00004_2 | Sox14_secon | NKBASACAA | 15 | 12 | 4.7e-006 | CENTRIMO | | http://the_brain.bwh.harvard.edu/uniprobe/details2.php?id=00004 |
| 183 | db/MOUSE/ur | UP00030_1 | Sox11_primar | HNDARAACA | 17 | 12 | 4.7e-006 | CENTRIMO | | http://the_brain.bwh.harvard.edu/uniprobe/details2.php?id=00030 |
| 184 | db/MOUSE/ur | UP00062_1 | Sox4_primary | HNNWRAAC | 17 | 12 | 4.7e-006 | CENTRIMO | | http://the_brain.bwh.harvard.edu/uniprobe/details2.php?id=00062 |
| 185 | db/JASPAR/J/ | MA0515.1 | Sox6 | CCWTTGTYY | 10 | 12 | 4.7e-006 | CENTRIMO | | http://jaspar2018.genereg.net/matrix/MA0515.1 |
| 186 | db/JASPAR/J/ | MA0514.1 | Sox3 | CCTTTGTYY | 10 | 12 | 4.7e-006 | CENTRIMO | | http://jaspar2018.genereg.net/matrix/MA0514.1 |
| 187 | db/JASPAR/J/ | MA1152.1 | SOX15 | CYWTTGTTH | 10 | 12 | 4.7e-006 | CENTRIMO | | http://jaspar2018.genereg.net/matrix/MA1152.1 |
| 188 | db/JASPAR/J/ | MA0084.1 | SRY | NWWAACAAT | 9 | 12 | 5.1e-006 | CENTRIMO | | http://jaspar2018.genereg.net/matrix/MA0084.1 |
| 189 | db/EUKARYO | FOXJ2_DBD_2 | | RTAAACAA | 8 | 12 | 5.2e-006 | CENTRIMO | | http://floresta.eead.csic.es/footprintdb/index.php?db=HumanTF:1.0&motif=FOXJ2_DBD_2 |
| 190 | db/EUKARYO | FOXJ3_DBD_1 | | RTAAACAA | 8 | 12 | 5.2e-006 | CENTRIMO | | http://floresta.eead.csic.es/footprintdb/index.php?db=HumanTF:1.0&motif=FOXJ3_DBD_1 |
| 191 | db/EUKARYO | FOXO1_DBD_1 | | GTAAACAW | 8 | 12 | 5.2e-006 | CENTRIMO | | http://floresta.eead.csic.es/footprintdb/index.php?db=HumanTF:1.0&motif=FOXO1_DBD_1 |
| 192 | db/EUKARYO | Foxj3_DBD_3 | | RTAAACAA | 8 | 12 | 5.2e-006 | CENTRIMO | | http://floresta.eead.csic.es/footprintdb/index.php?db=HumanTF:1.0&motif=Foxj3_DBD_3 |
| 193 | db/JASPAR/J/ | MA0852.2 | FOXK1 | NRDGTAAAC | 14 | 12 | 5.3e-006 | CENTRIMO | | http://jaspar2018.genereg.net/matrix/MA0852.2 |
| 194 | db/JASPAR/J/ | MA0593.1 | FOXP2 | RWGTAAACA | 11 | 12 | 5.4e-006 | CENTRIMO | | http://jaspar2018.genereg.net/matrix/MA0593.1 |
| 195 | db/JASPAR/J/ | MA0486.2 | HSF1 | TTCTAGAAYF | 13 | 12 | 6.0e-006 | CENTRIMO | | http://jaspar2018.genereg.net/matrix/MA0486.2 |
| 196 | db/MOUSE/ur | UP00089_1 | Tcf1_primary | NHTHRGTTA | 17 | 12 | 1.0e-005 | CENTRIMO | | http://the_brain.bwh.harvard.edu/uniprobe/details2.php?id=00089 |
| 197 | db/EUKARYO | RUNX3_DBD_3 | | HRACCGCAF | 18 | 12 | 1.1e-005 | CENTRIMO | | http://floresta.eead.csic.es/footprintdb/index.php?db=HumanTF:1.0&motif=RUNX3_DBD_3 |
| 198 | DREME | GGACAGCC | DREME-12 | GGACAGCC | 8 | 24 | 1.2e-005 | DREME | | |
| 199 | DREME | RGAA | DREME-5 | RGAA | 4 | 3608 | 1.8e-005 | DREME | | |
| 200 | db/MOUSE/ur | UP00090_2 | Elf3_seconda | GBTCHAAAA | 17 | 12 | 1.9e-005 | CENTRIMO | | http://the_brain.bwh.harvard.edu/uniprobe/details2.php?id=00090 |
| 201 | db/MOUSE/ur | UP00077_2 | Srf_secondary | STWNAAAAA | 17 | 12 | 1.9e-005 | CENTRIMO | | http://the_brain.bwh.harvard.edu/uniprobe/details2.php?id=00077 |
| 202 | db/JASPAR/J/ | MA1125.1 | ZNF384 | DNWMAAAA | 12 | 12 | 2.0e-005 | CENTRIMO | | http://jaspar2018.genereg.net/matrix/MA1125.1 |
| 203 | db/MOUSE/ur | UP00070_2 | Gcm1_second | DKSNNATAG | 17 | 12 | 2.1e-005 | CENTRIMO | | http://the_brain.bwh.harvard.edu/uniprobe/details2.php?id=00070 |
| 204 | db/MOUSE/ur | UP00058_2 | Tcf3_seconda | NSSSBAAWA | 15 | 12 | 2.2e-005 | CENTRIMO | | http://the_brain.bwh.harvard.edu/uniprobe/details2.php?id=00058 |
| 205 | db/EUKARYO | TEAD3_DBD_2 | | RCATTCCW | 8 | 12 | 2.4e-005 | CENTRIMO | | http://floresta.eead.csic.es/footprintdb/index.php?db=HumanTF:1.0&motif=TEAD3_DBD_2 |
| 206 | db/JASPAR/J/ | MA0808.1 | TEAD3 | RCATTCCW | 8 | 12 | 2.4e-005 | CENTRIMO | | http://jaspar2018.genereg.net/matrix/MA0808.1 |
| 207 | db/EUKARYO | TEAD1_full_1 | | YRCATTCCW | 10 | 12 | 2.4e-005 | CENTRIMO | | http://floresta.eead.csic.es/footprintdb/index.php?db=HumanTF:1.0&motif=TEAD1_full_1 |
| 208 | db/EUKARYO | TEAD4_DBD | | NRCATTCCW | 10 | 12 | 2.4e-005 | CENTRIMO | | http://floresta.eead.csic.es/footprintdb/index.php?db=HumanTF:1.0&motif=TEAD4_DBD |
| 209 | db/JASPAR/J/ | MA0090.2 | TEAD1 | YRCATTCCW | 10 | 12 | 2.4e-005 | CENTRIMO | | http://jaspar2018.genereg.net/matrix/MA0090.2 |
| 210 | db/JASPAR/J/ | MA0809.1 | TEAD4 | NRCATTCCW | 10 | 12 | 2.4e-005 | CENTRIMO | | http://jaspar2018.genereg.net/matrix/MA0809.1 |
| 211 | db/JASPAR/J/ | MA0057.1 | MZF1(var.2) | BKAGGGGKA | 10 | 12 | 2.8e-005 | CENTRIMO | | http://jaspar2018.genereg.net/matrix/MA0057.1 |
| 212 | db/MOUSE/ur | UP00040_2 | Irf5_secondar | BWNAYCGAC | 15 | 12 | 3.4e-005 | CENTRIMO | | http://the_brain.bwh.harvard.edu/uniprobe/details2.php?id=00040 |
| 213 | db/EUKARYO | IRF5_full_1 | | CCGAAACCG | 14 | 12 | 4.4e-005 | CENTRIMO | | http://floresta.eead.csic.es/footprintdb/index.php?db=HumanTF:1.0&motif=IRF5_full_1 |
| 214 | db/JASPAR/J/ | MA1420.1 | IRF5 | CCGAAACCG | 14 | 12 | 4.4e-005 | CENTRIMO | | http://jaspar2018.genereg.net/matrix/MA1420.1 |
| 215 | db/EUKARYO | TFCP2_full_1 | | AAACCGGTT | 10 | 12 | 4.7e-005 | CENTRIMO | | http://floresta.eead.csic.es/footprintdb/index.php?db=HumanTF:1.0&motif=TFCP2_full_1 |
| 216 | db/JASPAR/J/ | MA0092.1 | Hand1::Tcf3 | BRTCTGGMW | 10 | 12 | 4.7e-005 | CENTRIMO | | http://jaspar2018.genereg.net/matrix/MA0092.1 |
| 217 | db/JASPAR/J/ | MA0145.3 | TFCP2 | AAACCGGTT | 10 | 12 | 4.7e-005 | CENTRIMO | | http://jaspar2018.genereg.net/matrix/MA0145.3 |
| 218 | db/EUKARYO | GRHL1_DBD_2 | | AAACCGGTT | 10 | 12 | 5.9e-005 | CENTRIMO | | http://floresta.eead.csic.es/footprintdb/index.php?db=HumanTF:1.0&motif=GRHL1_DBD_2 |
| 219 | db/JASPAR/J/ | MA1105.1 | GRHL2 | GNMAAACCA | 15 | 12 | 6.1e-005 | CENTRIMO | | http://jaspar2018.genereg.net/matrix/MA1105.1 |
| 220 | db/MOUSE/ur | UP00034_2 | Sox7_second | GTVSWAATT | 22 | 12 | 6.6e-005 | CENTRIMO | | http://the_brain.bwh.harvard.edu/uniprobe/details2.php?id=00034 |
| 221 | db/EUKARYO | GRHL1_DBD_1 | | AACCGGTYT | 17 | 12 | 7.2e-005 | CENTRIMO | | http://floresta.eead.csic.es/footprintdb/index.php?db=HumanTF:1.0&motif=GRHL1_DBD_1 |
| 222 | db/EUKARYO | MYBL1_DBD_1 | | ACCGTTAAA | 12 | 12 | 7.3e-005 | CENTRIMO | | http://floresta.eead.csic.es/footprintdb/index.php?db=HumanTF:1.0&motif=MYBL1_DBD_1 |
| 223 | DREME | CTACARAG | DREME-6 | CTACARAG | 8 | 36 | 7.6e-005 | DREME | | |
| 224 | db/JASPAR/J/ | MA0066.1 | PPARG | STAGGTCAC | 20 | 12 | 8.4e-005 | CENTRIMO | | http://jaspar2018.genereg.net/matrix/MA0066.1 |
| 225 | db/EUKARYO | TFEC_DBD | | RTCACRTGA | 10 | 12 | 1.7e-004 | CENTRIMO | | http://floresta.eead.csic.es/footprintdb/index.php?db=HumanTF:1.0&motif=TFEC_DBD |
| 226 | db/JASPAR/J/ | MA0871.1 | TFEC | RTCACRTGA | 10 | 12 | 1.7e-004 | CENTRIMO | | http://jaspar2018.genereg.net/matrix/MA0871.1 |
| 227 | db/JASPAR/J/ | MA0620.2 | MITF | NNDRGTCAC | 18 | 12 | 1.7e-004 | CENTRIMO | | http://jaspar2018.genereg.net/matrix/MA0620.2 |
| 228 | db/MOUSE/ur | UP00050_1 | Bhlhb2_prima | RNHWNNDT | 22 | 12 | 1.7e-004 | CENTRIMO | | http://the_brain.bwh.harvard.edu/uniprobe/details2.php?id=00050 |
| 229 | db/EUKARYO | RARG_full_3 | | RAGGTCAHB | 17 | 12 | 1.8e-004 | CENTRIMO | | http://floresta.eead.csic.es/footprintdb/index.php?db=HumanTF:1.0&motif=RARG_full_3 |
| 230 | db/EUKARYO | POU3F1_DBD_1 | | WTATGCWA | 12 | 12 | 1.8e-004 | CENTRIMO | | http://floresta.eead.csic.es/footprintdb/index.php?db=HumanTF:1.0&motif=POU3F1_DBD_1 |
| 231 | db/JASPAR/J/ | MA0786.1 | POU3F1 | WTATGCWA | 12 | 12 | 1.8e-004 | CENTRIMO | | http://jaspar2018.genereg.net/matrix/MA0786.1 |
| 232 | db/EUKARYO | POU3F4_DBD_1 | | TATGCWAAT | 9 | 12 | 1.9e-004 | CENTRIMO | | http://floresta.eead.csic.es/footprintdb/index.php?db=HumanTF:1.0&motif=POU3F4_DBD_1 |
| 233 | db/EUKARYO | Pou2f2_DBD_2 | | TATGCAAAT | 9 | 12 | 1.9e-004 | CENTRIMO | | http://floresta.eead.csic.es/footprintdb/index.php?db=HumanTF:1.0&motif=Pou2f2_DBD_2 |
| 234 | db/JASPAR/J/ | MA0789.1 | POU3F4 | TATGCWAAT | 9 | 12 | 1.9e-004 | CENTRIMO | | http://jaspar2018.genereg.net/matrix/MA0789.1 |
| 235 | db/EUKARYO | POU2F1_DBD_1 | | AWTATGCWA | 12 | 12 | 1.9e-004 | CENTRIMO | | http://floresta.eead.csic.es/footprintdb/index.php?db=HumanTF:1.0&motif=POU2F1_DBD_1 |

| | | | | | | | | | | | |
|---|---|---|---|---|---|---|---|---|---|---|---|
| 236 | db/JASPAR/J | MA0785.1 | POU2F1 | AWTATGCWA | 12 | 12 | 1.9e-004 | CENTRIMO | | | http://jaspar2018.genereg.net/matrix/MA0785.1 |
| 237 | db/MOUSE/ur | UP00082_1 | Zfp187_prima | NTATGTACTA | 14 | 12 | 1.9e-004 | CENTRIMO | | | http://the_brain.bwh.harvard.edu/uniprobe/details2.php?id=00082 |
| 238 | db/EUKARYO | CUX1_DBD_2 | | ATCGATHMN | 18 | 12 | 1.9e-004 | CENTRIMO | | | http://floresta.eead.csic.es/footprintdb/index.php?db=HumanTF:1.0&motif=CUX1_DBD_2 |
| 239 | db/EUKARYO | CUX2_DBD_1 | | ATCRATAAWI | 18 | 12 | 1.9e-004 | CENTRIMO | | | http://floresta.eead.csic.es/footprintdb/index.php?db=HumanTF:1.0&motif=CUX2_DBD_1 |
| 240 | db/JASPAR/J | MA0609.1 | Crem | NATGACGTM | 10 | 12 | 2.0e-004 | CENTRIMO | | | http://jaspar2018.genereg.net/matrix/MA0609.1 |
| 241 | db/EUKARYO | XBP1_DBD_2 | | WDKGMCAC | 14 | 12 | 2.0e-004 | CENTRIMO | | | http://floresta.eead.csic.es/footprintdb/index.php?db=HumanTF:1.0&motif=XBP1_DBD_2 |
| 242 | db/JASPAR/J | MA0844.1 | XBP1 | WDKGMCAC | 14 | 12 | 2.0e-004 | CENTRIMO | | | http://jaspar2018.genereg.net/matrix/MA0844.1 |
| 243 | db/JASPAR/J | MA0615.1 | Gmeb1 | BHBBKKACG | 17 | 12 | 2.1e-004 | CENTRIMO | | | http://jaspar2018.genereg.net/matrix/MA0615.1 |
| 244 | db/MOUSE/ur | UP00084_1 | Gmeb1_prima | BHBBKKACG | 17 | 12 | 2.1e-004 | CENTRIMO | | | http://the_brain.bwh.harvard.edu/uniprobe/details2.php?id=00084 |
| 245 | db/JASPAR/J | MA1129.1 | FOSL1::JUN( | ATGACGTCA | 10 | 12 | 2.1e-004 | CENTRIMO | | | http://jaspar2018.genereg.net/matrix/MA1129.1 |
| 246 | db/JASPAR/J | MA1136.1 | FOSB::JUNB( | RTGACGTCA | 10 | 12 | 2.1e-004 | CENTRIMO | | | http://jaspar2018.genereg.net/matrix/MA1136.1 |
| 247 | db/JASPAR/J | MA1140.1 | JUNB(var.2) | RTGACGTCA | 10 | 12 | 2.1e-004 | CENTRIMO | | | http://jaspar2018.genereg.net/matrix/MA1140.1 |
| 248 | db/JASPAR/J | MA1143.1 | FOSL1::JUND | RTGACGTMA | 10 | 12 | 2.1e-004 | CENTRIMO | | | http://jaspar2018.genereg.net/matrix/MA1143.1 |
| 249 | db/EUKARYO | Creb3l2_DBD_2 | | TGCCACGTG | 12 | 12 | 2.1e-004 | CENTRIMO | | | http://floresta.eead.csic.es/footprintdb/index.php?db=HumanTF:1.0&motif=Creb3l2_DBD_2 |
| 250 | db/EUKARYO | Creb5_DBD | | NATGACGTC | 12 | 12 | 2.1e-004 | CENTRIMO | | | http://floresta.eead.csic.es/footprintdb/index.php?db=HumanTF:1.0&motif=Creb5_DBD |
| 251 | db/EUKARYO | DBP_DBD | | NRTTACGTA | 12 | 12 | 2.1e-004 | CENTRIMO | | | http://floresta.eead.csic.es/footprintdb/index.php?db=HumanTF:1.0&motif=DBP_DBD |
| 252 | db/EUKARYO | Dbp_DBD | | NRTTACGTA | 12 | 12 | 2.1e-004 | CENTRIMO | | | http://floresta.eead.csic.es/footprintdb/index.php?db=HumanTF:1.0&motif=Dbp_DBD |
| 253 | db/EUKARYO | JDP2_DBD_2 | | GATGACGTC | 12 | 12 | 2.1e-004 | CENTRIMO | | | http://floresta.eead.csic.es/footprintdb/index.php?db=HumanTF:1.0&motif=JDP2_DBD_2 |
| 254 | db/EUKARYO | JDP2_full_2 | | GATGACGTC | 12 | 12 | 2.1e-004 | CENTRIMO | | | http://floresta.eead.csic.es/footprintdb/index.php?db=HumanTF:1.0&motif=JDP2_full_2 |
| 255 | db/EUKARYO | Jdp2_DBD_2 | | GATGACGTC | 12 | 12 | 2.1e-004 | CENTRIMO | | | http://floresta.eead.csic.es/footprintdb/index.php?db=HumanTF:1.0&motif=Jdp2_DBD_2 |
| 256 | db/EUKARYO | XBP1_DBD_1 | | GATGACGTC | 12 | 12 | 2.1e-004 | CENTRIMO | | | http://floresta.eead.csic.es/footprintdb/index.php?db=HumanTF:1.0&motif=XBP1_DBD_1 |
| 257 | db/JASPAR/J | MA0656.1 | JDP2(var.2) | GATGACGTC | 12 | 12 | 2.1e-004 | CENTRIMO | | | http://jaspar2018.genereg.net/matrix/MA0656.1 |
| 258 | db/JASPAR/J | MA0840.1 | Creb5 | NATGACGTC | 12 | 12 | 2.1e-004 | CENTRIMO | | | http://jaspar2018.genereg.net/matrix/MA0840.1 |
| 259 | db/JASPAR/J | MA0018.3 | CREB1 | NVTGACGTC | 12 | 12 | 2.1e-004 | CENTRIMO | | | http://jaspar2018.genereg.net/matrix/MA0018.3 |
| 260 | db/JASPAR/J | MA1133.1 | JUN::JUNB(va | KRTGACGTC | 12 | 12 | 2.1e-004 | CENTRIMO | | | http://jaspar2018.genereg.net/matrix/MA1133.1 |
| 261 | db/JASPAR/J | MA1139.1 | FOSL2::JUNB | DATGACGTC | 12 | 12 | 2.1e-004 | CENTRIMO | | | http://jaspar2018.genereg.net/matrix/MA1139.1 |
| 262 | db/EUKARYO | ATF7_DBD | | NDATGACGT | 14 | 12 | 2.1e-004 | CENTRIMO | | | http://floresta.eead.csic.es/footprintdb/index.php?db=HumanTF:1.0&motif=ATF7_DBD |
| 263 | db/EUKARYO | BATF3_DBD | | TGATGACGT | 14 | 12 | 2.1e-004 | CENTRIMO | | | http://floresta.eead.csic.es/footprintdb/index.php?db=HumanTF:1.0&motif=BATF3_DBD |
| 264 | db/EUKARYO | CREB3_full_1 | | BNRTGACGT | 14 | 12 | 2.1e-004 | CENTRIMO | | | http://floresta.eead.csic.es/footprintdb/index.php?db=HumanTF:1.0&motif=CREB3_full_1 |
| 265 | db/EUKARYO | CREB3_full_2 | | VTGCCACGT | 14 | 12 | 2.1e-004 | CENTRIMO | | | http://floresta.eead.csic.es/footprintdb/index.php?db=HumanTF:1.0&motif=CREB3_full_2 |
| 266 | db/JASPAR/J | MA0638.1 | CREB3 | VTGCCACGT | 14 | 12 | 2.1e-004 | CENTRIMO | | | http://jaspar2018.genereg.net/matrix/MA0638.1 |
| 267 | db/JASPAR/J | MA0834.1 | ATF7 | NDATGACGT | 14 | 12 | 2.1e-004 | CENTRIMO | | | http://jaspar2018.genereg.net/matrix/MA0834.1 |
| 268 | db/JASPAR/J | MA0835.1 | BATF3 | TGATGACGT | 14 | 12 | 2.1e-004 | CENTRIMO | | | http://jaspar2018.genereg.net/matrix/MA0835.1 |
| 269 | db/EUKARYO | TBX21_DBD_1 | | GGTGTGAHV | 16 | 12 | 2.1e-004 | CENTRIMO | | | http://floresta.eead.csic.es/footprintdb/index.php?db=HumanTF:1.0&motif=TBX21_DBD_1 |
| 270 | db/MOUSE/ur | UP00020_1 | Atf1_primary | DYKRTGACG | 16 | 12 | 2.1e-004 | CENTRIMO | | | http://the_brain.bwh.harvard.edu/uniprobe/details2.php?id=00020 |
| 271 | db/MOUSE/ur | UP00103_1 | Jundm2_prim | NSGATGACG | 16 | 12 | 2.1e-004 | CENTRIMO | | | http://the_brain.bwh.harvard.edu/uniprobe/details2.php?id=00103 |
| 272 | db/JASPAR/J | MA1127.1 | FOSB::JUN | GATGACGTC | 11 | 12 | 2.1e-004 | CENTRIMO | | | http://jaspar2018.genereg.net/matrix/MA1127.1 |
| 273 | db/JASPAR/J | MA1131.1 | FOSL2::JUN( | GRTGACGTM | 11 | 12 | 2.1e-004 | CENTRIMO | | | http://jaspar2018.genereg.net/matrix/MA1131.1 |
| 274 | db/JASPAR/J | MA1145.1 | FOSL2::JUND | NNRTGACGT | 15 | 12 | 2.1e-004 | CENTRIMO | | | http://jaspar2018.genereg.net/matrix/MA1145.1 |
| 275 | db/MOUSE/ur | UP00223_2 | Irx3_2226.1 | WDDNTACAT | 17 | 12 | 2.1e-004 | CENTRIMO | | | http://the_brain.bwh.harvard.edu/uniprobe/details2.php?id=00223 |
| 276 | db/MOUSE/ur | UP00194_1 | Irx4_2242.3 | WNNNWACA | 17 | 12 | 2.1e-004 | CENTRIMO | | | http://the_brain.bwh.harvard.edu/uniprobe/details2.php?id=00194 |
| 277 | db/MOUSE/ur | UP00250_1 | Irx5_2385.1 | WWNNWACA | 17 | 12 | 2.1e-004 | CENTRIMO | | | http://the_brain.bwh.harvard.edu/uniprobe/details2.php?id=00250 |
| 278 | db/MOUSE/ur | UP00150_1 | Irx6_2623.2 | HAWNTACAT | 17 | 12 | 2.1e-004 | CENTRIMO | | | http://the_brain.bwh.harvard.edu/uniprobe/details2.php?id=00150 |
| 279 | db/EUKARYO | POU5F1P1_DBD_2 | | ATGMATATK | 12 | 12 | 2.2e-004 | CENTRIMO | | | http://floresta.eead.csic.es/footprintdb/index.php?db=HumanTF:1.0&motif=POU5F1P1_DBD_2 |
| 280 | db/EUKARYO | FOXC1_DBD_2 | | DGTMAATAT | 14 | 12 | 2.2e-004 | CENTRIMO | | | http://floresta.eead.csic.es/footprintdb/index.php?db=HumanTF:1.0&motif=FOXC1_DBD_2 |
| 281 | db/EUKARYO | FOXC2_DBD_1 | | DGTMAATAT | 14 | 12 | 2.2e-004 | CENTRIMO | | | http://floresta.eead.csic.es/footprintdb/index.php?db=HumanTF:1.0&motif=FOXC2_DBD_1 |
| 282 | db/EUKARYO | POU2F1_DBD_2 | | HATGMATAT | 14 | 12 | 2.2e-004 | CENTRIMO | | | http://floresta.eead.csic.es/footprintdb/index.php?db=HumanTF:1.0&motif=POU2F1_DBD_2 |
| 283 | db/EUKARYO | POU2F2_DBD_2 | | HWTRMATAT | 14 | 12 | 2.2e-004 | CENTRIMO | | | http://floresta.eead.csic.es/footprintdb/index.php?db=HumanTF:1.0&motif=POU2F2_DBD_2 |
| 284 | db/EUKARYO | Pou2f2_DBD_1 | | YWTGAATAT | 14 | 12 | 2.2e-004 | CENTRIMO | | | http://floresta.eead.csic.es/footprintdb/index.php?db=HumanTF:1.0&motif=Pou2f2_DBD_1 |
| 285 | db/EUKARYO | FOXB1_DBD_2 | | WATGTAAAT | 18 | 12 | 2.2e-004 | CENTRIMO | | | http://floresta.eead.csic.es/footprintdb/index.php?db=HumanTF:1.0&motif=FOXB1_DBD_2 |
| 286 | db/JASPAR/J | MA0488.1 | JUN | DDRATGATG | 13 | 12 | 2.2e-004 | CENTRIMO | | | http://jaspar2018.genereg.net/matrix/MA0488.1 |
| 287 | db/JASPAR/J | MA0492.1 | JUND(var.2) | DDDRATGAB | 15 | 12 | 2.2e-004 | CENTRIMO | | | http://jaspar2018.genereg.net/matrix/MA0492.1 |
| 288 | db/JASPAR/J | MA0605.1 | Atf3 | GATGACGT | 8 | 12 | 2.2e-004 | CENTRIMO | | | http://jaspar2018.genereg.net/matrix/MA0605.1 |
| 289 | db/JASPAR/J | MA1126.1 | FOS::JUN(var | DRTGACGTC | 16 | 12 | 2.3e-004 | CENTRIMO | | | http://jaspar2018.genereg.net/matrix/MA1126.1 |
| 290 | db/EUKARYO | CUX1_DBD_3 | | TRATCRATA | 10 | 12 | 2.4e-004 | CENTRIMO | | | http://floresta.eead.csic.es/footprintdb/index.php?db=HumanTF:1.0&motif=CUX1_DBD_3 |
| 291 | db/EUKARYO | CUX2_DBD_2 | | TRATCGATA | 10 | 12 | 2.4e-004 | CENTRIMO | | | http://floresta.eead.csic.es/footprintdb/index.php?db=HumanTF:1.0&motif=CUX2_DBD_2 |
| 292 | db/JASPAR/J | MA0754.1 | CUX1 | TRATCRATA | 10 | 12 | 2.4e-004 | CENTRIMO | | | http://jaspar2018.genereg.net/matrix/MA0754.1 |
| 293 | db/JASPAR/J | MA0755.1 | CUX2 | TRATCGATA | 10 | 12 | 2.4e-004 | CENTRIMO | | | http://jaspar2018.genereg.net/matrix/MA0755.1 |
| 294 | db/MOUSE/ur | UP00185_1 | Pbx1_3203.1 | HYAYMCATC | 17 | 12 | 2.5e-004 | CENTRIMO | | | http://the_brain.bwh.harvard.edu/uniprobe/details2.php?id=00185 |
| 295 | DREME | CMCGCCC | DREME-8 | CMCGCCC | 7 | 84 | 4.0e-004 | DREME | db/MOUSE/uniprobe | **UP00093_1 (Klf7_prir** | http://the_brain.bwh.harvard.edu/uniprobe/details2.php?id=00093 |
| 296 | db/EUKARYO | RUNX2_DBD_2 | | WRACCGCA | 18 | 12 | 4.1e-004 | CENTRIMO | | | http://floresta.eead.csic.es/footprintdb/index.php?db=HumanTF:1.0&motif=RUNX2_DBD_2 |

| | | | | | | | | | |
|---|---|---|---|---|---|---|---|---|---|
| 297 | db/JASPAR/J | MA0006.1 | Ahr::Arnt | YGCGTG | 6 | 12 | 4.6e-004 | CENTRIMO | http://jaspar2018.genereg.net/matrix/MA0006.1 |
| 298 | db/JASPAR/J | MA0062.2 | Gabpa | CCGGAAGTG | 11 | 12 | 5.7e-004 | CENTRIMO | http://jaspar2018.genereg.net/matrix/MA0062.2 |
| 299 | db/EUKARYO | ELK1_DBD_1 | | ACCGGAAGT | 10 | 12 | 6.0e-004 | CENTRIMO | http://floresta.eead.csic.es/footprintdb/index.php?db=HumanTF:1.0&motif=ELK1_DBD_1 |
| 300 | db/EUKARYO | ELK1_DBD_2 | | ACCGGAAGT | 10 | 12 | 6.0e-004 | CENTRIMO | http://floresta.eead.csic.es/footprintdb/index.php?db=HumanTF:1.0&motif=ELK1_DBD_2 |
| 301 | db/EUKARYO | ELK1_full_1 | | ACCGGAAGT | 10 | 12 | 6.0e-004 | CENTRIMO | http://floresta.eead.csic.es/footprintdb/index.php?db=HumanTF:1.0&motif=ELK1_full_1 |
| 302 | db/EUKARYO | ELK3_DBD | | ACCGGAAGT | 10 | 12 | 6.0e-004 | CENTRIMO | http://floresta.eead.csic.es/footprintdb/index.php?db=HumanTF:1.0&motif=ELK3_DBD |
| 303 | db/EUKARYO | ETV1_DBD | | ACCGGAAGT | 10 | 12 | 6.0e-004 | CENTRIMO | http://floresta.eead.csic.es/footprintdb/index.php?db=HumanTF:1.0&motif=ETV1_DBD |
| 304 | db/EUKARYO | ETV3_DBD | | ACCGGAAGT | 10 | 12 | 6.0e-004 | CENTRIMO | http://floresta.eead.csic.es/footprintdb/index.php?db=HumanTF:1.0&motif=ETV3_DBD |
| 305 | db/EUKARYO | ETV4_DBD | | ACCGGAAGT | 10 | 12 | 6.0e-004 | CENTRIMO | http://floresta.eead.csic.es/footprintdb/index.php?db=HumanTF:1.0&motif=ETV4_DBD |
| 306 | db/EUKARYO | Elk3_DBD | | ACCGGAAGT | 10 | 12 | 6.0e-004 | CENTRIMO | http://floresta.eead.csic.es/footprintdb/index.php?db=HumanTF:1.0&motif=Elk3_DBD |
| 307 | db/EUKARYO | GABPA_full | | ACCGGAAGT | 10 | 12 | 6.0e-004 | CENTRIMO | http://floresta.eead.csic.es/footprintdb/index.php?db=HumanTF:1.0&motif=GABPA_full |
| 308 | db/JASPAR/J | MA0759.1 | ELK3 | ACCGGAAGT | 10 | 12 | 6.0e-004 | CENTRIMO | http://jaspar2018.genereg.net/matrix/MA0759.1 |
| 309 | db/JASPAR/J | MA0761.1 | ETV1 | ACCGGAAGT | 10 | 12 | 6.0e-004 | CENTRIMO | http://jaspar2018.genereg.net/matrix/MA0761.1 |
| 310 | db/JASPAR/J | MA0763.1 | ETV3 | ACCGGAAGT | 10 | 12 | 6.0e-004 | CENTRIMO | http://jaspar2018.genereg.net/matrix/MA0763.1 |
| 311 | db/JASPAR/J | MA0764.1 | ETV4 | ACCGGAAGT | 10 | 12 | 6.0e-004 | CENTRIMO | http://jaspar2018.genereg.net/matrix/MA0764.1 |
| 312 | db/JASPAR/J | MA0028.2 | ELK1 | ACCGGAAGT | 10 | 12 | 6.0e-004 | CENTRIMO | http://jaspar2018.genereg.net/matrix/MA0028.2 |
| 313 | db/EUKARYO | SPDEF_full_1 | | AMCCGGATG | 11 | 12 | 6.1e-004 | CENTRIMO | http://floresta.eead.csic.es/footprintdb/index.php?db=HumanTF:1.0&motif=SPDEF_full_1 |
| 314 | db/JASPAR/J | MA0686.1 | SPDEF | AMCCGGATG | 11 | 12 | 6.1e-004 | CENTRIMO | http://jaspar2018.genereg.net/matrix/MA0686.1 |
| 315 | db/EUKARYO | ERF_DBD | | ACCGGAAGT | 10 | 12 | 6.5e-004 | CENTRIMO | http://floresta.eead.csic.es/footprintdb/index.php?db=HumanTF:1.0&motif=ERF_DBD |
| 316 | db/JASPAR/J | MA0760.1 | ERF | ACCGGAAGT | 10 | 12 | 6.5e-004 | CENTRIMO | http://jaspar2018.genereg.net/matrix/MA0760.1 |
| 317 | db/EUKARYO | SPDEF_DBD_1 | | AMCCGGATG | 11 | 12 | 6.6e-004 | CENTRIMO | http://floresta.eead.csic.es/footprintdb/index.php?db=HumanTF:1.0&motif=SPDEF_DBD_1 |
| 318 | db/MOUSE/ur | UP00018_2 | Irf4_secondar | NNNAYTCTC | 15 | 12 | 6.6e-004 | CENTRIMO | http://the_brain.bwh.harvard.edu/uniprobe/details2.php?id=00018 |
| 319 | db/MOUSE/ur | UP00011_2 | Irf6_secondar | NBBACTCYC | 15 | 12 | 6.6e-004 | CENTRIMO | http://the_brain.bwh.harvard.edu/uniprobe/details2.php?id=00011 |
| 320 | db/MOUSE/ur | UP00013_1 | Gabpa_primar | MNWWACCG | 17 | 12 | 6.7e-004 | CENTRIMO | http://the_brain.bwh.harvard.edu/uniprobe/details2.php?id=00013 |
| 321 | db/EUKARYO | NFATC1_full_2 | | TTTCCAYWR | 15 | 12 | 7.6e-004 | CENTRIMO | http://floresta.eead.csic.es/footprintdb/index.php?db=HumanTF:1.0&motif=NFATC1_full_2 |
| 322 | db/MOUSE/ur | UP00059_2 | Arid5a_secon | NNYNMAATA | 17 | 12 | 1.1e-003 | CENTRIMO | http://the_brain.bwh.harvard.edu/uniprobe/details2.php?id=00059 |
| 323 | db/MOUSE/ur | UP00065_2 | Zfp161_secon | GYCGCGCAR | 14 | 12 | 1.1e-003 | CENTRIMO | http://the_brain.bwh.harvard.edu/uniprobe/details2.php?id=00065 |
| 324 | db/EUKARYO | CEBPB_DBD | | RTTRCGCAA | 10 | 12 | 1.3e-003 | CENTRIMO | http://floresta.eead.csic.es/footprintdb/index.php?db=HumanTF:1.0&motif=CEBPB_DBD |
| 325 | db/EUKARYO | CEBPB_full | | ATTRCGCAA | 10 | 12 | 1.3e-003 | CENTRIMO | http://floresta.eead.csic.es/footprintdb/index.php?db=HumanTF:1.0&motif=CEBPB_full |
| 326 | db/EUKARYO | CEBPD_DBD | | RTTRCGCAA | 10 | 12 | 1.3e-003 | CENTRIMO | http://floresta.eead.csic.es/footprintdb/index.php?db=HumanTF:1.0&motif=CEBPD_DBD |
| 327 | db/EUKARYO | CEBPE_DBD | | VTTRCGCAA | 10 | 12 | 1.3e-003 | CENTRIMO | http://floresta.eead.csic.es/footprintdb/index.php?db=HumanTF:1.0&motif=CEBPE_DBD |
| 328 | db/EUKARYO | CEBPG_DBD | | ATTRCGCAA | 10 | 12 | 1.3e-003 | CENTRIMO | http://floresta.eead.csic.es/footprintdb/index.php?db=HumanTF:1.0&motif=CEBPG_DBD |
| 329 | db/EUKARYO | CEBPG_full | | ATTRCGCAA | 10 | 12 | 1.3e-003 | CENTRIMO | http://floresta.eead.csic.es/footprintdb/index.php?db=HumanTF:1.0&motif=CEBPG_full |
| 330 | db/EUKARYO | Cebpb_DBD | | RTTRCGCAA | 10 | 12 | 1.3e-003 | CENTRIMO | http://floresta.eead.csic.es/footprintdb/index.php?db=HumanTF:1.0&motif=Cebpb_DBD |
| 331 | db/JASPAR/J | MA0466.2 | CEBPB | RTTRCGCAA | 10 | 12 | 1.3e-003 | CENTRIMO | http://jaspar2018.genereg.net/matrix/MA0466.2 |
| 332 | db/JASPAR/J | MA0836.1 | CEBPD | RTTRCGCAA | 10 | 12 | 1.3e-003 | CENTRIMO | http://jaspar2018.genereg.net/matrix/MA0836.1 |
| 333 | db/JASPAR/J | MA0837.1 | CEBPE | VTTRCGCAA | 10 | 12 | 1.3e-003 | CENTRIMO | http://jaspar2018.genereg.net/matrix/MA0837.1 |
| 334 | db/JASPAR/J | MA0838.1 | CEBPG | ATTRCGCAA | 10 | 12 | 1.3e-003 | CENTRIMO | http://jaspar2018.genereg.net/matrix/MA0838.1 |
| 335 | db/EUKARYO | DBP_full | | NRTTACGTA | 12 | 12 | 1.3e-003 | CENTRIMO | http://floresta.eead.csic.es/footprintdb/index.php?db=HumanTF:1.0&motif=DBP_full |
| 336 | db/EUKARYO | HLF_full | | NVTTACRTA | 12 | 12 | 1.3e-003 | CENTRIMO | http://floresta.eead.csic.es/footprintdb/index.php?db=HumanTF:1.0&motif=HLF_full |
| 337 | db/EUKARYO | NFIL3_DBD | | NRTTACRTA | 12 | 12 | 1.3e-003 | CENTRIMO | http://floresta.eead.csic.es/footprintdb/index.php?db=HumanTF:1.0&motif=NFIL3_DBD |
| 338 | db/JASPAR/J | MA0639.1 | DBP | NRTTACGTA | 12 | 12 | 1.3e-003 | CENTRIMO | http://jaspar2018.genereg.net/matrix/MA0639.1 |
| 339 | db/JASPAR/J | MA0043.2 | HLF | NVTTACRTA | 12 | 12 | 1.3e-003 | CENTRIMO | http://jaspar2018.genereg.net/matrix/MA0043.2 |
| 340 | db/JASPAR/J | MA0036.3 | GATA2 | NBCTTATCTN | 11 | 12 | 1.3e-003 | CENTRIMO | http://jaspar2018.genereg.net/matrix/MA0036.3 |
| 341 | db/EUKARYO | ATF4_DBD | | GKATGAYGC | 13 | 12 | 1.4e-003 | CENTRIMO | http://floresta.eead.csic.es/footprintdb/index.php?db=HumanTF:1.0&motif=ATF4_DBD |
| 342 | db/JASPAR/J | MA0833.1 | ATF4 | GKATGAYGC | 13 | 12 | 1.4e-003 | CENTRIMO | http://jaspar2018.genereg.net/matrix/MA0833.1 |
| 343 | db/JASPAR/J | MA1104.1 | GATA6 | NDNAGATAA | 13 | 12 | 1.4e-003 | CENTRIMO | http://jaspar2018.genereg.net/matrix/MA1104.1 |
| 344 | db/JASPAR/J | MA0604.1 | Atf1 | RTGACGTA | 8 | 12 | 1.4e-003 | CENTRIMO | http://jaspar2018.genereg.net/matrix/MA0604.1 |
| 345 | db/JASPAR/J | MA1132.1 | JUN::JUNB | KATGACKCA | 10 | 12 | 1.4e-003 | CENTRIMO | http://jaspar2018.genereg.net/matrix/MA1132.1 |
| 346 | db/MOUSE/ur | UP00080_1 | Gata5_primar | HWWRCTGA | 17 | 12 | 1.4e-003 | CENTRIMO | http://the_brain.bwh.harvard.edu/uniprobe/details2.php?id=00080 |
| 347 | db/EUKARYO | Atf4_DBD | | AGGATGATG | 14 | 12 | 1.4e-003 | CENTRIMO | http://floresta.eead.csic.es/footprintdb/index.php?db=HumanTF:1.0&motif=Atf4_DBD |
| 348 | db/MOUSE/ur | UP00032_1 | Gata3_primar | BDWDDAKA | 22 | 12 | 1.4e-003 | CENTRIMO | http://the_brain.bwh.harvard.edu/uniprobe/details2.php?id=00032 |
| 349 | db/MOUSE/ur | UP00073_2 | Foxa2_secon | VVWAATAAC | 15 | 12 | 1.5e-003 | CENTRIMO | http://the_brain.bwh.harvard.edu/uniprobe/details2.php?id=00073 |
| 350 | db/JASPAR/J | MA0831.2 | TFE3 | CACGTGAY | 8 | 12 | 1.5e-003 | CENTRIMO | http://jaspar2018.genereg.net/matrix/MA0831.2 |
| 351 | db/EUKARYO | LBX2_DBD_1 | | CTBRANSTR | 13 | 12 | 1.6e-003 | CENTRIMO | http://floresta.eead.csic.es/footprintdb/index.php?db=HumanTF:1.0&motif=LBX2_DBD_1 |
| 352 | db/EUKARYO | IRF3_full | | NSRRAAMGC | 21 | 12 | 2.8e-003 | CENTRIMO | http://floresta.eead.csic.es/footprintdb/index.php?db=HumanTF:1.0&motif=IRF3_full |
| 353 | DREME | AAYCCCAG | DREME-7 | AAYCCCAG | 8 | 43 | 2.9e-003 | DREME | |
| 354 | db/JASPAR/J | MA0111.1 | Spz1 | AGGGTWWC | 11 | 12 | 3.3e-003 | CENTRIMO | http://jaspar2018.genereg.net/matrix/MA0111.1 |
| 355 | db/JASPAR/J | MA0470.1 | E2F4 | GGGCGGGA | 11 | 12 | 3.7e-003 | CENTRIMO | http://jaspar2018.genereg.net/matrix/MA0470.1 |
| 356 | db/JASPAR/J | MA1122.1 | TFDP1 | VSGCGGGA | 11 | 12 | 3.7e-003 | CENTRIMO | http://jaspar2018.genereg.net/matrix/MA1122.1 |
| 357 | db/JASPAR/J | MA0089.1 | MAFG::NFE2I | NATGAC | 6 | 12 | 3.8e-003 | CENTRIMO | http://jaspar2018.genereg.net/matrix/MA0089.1 |

| | | | | | | | | | | | |
|---|---|---|---|---|---|---|---|---|---|---|---|
| 358 | db/EUKARYO | SP1_DBD | | RCCMCRCC( | 11 | 12 | 3.9e-003 | CENTRIMO | | | http://floresta.eead.csic.es/footprintdb/index.php?db=HumanTF:1.0&motif=SP1_DBD |
| 359 | db/EUKARYO | SP4_full | | BWRGCCAC( | 17 | 12 | 4.0e-003 | CENTRIMO | | | http://floresta.eead.csic.es/footprintdb/index.php?db=HumanTF:1.0&motif=SP4_full |
| 360 | db/JASPAR/J/ | MA0685.1 | SP4 | BWRGCCAC( | 17 | 12 | 4.0e-003 | CENTRIMO | | | http://jaspar2018.genereg.net/matrix/MA0685.1 |
| 361 | db/EUKARYO | SP8_DBD | | RCCACGCC( | 12 | 12 | 4.0e-003 | CENTRIMO | | | http://floresta.eead.csic.es/footprintdb/index.php?db=HumanTF:1.0&motif=SP8_DBD |
| 362 | db/JASPAR/J/ | MA0747.1 | SP8 | RCCACGCC( | 12 | 12 | 4.0e-003 | CENTRIMO | | | http://jaspar2018.genereg.net/matrix/MA0747.1 |
| 363 | db/EUKARYO | KLF14_DBD | | KRCCACGC( | 14 | 12 | 4.0e-003 | CENTRIMO | | | http://floresta.eead.csic.es/footprintdb/index.php?db=HumanTF:1.0&motif=KLF14_DBD |
| 364 | db/JASPAR/J/ | MA0740.1 | KLF14 | KRCCACGC( | 14 | 12 | 4.0e-003 | CENTRIMO | | | http://jaspar2018.genereg.net/matrix/MA0740.1 |
| 365 | db/JASPAR/J/ | MA0516.1 | SP2 | GYCCCGCCY | 15 | 12 | 4.2e-003 | CENTRIMO | | | http://jaspar2018.genereg.net/matrix/MA0516.1 |
| 366 | db/MOUSE/ur | UP00002_1 | Sp4_primary | BDHMMCGC( | 17 | 12 | 4.2e-003 | CENTRIMO | | | http://the_brain.bwh.harvard.edu/uniprobe/details2.php?id=00002 |
| 367 | db/EUKARYO | EGR2_DBD | | MCGCCCAC( | 11 | 12 | 4.3e-003 | CENTRIMO | | | http://floresta.eead.csic.es/footprintdb/index.php?db=HumanTF:1.0&motif=EGR2_DBD |
| 368 | db/JASPAR/J/ | MA0472.2 | EGR2 | MCGCCCAC( | 11 | 12 | 4.3e-003 | CENTRIMO | | | http://jaspar2018.genereg.net/matrix/MA0472.2 |
| 369 | db/EUKARYO | EGR2_full | | NHMCGCCC/ | 15 | 12 | 4.3e-003 | CENTRIMO | | | http://floresta.eead.csic.es/footprintdb/index.php?db=HumanTF:1.0&motif=EGR2_full |
| 370 | db/EUKARYO | GLI2_DBD_2 | | GCGACCACN | 12 | 12 | 4.4e-003 | CENTRIMO | | | http://floresta.eead.csic.es/footprintdb/index.php?db=HumanTF:1.0&motif=GLI2_DBD_2 |
| 371 | db/JASPAR/J/ | MA0734.1 | GLI2 | GCGACCACN | 12 | 12 | 4.4e-003 | CENTRIMO | | | http://jaspar2018.genereg.net/matrix/MA0734.1 |
| 372 | db/EUKARYO | EGR1_DBD | | HMCGCCCM( | 14 | 12 | 4.4e-003 | CENTRIMO | | | http://floresta.eead.csic.es/footprintdb/index.php?db=HumanTF:1.0&motif=EGR1_DBD |
| 373 | db/EUKARYO | Egr1_mouse_DBD_mutant_I | | HHMCGCCCN | 16 | 12 | 4.4e-003 | CENTRIMO | | | http://floresta.eead.csic.es/footprintdb/index.php?db=HumanTF:1.0&motif=Egr1_mouse_DBD_mutant_DBD |
| 374 | db/EUKARYO | ZBTB7B_full | | RCGACCACC | 12 | 12 | 4.6e-003 | CENTRIMO | | | http://floresta.eead.csic.es/footprintdb/index.php?db=HumanTF:1.0&motif=ZBTB7B_full |
| 375 | db/EUKARYO | ZBTB7C_full | | RCGACCACC | 12 | 12 | 4.6e-003 | CENTRIMO | | | http://floresta.eead.csic.es/footprintdb/index.php?db=HumanTF:1.0&motif=ZBTB7C_full |
| 376 | db/JASPAR/J/ | MA0694.1 | ZBTB7B | RCGACCACC | 12 | 12 | 4.6e-003 | CENTRIMO | | | http://jaspar2018.genereg.net/matrix/MA0694.1 |
| 377 | db/JASPAR/J/ | MA0695.1 | ZBTB7C | RCGACCACC | 12 | 12 | 4.6e-003 | CENTRIMO | | | http://jaspar2018.genereg.net/matrix/MA0695.1 |
| 378 | db/EUKARYO | TCF4_full | | HRCACCTGB | 10 | 12 | 5.9e-003 | CENTRIMO | | | http://floresta.eead.csic.es/footprintdb/index.php?db=HumanTF:1.0&motif=TCF4_full |
| 379 | db/EUKARYO | TBX21_full_2 | | AAGGTGTGA | 10 | 12 | 6.2e-003 | CENTRIMO | | | http://floresta.eead.csic.es/footprintdb/index.php?db=HumanTF:1.0&motif=TBX21_full_2 |
| 380 | db/EUKARYO | TBR1_full | | AAGGTGTGA | 11 | 12 | 6.3e-003 | CENTRIMO | | | http://floresta.eead.csic.es/footprintdb/index.php?db=HumanTF:1.0&motif=TBR1_full |
| 381 | db/EUKARYO | TBX20_full_1 | | WAGGTGTG/ | 11 | 12 | 6.3e-003 | CENTRIMO | | | http://floresta.eead.csic.es/footprintdb/index.php?db=HumanTF:1.0&motif=TBX20_full_1 |
| 382 | db/JASPAR/J/ | MA0689.1 | TBX20 | WAGGTGTG/ | 11 | 12 | 6.3e-003 | CENTRIMO | | | http://jaspar2018.genereg.net/matrix/MA0689.1 |
| 383 | db/EUKARYO | SPDEF_DBD_3 | | RCAGDAAGA | 16 | 12 | 7.5e-003 | CENTRIMO | | | http://floresta.eead.csic.es/footprintdb/index.php?db=HumanTF:1.0&motif=SPDEF_DBD_3 |
| 384 | db/EUKARYO | SPDEF_full_3 | | GCAGDAAGA | 16 | 12 | 7.5e-003 | CENTRIMO | | | http://floresta.eead.csic.es/footprintdb/index.php?db=HumanTF:1.0&motif=SPDEF_full_3 |
| 385 | db/EUKARYO | TFAP2A_DBD_2 | | HGCCYSAGC | 11 | 12 | 8.4e-003 | CENTRIMO | | | http://floresta.eead.csic.es/footprintdb/index.php?db=HumanTF:1.0&motif=TFAP2A_DBD_2 |
| 386 | db/JASPAR/J/ | MA0003.3 | TFAP2A | HGCCYSAGC | 11 | 12 | 8.4e-003 | CENTRIMO | | | http://jaspar2018.genereg.net/matrix/MA0003.3 |
| 387 | DREME | HTTTAAAA | DREME-9 | HTTTAAAA | 8 | 66 | 8.9e-003 | DREME | | | |
| 388 | db/EUKARYO | E2F1_DBD_1 | | WWTGGCGC | 12 | 12 | 1.6e-002 | CENTRIMO | | | http://floresta.eead.csic.es/footprintdb/index.php?db=HumanTF:1.0&motif=E2F1_DBD_1 |
| 389 | db/EUKARYO | E2F1_DBD_2 | | TTTGGCGCC | 12 | 12 | 1.6e-002 | CENTRIMO | | | http://floresta.eead.csic.es/footprintdb/index.php?db=HumanTF:1.0&motif=E2F1_DBD_2 |
| 390 | db/JASPAR/J/ | MA0024.3 | E2F1 | TTTGGCGCC | 12 | 12 | 1.6e-002 | CENTRIMO | | | http://jaspar2018.genereg.net/matrix/MA0024.3 |
| 391 | db/EUKARYO | E2F1_DBD_3 | | WWTTGGCG | 14 | 12 | 1.6e-002 | CENTRIMO | | | http://floresta.eead.csic.es/footprintdb/index.php?db=HumanTF:1.0&motif=E2F1_DBD_3 |
| 392 | db/MOUSE/ur | UP00001_1 | E2F2_primary | NHWARGGC( | 15 | 12 | 1.6e-002 | CENTRIMO | | | http://the_brain.bwh.harvard.edu/uniprobe/details2.php?id=00001 |
| 393 | db/MOUSE/ur | UP00003_1 | E2F3_primary | VHDADGGC( | 15 | 12 | 1.6e-002 | CENTRIMO | | | http://the_brain.bwh.harvard.edu/uniprobe/details2.php?id=00003 |
| 394 | db/EUKARYO | YY1_full | | NCCGCCATT | 11 | 12 | 1.7e-002 | CENTRIMO | | | http://floresta.eead.csic.es/footprintdb/index.php?db=HumanTF:1.0&motif=YY1_full |
| 395 | db/JASPAR/J/ | MA0107.1 | RELA | GGGRATTTC | 10 | 12 | 1.8e-002 | CENTRIMO | | | http://jaspar2018.genereg.net/matrix/MA0107.1 |
| 396 | DREME | GGCTGCTM | DREME-10 | GGCTGCTM | 8 | 30 | 2.1e-002 | DREME | | | |
| 397 | db/MOUSE/ur | UP00101_2 | Sox12_secon( | VVNBASACA/ | 16 | 12 | 3.1e-002 | CENTRIMO | | | http://the_brain.bwh.harvard.edu/uniprobe/details2.php?id=00101 |
| 398 | DREME | ACRTGG | DREME-11 | ACRTGG | 6 | 142 | 3.6e-002 | DREME | | | |
| 399 | db/MOUSE/ur | UP00074_1 | Isgf3g_primar | NDRAAWCG/ | 15 | 12 | 3.9e-002 | CENTRIMO | | | http://the_brain.bwh.harvard.edu/uniprobe/details2.php?id=00074 |
| 400 | db/EUKARYO | ETV5_DBD | | ACCGGAWG' | 10 | 12 | 4.8e-002 | CENTRIMO | | | http://floresta.eead.csic.es/footprintdb/index.php?db=HumanTF:1.0&motif=ETV5_DBD |
| 401 | db/JASPAR/J/ | MA0765.1 | ETV5 | ACCGGAWG' | 10 | 12 | 4.8e-002 | CENTRIMO | | | http://jaspar2018.genereg.net/matrix/MA0765.1 |
| 402 | db/MOUSE/ur | UP00071_2 | Sox21_secon( | NVYSWATTG | 17 | 12 | 4.9e-002 | CENTRIMO | | | http://the_brain.bwh.harvard.edu/uniprobe/details2.php?id=00071 |

### MEME-ChIP (Motif Analysis of Large Nucleotide Datasets): Version 5.0.3 released on Sun Dec 02 18:41:45 2018 -0800
### The format of this file is described at http://meme-suite.org//doc/meme-chip-output-format.html
### meme-chip -oc . -time 300 -ccut 100 -order 1 -db db/EUKARYOTE/jolma2013.meme -db db/JASPAR/JASPAR2018_CORE_vertebrates_non-redundant.meme -db db/MOUSE/uniprobe_mouse.meme -meme-mod zoops -meme-minw 6 -meme-maxw 30 -meme-nmotifs 3 -meme-searchsize 1000(

**ZT12 dynamic**

| MOTIF_INDEX | MOTIF_SOUF | MOTIF_ID | ALT_ID | CONSENSUS | WIDTH | SITES | E-VALUE | E-VALUE_SO | MOST_SIMILAR_MC | **MOST_SIMILAR_MO'** | URL |
|---|---|---|---|---|---|---|---|---|---|---|---|
| 1 | MEME | CWCTDTGTA | MEME-1 | CWCTDTGTA | 28 | 87 | 8.0e-223 | MEME | db/JASPAR/JASPAR | **MA0505.1 (Nr5a2)** | http://jaspar2018.genereg.net/matrix/MA0505.1 |
| 2 | MEME | TKTDTKTDTK | MEME-2 | TKTDTKTDTK | 21 | 306 | 1.0e-144 | MEME | db/JASPAR/JASPAR | **MA1125.1 (ZNF384)** | http://jaspar2018.genereg.net/matrix/MA1125.1 |
| 3 | MEME | GTWGCTGT( | MEME-3 | GTWGCTGT( | 30 | 13 | 5.2e-070 | MEME | | | |
| 4 | DREME | AGRMAG | DREME-1 | AGRMAG | 6 | 1358 | 8.8e-013 | DREME | | | |
| 5 | DREME | AGGCCAGM | DREME-2 | AGGCCAGM | 8 | 105 | 1.0e-011 | DREME | db/JASPAR/JASPAR | **MA0505.1 (Nr5a2)** | http://jaspar2018.genereg.net/matrix/MA0505.1 |
| 6 | DREME | GGKCTACA | DREME-3 | GGKCTACA | 8 | 65 | 1.7e-010 | DREME | | | |
| 7 | DREME | AAAANAAA | DREME-4 | AAAANAAA | 8 | 319 | 4.0e-008 | DREME | db/JASPAR/JASPAR | **MA1125.1 (ZNF384)** | http://jaspar2018.genereg.net/matrix/MA1125.1 |
| 8 | DREME | AWTCCCAG | DREME-5 | AWTCCCAG | 8 | 84 | 2.8e-007 | DREME | | | |

| | | | | | | | | | | | |
|---|---|---|---|---|---|---|---|---|---|---|---|
| 9 | DREME | AGATGGYT | DREME-6 | AGATGGYT | 8 | 61 | 1.9e-006 | DREME | | | |
| 10 | DREME | AHAYA | DREME-7 | AHAYA | 5 | 3570 | 3.5e-005 | DREME | | | |
| 11 | DREME | GAACTCA | DREME-8 | GAACTCA | 7 | 122 | 1.1e-004 | DREME | db/JASPAR/JASPAR | **MA0693.2 (VDR)** | http://jaspar2018.genereg.net/matrix/MA0693.2 |
| 12 | DREME | CACCATGY | DREME-9 | CACCATGY | 8 | 56 | 4.2e-003 | DREME | db/JASPAR/JASPAR | **MA0119.1 (NFIC::TLX** | http://jaspar2018.genereg.net/matrix/MA0119.1 |
| 13 | DREME | CAGKACAG | DREME-10 | CAGKACAG | 8 | 65 | 3.2e-002 | DREME | | | |

### MEME-ChIP (Motif Analysis of Large Nucleotide Datasets): Version 5.0.3 released on Sun Dec 02 18:41:45 2018 -0800
### The format of this file is described at http://meme-suite.org//doc/meme-chip-output-format.html
### meme-chip -oc . -time 300 -ccut 100 -order 1 -db db/EUKARYOTE/jolma2013.meme -db db/JASPAR/JASPAR2018_CORE_vertebrates_non-redundant.meme -db db/MOUSE/uniprobe_mouse.meme -meme-mod zoops -meme-minw 6 -meme-maxw 30 -meme-nmotifs 3 -meme-searchsize 1000(

**ZT12 static**

| MOTIF_INDEX | MOTIF_SOUF | MOTIF_ID | ALT_ID | CONSENSUS | WIDTH | SITES | E-VALUE | E-VALUE_SO | MOST_SIMILAR_MC | **MOST_SIMILAR_MO** | URL |
|---|---|---|---|---|---|---|---|---|---|---|---|
| 1 | MEME | TTTTWTTTTT | MEME-1 | TTTTWTTTTT | 21 | 415 | 3.5e-097 | MEME | db/JASPAR/JASPAR | **MA1125.1 (ZNF384)** | http://jaspar2018.genereg.net/matrix/MA1125.1 |
| 2 | MEME | CYWCCYGM | MEME-2 | CYWCCYGM | 29 | 24 | 9.9e-051 | MEME | db/MOUSE/uniprobe_ | **UP00043_2 (Bcl6b_s** | http://the_brain.bwh.harvard.edu/uniprobe/details2.php?id=00043 |
| 3 | MEME | SMCHVKGYA | MEME-3 | SMCHVKGYA | 27 | 93 | 4.3e-048 | MEME | | | |
| 4 | db/MOUSE/ur | UP00034_2 | Sox7_second | GTVSWAATT | 22 | 10 | 4.3e-027 | CENTRIMO | | | http://the_brain.bwh.harvard.edu/uniprobe/details2.php?id=00034 |
| 5 | db/MOUSE/ur | UP00138_1 | Bsx_3483.2 | VVRGYAATTI | 16 | 10 | 2.5e-022 | CENTRIMO | | | http://the_brain.bwh.harvard.edu/uniprobe/details2.php?id=00138 |
| 6 | db/MOUSE/ur | UP00230_1 | Dlx5_3419.2 | BSRGYAATTI | 16 | 10 | 2.5e-022 | CENTRIMO | | | http://the_brain.bwh.harvard.edu/uniprobe/details2.php?id=00230 |
| 7 | db/MOUSE/ur | UP00086_2 | Irf3_secondar | RKAGAAWG | 14 | 10 | 4.1e-020 | CENTRIMO | | | http://the_brain.bwh.harvard.edu/uniprobe/details2.php?id=00086 |
| 8 | db/EUKARYO | LBX2_DBD_1 | | CTBRANSTR | 13 | 10 | 2.5e-019 | CENTRIMO | | | http://floresta.eead.csic.es/footprintdb/index.php?db=HumanTF:1.0&motif=LBX2_DBD_1 |
| 9 | db/MOUSE/ur | UP00027_2 | Osr1_second | RHHHGCTAC | 16 | 10 | 3.0e-018 | CENTRIMO | | | http://the_brain.bwh.harvard.edu/uniprobe/details2.php?id=00027 |
| 10 | db/MOUSE/ur | UP00052_2 | Osr2_second | ANHTGCTAC | 16 | 10 | 3.0e-018 | CENTRIMO | | | http://the_brain.bwh.harvard.edu/uniprobe/details2.php?id=00052 |
| 11 | db/MOUSE/ur | UP00036_2 | Myf6_second | VNNRACAGV | 15 | 10 | 2.6e-016 | CENTRIMO | | | http://the_brain.bwh.harvard.edu/uniprobe/details2.php?id=00036 |
| 12 | db/MOUSE/ur | UP00001_1 | E2F2_primary | NHWARGGC | 15 | 10 | 3.0e-015 | CENTRIMO | | | http://the_brain.bwh.harvard.edu/uniprobe/details2.php?id=00001 |
| 13 | db/MOUSE/ur | UP00003_1 | E2F3_primary | VHDADGGC | 15 | 10 | 3.0e-015 | CENTRIMO | | | http://the_brain.bwh.harvard.edu/uniprobe/details2.php?id=00003 |
| 14 | db/MOUSE/ur | UP00043_2 | Bcl6b_second | HNNBCCGC | 16 | 10 | 3.8e-015 | CENTRIMO | | | http://the_brain.bwh.harvard.edu/uniprobe/details2.php?id=00043 |
| 15 | DREME | AACRCCG | DREME-15 | AACRCCG | 7 | 40 | 1.1e-013 | DREME | | | |
| 16 | db/EUKARYO | MAFK_DBD_1 | | DWWWDTGC | 12 | 10 | 8.7e-013 | CENTRIMO | | | http://floresta.eead.csic.es/footprintdb/index.php?db=HumanTF:1.0&motif=MAFK_DBD_1 |
| 17 | DREME | RCCTTTAA | DREME-19 | RCCTTTAA | 8 | 42 | 2.2e-010 | DREME | db/EUKARYOTE/joln | **NR4A2_full_3** | http://floresta.eead.csic.es/footprintdb/index.php?db=HumanTF:1.0&motif=NR4A2_full_3 |
| 18 | db/EUKARYO | ZIC1_full | | GACCCCCYG | 14 | 10 | 2.2e-010 | CENTRIMO | | | http://floresta.eead.csic.es/footprintdb/index.php?db=HumanTF:1.0&motif=ZIC1_full |
| 19 | db/JASPAR/J/ | MA0696.1 | ZIC1 | GACCCCCYG | 14 | 10 | 2.2e-010 | CENTRIMO | | | http://jaspar2018.genereg.net/matrix/MA0696.1 |
| 20 | db/EUKARYO | ZIC3_full | | GACCCCCCC | 15 | 10 | 2.4e-010 | CENTRIMO | | | http://floresta.eead.csic.es/footprintdb/index.php?db=HumanTF:1.0&motif=ZIC3_full |
| 21 | db/JASPAR/J/ | MA0697.1 | ZIC3 | GACCCCCCC | 15 | 10 | 2.4e-010 | CENTRIMO | | | http://jaspar2018.genereg.net/matrix/MA0697.1 |
| 22 | DREME | AWAYAAA | DREME-1 | AWAYAAA | 7 | 456 | 1.7e-009 | DREME | db/EUKARYOTE/joln | **Foxc1_DBD_1** | http://floresta.eead.csic.es/footprintdb/index.php?db=HumanTF:1.0&motif=Foxc1_DBD_1 |
| 23 | db/EUKARYO | FOXB1_DBD_2 | | WATGTAAAT/ | 18 | 10 | 2.7e-009 | CENTRIMO | | | http://floresta.eead.csic.es/footprintdb/index.php?db=HumanTF:1.0&motif=FOXB1_DBD_2 |
| 24 | DREME | GAGGCAGR | DREME-2 | GAGGCAGR | 8 | 110 | 2.8e-009 | DREME | | | |
| 25 | db/EUKARYO | OLIG2_DBD | | AMCATATGK | 10 | 10 | 2.4e-008 | CENTRIMO | | | http://floresta.eead.csic.es/footprintdb/index.php?db=HumanTF:1.0&motif=OLIG2_DBD |
| 26 | db/MOUSE/ur | UP00180_1 | Hoxd13_2356 | NNNYYAATA/ | 16 | 10 | 3.7e-008 | CENTRIMO | | | http://the_brain.bwh.harvard.edu/uniprobe/details2.php?id=00180 |
| 27 | db/MOUSE/ur | UP00035_1 | Hic1_primary | NBBATGCCA | 16 | 10 | 4.0e-008 | CENTRIMO | | | http://the_brain.bwh.harvard.edu/uniprobe/details2.php?id=00035 |
| 28 | DREME | ACAGWRA | DREME-3 | ACAGWRA | 7 | 331 | 4.2e-008 | DREME | | | |
| 29 | DREME | AGCCSTGG | DREME-4 | AGCCSTGG | 8 | 73 | 1.1e-007 | DREME | | | |
| 30 | db/EUKARYO | YY2_full_2 | | CCATGCCGC | 12 | 10 | 1.2e-007 | CENTRIMO | | | http://floresta.eead.csic.es/footprintdb/index.php?db=HumanTF:1.0&motif=YY2_full_2 |
| 31 | db/MOUSE/ur | UP00001_2 | E2F2_second | NNNWYGGC | 17 | 10 | 1.3e-007 | CENTRIMO | | | http://the_brain.bwh.harvard.edu/uniprobe/details2.php?id=00001 |
| 32 | db/MOUSE/ur | UP00003_2 | E2F3_second | NNYWYGGC | 17 | 10 | 1.3e-007 | CENTRIMO | | | http://the_brain.bwh.harvard.edu/uniprobe/details2.php?id=00003 |
| 33 | db/EUKARYO | E2F1_DBD_1 | | WWTGGCGC | 12 | 10 | 1.3e-007 | CENTRIMO | | | http://floresta.eead.csic.es/footprintdb/index.php?db=HumanTF:1.0&motif=E2F1_DBD_1 |
| 34 | db/EUKARYO | E2F1_DBD_2 | | TTTGGCGCC | 12 | 10 | 1.3e-007 | CENTRIMO | | | http://floresta.eead.csic.es/footprintdb/index.php?db=HumanTF:1.0&motif=E2F1_DBD_2 |
| 35 | db/JASPAR/J/ | MA0024.3 | E2F1 | TTTGGCGCC | 12 | 10 | 1.3e-007 | CENTRIMO | | | http://jaspar2018.genereg.net/matrix/MA0024.3 |
| 36 | db/MOUSE/ur | UP00055_1 | Hbp1_primary | HNNWTSAAT | 16 | 10 | 1.3e-007 | CENTRIMO | | | http://the_brain.bwh.harvard.edu/uniprobe/details2.php?id=00055 |
| 37 | db/EUKARYO | YY1_full | | NCCGCCATT | 11 | 10 | 1.6e-007 | CENTRIMO | | | http://floresta.eead.csic.es/footprintdb/index.php?db=HumanTF:1.0&motif=YY1_full |
| 38 | DREME | AGTSCTGG | DREME-5 | AGTSCTGG | 8 | 78 | 6.3e-007 | DREME | | | |
| 39 | db/MOUSE/ur | UP00084_2 | Gmeb1_secor | KGRBCRACG | 16 | 10 | 6.5e-007 | CENTRIMO | | | http://the_brain.bwh.harvard.edu/uniprobe/details2.php?id=00084 |
| 40 | db/MOUSE/ur | UP00090_1 | Elf3_primary | DAHVMGGA/ | 13 | 10 | 9.7e-007 | CENTRIMO | | | http://the_brain.bwh.harvard.edu/uniprobe/details2.php?id=00090 |
| 41 | db/MOUSE/ur | UP00059_2 | Arid5a_secon | NNYNMAATA | 17 | 10 | 1.2e-006 | CENTRIMO | | | http://the_brain.bwh.harvard.edu/uniprobe/details2.php?id=00059 |
| 42 | db/MOUSE/ur | UP00009_1 | Nr2f2_primary | NHBNAAAGG | 16 | 10 | 1.4e-006 | CENTRIMO | | | http://the_brain.bwh.harvard.edu/uniprobe/details2.php?id=00009 |
| 43 | db/EUKARYO | NR2F1_full | | GRGGTCAA | 8 | 10 | 1.4e-006 | CENTRIMO | | | http://floresta.eead.csic.es/footprintdb/index.php?db=HumanTF:1.0&motif=NR2F1_full |
| 44 | db/EUKARYO | NR2F1_DBD_3 | | CARAGGTCA | 13 | 10 | 1.5e-006 | CENTRIMO | | | http://floresta.eead.csic.es/footprintdb/index.php?db=HumanTF:1.0&motif=NR2F1_DBD_3 |
| 45 | db/JASPAR/J/ | MA0017.2 | NR2F1 | CARAGGTCA | 13 | 10 | 1.5e-006 | CENTRIMO | | | http://jaspar2018.genereg.net/matrix/MA0017.2 |

| | | | | | | | | | | |
|---|---|---|---|---|---|---|---|---|---|---|
| 46 | db/MOUSE/ur | UP00069_1 | Sox1_primary | HWDYAATTC | 16 | 10 | 1.5e-006 | CENTRIMO | | http://the_brain.bwh.harvard.edu/uniprobe/details2.php?id=00069 |
| 47 | db/EUKARYO | OTX2_DBD_2 | | YTAATCCB | 8 | 10 | 3.0e-006 | CENTRIMO | | http://floresta.eead.csic.es/footprintdb/index.php?db=HumanTF:1.0&motif=OTX2_DBD_2 |
| 48 | db/EUKARYO | Otx1_DBD_2 | | HTAATCCB | 8 | 10 | 3.0e-006 | CENTRIMO | | http://floresta.eead.csic.es/footprintdb/index.php?db=HumanTF:1.0&motif=Otx1_DBD_2 |
| 49 | db/JASPAR/J/ | MA0712.1 | OTX2 | YTAATCCB | 8 | 10 | 3.0e-006 | CENTRIMO | | http://jaspar2018.genereg.net/matrix/MA0712.1 |
| 50 | DREME | CCTGGAAC | DREME-6 | CCTGGAAC | 8 | 41 | 3.6e-006 | DREME | | |
| 51 | db/MOUSE/ur | UP00099_2 | Ascl2_second | BYVWCCCCD | 16 | 10 | 4.6e-006 | CENTRIMO | | http://the_brain.bwh.harvard.edu/uniprobe/details2.php?id=00099 |
| 52 | db/EUKARYO | KLF16_DBD | | GMCACGCC( | 11 | 10 | 4.7e-006 | CENTRIMO | | http://floresta.eead.csic.es/footprintdb/index.php?db=HumanTF:1.0&motif=KLF16_DBD |
| 53 | db/EUKARYO | SP3_DBD | | VCCACGCCC | 11 | 10 | 4.7e-006 | CENTRIMO | | http://floresta.eead.csic.es/footprintdb/index.php?db=HumanTF:1.0&motif=SP3_DBD |
| 54 | db/JASPAR/J/ | MA0079.3 | SP1 | GCCCCKCCC | 11 | 10 | 4.7e-006 | CENTRIMO | | http://jaspar2018.genereg.net/matrix/MA0079.3 |
| 55 | db/JASPAR/J/ | MA0741.1 | KLF16 | GMCACGCC( | 11 | 10 | 4.7e-006 | CENTRIMO | | http://jaspar2018.genereg.net/matrix/MA0741.1 |
| 56 | db/JASPAR/J/ | MA0746.1 | SP3 | VCCACGCCC | 11 | 10 | 4.7e-006 | CENTRIMO | | http://jaspar2018.genereg.net/matrix/MA0746.1 |
| 57 | db/MOUSE/ur | UP00000_2 | Smad3_secon | KHHNCCCCG | 17 | 10 | 4.8e-006 | CENTRIMO | | http://the_brain.bwh.harvard.edu/uniprobe/details2.php?id=00000 |
| 58 | db/EUKARYO | EGR2_full | | NHMCGCCC/ | 15 | 10 | 5.5e-006 | CENTRIMO | | http://floresta.eead.csic.es/footprintdb/index.php?db=HumanTF:1.0&motif=EGR2_full |
| 59 | db/EUKARYO | EGR1_DBD | | HMCGCCCM( | 14 | 10 | 5.7e-006 | CENTRIMO | | http://floresta.eead.csic.es/footprintdb/index.php?db=HumanTF:1.0&motif=EGR1_DBD |
| 60 | db/MOUSE/ur | UP00054_2 | Tcf7_seconda | MBKTATTAN/ | 15 | 10 | 5.8e-006 | CENTRIMO | | http://the_brain.bwh.harvard.edu/uniprobe/details2.php?id=00054 |
| 61 | db/JASPAR/J/ | MA0163.1 | PLAG1 | GGGGCCCW | 14 | 10 | 7.0e-006 | CENTRIMO | | http://jaspar2018.genereg.net/matrix/MA0163.1 |
| 62 | db/EUKARYO | MLX_full | | RTCACGTGA | 10 | 10 | 8.9e-006 | CENTRIMO | | http://floresta.eead.csic.es/footprintdb/index.php?db=HumanTF:1.0&motif=MLX_full |
| 63 | db/EUKARYO | Mlx_DBD | | RTCACGTGA | 10 | 10 | 8.9e-006 | CENTRIMO | | http://floresta.eead.csic.es/footprintdb/index.php?db=HumanTF:1.0&motif=Mlx_DBD |
| 64 | db/EUKARYO | SREBF2_DBD | | ATCACGTGA' | 10 | 10 | 8.9e-006 | CENTRIMO | | http://floresta.eead.csic.es/footprintdb/index.php?db=HumanTF:1.0&motif=SREBF2_DBD |
| 65 | db/JASPAR/J/ | MA0663.1 | MLX | RTCACGTGA | 10 | 10 | 8.9e-006 | CENTRIMO | | http://jaspar2018.genereg.net/matrix/MA0663.1 |
| 66 | db/EUKARYO | HOXD12_DBD_4 | | GYAATAAAA | 9 | 10 | 1.1e-005 | CENTRIMO | | http://floresta.eead.csic.es/footprintdb/index.php?db=HumanTF:1.0&motif=HOXD12_DBD_4 |
| 67 | db/JASPAR/J/ | MA0493.1 | Klf1 | RRCCACACC | 11 | 10 | 1.2e-005 | CENTRIMO | | http://jaspar2018.genereg.net/matrix/MA0493.1 |
| 68 | db/MOUSE/ur | UP00002_2 | Sp4_seconda | BWWAGGCG | 15 | 10 | 1.2e-005 | CENTRIMO | | http://the_brain.bwh.harvard.edu/uniprobe/details2.php?id=00002 |
| 69 | DREME | CCCCRCCC | DREME-11 | CCCCRCCC | 8 | 83 | 1.2e-005 | DREME | db/JASPAR/JASPAR **MA0599.1 (KLF5)** | http://jaspar2018.genereg.net/matrix/MA0599.1 |
| 70 | db/JASPAR/J/ | MA0599.1 | KLF5 | GCCCCDCCC | 10 | 10 | 1.2e-005 | CENTRIMO | | http://jaspar2018.genereg.net/matrix/MA0599.1 |
| 71 | db/MOUSE/ur | UP00093_1 | Klf7_primary | TNRRCCMC( | 16 | 10 | 1.3e-005 | CENTRIMO | | http://the_brain.bwh.harvard.edu/uniprobe/details2.php?id=00093 |
| 72 | db/EUKARYO | SP1_DBD | | RCCMCRCC( | 11 | 10 | 1.3e-005 | CENTRIMO | | http://floresta.eead.csic.es/footprintdb/index.php?db=HumanTF:1.0&motif=SP1_DBD |
| 73 | db/EUKARYO | SP4_full | | BWRGCCAC( | 17 | 10 | 1.3e-005 | CENTRIMO | | http://floresta.eead.csic.es/footprintdb/index.php?db=HumanTF:1.0&motif=SP4_full |
| 74 | db/JASPAR/J/ | MA0685.1 | SP4 | BWRGCCAC( | 17 | 10 | 1.3e-005 | CENTRIMO | | http://jaspar2018.genereg.net/matrix/MA0685.1 |
| 75 | db/EUKARYO | SP8_DBD | | RCCACGCCC | 12 | 10 | 1.3e-005 | CENTRIMO | | http://floresta.eead.csic.es/footprintdb/index.php?db=HumanTF:1.0&motif=SP8_DBD |
| 76 | db/JASPAR/J/ | MA0747.1 | SP8 | RCCACGCCC | 12 | 10 | 1.3e-005 | CENTRIMO | | http://jaspar2018.genereg.net/matrix/MA0747.1 |
| 77 | db/EUKARYO | KLF14_DBD | | KRCCACGCC | 14 | 10 | 1.3e-005 | CENTRIMO | | http://floresta.eead.csic.es/footprintdb/index.php?db=HumanTF:1.0&motif=KLF14_DBD |
| 78 | db/JASPAR/J/ | MA0740.1 | KLF14 | KRCCACGCC | 14 | 10 | 1.3e-005 | CENTRIMO | | http://jaspar2018.genereg.net/matrix/MA0740.1 |
| 79 | db/EUKARYO | Klf12_DBD | | GRCCACGC( | 15 | 10 | 1.3e-005 | CENTRIMO | | http://floresta.eead.csic.es/footprintdb/index.php?db=HumanTF:1.0&motif=Klf12_DBD |
| 80 | db/JASPAR/J/ | MA0742.1 | Klf12 | GRCCACGC( | 15 | 10 | 1.3e-005 | CENTRIMO | | http://jaspar2018.genereg.net/matrix/MA0742.1 |
| 81 | db/JASPAR/J/ | MA0516.1 | SP2 | GYCCCGCCY | 15 | 10 | 1.4e-005 | CENTRIMO | | http://jaspar2018.genereg.net/matrix/MA0516.1 |
| 82 | db/EUKARYO | CLOCK_DBD | | AACACGTGT | 10 | 10 | 1.6e-005 | CENTRIMO | | http://floresta.eead.csic.es/footprintdb/index.php?db=HumanTF:1.0&motif=CLOCK_DBD |
| 83 | db/JASPAR/J/ | MA0819.1 | CLOCK | AACACGTGT | 10 | 10 | 1.6e-005 | CENTRIMO | | http://jaspar2018.genereg.net/matrix/MA0819.1 |
| 84 | db/JASPAR/J/ | MA0825.1 | MNT | RVCACGTGM | 10 | 10 | 1.6e-005 | CENTRIMO | | http://jaspar2018.genereg.net/matrix/MA0825.1 |
| 85 | db/EUKARYO | GRHL1_full | | AAAACCGGT | 12 | 10 | 1.7e-005 | CENTRIMO | | http://floresta.eead.csic.es/footprintdb/index.php?db=HumanTF:1.0&motif=GRHL1_full |
| 86 | db/JASPAR/J/ | MA0647.1 | GRHL1 | AAAACCGGT | 12 | 10 | 1.7e-005 | CENTRIMO | | http://jaspar2018.genereg.net/matrix/MA0647.1 |
| 87 | db/EUKARYO | FOXO1_DBD_2 | | GTAAACATG' | 14 | 10 | 1.7e-005 | CENTRIMO | | http://floresta.eead.csic.es/footprintdb/index.php?db=HumanTF:1.0&motif=FOXO1_DBD_2 |
| 88 | db/EUKARYO | FOXO3_full_1 | | GTAAACATG' | 14 | 10 | 1.7e-005 | CENTRIMO | | http://floresta.eead.csic.es/footprintdb/index.php?db=HumanTF:1.0&motif=FOXO3_full_1 |
| 89 | db/EUKARYO | FOXO4_DBD_1 | | GTMAACATG | 14 | 10 | 1.7e-005 | CENTRIMO | | http://floresta.eead.csic.es/footprintdb/index.php?db=HumanTF:1.0&motif=FOXO4_DBD_1 |
| 90 | db/EUKARYO | FOXO6_DBD_1 | | GTAAACATG' | 14 | 10 | 1.7e-005 | CENTRIMO | | http://floresta.eead.csic.es/footprintdb/index.php?db=HumanTF:1.0&motif=FOXO6_DBD_1 |
| 91 | db/MOUSE/ur | UP00002_1 | Sp4_primary | BDHMMCGC( | 17 | 10 | 3.9e-005 | CENTRIMO | | http://the_brain.bwh.harvard.edu/uniprobe/details2.php?id=00002 |
| 92 | db/JASPAR/J/ | MA0732.1 | EGR3 | HHMCGCCC/ | 15 | 10 | 4.1e-005 | CENTRIMO | | http://jaspar2018.genereg.net/matrix/MA0732.1 |
| 93 | db/MOUSE/ur | UP00007_1 | Egr1_primary | NCCGCCCC( | 14 | 10 | 4.2e-005 | CENTRIMO | | http://the_brain.bwh.harvard.edu/uniprobe/details2.php?id=00007 |
| 94 | db/EUKARYO | EGR4_DBD_1 | | NHMCGCCC/ | 16 | 10 | 4.2e-005 | CENTRIMO | | http://floresta.eead.csic.es/footprintdb/index.php?db=HumanTF:1.0&motif=EGR4_DBD_1 |
| 95 | db/EUKARYO | Egr1_mouse_DBD_mutant_I | | HHMCGCCCI | 16 | 10 | 4.2e-005 | CENTRIMO | | http://floresta.eead.csic.es/footprintdb/index.php?db=HumanTF:1.0&motif=Egr1_mouse_DBD_mutant_DBD |
| 96 | db/JASPAR/J/ | MA0733.1 | EGR4 | NHMCGCCC/ | 16 | 10 | 4.2e-005 | CENTRIMO | | http://jaspar2018.genereg.net/matrix/MA0733.1 |
| 97 | db/MOUSE/ur | UP00024_1 | Glis2_primary | YWHNGACC( | 16 | 10 | 4.2e-005 | CENTRIMO | | http://the_brain.bwh.harvard.edu/uniprobe/details2.php?id=00024 |
| 98 | db/MOUSE/ur | UP00022_1 | Zfp740_prima | YSNCCCCCC | 16 | 10 | 4.2e-005 | CENTRIMO | | http://the_brain.bwh.harvard.edu/uniprobe/details2.php?id=00022 |
| 99 | db/EUKARYO | ZNF740_DBD | | MCCCCCCC/ | 10 | 10 | 4.3e-005 | CENTRIMO | | http://floresta.eead.csic.es/footprintdb/index.php?db=HumanTF:1.0&motif=ZNF740_DBD |
| 100 | db/EUKARYO | ZNF740_full | | CCCCCCCCA | 10 | 10 | 4.3e-005 | CENTRIMO | | http://floresta.eead.csic.es/footprintdb/index.php?db=HumanTF:1.0&motif=ZNF740_full |
| 101 | db/EUKARYO | Zfp740_DBD | | VCCCCCCCA | 10 | 10 | 4.3e-005 | CENTRIMO | | http://floresta.eead.csic.es/footprintdb/index.php?db=HumanTF:1.0&motif=Zfp740_DBD |
| 102 | db/JASPAR/J/ | MA0753.1 | ZNF740 | MCCCCCCC/ | 10 | 10 | 4.3e-005 | CENTRIMO | | http://jaspar2018.genereg.net/matrix/MA0753.1 |
| 103 | db/EUKARYO | Zic3_DBD | | GACCCCCYC | 15 | 10 | 4.8e-005 | CENTRIMO | | http://floresta.eead.csic.es/footprintdb/index.php?db=HumanTF:1.0&motif=Zic3_DBD |
| 104 | db/MOUSE/ur | UP00070_2 | Gcm1_second | DKSNNATAG | 17 | 10 | 4.9e-005 | CENTRIMO | | http://the_brain.bwh.harvard.edu/uniprobe/details2.php?id=00070 |
| 105 | db/JASPAR/J/ | MA0158.1 | HOXA5 | CDBWAATK | 8 | 10 | 4.9e-005 | CENTRIMO | | http://jaspar2018.genereg.net/matrix/MA0158.1 |
| 106 | db/MOUSE/ur | UP00040_2 | Irf5_secondar | BWNAYCGAC | 15 | 10 | 6.2e-005 | CENTRIMO | | http://the_brain.bwh.harvard.edu/uniprobe/details2.php?id=00040 |

| | | | | | | | | | | | |
|---|---|---|---|---|---|---|---|---|---|---|---|
| 107 | db/JASPAR/J/ | MA0119.1 | NFIC::TLX1 | TGGCASSRW | 14 | 10 | 6.8e-005 | CENTRIMO | | | http://jaspar2018.genereg.net/matrix/MA0119.1 |
| 108 | db/EUKARYO | NFIX_full_4 | | YTGGCAHND | 15 | 10 | 7.0e-005 | CENTRIMO | | | http://floresta.eead.csic.es/footprintdb/index.php?db=HumanTF:1.0&motif=NFIX_full_4 |
| 109 | db/EUKARYO | HEY2_DBD | | GRCACGTGC | 10 | 10 | 7.1e-005 | CENTRIMO | | | http://floresta.eead.csic.es/footprintdb/index.php?db=HumanTF:1.0&motif=HEY2_DBD |
| 110 | db/JASPAR/J/ | MA0147.3 | MYC | NNCCACGTG | 12 | 10 | 7.1e-005 | CENTRIMO | | | http://jaspar2018.genereg.net/matrix/MA0147.3 |
| 111 | db/MOUSE/ur | UP00093_2 | Klf7_secondar | VRNBATACG | 17 | 10 | 8.9e-005 | CENTRIMO | | | http://the_brain.bwh.harvard.edu/uniprobe/details2.php?id=00093 |
| 112 | db/EUKARYO | ZBTB7A_DBD | | BGCGACCAC | 12 | 10 | 9.7e-005 | CENTRIMO | | | http://floresta.eead.csic.es/footprintdb/index.php?db=HumanTF:1.0&motif=ZBTB7A_DBD |
| 113 | DREME | GTGTGTGY | DREME-7 | GTGTGTGY | 8 | 207 | 1.0e-004 | DREME | db/MOUSE/uniprobe | **UP00042_2 (Gm397_** | http://the_brain.bwh.harvard.edu/uniprobe/details2.php?id=00042 |
| 114 | db/JASPAR/J/ | MA1102.1 | CTCFL | CRSCAGGGC | 14 | 10 | 1.0e-004 | CENTRIMO | | | http://jaspar2018.genereg.net/matrix/MA1102.1 |
| 115 | db/EUKARYO | TFAP2A_DBD_1 | | YGCCCBVRG | 12 | 10 | 1.1e-004 | CENTRIMO | | | http://floresta.eead.csic.es/footprintdb/index.php?db=HumanTF:1.0&motif=TFAP2A_DBD_1 |
| 116 | db/EUKARYO | TFAP2A_DBD_5 | | TGCCCBVRG | 12 | 10 | 1.1e-004 | CENTRIMO | | | http://floresta.eead.csic.es/footprintdb/index.php?db=HumanTF:1.0&motif=TFAP2A_DBD_5 |
| 117 | db/EUKARYO | TFAP2B_DBD_1 | | YGCCCBVRG | 12 | 10 | 1.1e-004 | CENTRIMO | | | http://floresta.eead.csic.es/footprintdb/index.php?db=HumanTF:1.0&motif=TFAP2B_DBD_1 |
| 118 | db/EUKARYO | TFAP2C_DBD_1 | | YGCCYBVRG | 12 | 10 | 1.1e-004 | CENTRIMO | | | http://floresta.eead.csic.es/footprintdb/index.php?db=HumanTF:1.0&motif=TFAP2C_DBD_1 |
| 119 | db/EUKARYO | TFAP2C_full_1 | | HGCCCBVRG | 12 | 10 | 1.1e-004 | CENTRIMO | | | http://floresta.eead.csic.es/footprintdb/index.php?db=HumanTF:1.0&motif=TFAP2C_full_1 |
| 120 | db/EUKARYO | Tcfap2a_DBD_1 | | TGCCCYVRG | 12 | 10 | 1.1e-004 | CENTRIMO | | | http://floresta.eead.csic.es/footprintdb/index.php?db=HumanTF:1.0&motif=Tcfap2a_DBD_1 |
| 121 | db/JASPAR/J/ | MA0810.1 | TFAP2A(var.2 | YGCCCBVRG | 12 | 10 | 1.1e-004 | CENTRIMO | | | http://jaspar2018.genereg.net/matrix/MA0810.1 |
| 122 | db/JASPAR/J/ | MA0811.1 | TFAP2B | YGCCCBVRG | 12 | 10 | 1.1e-004 | CENTRIMO | | | http://jaspar2018.genereg.net/matrix/MA0811.1 |
| 123 | db/JASPAR/J/ | MA0524.2 | TFAP2C | YGCCYBVRG | 12 | 10 | 1.1e-004 | CENTRIMO | | | http://jaspar2018.genereg.net/matrix/MA0524.2 |
| 124 | db/MOUSE/ur | UP00005_2 | Tcfap2a_seco | YHRCCYBWC | 14 | 10 | 1.1e-004 | CENTRIMO | | | http://the_brain.bwh.harvard.edu/uniprobe/details2.php?id=00005 |
| 125 | db/MOUSE/ur | UP00087_2 | Tcfap2c_seco | NYGCCCDAC | 14 | 10 | 1.1e-004 | CENTRIMO | | | http://the_brain.bwh.harvard.edu/uniprobe/details2.php?id=00087 |
| 126 | db/MOUSE/ur | UP00028_1 | Tcfap2e_prima | WNHGCCTS/ | 15 | 10 | 1.1e-004 | CENTRIMO | | | http://the_brain.bwh.harvard.edu/uniprobe/details2.php?id=00028 |
| 127 | db/EUKARYO | THRB_DBD_3 | | DTGACCTYAI | 20 | 10 | 1.2e-004 | CENTRIMO | | | http://floresta.eead.csic.es/footprintdb/index.php?db=HumanTF:1.0&motif=THRB_DBD_3 |
| 128 | db/JASPAR/J/ | MA0609.1 | Crem | NATGACGTM | 10 | 10 | 1.6e-004 | CENTRIMO | | | http://jaspar2018.genereg.net/matrix/MA0609.1 |
| 129 | db/MOUSE/ur | UP00147_1 | Nkx2-6_3437. | NRADCCACT | 16 | 10 | 1.7e-004 | CENTRIMO | | | http://the_brain.bwh.harvard.edu/uniprobe/details2.php?id=00147 |
| 130 | db/MOUSE/ur | UP00165_1 | Titf1_1722.2 | YVAGCCACT | 16 | 10 | 1.7e-004 | CENTRIMO | | | http://the_brain.bwh.harvard.edu/uniprobe/details2.php?id=00165 |
| 131 | DREME | GAGCCABC | DREME-8 | GAGCCABC | 8 | 89 | 2.6e-004 | DREME | db/JASPAR/JASPAR | **MA0099.3 (FOS::JUN** | http://jaspar2018.genereg.net/matrix/MA0099.3 |
| 132 | DREME | GGCTGGM | DREME-9 | GGCTGGM | 7 | 170 | 2.7e-004 | DREME | db/JASPAR/JASPAR | **MA1121.1 (TEAD2)** | http://jaspar2018.genereg.net/matrix/MA1121.1 |
| 133 | db/EUKARYO | EGR3_DBD | | HHMCGCCC/ | 15 | 10 | 3.7e-004 | CENTRIMO | | | http://floresta.eead.csic.es/footprintdb/index.php?db=HumanTF:1.0&motif=EGR3_DBD |
| 134 | db/EUKARYO | EGR1_full | | HACGCCCAC | 14 | 10 | 3.7e-004 | CENTRIMO | | | http://floresta.eead.csic.es/footprintdb/index.php?db=HumanTF:1.0&motif=EGR1_full |
| 135 | db/JASPAR/J/ | MA0162.3 | EGR1 | HACGCCCAC | 14 | 10 | 3.7e-004 | CENTRIMO | | | http://jaspar2018.genereg.net/matrix/MA0162.3 |
| 136 | db/EUKARYO | Egr3_DBD | | HACGCCCAC | 15 | 10 | 3.8e-004 | CENTRIMO | | | http://floresta.eead.csic.es/footprintdb/index.php?db=HumanTF:1.0&motif=Egr3_DBD |
| 137 | db/EUKARYO | TFCP2_full_2 | | ACCGGTTYA. | 16 | 10 | 5.7e-004 | CENTRIMO | | | http://floresta.eead.csic.es/footprintdb/index.php?db=HumanTF:1.0&motif=TFCP2_full_2 |
| 138 | db/MOUSE/ur | UP00053_2 | Rxra_seconda | KCRCRWAGK | 16 | 10 | 6.7e-004 | CENTRIMO | | | http://the_brain.bwh.harvard.edu/uniprobe/details2.php?id=00053 |
| 139 | DREME | CAGAA | DREME-10 | CAGAA | 5 | 791 | 9.6e-004 | DREME | | | |
| 140 | DREME | CYGGGTCC | DREME-16 | CYGGGTCC | 8 | 29 | 1.0e-003 | DREME | db/MOUSE/uniprobe | **UP00015_1 (Ehf_prim** | http://the_brain.bwh.harvard.edu/uniprobe/details2.php?id=00015 |
| 141 | db/EUKARYO | FOXJ2_DBD_2 | | RTAAACAA | 8 | 10 | 1.1e-003 | CENTRIMO | | | http://floresta.eead.csic.es/footprintdb/index.php?db=HumanTF:1.0&motif=FOXJ2_DBD_2 |
| 142 | db/EUKARYO | FOXO1_DBD_1 | | GTAAACAW | 8 | 10 | 1.1e-003 | CENTRIMO | | | http://floresta.eead.csic.es/footprintdb/index.php?db=HumanTF:1.0&motif=FOXO1_DBD_1 |
| 143 | db/EUKARYO | Foxj3_DBD_3 | | RTAAACAA | 8 | 10 | 1.1e-003 | CENTRIMO | | | http://floresta.eead.csic.es/footprintdb/index.php?db=HumanTF:1.0&motif=Foxj3_DBD_3 |
| 144 | db/JASPAR/J/ | MA0613.1 | FOXG1 | RTAAACAW | 8 | 10 | 1.1e-003 | CENTRIMO | | | http://jaspar2018.genereg.net/matrix/MA0613.1 |
| 145 | db/JASPAR/J/ | MA0614.1 | Foxj2 | RTAAACAA | 8 | 10 | 1.1e-003 | CENTRIMO | | | http://jaspar2018.genereg.net/matrix/MA0614.1 |
| 146 | db/EUKARYO | FOXD2_DBD_2 | | GTAAACA | 7 | 10 | 1.1e-003 | CENTRIMO | | | http://floresta.eead.csic.es/footprintdb/index.php?db=HumanTF:1.0&motif=FOXD2_DBD_2 |
| 147 | db/EUKARYO | FOXL1_full_1 | | RTAAACA | 7 | 10 | 1.1e-003 | CENTRIMO | | | http://floresta.eead.csic.es/footprintdb/index.php?db=HumanTF:1.0&motif=FOXL1_full_1 |
| 148 | db/EUKARYO | FOXO4_DBD_2 | | GTAAACA | 7 | 10 | 1.1e-003 | CENTRIMO | | | http://floresta.eead.csic.es/footprintdb/index.php?db=HumanTF:1.0&motif=FOXO4_DBD_2 |
| 149 | db/EUKARYO | FOXO6_DBD_2 | | GTAAACA | 7 | 10 | 1.1e-003 | CENTRIMO | | | http://floresta.eead.csic.es/footprintdb/index.php?db=HumanTF:1.0&motif=FOXO6_DBD_2 |
| 150 | db/EUKARYO | FOXP3_DBD | | RTAAACA | 7 | 10 | 1.1e-003 | CENTRIMO | | | http://floresta.eead.csic.es/footprintdb/index.php?db=HumanTF:1.0&motif=FOXP3_DBD |
| 151 | db/EUKARYO | Foxc1_DBD_2 | | RTAAACA | 7 | 10 | 1.1e-003 | CENTRIMO | | | http://floresta.eead.csic.es/footprintdb/index.php?db=HumanTF:1.0&motif=Foxc1_DBD_2 |
| 152 | db/EUKARYO | Foxg1_DBD_3 | | GTAAACA | 7 | 10 | 1.1e-003 | CENTRIMO | | | http://floresta.eead.csic.es/footprintdb/index.php?db=HumanTF:1.0&motif=Foxg1_DBD_3 |
| 153 | db/EUKARYO | Foxk1_DBD_2 | | RTAAACA | 7 | 10 | 1.1e-003 | CENTRIMO | | | http://floresta.eead.csic.es/footprintdb/index.php?db=HumanTF:1.0&motif=Foxk1_DBD_2 |
| 154 | db/JASPAR/J/ | MA0033.2 | FOXL1 | RTAAACA | 7 | 10 | 1.1e-003 | CENTRIMO | | | http://jaspar2018.genereg.net/matrix/MA0033.2 |
| 155 | db/JASPAR/J/ | MA0847.1 | FOXD2 | GTAAACA | 7 | 10 | 1.1e-003 | CENTRIMO | | | http://jaspar2018.genereg.net/matrix/MA0847.1 |
| 156 | db/JASPAR/J/ | MA0848.1 | FOXO4 | GTAAACA | 7 | 10 | 1.1e-003 | CENTRIMO | | | http://jaspar2018.genereg.net/matrix/MA0848.1 |
| 157 | db/JASPAR/J/ | MA0849.1 | FOXO6 | GTAAACA | 7 | 10 | 1.1e-003 | CENTRIMO | | | http://jaspar2018.genereg.net/matrix/MA0849.1 |
| 158 | db/JASPAR/J/ | MA0850.1 | FOXP3 | RTAAACA | 7 | 10 | 1.1e-003 | CENTRIMO | | | http://jaspar2018.genereg.net/matrix/MA0850.1 |
| 159 | db/EUKARYO | RARG_full_3 | | RAGGTCAHB | 17 | 10 | 1.5e-003 | CENTRIMO | | | http://floresta.eead.csic.es/footprintdb/index.php?db=HumanTF:1.0&motif=RARG_full_3 |
| 160 | db/EUKARYO | RXRA_full_2 | | RRGGTCATG | 14 | 10 | 1.6e-003 | CENTRIMO | | | http://floresta.eead.csic.es/footprintdb/index.php?db=HumanTF:1.0&motif=RXRA_full_2 |
| 161 | db/EUKARYO | GCM1_full_2 | | BATGCGGGT | 11 | 10 | 1.6e-003 | CENTRIMO | | | http://floresta.eead.csic.es/footprintdb/index.php?db=HumanTF:1.0&motif=GCM1_full_2 |
| 162 | db/JASPAR/J/ | MA0646.1 | GCM1 | BATGCGGGT | 11 | 10 | 1.6e-003 | CENTRIMO | | | http://jaspar2018.genereg.net/matrix/MA0646.1 |
| 163 | db/JASPAR/J/ | MA0597.1 | THAP1 | YTGCCCDBA | 9 | 10 | 1.6e-003 | CENTRIMO | | | http://jaspar2018.genereg.net/matrix/MA0597.1 |
| 164 | db/EUKARYO | ESRRB_DBD | | TCAAGGTCA | 11 | 10 | 1.7e-003 | CENTRIMO | | | http://floresta.eead.csic.es/footprintdb/index.php?db=HumanTF:1.0&motif=ESRRB_DBD |
| 165 | db/JASPAR/J/ | MA0141.3 | ESRRB | TCAAGGTCA | 11 | 10 | 1.7e-003 | CENTRIMO | | | http://jaspar2018.genereg.net/matrix/MA0141.3 |
| 166 | db/JASPAR/J/ | MA1111.1 | NR2F2 | NAAAGGTCA | 11 | 10 | 1.7e-003 | CENTRIMO | | | http://jaspar2018.genereg.net/matrix/MA1111.1 |
| 167 | db/MOUSE/ur | UP00079_1 | Esrra_primary | NMNYCAAGC | 17 | 10 | 1.8e-003 | CENTRIMO | | | http://the_brain.bwh.harvard.edu/uniprobe/details2.php?id=00079 |

| | | | | | | | | | | | |
|---|---|---|---|---|---|---|---|---|---|---|---|
| 168 | db/EUKARYO | ESRRG_full_3 | | TCAAGGTCA | 10 | 10 | 1.8e-003 | CENTRIMO | | | http://floresta.eead.csic.es/footprintdb/index.php?db=HumanTF:1.0&motif=ESRRG_full_3 |
| 169 | db/JASPAR/J | MA0643.1 | Esrrg | TCAAGGTCA | 10 | 10 | 1.8e-003 | CENTRIMO | | | http://jaspar2018.genereg.net/matrix/MA0643.1 |
| 170 | db/JASPAR/J | MA1112.1 | NR4A1 | NAAAGGTCA | 10 | 10 | 1.8e-003 | CENTRIMO | | | http://jaspar2018.genereg.net/matrix/MA1112.1 |
| 171 | db/EUKARYO | ESRRA_DBD_1 | | BTCAAGGTC | 11 | 10 | 1.8e-003 | CENTRIMO | | | http://floresta.eead.csic.es/footprintdb/index.php?db=HumanTF:1.0&motif=ESRRA_DBD_1 |
| 172 | db/EUKARYO | ESRRA_DBD_4 | | BTCAAGGTC | 11 | 10 | 1.8e-003 | CENTRIMO | | | http://floresta.eead.csic.es/footprintdb/index.php?db=HumanTF:1.0&motif=ESRRA_DBD_4 |
| 173 | db/EUKARYO | Esrra_DBD_2 | | TTCAAGGTC | 11 | 10 | 1.8e-003 | CENTRIMO | | | http://floresta.eead.csic.es/footprintdb/index.php?db=HumanTF:1.0&motif=Esrra_DBD_2 |
| 174 | db/JASPAR/J | MA0592.2 | Esrra | TTCAAGGTC | 11 | 10 | 1.8e-003 | CENTRIMO | | | http://jaspar2018.genereg.net/matrix/MA0592.2 |
| 175 | db/JASPAR/J | MA0071.1 | RORA | AWMWAGGT | 10 | 10 | 1.8e-003 | CENTRIMO | | | http://jaspar2018.genereg.net/matrix/MA0071.1 |
| 176 | db/EUKARYO | NR4A2_full_3 | | TTTAAAGGT | 11 | 10 | 1.8e-003 | CENTRIMO | | | http://floresta.eead.csic.es/footprintdb/index.php?db=HumanTF:1.0&motif=NR4A2_full_3 |
| 177 | db/JASPAR/J | MA0505.1 | Nr5a2 | RAGTTCAAG | 15 | 10 | 1.9e-003 | CENTRIMO | | | http://jaspar2018.genereg.net/matrix/MA0505.1 |
| 178 | DREME | TACATW | DREME-12 | TACATW | 6 | 300 | 1.9e-003 | DREME | db/EUKARYOTE/joln | **FOXB1_DBD_3** | http://floresta.eead.csic.es/footprintdb/index.php?db=HumanTF:1.0&motif=FOXB1_DBD_3 |
| 179 | db/EUKARYO | NR2F1_DBD_1 | | RRGGTCAAA | 15 | 10 | 1.9e-003 | CENTRIMO | | | http://floresta.eead.csic.es/footprintdb/index.php?db=HumanTF:1.0&motif=NR2F1_DBD_1 |
| 180 | db/EUKARYO | NR2F6_full | | RRGGTCAAA | 14 | 10 | 1.9e-003 | CENTRIMO | | | http://floresta.eead.csic.es/footprintdb/index.php?db=HumanTF:1.0&motif=NR2F6_full |
| 181 | db/EUKARYO | RXRA_full_1 | | GRGGTCAAA | 14 | 10 | 1.9e-003 | CENTRIMO | | | http://floresta.eead.csic.es/footprintdb/index.php?db=HumanTF:1.0&motif=RXRA_full_1 |
| 182 | db/EUKARYO | Rxra_DBD_1 | | RRGGTCAAA | 14 | 10 | 1.9e-003 | CENTRIMO | | | http://floresta.eead.csic.es/footprintdb/index.php?db=HumanTF:1.0&motif=Rxra_DBD_1 |
| 183 | db/JASPAR/J | MA0677.1 | Nr2f6 | RRGGTCAAA | 14 | 10 | 1.9e-003 | CENTRIMO | | | http://jaspar2018.genereg.net/matrix/MA0677.1 |
| 184 | DREME | CCACGYG | DREME-13 | CCACGYG | 7 | 35 | 2.2e-003 | DREME | db/JASPAR/JASPAR | **MA0059.1 (MAX::MY** | http://jaspar2018.genereg.net/matrix/MA0059.1 |
| 185 | db/JASPAR/J | MA0159.1 | RARA::RXRA | RGKTCANVG | 17 | 10 | 2.4e-003 | CENTRIMO | | | http://jaspar2018.genereg.net/matrix/MA0159.1 |
| 186 | db/EUKARYO | ESRRG_full_1 | | AAGGTCAYY | 18 | 10 | 3.4e-003 | CENTRIMO | | | http://floresta.eead.csic.es/footprintdb/index.php?db=HumanTF:1.0&motif=ESRRG_full_1 |
| 187 | db/EUKARYO | ESRRA_DBD_3 | | CAAGGTCAH | 19 | 10 | 3.5e-003 | CENTRIMO | | | http://floresta.eead.csic.es/footprintdb/index.php?db=HumanTF:1.0&motif=ESRRA_DBD_3 |
| 188 | db/EUKARYO | ESRRA_DBD_6 | | CAAGGTCAH | 19 | 10 | 3.5e-003 | CENTRIMO | | | http://floresta.eead.csic.es/footprintdb/index.php?db=HumanTF:1.0&motif=ESRRA_DBD_6 |
| 189 | db/EUKARYO | NR2E1_full_2 | | AAGTCAAWA | 14 | 10 | 4.0e-003 | CENTRIMO | | | http://floresta.eead.csic.es/footprintdb/index.php?db=HumanTF:1.0&motif=NR2E1_full_2 |
| 190 | db/EUKARYO | Nr2e1_DBD_2 | | AAGTCAADA | 14 | 10 | 4.0e-003 | CENTRIMO | | | http://floresta.eead.csic.es/footprintdb/index.php?db=HumanTF:1.0&motif=Nr2e1_DBD_2 |
| 191 | DREME | AGWCAGGG | DREME-14 | AGWCAGGG | 8 | 60 | 4.0e-003 | DREME | | | |
| 192 | db/MOUSE/ur | UP00032_2 | Gata3_second | YTHNVYWGA | 22 | 10 | 8.1e-003 | CENTRIMO | | | http://the_brain.bwh.harvard.edu/uniprobe/details2.php?id=00032 |
| 193 | db/JASPAR/J | MA0831.2 | TFE3 | CACGTGAY | 8 | 10 | 8.3e-003 | CENTRIMO | | | http://jaspar2018.genereg.net/matrix/MA0831.2 |
| 194 | db/JASPAR/J | MA0603.1 | Arntl | NGTCACGTG | 10 | 10 | 8.3e-003 | CENTRIMO | | | http://jaspar2018.genereg.net/matrix/MA0603.1 |
| 195 | db/JASPAR/J | MA0093.2 | USF1 | VYCAYGTGA | 11 | 10 | 8.4e-003 | CENTRIMO | | | http://jaspar2018.genereg.net/matrix/MA0093.2 |
| 196 | db/EUKARYO | MLXIPL_full | | RTCACGTGA | 10 | 10 | 8.5e-003 | CENTRIMO | | | http://floresta.eead.csic.es/footprintdb/index.php?db=HumanTF:1.0&motif=MLXIPL_full |
| 197 | db/EUKARYO | TFEB_full | | RYCACGTGA | 10 | 10 | 8.5e-003 | CENTRIMO | | | http://floresta.eead.csic.es/footprintdb/index.php?db=HumanTF:1.0&motif=TFEB_full |
| 198 | db/EUKARYO | TFEC_DBD | | RTCACRTGA | 10 | 10 | 8.5e-003 | CENTRIMO | | | http://floresta.eead.csic.es/footprintdb/index.php?db=HumanTF:1.0&motif=TFEC_DBD |
| 199 | db/EUKARYO | USF1_DBD | | RHCACGTGA | 10 | 10 | 8.5e-003 | CENTRIMO | | | http://floresta.eead.csic.es/footprintdb/index.php?db=HumanTF:1.0&motif=USF1_DBD |
| 200 | db/JASPAR/J | MA0664.1 | MLXIPL | RTCACGTGA | 10 | 10 | 8.5e-003 | CENTRIMO | | | http://jaspar2018.genereg.net/matrix/MA0664.1 |
| 201 | db/JASPAR/J | MA0692.1 | TFEB | RYCACGTGA | 10 | 10 | 8.5e-003 | CENTRIMO | | | http://jaspar2018.genereg.net/matrix/MA0692.1 |
| 202 | db/JASPAR/J | MA0871.1 | TFEC | RTCACRTGA | 10 | 10 | 8.5e-003 | CENTRIMO | | | http://jaspar2018.genereg.net/matrix/MA0871.1 |
| 203 | db/EUKARYO | MEIS3_DBD_2 | | TGACAGSTG | 12 | 10 | 8.6e-003 | CENTRIMO | | | http://floresta.eead.csic.es/footprintdb/index.php?db=HumanTF:1.0&motif=MEIS3_DBD_2 |
| 204 | db/EUKARYO | Meis2_DBD_2 | | TGACAGSTG | 12 | 10 | 8.6e-003 | CENTRIMO | | | http://floresta.eead.csic.es/footprintdb/index.php?db=HumanTF:1.0&motif=Meis2_DBD_2 |
| 205 | db/EUKARYO | TGIF2_DBD | | TGACAGSTG | 12 | 10 | 8.6e-003 | CENTRIMO | | | http://floresta.eead.csic.es/footprintdb/index.php?db=HumanTF:1.0&motif=TGIF2_DBD |
| 206 | db/JASPAR/J | MA0797.1 | TGIF2 | TGACAGSTG | 12 | 10 | 8.6e-003 | CENTRIMO | | | http://jaspar2018.genereg.net/matrix/MA0797.1 |
| 207 | db/MOUSE/ur | UP00060_2 | Max_seconda | BYGNCACGC | 14 | 10 | 8.6e-003 | CENTRIMO | | | http://the_brain.bwh.harvard.edu/uniprobe/details2.php?id=00060 |
| 208 | db/JASPAR/J | MA0620.2 | MITF | NNDRGTCAC | 18 | 10 | 8.6e-003 | CENTRIMO | | | http://jaspar2018.genereg.net/matrix/MA0620.2 |
| 209 | db/EUKARYO | HOMEZ_DBD | | AAAACGATT | 12 | 10 | 8.8e-003 | CENTRIMO | | | http://floresta.eead.csic.es/footprintdb/index.php?db=HumanTF:1.0&motif=HOMEZ_DBD |
| 210 | db/MOUSE/ur | UP00114_1 | Homez_1063. | AWWWMATC | 17 | 10 | 9.2e-003 | CENTRIMO | | | http://the_brain.bwh.harvard.edu/uniprobe/details2.php?id=00114 |
| 211 | db/EUKARYO | HOXD12_DBD_2 | | RGTCGTAAA | 11 | 10 | 1.0e-002 | CENTRIMO | | | http://floresta.eead.csic.es/footprintdb/index.php?db=HumanTF:1.0&motif=HOXD12_DBD_2 |
| 212 | DREME | ATGRGATC | DREME-17 | ATGRGATC | 8 | 34 | 1.1e-002 | DREME | | | |
| 213 | db/EUKARYO | T_full | | TCACACMTA | 16 | 10 | 1.2e-002 | CENTRIMO | | | http://floresta.eead.csic.es/footprintdb/index.php?db=HumanTF:1.0&motif=T_full |
| 214 | db/JASPAR/J | MA0009.2 | T | TCACACMTA | 16 | 10 | 1.2e-002 | CENTRIMO | | | http://jaspar2018.genereg.net/matrix/MA0009.2 |
| 215 | db/EUKARYO | TBX19_DBD | | DTTMRCACM | 20 | 10 | 1.2e-002 | CENTRIMO | | | http://floresta.eead.csic.es/footprintdb/index.php?db=HumanTF:1.0&motif=TBX19_DBD |
| 216 | db/JASPAR/J | MA0804.1 | TBX19 | DTTMRCACM | 20 | 10 | 1.2e-002 | CENTRIMO | | | http://jaspar2018.genereg.net/matrix/MA0804.1 |
| 217 | db/MOUSE/ur | UP00100_2 | Gata6_second | GSBNCGATA | 17 | 10 | 1.2e-002 | CENTRIMO | | | http://the_brain.bwh.harvard.edu/uniprobe/details2.php?id=00100 |
| 218 | db/JASPAR/J | MA0500.1 | Myog | RRCAGCTGS | 11 | 10 | 1.2e-002 | CENTRIMO | | | http://jaspar2018.genereg.net/matrix/MA0500.1 |
| 219 | db/JASPAR/J | MA0521.1 | Tcf12 | RACAGCTGS | 11 | 10 | 1.2e-002 | CENTRIMO | | | http://jaspar2018.genereg.net/matrix/MA0521.1 |
| 220 | db/EUKARYO | ZNF238_full | | NATCCAGAT | 13 | 10 | 1.2e-002 | CENTRIMO | | | http://floresta.eead.csic.es/footprintdb/index.php?db=HumanTF:1.0&motif=ZNF238_full |
| 221 | db/JASPAR/J | MA0698.1 | ZBTB18 | NATCCAGAT | 13 | 10 | 1.2e-002 | CENTRIMO | | | http://jaspar2018.genereg.net/matrix/MA0698.1 |
| 222 | db/EUKARYO | FIGLA_DBD | | WMCACCTG | 10 | 10 | 1.3e-002 | CENTRIMO | | | http://floresta.eead.csic.es/footprintdb/index.php?db=HumanTF:1.0&motif=FIGLA_DBD |
| 223 | db/EUKARYO | MSC_full | | AACAGCTGT | 10 | 10 | 1.3e-002 | CENTRIMO | | | http://floresta.eead.csic.es/footprintdb/index.php?db=HumanTF:1.0&motif=MSC_full |
| 224 | db/JASPAR/J | MA0665.1 | MSC | AACAGCTGT | 10 | 10 | 1.3e-002 | CENTRIMO | | | http://jaspar2018.genereg.net/matrix/MA0665.1 |
| 225 | db/JASPAR/J | MA0820.1 | FIGLA | WMCACCTG | 10 | 10 | 1.3e-002 | CENTRIMO | | | http://jaspar2018.genereg.net/matrix/MA0820.1 |
| 226 | db/EUKARYO | Tcf21_DBD | | RYAACAGCT | 14 | 10 | 1.3e-002 | CENTRIMO | | | http://floresta.eead.csic.es/footprintdb/index.php?db=HumanTF:1.0&motif=Tcf21_DBD |
| 227 | db/JASPAR/J | MA0832.1 | Tcf21 | RYAACAGCT | 14 | 10 | 1.3e-002 | CENTRIMO | | | http://jaspar2018.genereg.net/matrix/MA0832.1 |
| 228 | db/JASPAR/J | MA1100.1 | ASCL1 | VSAGCAGCT | 13 | 10 | 1.3e-002 | CENTRIMO | | | http://jaspar2018.genereg.net/matrix/MA1100.1 |

| | | | | | | | | | |
|---|---|---|---|---|---|---|---|---|---|
| 229 | db/JASPAR/J | MA0499.1 | Myod1 | NSCAGCTGY | 13 | 10 | 1.3e-002 | CENTRIMO | http://jaspar2018.genereg.net/matrix/MA0499.1 |
| 230 | db/EUKARYO | EGR2_DBD | | MCGCCCAC | 11 | 10 | 1.8e-002 | CENTRIMO | http://floresta.eead.csic.es/footprintdb/index.php?db=HumanTF:1.0&motif=EGR2_DBD |
| 231 | db/JASPAR/J | MA0472.2 | EGR2 | MCGCCCAC | 11 | 10 | 1.8e-002 | CENTRIMO | http://jaspar2018.genereg.net/matrix/MA0472.2 |
| 232 | db/JASPAR/J | MA0601.1 | Arid3b | ATATTAATWA | 11 | 10 | 1.9e-002 | CENTRIMO | http://jaspar2018.genereg.net/matrix/MA0601.1 |
| 233 | db/EUKARYO | HOXC10_DBD_1 | | VCMATWAAA | 10 | 10 | 1.9e-002 | CENTRIMO | http://floresta.eead.csic.es/footprintdb/index.php?db=HumanTF:1.0&motif=HOXC10_DBD_1 |
| 234 | db/EUKARYO | Hoxc10_DBD_2 | | GYAATAAAA | 10 | 10 | 1.9e-002 | CENTRIMO | http://floresta.eead.csic.es/footprintdb/index.php?db=HumanTF:1.0&motif=Hoxc10_DBD_2 |
| 235 | db/EUKARYO | Hoxd9_DBD_2 | | GYAATWAAA | 10 | 10 | 1.9e-002 | CENTRIMO | http://floresta.eead.csic.es/footprintdb/index.php?db=HumanTF:1.0&motif=Hoxd9_DBD_2 |
| 236 | db/JASPAR/J | MA0913.1 | Hoxd9 | GYAATWAAA | 10 | 10 | 1.9e-002 | CENTRIMO | http://jaspar2018.genereg.net/matrix/MA0913.1 |
| 237 | db/JASPAR/J | MA0063.1 | Nkx2-5 | WTAAKTG | 7 | 10 | 1.9e-002 | CENTRIMO | http://jaspar2018.genereg.net/matrix/MA0063.1 |
| 238 | db/MOUSE/ur | UP00207_1 | Hoxb9_3413. | VRRGCMATA | 16 | 10 | 1.9e-002 | CENTRIMO | http://the_brain.bwh.harvard.edu/uniprobe/details2.php?id=00207 |
| 239 | db/EUKARYO | HMX1_DBD | | ANCAATTAA | 11 | 10 | 1.9e-002 | CENTRIMO | http://floresta.eead.csic.es/footprintdb/index.php?db=HumanTF:1.0&motif=HMX1_DBD |
| 240 | db/EUKARYO | HMX2_DBD | | AMCAMTTAA | 11 | 10 | 1.9e-002 | CENTRIMO | http://floresta.eead.csic.es/footprintdb/index.php?db=HumanTF:1.0&motif=HMX2_DBD |
| 241 | db/EUKARYO | HMX3_DBD | | RVCAMTTAA | 11 | 10 | 1.9e-002 | CENTRIMO | http://floresta.eead.csic.es/footprintdb/index.php?db=HumanTF:1.0&motif=HMX3_DBD |
| 242 | db/EUKARYO | HOXA10_DBD_2 | | DGYMATAAA | 11 | 10 | 1.9e-002 | CENTRIMO | http://floresta.eead.csic.es/footprintdb/index.php?db=HumanTF:1.0&motif=HOXA10_DBD_2 |
| 243 | db/EUKARYO | HOXC11_DBD_2 | | DGYMATAAA | 11 | 10 | 1.9e-002 | CENTRIMO | http://floresta.eead.csic.es/footprintdb/index.php?db=HumanTF:1.0&motif=HOXC11_DBD_2 |
| 244 | db/EUKARYO | HOXC11_full_2 | | RGYMATAAA | 11 | 10 | 1.9e-002 | CENTRIMO | http://floresta.eead.csic.es/footprintdb/index.php?db=HumanTF:1.0&motif=HOXC11_full_2 |
| 245 | db/JASPAR/J | MA0899.1 | HOXA10 | DGYMATAAA | 11 | 10 | 1.9e-002 | CENTRIMO | http://jaspar2018.genereg.net/matrix/MA0899.1 |
| 246 | db/EUKARYO | BARX1_DBD_2 | | VCMATTAV | 8 | 10 | 1.9e-002 | CENTRIMO | http://floresta.eead.csic.es/footprintdb/index.php?db=HumanTF:1.0&motif=BARX1_DBD_2 |
| 247 | db/EUKARYO | LHX9_DBD_1 | | BYAATTAR | 8 | 10 | 1.9e-002 | CENTRIMO | http://floresta.eead.csic.es/footprintdb/index.php?db=HumanTF:1.0&motif=LHX9_DBD_1 |
| 248 | db/EUKARYO | LMX1B_DBD | | NYAATTAA | 8 | 10 | 1.9e-002 | CENTRIMO | http://floresta.eead.csic.es/footprintdb/index.php?db=HumanTF:1.0&motif=LMX1B_DBD |
| 249 | db/EUKARYO | MSX1_DBD_2 | | SCAATTAV | 8 | 10 | 1.9e-002 | CENTRIMO | http://floresta.eead.csic.es/footprintdb/index.php?db=HumanTF:1.0&motif=MSX1_DBD_2 |
| 250 | db/EUKARYO | MSX2_DBD_2 | | CCAATTAV | 8 | 10 | 1.9e-002 | CENTRIMO | http://floresta.eead.csic.es/footprintdb/index.php?db=HumanTF:1.0&motif=MSX2_DBD_2 |
| 251 | db/EUKARYO | Msx3_DBD_2 | | SCAATTAN | 8 | 10 | 1.9e-002 | CENTRIMO | http://floresta.eead.csic.es/footprintdb/index.php?db=HumanTF:1.0&motif=Msx3_DBD_2 |
| 252 | db/EUKARYO | PRRX1_DBD | | YYAATTAR | 8 | 10 | 1.9e-002 | CENTRIMO | http://floresta.eead.csic.es/footprintdb/index.php?db=HumanTF:1.0&motif=PRRX1_DBD |
| 253 | db/EUKARYO | SHOX2_DBD | | YYAATTAR | 8 | 10 | 1.9e-002 | CENTRIMO | http://floresta.eead.csic.es/footprintdb/index.php?db=HumanTF:1.0&motif=SHOX2_DBD |
| 254 | db/EUKARYO | SHOX_DBD | | YYAATTAR | 8 | 10 | 1.9e-002 | CENTRIMO | http://floresta.eead.csic.es/footprintdb/index.php?db=HumanTF:1.0&motif=SHOX_DBD |
| 255 | db/JASPAR/J | MA0701.1 | LHX9 | BYAATTAR | 8 | 10 | 1.9e-002 | CENTRIMO | http://jaspar2018.genereg.net/matrix/MA0701.1 |
| 256 | db/JASPAR/J | MA0703.1 | LMX1B | NYAATTAA | 8 | 10 | 1.9e-002 | CENTRIMO | http://jaspar2018.genereg.net/matrix/MA0703.1 |
| 257 | db/JASPAR/J | MA0708.1 | MSX2 | CCAATTAV | 8 | 10 | 1.9e-002 | CENTRIMO | http://jaspar2018.genereg.net/matrix/MA0708.1 |
| 258 | db/JASPAR/J | MA0709.1 | Msx3 | SCAATTAN | 8 | 10 | 1.9e-002 | CENTRIMO | http://jaspar2018.genereg.net/matrix/MA0709.1 |
| 259 | db/JASPAR/J | MA0875.1 | BARX1 | VCMATTAV | 8 | 10 | 1.9e-002 | CENTRIMO | http://jaspar2018.genereg.net/matrix/MA0875.1 |
| 260 | db/JASPAR/J | MA0896.1 | Hmx1 | VSVAGCAAT | 17 | 10 | 1.9e-002 | CENTRIMO | http://jaspar2018.genereg.net/matrix/MA0896.1 |
| 261 | db/MOUSE/ur | UP00078_1 | Arid3a_primar | SNNHTTAAT | 17 | 10 | 1.9e-002 | CENTRIMO | http://the_brain.bwh.harvard.edu/uniprobe/details2.php?id=00078 |
| 262 | db/MOUSE/ur | UP00104_1 | Hmx1_3423.1 | VSVAGCAAT | 17 | 10 | 1.9e-002 | CENTRIMO | http://the_brain.bwh.harvard.edu/uniprobe/details2.php?id=00104 |
| 263 | db/EUKARYO | HOXD11_DBD_2 | | RRYMATAAA | 10 | 10 | 1.9e-002 | CENTRIMO | http://floresta.eead.csic.es/footprintdb/index.php?db=HumanTF:1.0&motif=HOXD11_DBD_2 |
| 264 | db/EUKARYO | LBX2_DBD_2 | | NCYAATTAR | 10 | 10 | 1.9e-002 | CENTRIMO | http://floresta.eead.csic.es/footprintdb/index.php?db=HumanTF:1.0&motif=LBX2_DBD_2 |
| 265 | db/EUKARYO | MIXL1_full | | NBYAATTAVN | 10 | 10 | 1.9e-002 | CENTRIMO | http://floresta.eead.csic.es/footprintdb/index.php?db=HumanTF:1.0&motif=MIXL1_full |
| 266 | db/EUKARYO | MNX1_DBD | | DNYAATTAA | 10 | 10 | 1.9e-002 | CENTRIMO | http://floresta.eead.csic.es/footprintdb/index.php?db=HumanTF:1.0&motif=MNX1_DBD |
| 267 | db/EUKARYO | RAX_DBD | | RYYAATTAR | 10 | 10 | 1.9e-002 | CENTRIMO | http://floresta.eead.csic.es/footprintdb/index.php?db=HumanTF:1.0&motif=RAX_DBD |
| 268 | db/JASPAR/J | MA0662.1 | MIXL1 | NBYAATTAVN | 10 | 10 | 1.9e-002 | CENTRIMO | http://jaspar2018.genereg.net/matrix/MA0662.1 |
| 269 | db/JASPAR/J | MA0699.1 | LBX2 | NCYAATTAR | 10 | 10 | 1.9e-002 | CENTRIMO | http://jaspar2018.genereg.net/matrix/MA0699.1 |
| 270 | db/JASPAR/J | MA0707.1 | MNX1 | DNYAATTAA | 10 | 10 | 1.9e-002 | CENTRIMO | http://jaspar2018.genereg.net/matrix/MA0707.1 |
| 271 | db/JASPAR/J | MA0718.1 | RAX | RYYAATTAR | 10 | 10 | 1.9e-002 | CENTRIMO | http://jaspar2018.genereg.net/matrix/MA0718.1 |
| 272 | db/MOUSE/ur | UP00120_1 | Lbx2_3869.2 | TVHHYTAAT | 17 | 10 | 2.0e-002 | CENTRIMO | http://the_brain.bwh.harvard.edu/uniprobe/details2.php?id=00120 |
| 273 | db/MOUSE/ur | UP00175_1 | Lhx9_3492.1 | CBYATTAATT | 17 | 10 | 2.0e-002 | CENTRIMO | http://the_brain.bwh.harvard.edu/uniprobe/details2.php?id=00175 |
| 274 | db/MOUSE/ur | UP00139_1 | Nkx1-2_3214. | NKBCRYTAA | 17 | 10 | 2.0e-002 | CENTRIMO | http://the_brain.bwh.harvard.edu/uniprobe/details2.php?id=00139 |
| 275 | db/MOUSE/ur | UP00253_1 | Rax_3443.1 | WSCAYTAAT | 17 | 10 | 2.0e-002 | CENTRIMO | http://the_brain.bwh.harvard.edu/uniprobe/details2.php?id=00253 |
| 276 | db/MOUSE/ur | UP00023_1 | Sox30_primar | DHDSAACAA | 16 | 10 | 2.0e-002 | CENTRIMO | http://the_brain.bwh.harvard.edu/uniprobe/details2.php?id=00023 |
| 277 | db/EUKARYO | YY2_DBD | | RWCCGCCA | 11 | 10 | 2.3e-002 | CENTRIMO | http://floresta.eead.csic.es/footprintdb/index.php?db=HumanTF:1.0&motif=YY2_DBD |
| 278 | db/JASPAR/J | MA0748.1 | YY2 | RWCCGCCA | 11 | 10 | 2.3e-002 | CENTRIMO | http://jaspar2018.genereg.net/matrix/MA0748.1 |
| 279 | db/JASPAR/J | MA0057.1 | MZF1(var.2) | BKAGGGGK | 10 | 10 | 2.3e-002 | CENTRIMO | http://jaspar2018.genereg.net/matrix/MA0057.1 |
| 280 | db/EUKARYO | GLIS3_DBD | | GACCCCCC | 14 | 10 | 2.3e-002 | CENTRIMO | http://floresta.eead.csic.es/footprintdb/index.php?db=HumanTF:1.0&motif=GLIS3_DBD |
| 281 | db/JASPAR/J | MA0737.1 | GLIS3 | GACCCCCC | 14 | 10 | 2.3e-002 | CENTRIMO | http://jaspar2018.genereg.net/matrix/MA0737.1 |
| 282 | db/MOUSE/ur | UP00390_1 | Tcf1_2666.2 | YSKNRGTTA | 17 | 10 | 2.3e-002 | CENTRIMO | http://the_brain.bwh.harvard.edu/uniprobe/details2.php?id=00390 |
| 283 | db/MOUSE/ur | UP00113_1 | Hoxc4_3491.1 | CRMRTTAAT | 17 | 10 | 2.4e-002 | CENTRIMO | http://the_brain.bwh.harvard.edu/uniprobe/details2.php?id=00113 |
| 284 | db/MOUSE/ur | UP00222_1 | Tcf2_0913.2 | DBYNGTTAA | 17 | 10 | 2.4e-002 | CENTRIMO | http://the_brain.bwh.harvard.edu/uniprobe/details2.php?id=00222 |
| 285 | db/EUKARYO | HMBOX1_DBD | | MYTAGTTAM | 10 | 10 | 2.4e-002 | CENTRIMO | http://floresta.eead.csic.es/footprintdb/index.php?db=HumanTF:1.0&motif=HMBOX1_DBD |
| 286 | db/JASPAR/J | MA0895.1 | HMBOX1 | MYTAGTTAM | 10 | 10 | 2.4e-002 | CENTRIMO | http://jaspar2018.genereg.net/matrix/MA0895.1 |
| 287 | db/EUKARYO | Hoxd9_DBD_1 | | SCCATWAAA | 9 | 10 | 2.4e-002 | CENTRIMO | http://floresta.eead.csic.es/footprintdb/index.php?db=HumanTF:1.0&motif=Hoxd9_DBD_1 |
| 288 | db/MOUSE/ur | UP00161_1 | Hmbox1_267 | NHRAWCTA | 17 | 10 | 2.4e-002 | CENTRIMO | http://the_brain.bwh.harvard.edu/uniprobe/details2.php?id=00161 |
| 289 | DREME | GARGAAAA | DREME-18 | GARGAAAA | 8 | 64 | 2.5e-002 | DREME | |

| | | | | | | | | | |
|---|---|---|---|---|---|---|---|---|---|
| 290 | db/JASPAR/JA | MA0164.1 | Nr2e3 | CAAGCTT | 7 | 10 | 2.7e-002 | CENTRIMO | http://jaspar2018.genereg.net/matrix/MA0164.1 |
| 291 | db/MOUSE/ur | UP00068_1 | Eomes_prima | NNNRAGGTG | 17 | 10 | 2.9e-002 | CENTRIMO | http://the_brain.bwh.harvard.edu/uniprobe/details2.php?id=00068 |
| 292 | db/JASPAR/JA | MA0102.3 | CEBPA | ATTGCAYAAY | 11 | 10 | 3.0e-002 | CENTRIMO | http://jaspar2018.genereg.net/matrix/MA0102.3 |
| 293 | db/EUKARYO | CEBPB_DBD | | RTTRCGCAA | 10 | 10 | 3.0e-002 | CENTRIMO | http://floresta.eead.csic.es/footprintdb/index.php?db=HumanTF:1.0&motif=CEBPB_DBD |
| 294 | db/EUKARYO | CEBPD_DBD | | RTTRCGCAA | 10 | 10 | 3.0e-002 | CENTRIMO | http://floresta.eead.csic.es/footprintdb/index.php?db=HumanTF:1.0&motif=CEBPD_DBD |
| 295 | db/EUKARYO | CEBPE_DBD | | VTTRCGCAA | 10 | 10 | 3.0e-002 | CENTRIMO | http://floresta.eead.csic.es/footprintdb/index.php?db=HumanTF:1.0&motif=CEBPE_DBD |
| 296 | db/EUKARYO | Cebpb_DBD | | RTTRCGCAA | 10 | 10 | 3.0e-002 | CENTRIMO | http://floresta.eead.csic.es/footprintdb/index.php?db=HumanTF:1.0&motif=Cebpb_DBD |
| 297 | db/JASPAR/JA | MA0466.2 | CEBPB | RTTRCGCAA | 10 | 10 | 3.0e-002 | CENTRIMO | http://jaspar2018.genereg.net/matrix/MA0466.2 |
| 298 | db/JASPAR/JA | MA0836.1 | CEBPD | RTTRCGCAA | 10 | 10 | 3.0e-002 | CENTRIMO | http://jaspar2018.genereg.net/matrix/MA0836.1 |
| 299 | db/JASPAR/JA | MA0837.1 | CEBPE | VTTRCGCAA | 10 | 10 | 3.0e-002 | CENTRIMO | http://jaspar2018.genereg.net/matrix/MA0837.1 |
| 300 | db/EUKARYO | MAFF_DBD | | WTGCTGAST | 15 | 10 | 3.0e-002 | CENTRIMO | http://floresta.eead.csic.es/footprintdb/index.php?db=HumanTF:1.0&motif=MAFF_DBD |
| 301 | db/EUKARYO | MAFK_full_2 | | DTGCTGAST | 15 | 10 | 3.0e-002 | CENTRIMO | http://floresta.eead.csic.es/footprintdb/index.php?db=HumanTF:1.0&motif=MAFK_full_2 |
| 302 | db/EUKARYO | Mafb_DBD_2 | | WNTGCTGAS | 17 | 10 | 3.1e-002 | CENTRIMO | http://floresta.eead.csic.es/footprintdb/index.php?db=HumanTF:1.0&motif=Mafb_DBD_2 |
| 303 | db/EUKARYO | MAFG_full | | WWWNTGCT | 21 | 10 | 3.1e-002 | CENTRIMO | http://floresta.eead.csic.es/footprintdb/index.php?db=HumanTF:1.0&motif=MAFG_full |
| 304 | db/EUKARYO | MAFK_DBD_2 | | DWWNTGCT | 21 | 10 | 3.1e-002 | CENTRIMO | http://floresta.eead.csic.es/footprintdb/index.php?db=HumanTF:1.0&motif=MAFK_DBD_2 |
| 305 | db/JASPAR/JA | MA0659.1 | MAFG | WWWNTGCT | 21 | 10 | 3.1e-002 | CENTRIMO | http://jaspar2018.genereg.net/matrix/MA0659.1 |
| 306 | db/MOUSE/ur | UP00018_1 | Irf4_primary | MRWAYCGAA | 15 | 10 | 3.1e-002 | CENTRIMO | http://the_brain.bwh.harvard.edu/uniprobe/details2.php?id=00018 |
| 307 | db/EUKARYO | IRF5_full_1 | | CCGAAACCG | 14 | 10 | 3.4e-002 | CENTRIMO | http://floresta.eead.csic.es/footprintdb/index.php?db=HumanTF:1.0&motif=IRF5_full_1 |
| 308 | db/EUKARYO | IRF8_DBD | | HCGAAACCG | 14 | 10 | 3.4e-002 | CENTRIMO | http://floresta.eead.csic.es/footprintdb/index.php?db=HumanTF:1.0&motif=IRF8_DBD |
| 309 | db/JASPAR/JA | MA1420.1 | IRF5 | CCGAAACCG | 14 | 10 | 3.4e-002 | CENTRIMO | http://jaspar2018.genereg.net/matrix/MA1420.1 |
| 310 | db/EUKARYO | POU3F1_DBD_1 | | WTATGCWAA | 12 | 10 | 3.8e-002 | CENTRIMO | http://floresta.eead.csic.es/footprintdb/index.php?db=HumanTF:1.0&motif=POU3F1_DBD_1 |
| 311 | db/JASPAR/JA | MA0786.1 | POU3F1 | WTATGCWAA | 12 | 10 | 3.8e-002 | CENTRIMO | http://jaspar2018.genereg.net/matrix/MA0786.1 |
| 312 | db/JASPAR/JA | MA0627.1 | Pou2f3 | THKTATGCAA | 16 | 10 | 3.8e-002 | CENTRIMO | http://jaspar2018.genereg.net/matrix/MA0627.1 |
| 313 | db/MOUSE/ur | UP00179_1 | Pou2f3_3986. | THKTATGCAA | 16 | 10 | 3.8e-002 | CENTRIMO | http://the_brain.bwh.harvard.edu/uniprobe/details2.php?id=00179 |
| 314 | db/EUKARYO | POU2F3_DBD_1 | | TATGCWAAT | 9 | 10 | 3.8e-002 | CENTRIMO | http://floresta.eead.csic.es/footprintdb/index.php?db=HumanTF:1.0&motif=POU2F3_DBD_1 |
| 315 | db/EUKARYO | POU3F4_DBD_1 | | TATGCWAAT | 9 | 10 | 3.8e-002 | CENTRIMO | http://floresta.eead.csic.es/footprintdb/index.php?db=HumanTF:1.0&motif=POU3F4_DBD_1 |
| 316 | db/EUKARYO | POU5F1P1_DBD_1 | | TATGCWAAT | 9 | 10 | 3.8e-002 | CENTRIMO | http://floresta.eead.csic.es/footprintdb/index.php?db=HumanTF:1.0&motif=POU5F1P1_DBD_1 |
| 317 | db/JASPAR/JA | MA0789.1 | POU3F4 | TATGCWAAT | 9 | 10 | 3.8e-002 | CENTRIMO | http://jaspar2018.genereg.net/matrix/MA0789.1 |
| 318 | db/JASPAR/JA | MA0792.1 | POU5F1B | TATGCWAAT | 9 | 10 | 3.8e-002 | CENTRIMO | http://jaspar2018.genereg.net/matrix/MA0792.1 |
| 319 | db/EUKARYO | POU2F2_DBD_1 | | DTATGCWAA | 11 | 10 | 3.8e-002 | CENTRIMO | http://floresta.eead.csic.es/footprintdb/index.php?db=HumanTF:1.0&motif=POU2F2_DBD_1 |
| 320 | db/JASPAR/JA | MA1115.1 | POU5F1 | WHATGCAAA | 11 | 10 | 3.8e-002 | CENTRIMO | http://jaspar2018.genereg.net/matrix/MA1115.1 |
| 321 | db/EUKARYO | TBX2_full_1 | | GGTGTGARA | 18 | 10 | 3.9e-002 | CENTRIMO | http://floresta.eead.csic.es/footprintdb/index.php?db=HumanTF:1.0&motif=TBX2_full_1 |
| 322 | db/EUKARYO | TBX4_DBD_2 | | AGGTGTGAD | 20 | 10 | 3.9e-002 | CENTRIMO | http://floresta.eead.csic.es/footprintdb/index.php?db=HumanTF:1.0&motif=TBX4_DBD_2 |
| 323 | db/JASPAR/JA | MA0031.1 | FOXD1 | GTAAACAW | 8 | 10 | 4.0e-002 | CENTRIMO | http://jaspar2018.genereg.net/matrix/MA0031.1 |
| 324 | db/EUKARYO | EOMES_DBD_1 | | RAGGTGTGA | 13 | 10 | 4.3e-002 | CENTRIMO | http://floresta.eead.csic.es/footprintdb/index.php?db=HumanTF:1.0&motif=EOMES_DBD_1 |
| 325 | db/JASPAR/JA | MA0800.1 | EOMES | RAGGTGTGA | 13 | 10 | 4.3e-002 | CENTRIMO | http://jaspar2018.genereg.net/matrix/MA0800.1 |
| 326 | db/EUKARYO | TBR1_DBD | | AGGTGTGAA | 10 | 10 | 4.3e-002 | CENTRIMO | http://floresta.eead.csic.es/footprintdb/index.php?db=HumanTF:1.0&motif=TBR1_DBD |
| 327 | db/JASPAR/JA | MA0802.1 | TBR1 | AGGTGTGAA | 10 | 10 | 4.3e-002 | CENTRIMO | http://jaspar2018.genereg.net/matrix/MA0802.1 |
| 328 | db/EUKARYO | TBR1_full | | AAGGTGTGA | 11 | 10 | 4.4e-002 | CENTRIMO | http://floresta.eead.csic.es/footprintdb/index.php?db=HumanTF:1.0&motif=TBR1_full |
| 329 | db/EUKARYO | TBX20_full_1 | | WAGGTGTGA | 11 | 10 | 4.4e-002 | CENTRIMO | http://floresta.eead.csic.es/footprintdb/index.php?db=HumanTF:1.0&motif=TBX20_full_1 |
| 330 | db/EUKARYO | TBX2_full_2 | | DAGGTGTGA | 11 | 10 | 4.4e-002 | CENTRIMO | http://floresta.eead.csic.es/footprintdb/index.php?db=HumanTF:1.0&motif=TBX2_full_2 |
| 331 | db/JASPAR/JA | MA0688.1 | TBX2 | DAGGTGTGA | 11 | 10 | 4.4e-002 | CENTRIMO | http://jaspar2018.genereg.net/matrix/MA0688.1 |
| 332 | db/JASPAR/JA | MA0689.1 | TBX20 | WAGGTGTGA | 11 | 10 | 4.4e-002 | CENTRIMO | http://jaspar2018.genereg.net/matrix/MA0689.1 |
| 333 | db/EUKARYO | MGA_DBD_1 | | AGGTGTGA | 8 | 10 | 4.4e-002 | CENTRIMO | http://floresta.eead.csic.es/footprintdb/index.php?db=HumanTF:1.0&motif=MGA_DBD_1 |
| 334 | db/EUKARYO | TBX15_DBD_2 | | AGGTGTGA | 8 | 10 | 4.4e-002 | CENTRIMO | http://floresta.eead.csic.es/footprintdb/index.php?db=HumanTF:1.0&motif=TBX15_DBD_2 |
| 335 | db/EUKARYO | TBX1_DBD_3 | | AGGTGTGA | 8 | 10 | 4.4e-002 | CENTRIMO | http://floresta.eead.csic.es/footprintdb/index.php?db=HumanTF:1.0&motif=TBX1_DBD_3 |
| 336 | db/EUKARYO | TBX4_DBD_1 | | AGGTGTGA | 8 | 10 | 4.4e-002 | CENTRIMO | http://floresta.eead.csic.es/footprintdb/index.php?db=HumanTF:1.0&motif=TBX4_DBD_1 |
| 337 | db/EUKARYO | TBX5_DBD_1 | | AGGTGTKA | 8 | 10 | 4.4e-002 | CENTRIMO | http://floresta.eead.csic.es/footprintdb/index.php?db=HumanTF:1.0&motif=TBX5_DBD_1 |
| 338 | db/JASPAR/JA | MA0801.1 | MGA | AGGTGTGA | 8 | 10 | 4.4e-002 | CENTRIMO | http://jaspar2018.genereg.net/matrix/MA0801.1 |
| 339 | db/JASPAR/JA | MA0803.1 | TBX15 | AGGTGTGA | 8 | 10 | 4.4e-002 | CENTRIMO | http://jaspar2018.genereg.net/matrix/MA0803.1 |
| 340 | db/JASPAR/JA | MA0805.1 | TBX1 | AGGTGTGA | 8 | 10 | 4.4e-002 | CENTRIMO | http://jaspar2018.genereg.net/matrix/MA0805.1 |
| 341 | db/JASPAR/JA | MA0806.1 | TBX4 | AGGTGTGA | 8 | 10 | 4.4e-002 | CENTRIMO | http://jaspar2018.genereg.net/matrix/MA0806.1 |
| 342 | db/JASPAR/JA | MA0807.1 | TBX5 | AGGTGTKA | 8 | 10 | 4.4e-002 | CENTRIMO | http://jaspar2018.genereg.net/matrix/MA0807.1 |
| 343 | db/EUKARYO | TBX21_DBD_2 | | DAGGTGTGA | 10 | 10 | 4.4e-002 | CENTRIMO | http://floresta.eead.csic.es/footprintdb/index.php?db=HumanTF:1.0&motif=TBX21_DBD_2 |
| 344 | db/EUKARYO | TBX21_full_2 | | AAGGTGTGA | 10 | 10 | 4.4e-002 | CENTRIMO | http://floresta.eead.csic.es/footprintdb/index.php?db=HumanTF:1.0&motif=TBX21_full_2 |
| 345 | db/JASPAR/JA | MA0690.1 | TBX21 | AAGGTGTGA | 10 | 10 | 4.4e-002 | CENTRIMO | http://jaspar2018.genereg.net/matrix/MA0690.1 |
| 346 | db/EUKARYO | TBX20_DBD_3 | | GNDAAGGTG | 15 | 10 | 4.5e-002 | CENTRIMO | http://floresta.eead.csic.es/footprintdb/index.php?db=HumanTF:1.0&motif=TBX20_DBD_3 |

### MEME-ChIP (Motif Analysis of Large Nucleotide Datasets): Version 5.0.3 released on Sun Dec 02 18:41:45 2018 -0800
### The format of this file is described at http://meme-suite.org//doc/meme-chip-output-format.html
### meme-chip -oc . -time 300 -ccut 100 -order 1 -db db/EUKARYOTE/jolma2013.meme -db db/JASPAR/JASPAR2018_CORE_vertebrates_non-redundant.meme -db db/MOUSE/uniprobe_mouse.meme -meme-mod zoops -meme-minw 6 -meme-maxw 30 -meme-nmotifs 3 -meme-searchsize 1000

**ZT18 dynamic**

| MOTIF_INDEX | MOTIF_SOUF | MOTIF_ID | ALT_ID | CONSENSUS | WIDTH | SITES | E-VALUE | E-VALUE_SO | MOST_SIMILAR_MC | **MOST_SIMILAR_MO** | URL |
|---|---|---|---|---|---|---|---|---|---|---|---|
| 1 | MEME | TWTDTTTKT | MEME-3 | TWTDTTTKT | 30 | 233 | 1.1e-190 | MEME | db/JASPAR/JASPAR | **MA1125.1 (ZNF384)** | http://jaspar2018.genereg.net/matrix/MA1125.1 |
| 2 | MEME | TGCCTCTGC | MEME-1 | TGCCTCTGC | 26 | 37 | 1.9e-186 | MEME | db/MOUSE/uniprobe | **UP00087_2 (Tcfap2c_** | http://the_brain.bwh.harvard.edu/uniprobe/details2.php?id=00087 |
| 3 | MEME | AGGCCAGC | MEME-2 | AGGCCAGC | 21 | 54 | 1.9e-181 | MEME | db/JASPAR/JASPAR | **MA0258.2 (ESR2)** | http://jaspar2018.genereg.net/matrix/MA0258.2 |
| 4 | DREME | GAGTTCNA | DREME-1 | GAGTTCNA | 8 | 78 | 2.2e-007 | DREME | db/JASPAR/JASPAR | **MA0505.1 (Nr5a2)** | http://jaspar2018.genereg.net/matrix/MA0505.1 |
| 5 | DREME | CTCTGB | DREME-2 | CTCTGB | 6 | 409 | 5.4e-006 | DREME | | | |
| 6 | DREME | GCTGGGAW | DREME-3 | GCTGGGAW | 8 | 48 | 3.5e-005 | DREME | | | |
| 7 | DREME | RTGGCTCA | DREME-5 | RTGGCTCA | 8 | 41 | 7.1e-004 | DREME | | | |
| 8 | DREME | GTGTGTS | DREME-4 | GTGTGTS | 7 | 120 | 2.5e-003 | DREME | db/MOUSE/uniprobe | **UP00042_2 (Gm397_** | http://the_brain.bwh.harvard.edu/uniprobe/details2.php?id=00042 |
| 9 | DREME | AAAWAAAA | DREME-6 | AAAWAAAA | 8 | 107 | 1.8e-002 | DREME | db/JASPAR/JASPAR | **MA1125.1 (ZNF384)** | http://jaspar2018.genereg.net/matrix/MA1125.1 |
| 10 | DREME | AAGAAGMA | DREME-7 | AAGAAGMA | 8 | 34 | 3.7e-002 | DREME | | | |

### MEME-ChIP (Motif Analysis of Large Nucleotide Datasets): Version 5.0.3 released on Sun Dec 02 18:41:45 2018 -0800
### The format of this file is described at http://meme-suite.org//doc/meme-chip-output-format.html
### meme-chip -oc . -time 300 -ccut 100 -order 1 -db db/EUKARYOTE/jolma2013.meme -db db/JASPAR/JASPAR2018_CORE_vertebrates_non-redundant.meme -db db/MOUSE/uniprobe_mouse.meme -meme-mod zoops -meme-minw 6 -meme-maxw 30 -meme-nmotifs 3 -meme-searchsize 1000

**ZT18 static**

| MOTIF_INDEX | MOTIF_SOUF | MOTIF_ID | ALT_ID | CONSENSUS | WIDTH | SITES | E-VALUE | E-VALUE_SO | MOST_SIMILAR_MC | **MOST_SIMILAR_MO** | URL |
|---|---|---|---|---|---|---|---|---|---|---|---|
| 1 | MEME | GSCKGGAG | MEME-1 | GSCKGGAG | 29 | 101 | 1.5e-182 | MEME | db/JASPAR/JASPAR | **MA0471.1 (E2F6)** | http://jaspar2018.genereg.net/matrix/MA0471.1 |
| 2 | MEME | YGAGTTCGA | MEME-2 | YGAGTTCGA | 29 | 54 | 1.0e-163 | MEME | db/JASPAR/JASPAR | **MA0505.1 (Nr5a2)** | http://jaspar2018.genereg.net/matrix/MA0505.1 |
| 3 | MEME | TRCRCMKKC | MEME-3 | TRCRCMKKC | 30 | 18 | 7.2e-115 | MEME | | | |
| 4 | DREME | RTGGCTCA | DREME-1 | RTGGCTCA | 8 | 56 | 1.0e-007 | DREME | db/JASPAR/JASPAR | **MA0099.3 (FOS::JUN** | http://jaspar2018.genereg.net/matrix/MA0099.3 |
| 5 | DREME | TAHATA | DREME-2 | TAHATA | 6 | 191 | 4.5e-006 | DREME | | | |
| 6 | DREME | ACHCACA | DREME-4 | ACHCACA | 7 | 148 | 7.4e-005 | DREME | | | |
| 7 | DREME | GGCTGK | DREME-3 | GGCTGK | 6 | 220 | 1.6e-004 | DREME | | | |
| 8 | DREME | GAGGWTC | DREME-5 | GAGGWTC | 7 | 52 | 1.0e-003 | DREME | | | |
| 9 | DREME | GAACTCR | DREME-6 | GAACTCR | 7 | 68 | 1.9e-003 | DREME | db/JASPAR/JASPAR | **MA0693.2 (VDR)** | http://jaspar2018.genereg.net/matrix/MA0693.2 |
| 10 | DREME | GCAARCA | DREME-7 | GCAARCA | 7 | 61 | 5.6e-003 | DREME | | | |
| 11 | DREME | CCGGGASG | DREME-8 | CCGGGASG | 8 | 22 | 7.0e-003 | DREME | db/MOUSE/uniprobe | **UP00022_2 (Zfp740_** | http://the_brain.bwh.harvard.edu/uniprobe/details2.php?id=00022 |
| 12 | DREME | CGGCATCK | DREME-9 | CGGCATCK | 8 | 22 | 7.0e-003 | DREME | | | |
| 13 | DREME | GGAAGMA | DREME-10 | GGAAGMA | 7 | 85 | 1.0e-002 | DREME | | | |
| 14 | DREME | KCTGGGA | DREME-11 | KCTGGGA | 7 | 79 | 3.8e-002 | DREME | db/JASPAR/JASPAR | **MA0144.2 (STAT3)** | http://jaspar2018.genereg.net/matrix/MA0144.2 |
| 15 | DREME | ATAAWTC | DREME-12 | ATAAWTC | 7 | 42 | 4.2e-002 | DREME | | | |

### MEME-ChIP (Motif Analysis of Large Nucleotide Datasets): Version 5.0.3 released on Sun Dec 02 18:41:45 2018 -0800
### The format of this file is described at http://meme-suite.org//doc/meme-chip-output-format.html
### meme-chip -oc . -time 300 -ccut 100 -order 1 -db db/EUKARYOTE/jolma2013.meme -db db/JASPAR/JASPAR2018_CORE_vertebrates_non-redundant.meme -db db/MOUSE/uniprobe_mouse.meme -meme-mod zoops -meme-minw 6 -meme-maxw 30 -meme-nmotifs 3 -meme-searchsize 1000
